# *Trypanosoma cruzi* has 32 Chromosomes: A Telomere-to-Telomere Assembly Defines Its Karyotype

**DOI:** 10.1101/2025.03.27.645724

**Authors:** G. Greif, M.L. Chiribao, F. Díaz-Viraqué, Carlos E. Sanz-Rodríguez, C. Robello

## Abstract

*Trypanosoma cruzi*, the causative agent of Chagas disease, exhibits remarkable genomic variability and possesses an expanded genome rich in multigene families. However, its precise chromosomal composition has remained elusive due to the challenges posed by extensive repetitive regions. Using PacBio HiFi long-read sequencing, we assembled the complete nuclear genome of the Dm28c strain into 32 telomere-to-telomere chromosomes. This assembly revealed conserved chromosomal structures and synteny patterns when compared to the independently sequenced Dm25 strain, indicating that the 32-chromosome karyotype is a stable and conserved feature of *T. cruzi*. The species is diploid for all chromosomes, except for chromosome 16, which is consistently tetraploid in all strains analyzed, and homologous to chromosome 31 of Leishmania, also tetraploid. Through a comprehensive annotation pipeline, we refined gene content and resolved haplotypes, resulting in an approximately 25% reduction in core gene redundancy. All chromosomes display a conserved, compartmentalized architecture comprising core, disruptive, and newly defined subtelomeric regions. Subtelomeres, enriched in RHS, TS, and DGF-1 genes and depleted in MASP and mucins, constitute a distinct third genomic compartment, that is transcriptionally active, and represents the main source of genome variability among strains.

Altogether, this work defines the complete chromosomal complement of *T. cruzi*, establishing a robust framework for comparative genomics and enabling detailed studies into genome organization, antigenic variability, and evolutionary dynamics across strains and clades. This assembly redefines the genomic reference landscape and opens new avenues for exploring *T. cruzi* biology, pathogenic diversity, and adaptive mechanisms.

## Introduction

*Trypanosoma cruzi,* the causative agent of Chagas disease, belongs to the family Trypanosomatidae, which is composed exclusively of parasitic organisms. Three of these, known as the “TriTryps”*—Leishmania spp.*, *Trypanosoma brucei*, and *T. cruzi*—pose significant challenges to human and animal health and have been extensively studied. They share several biological features, including RNA editing, polycistronic transcription, a single mitochondrion with kinetoplast DNA (kDNA), *trans*-splicing, and the absence of chromosomal condensation during mitosis. Additionally, each of these species possesses unique characteristics. In *T. cruzi*, the most notable is its genome expansion due to multigene families, the majority of which encode surface GPI-anchored proteins. These include trans-sialidases (TS; Schenkman et al., 1991; Cazzulo and Frasch, 1992; Buscaglia et al., 1999; Freitas et al., 2011; Rubin and Schenkman, 2012; Freire-de-Lima et al., 2015), mucins (Frasch, 2000; Pollevick et al., 2000; Buscaglia et al., 2004; Buscaglia et al., 2006), mucin associated surface proteins (MASP; De Pablos and Osuna, 2012; Leão et al., 2022; Dean et al., 2025), GP63 (Alvarez et al., 2012; Berná et al., 2025), disperse gene family 1 (DGF-1; Wincker et al., 1992; Ramirez, 2023), and retroposon hot spot proteins (RHS; Bringaud et al., 2002; Bernardo et al., 2020). This expansion in *T. cruzi* represents an evolutionary acquisition linked to antigenic variability, invasion, infectivity, and immune system evasion, among other factors.

Regarding the karyotype of *T. cruzi*, the absence of chromosomal condensation during mitosis has impeded the use of classical cytogenetic studies. Instead, pulsed-field gel electrophoresis (PFGE) emerged decades ago as a valuable tool for determining their karyotypes. Using this approach, it was established that *T. brucei* has 11 chromosomes (Melville et al., 1998; Berriman et al., 2005), *Leishmania major*, *L. infantum*, and *L. donovani* each have 36 (Wincker et al., 1996), and *L. mexicana* and *L. braziliensis* contain 34 and 35 chromosomes, respectively (Britto et al., 1998). However, the exact chromosome number of *T. cruzi* remains unknown. PFGE studies consistently indicate that chromosomal sizes vary among different strains, ranging from 0.5 to 3 Mb, and estimate that it may have between 20 and 40 chromosomes (Henriksson et al., 1990, 1995, 1996; Santos et al., 1997; Vargas et al., 2004; Galindo et al., 2007; Souza et al., 2011). The challenge in determining its precise number, compared to *Leishmania* and *T. brucei*, arises from several factors. First, *T. cruzi* shows significant variability in chromosomal sizes across strains, which limits the exact resolution of well-defined bands in PFGE. Additionally, its genome exhibits extreme plasticity, characterized by multigene families, repetitive elements, and retroelements (Pita et al., 2019). Furthermore, homologous chromosome pairs often differ in size, leading to a complex and unresolved banding pattern.

After completing the genome projects for these three organisms - the "TriTryps" - many results obtained through PFGE were confirmed for *L. major* (Ivens et al., 2005) and *T. brucei* (Berriman et al., 2005). These studies also established a comparative chromosomal map between these two species, revealing regions of high synteny and providing insights into their common ancestor (El-Sayed et al., 2005b). However, *T. cruzi* was not included in this comparison since the scaffolds obtained were highly fragmented due to the repetitive nature of its genome, leading to a high incidence of regional collapse (El-Sayed et al., 2005a). Consequently, although syntenic regions were identified as shared among the TriTryps, chromosomal mappings systematically excluded *T. cruzi*. Later, Weatherly et al. (2009) proposed that *T. cruzi* possesses 41 chromosomes, based on BAC-end sequencing combined with chromosomal co-location and synteny with *T. brucei* and *Leishmania*, enabling the assembly of 41 *in silico* chromosomes. Although this estimation was derived from genome sequences prone to collapse due to repetitive regions, it has served as a valuable reference for subsequent studies.

The advent of long-read sequencing marked a significant advancement in studying complex genomes, such as *T. cruzi*, which contain numerous repetitive elements and multicopy genes that are highly similar or even identical and arranged in tandem. The use of PacBio or Nanopore methodologies enabled researchers to determine a more complete genome landscape by overcoming repetitive regions and generating scaffolds that, in some cases, have lengths compatible with the expected size of chromosomes (Berná et al., 2018; Callejas-Hernandez et al., 2018; Díaz-Viraqué et al., 2019; Wang et al 2021; Hakim et al., 2024). These studies provided a comprehensive view of genome organization and opened the door for comparative genomic analyses among the TriTryps. For instance, it was determined that multigene families encoding surface proteins are not situated in subtelomeric regions, as previously proposed, but clustered along different chromosomes. In this regard, a key discovery was the identification of two distinct genomic compartments in *T. cruzi*: the core and disruptive regions (Berná et al., 2018), the latter being non-syntenic with other trypanosomatids and housing hundreds of gene variants encoding surface proteins such as TS, mucins, and MASP. Furthermore, recent findings have shown that these genomic compartments correlate with specific three-dimensional chromatin organizations designated as C and D, which represent lower and higher levels of chromatin compaction, respectively, which in addition affects global gene expression (Lima et al., 2022; Díaz-Viraqué et al., 2023).

However, the lack of telomeric sequences at their ends prevents their classification as chromosomes. The sole exception is the genome of the Dm25 strain, which consists of 24 scaffolds with telomeres at both ends, confirming their identity as complete chromosomes (Sanchez et al., 2024). This significant advancement is primarily attributed to PacBio HiFi technology. Considering these precedents, we aimed to determine the complete karyotype of *T. cruzi*, elucidate the chromosomal organization of its core and disruptive compartments, and characterize the structural features of its telomeric and subtelomeric regions.

## Results

### 1. Chromosome-Level Assembly Defines the 32-Chromosome Karyotype of *T. cruzi*

The reference strain Dm28c was selected for this study because of its extensive characterization regarding cell cycle dynamics (Contreras et al., 1988) and prior genomic studies (Berná et al., 2018; Díaz-Viraqué et al., 2023; Grisard et al., 2014). Total genomic DNA was sequenced using PacBio HiFi technology, generating 301,047 reads, 2.93 Gb of sequenced bases, and an average read quality of Q29 (**Table 1**). The assembly was performed with the HiFiAsm algorithm (Cheng et al., 2021), resulting in an estimated genome size of 36.18 Mb with approximately 80× coverage. The assembly comprises 32 scaffolds, with an average GC content of 51.06% (**Table 1**). Analysis of scaffold ends revealed the presence of telomeric repeats (CCCTAA/TTAGGG) (Barros et al., 2012) at both ends of all 32 scaffolds, immediately adjacent to the conserved 189 bp junction sequence (Chiurillo et al., 1999), indicating that each scaffold represents a complete chromosome. Therefore, we determined that *T. cruzi* has 32 chromosomes, numbered sequentially from 1 to 32 in order of decreasing size (**Figure 1A and B**). Chromosome lengths range from 0.58 to 2.49 Mb (**Figure 1C and Table 1**). Finally, building on our previous development of an accessible website (Berná et al., 2018), we expanded that platform to include the Dm28cT2T genome. The updated platform, available at https://cruzi.pasteur.uy, maintains the original color scheme with minor modifications: RHS genes are shown in brown, TS in orange-red, DGF-1 in red, mucins and MASP in shades of blue, GP63 in orange, conserved genes in green, and pseudogenes in magenta (Supplementary Figure 1). This color scheme is consistently used throughout the manuscript. Likewise, chromosome orientation is displayed from 5ʹ to 3ʹ (left to right), and this convention is followed in all figures and descriptions involving chromosomal organization, directional gene clusters (DGC), telomeric and subtelomeric features, and comparative analyses.

**Figure 1.**
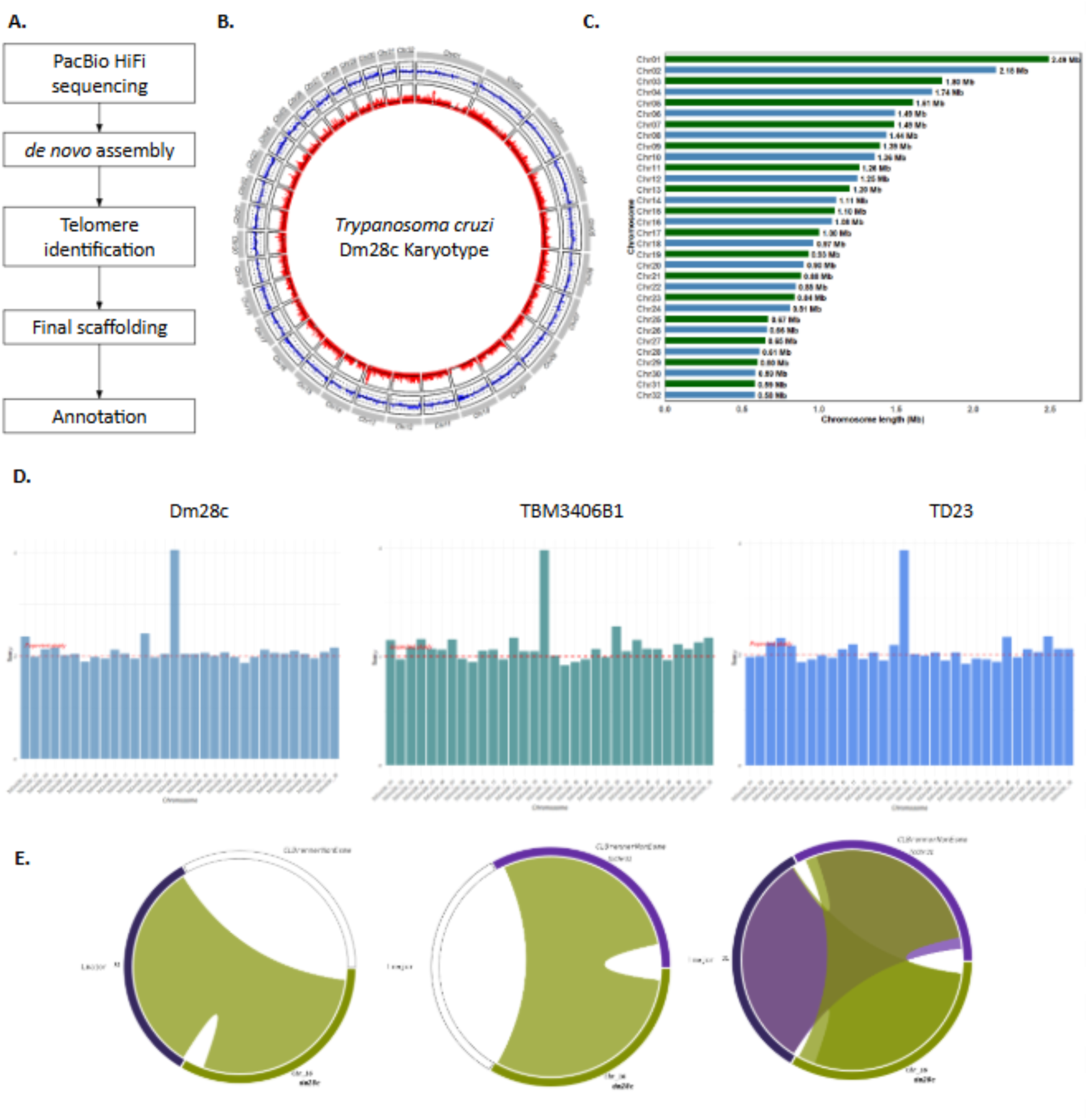
**The karyotype of *T. cruzi***. **A.** Assembly pipeline. **B.** Circos plot karyotype representation. The outer grey lines represent chromosomes; the middle circle represents the percentage of GC content along chromosomes (the black line indicates 50% GC, and the dotted black lines indicate 25% and 75% GC content). The inner circle in red indicates gene density calculated using overlapping windows of 5000 bases, sliding by 500 bases. **C.** Bar charts showing the *T. cruzi* chromosome lengths in Mb. **D.** Ploidy analysis for Dm28c (left), TBM3406B1 (center) and TD23 (right) strains. E. Syntenic regions (Symap representation) between *T. cruzi* chromosome 16, chromosome 31 *L. major* and *T. cruzi* CL Brenner strain..

**Table 1.** Sequencing, genome assembly statistics. Final genome assembly features and annotation.

| Elements | Metric | Value |
| --- | --- | --- |
| Sequencing Statistics | Number of reads | 301047 |
|  | Number of bases sequenced | 2.92 Gb |
|  | N50 | 10.68 Kb |
|  | Average read quality | Q29 |
| Genome Assembly Statistics | Number of contigs | 51 |
|  | Total length of assembly | 46.64 Mb |
|  | G+C content | 51.35 |
|  | Average coverage HiFi reads | 62.7 |
|  | N50 | 1.2 |
| Genome | Number of chromosomes | 32 |
|  | Genome length | 36.18 Mb |
|  | Longest chromosome | 2.49 Mb |
|  | Shortest chromosome | 0.58 Mb |
|  | Average coverage HiFi reads | 80.80 |
|  | N50 | 1.26 Mb |
|  | G+C content | 51.06 |
| Number of genes | Protein-coding genes | 13825 |
|  | RHS | 683 |
|  | Trans-sialidases | 1185 |
|  | Mucin | 384 |
|  | MASP | 530 |
|  | DGF-1 | 248 |
|  | Transposable elements | 1816 |
|  | Putative pseudogenes | 3218 |

Overall, these results provide a definitive resolution of the T. cruzi karyotype, establishing that its genome is organized into 32 complete chromosomes. This finding represents a significant milestone in T. cruzi genomics, as it definitively clarifies the parasite’s chromosomal complement, a question that has remained unresolved until now.

### 2. Gene Annotation of the T2T Genome of *T. cruzi*

The annotation pipeline (**Supplementary** Figure 2A) enabled the re-annotation of the Dm28c genome, revealing a haploid gene content of 13,766 protein-coding genes, including 1,781 putative pseudogenes (**Supplementary Table 1**). Approximately 22% of these genes belong to the most expanded multigene families: TS, mucins, MASP, RHS, GP63, and DGF-1 (**Table 1**). Compared to our previous annotation of Dm28c (Berná et al., 2018), the most significant difference is a ∼25% reduction in the number of core genes, from 12,230 to 9,916 (**Supplementary Table 1)**. This reduction results from methodological differences: in earlier assemblies, haplotypes could not be fully resolved. Consequently, genes in regions where both haplotypes were distinguishable were counted twice, whereas those in collapsed regions were counted only once. In contrast, the current annotation, based on PacBio HiFi sequencing, benefits from improved haplotype resolution. Gene counts for other multigene families varied, with some increasing and others decreasing (**Supplementary Table 1**), reflecting the enhanced continuity and resolution of repetitive regions (e.g., tandem, direct, and inverted repeats) and more accurate gene copy number estimation enabled by haplotype phasing.

For multigene families, we performed manual curation based on sequence alignments, cluster analysis, and the evaluation of coding sequence completeness. Although a detailed analysis exceeds the scope of this work, it is important to note that in one case, this process led to the identification of a previously unrecognized clade within the TS family. Further sequence analysis revealed that this clade corresponds to L1Tc, which had been misannotated as TS in UniProt (https://www.uniprot.org). As a result, 59 coding sequences originally classified as TS were removed from the annotation because they are part of the L1Tc retrotransposon family (**Supplementary** Figure 3). This finding underscores the necessity for comprehensive manual curation of the entire genome annotation and the revision of misclassified proteins in public repositories.

### 3- The *T. cruzi* Genome Is Diploid with Conserved Tetrasomy of Chromosome 16

We then determined the ploidy of *T. cruzi* by performing Illumina sequencing and mapping the reads onto the 32 chromosomes. Our analysis showed a diploid state across all chromosomes, except for chromosome 16, which consistently exhibited tetrasomy (**Figure 1D**). To determine if this pattern is unique to the Dm28c strain or a broader characteristic of *T. cruzi*, we analyzed Illumina read datasets from two additional strains: TBM3406B1 (Ecuador; SRA: SRR3676268) and TD23 (Texas; SRA: SRR3676272). In both cases, diploidy was maintained across all chromosomes except chromosome 16, which again showed clear evidence of tetrasomy (**Figure 1D**). Since tetrasomy has also been reported in the chromosome 31 of Leishmania and proposed to be conserved in trypanosomatids (Akopantis et al., 2009; Rogers et al., 2011; Reis-Cunha et al., 2024), we explored potential homology between these chromosomes. Remarkably, synteny analysis revealed that *L. major* chromosome 31 is homologous to chromosome 16 in our assembly (**Figure 1E**). Additionally, a putative tetraploid chromosome 31 has been identified in the *T. cruzi* CL Brener strain (Reis-Cunha et al., 2024), and comparative analysis has shown that it is also homologous to chromosome 16 (**Figure 1E**). These findings support the notion that this tetrasomic chromosome reflects a conserved ancestral tetraploidy shared among trypanosomatids.

### 4- Chromosome-Level Conservation in *T. cruzi*: A Stable Karyotype and Shared Synteny with *L. major*

It is widely accepted that *T. cruzi* exhibits significant genomic plasticity (Reis-Cunha et al., 2015; Cruz-Saavedra et al., 2022; Herreros-Cabello et al., 2025). In this context, we aimed to determine whether the complete karyotype described in this study is specific to the analyzed strain or represents a broader feature across the species. To investigate this, we used the Dm28c karyotype as a reference and compared it to haplotype 1 of the Dm25 strain. Although their names are similar, it’s important to note that Dm28c and Dm25 are not from the same clonal origin, as they were isolated in different regions and at different times (Contreras et al., 1988; Sanchez et al., 2024). Using this comparison, we could reassign the pseudo-chromosomes from the Dm25 molecular karyotype, especially those fragmented into multiple contigs and lacking telomeric regions. We found complete chromosomal correspondence in all cases (**Figure 2A**), with synteny and structural conservation clearly confirming homology (**Figures 2B, 2C, 2D**, and **Supplementary Figure 4**). For example, chromosome 13 maps to two different pseudo-chromosomes in Dm25 (contig c1 of pseudo-chromosome 16 and contig c2 of pseudo-chromosome 30), but maintains a highly conserved structure (**Figure 2B**). In cases where homologous chromosomes are fully assembled, we show two representative examples: one mainly composed of core genes and another enriched in disruptive genes. In both cases, the Dm28c chromosome has a clear homolog in Dm25, exhibiting remarkable similarity and conservation of gene content (**Figures 2C** and **2D**, **Supplementary Figure 4**). Synteny is well preserved in the core-enriched chromosomes (**Figure 2C** and **Supplementary Figure 4**), whereas the disruptive-enriched chromosomes, despite some differences—probably due to gene duplications or rearrangements—still retain their general structure (**Figure 2D** and **Supplementary Figure 4**). This approach allowed us to reconstruct the complete karyotype of the Dm25 strain, revealing that it also has 32 chromosomes ranging from 0.77 to 2.85 Mbp (**Figure 2A, Table 2**, and https://cruzi.pasteur.uy). The high similarity in chromosome size between homologous pairs, especially those resolved from telomere to telomere, is striking, and the most significant size differences were observed in those Dm25 chromosomes missing at least one telomeric end (Figure 2D). This high level of conservation across strains strongly suggests that a chromosome complement of 32 chromosomes is a characteristic feature of the *T. cruzi* species. However, analyzing additional clades is necessary to determine if all major *T. cruzi* lineages share this karyotype.

**Figure 2.**
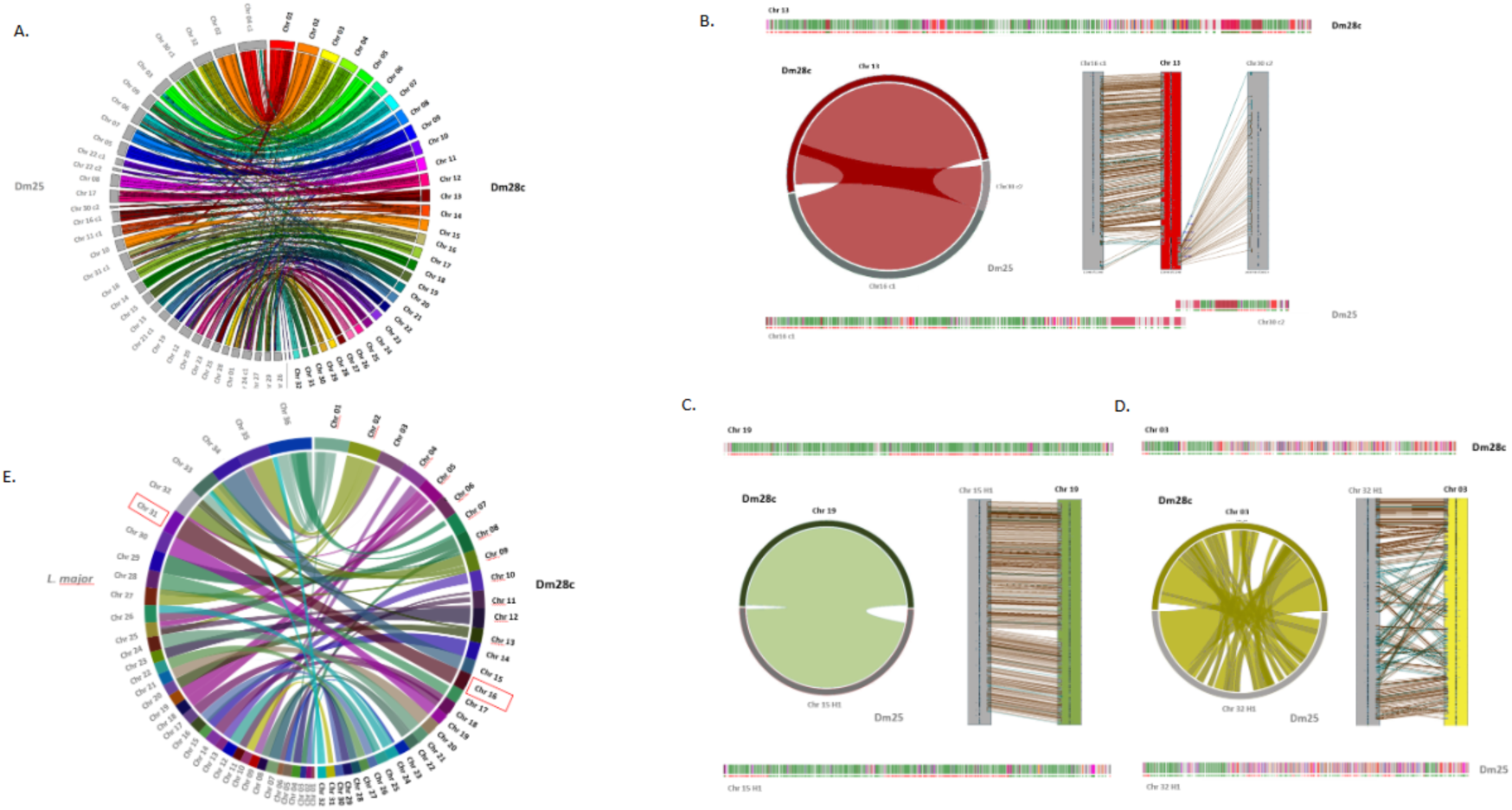
Comparison of the Dm28c and Dm25 genomes. **A.** The 32 chromosomes of *Dm28c* (colored) were mapped against all contigs present in haplotype 1 of *Dm25* (in gray). All contigs in the Dm25 assembly are represented in the 32 chromosomes of the Dm28c strain. **B.** The fragmented chromosome in Dm25 is assembled into the T2T chromosome in Dm28c. At the top (Dm28c) and bottom (Dm25) are schematic representations of chromosomes retrieved from the web page. In the middle, Symap results: Left panel: circos synteny plot; Right panel: synteny blocks. **C.** Example of synteny in the core chromosome and **D.** in the disruptive chromosome. At the top and bottom are schematic representations of chromosomes. In the middle, the results of synteny blocks obtained with Symap are represented by a circos plot and chromosomal representation similar to **C.** In all figures, gene representation colors align with the web interface’s color code (https://cruzi.pasteur.uy/): RHS (brown), TS (orange-red), DGF-1 (red), Mucin and MASP (shades of blue), GP63 (orange), Conserved genes (green), and Pseudogenes (magenta). E. Circos representation of synthenic regions between the Dm28c, *CLBrenner* and *L. major* determined with Symap. Red square shows proposed conserval aneuplody in trypanosomatids.

**Table 2.**
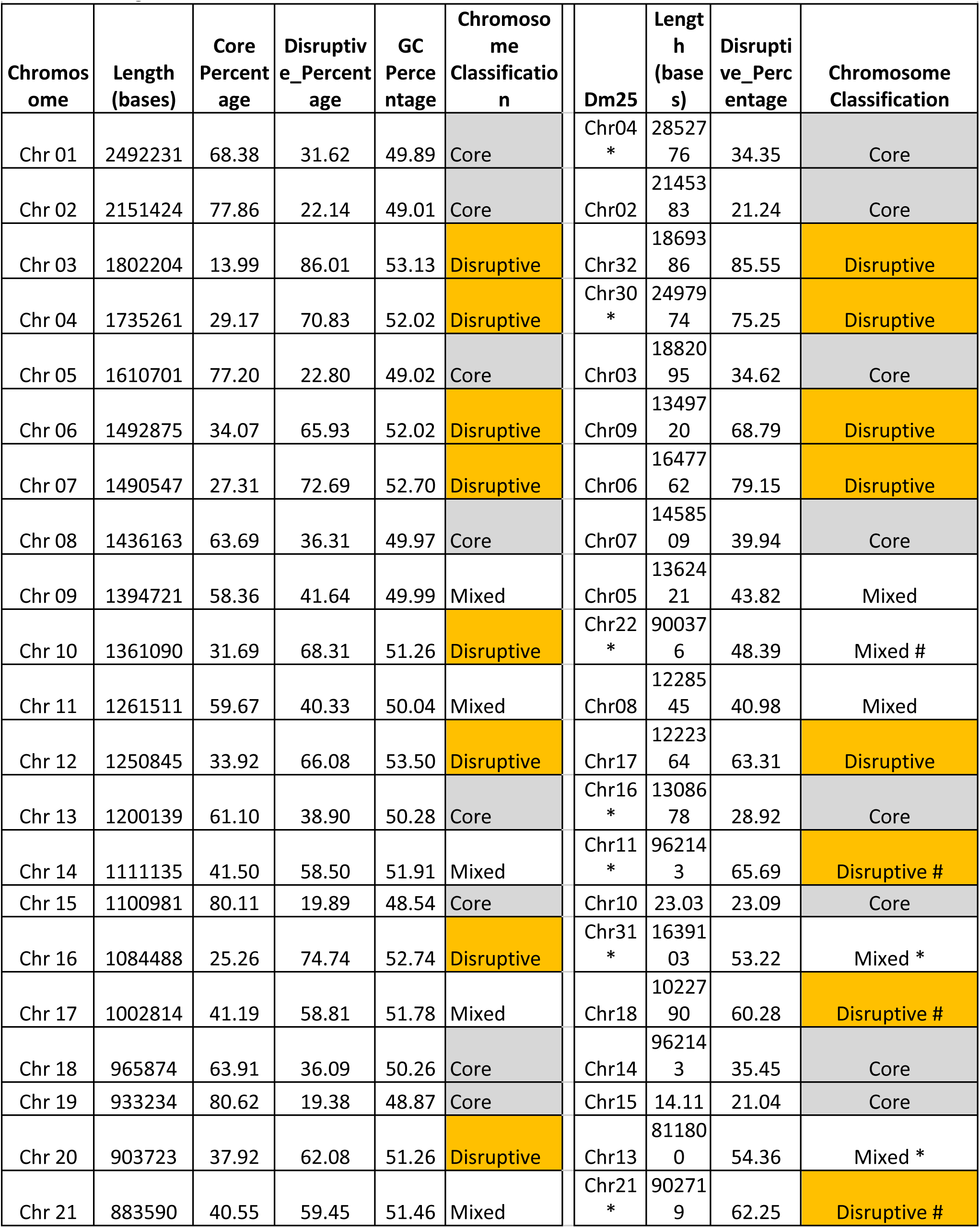
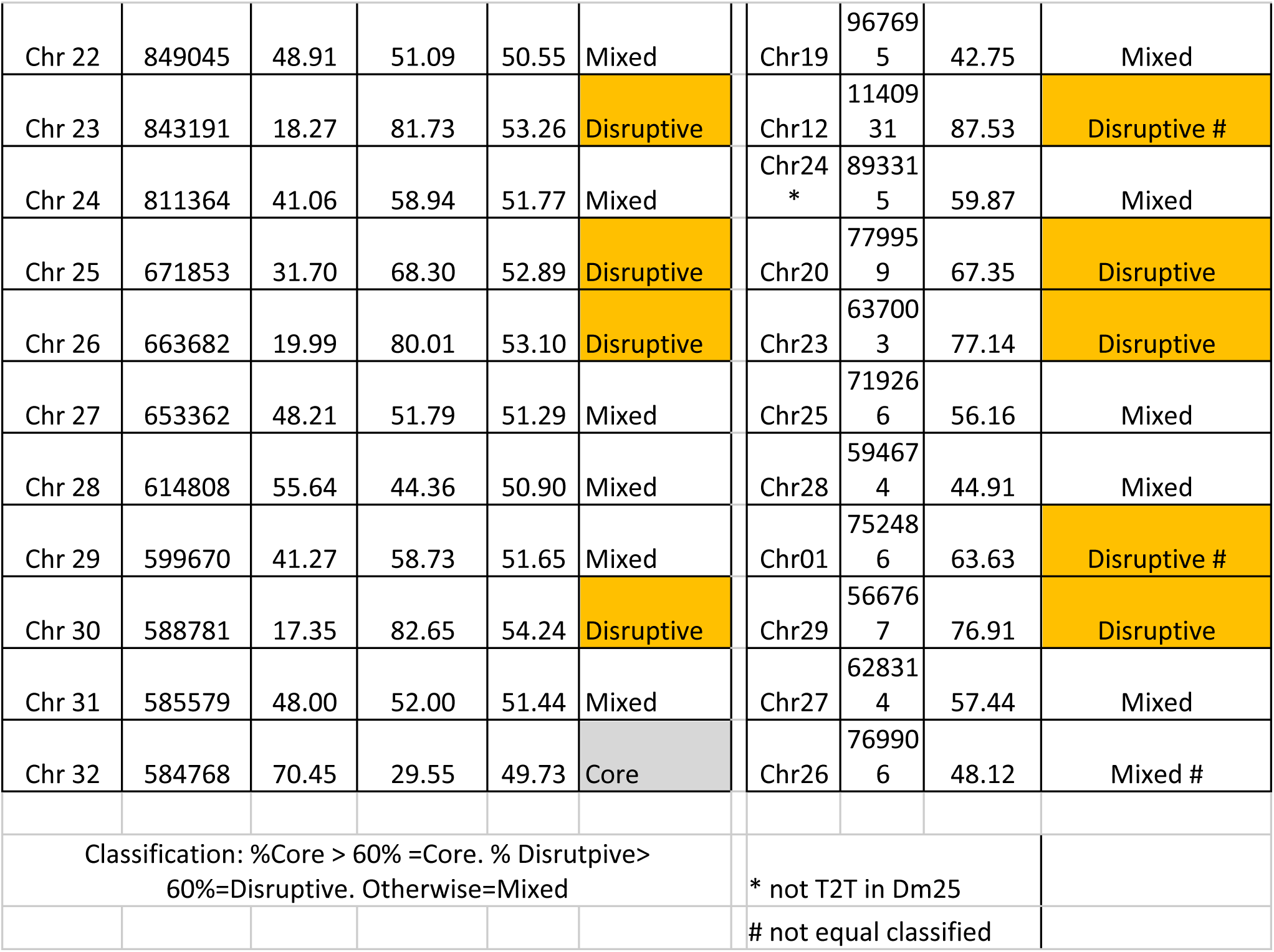
Chromosome classification by structure. Chromosome name, length in bases, core, disruptive and GC percentages and chromosome classification were included. Dm25 homolog chromosomes and their classification.

| Chromosome | Length (bases) | Core Percentage | Disruptive_Percentage | GC Percentage | Chromosome Classification | Dm25 | Length (bases) | Disruptive_Percentage | Chromosome Classification |
| --- | --- | --- | --- | --- | --- | --- | --- | --- | --- |
| Chr 01 | 2492231 | 68.38 | 31.62 | 49.89 | Core | Chr04 * | 2852776 | 34.35 | Core |
| Chr 02 | 2151424 | 77.86 | 22.14 | 49.01 | Core | Chr02 | 2145383 | 21.24 | Core |
| Chr 03 | 1802204 | 13.99 | 86.01 | 53.13 | Disruptive | Chr32 | 1869386 | 85.55 | Disruptive |
| Chr 04 | 1735261 | 29.17 | 70.83 | 52.02 | Disruptive | Chr30 * | 2497974 | 75.25 | Disruptive |
| Chr 05 | 1610701 | 77.20 | 22.80 | 49.02 | Core | Chr03 | 1882095 | 34.62 | Core |
| Chr 06 | 1492875 | 34.07 | 65.93 | 52.02 | Disruptive | Chr09 | 1349720 | 68.79 | Disruptive |
| Chr 07 | 1490547 | 27.31 | 72.69 | 52.70 | Disruptive | Chr06 | 1647762 | 79.15 | Disruptive |
| Chr 08 | 1436163 | 63.69 | 36.31 | 49.97 | Core | Chr07 | 1458509 | 39.94 | Core |
| Chr 09 | 1394721 | 58.36 | 41.64 | 49.99 | Mixed | Chr05 | 1362421 | 43.82 | Mixed |
| Chr 10 | 1361090 | 31.69 | 68.31 | 51.26 | Disruptive | Chr22 * | 900376 | 48.39 | Mixed # |
| Chr 11 | 1261511 | 59.67 | 40.33 | 50.04 | Mixed | Chr08 | 1228545 | 40.98 | Mixed |
| Chr 12 | 1250845 | 33.92 | 66.08 | 53.50 | Disruptive | Chr17 | 1222364 | 63.31 | Disruptive |
| Chr 13 | 1200139 | 61.10 | 38.90 | 50.28 | Core | Chr16 * | 1308678 | 28.92 | Core |
| Chr 14 | 1111135 | 41.50 | 58.50 | 51.91 | Mixed | Chr11 * | 962143 | 65.69 | Disruptive # |
| Chr 15 | 1100981 | 80.11 | 19.89 | 48.54 | Core | Chr10 | 23.03 | 23.09 | Core |
| Chr 16 | 1084488 | 25.26 | 74.74 | 52.74 | Disruptive | Chr31 * | 1639103 | 53.22 | Mixed * |
| Chr 17 | 1002814 | 41.19 | 58.81 | 51.78 | Mixed | Chr18 | 1022790 | 60.28 | Disruptive # |
| Chr 18 | 965874 | 63.91 | 36.09 | 50.26 | Core | Chr14 | 962143 | 35.45 | Core |
| Chr 19 | 933234 | 80.62 | 19.38 | 48.87 | Core | Chr15 | 14.11 | 21.04 | Core |
| Chr 20 | 903723 | 37.92 | 62.08 | 51.26 | Disruptive | Chr13 | 811800 | 54.36 | Mixed * |
| Chr 21 | 883590 | 40.55 | 59.45 | 51.46 | Mixed | Chr21 * | 902719 | 62.25 | Disruptive # |
| Chr 22 | 849045 | 48.91 | 51.09 | 50.55 | Mixed | Chr19 | 96769<br>5 | 42.75 | Mixed |
| Chr 23 | 843191 | 18.27 | 81.73 | 53.26 | Disruptive | Chr12 | 11409<br>31 | 87.53 | Disruptive # |
| Chr 24 | 811364 | 41.06 | 58.94 | 51.77 | Mixed | Chr24<br>* | 89331<br>5 | 59.87 | Mixed |
| Chr 25 | 671853 | 31.70 | 68.30 | 52.89 | Disruptive | Chr20 | 77995<br>9 | 67.35 | Disruptive |
| Chr 26 | 663682 | 19.99 | 80.01 | 53.10 | Disruptive | Chr23 | 63700<br>3 | 77.14 | Disruptive |
| Chr 27 | 653362 | 48.21 | 51.79 | 51.29 | Mixed | Chr25 | 71926<br>6 | 56.16 | Mixed |
| Chr 28 | 614808 | 55.64 | 44.36 | 50.90 | Mixed | Chr28 | 59467<br>4 | 44.91 | Mixed |
| Chr 29 | 599670 | 41.27 | 58.73 | 51.65 | Mixed | Chr01 | 75248<br>6 | 63.63 | Disruptive # |
| Chr 30 | 588781 | 17.35 | 82.65 | 54.24 | Disruptive | Chr29 | 56676<br>7 | 76.91 | Disruptive |
| Chr 31 | 585579 | 48.00 | 52.00 | 51.44 | Mixed | Chr27 | 62831<br>4 | 57.44 | Mixed |
| Chr 32 | 584768 | 70.45 | 29.55 | 49.73 | Core | Chr26 | 76990<br>6 | 48.12 | Mixed # |
| Classification: %Core > 60% =Core. % Disruptive><br>60%=Disruptive. Otherwise=Mixed |  |  |  |  |  | * not T2T in Dm25 |  |  |  |
|  |  |  |  |  |  | # not equal classified |  |  |  |

Based on the above-mentioned homology between *T. cruzi* chromosome 16 and *L. major* chromosome 31, we asked whether this interspecies conservation extends beyond this single example. To explore this, we broadened our comparative analysis to include all *L. major* chromosomes. This revealed a striking pattern: each core chromosome of *T. cruzi* has a homologous counterpart in L. major (Figure 2E), suggesting that the core chromosomal set is deeply conserved across trypanosomatids (**Figure 2E**).

Determining the complete chromosomal complement of *T. cruzi* constitutes a milestone that provides a reliable basis for investigating genome organization, synteny, and evolutionary dynamics, and will potentiate future studies on genome plasticity and comparative genomics.

### 5- Conservation of *T. cruzi* Chromosomal Architecture

To delve deeper into the conservation of chromosomal architecture, we focused on the organization of compartments. A significant difference previously reported is that the core and disruptive compartments differ markedly in their GC content, with the disruptive compartment showing much higher GC levels (Berná et al., 2018). In fact, GC content alone is often enough to predict whether a region belongs to the core or disruptive compartment. To measure the distribution of core and disruptive regions across chromosomes in relation to GC content, we utilized the disruptomics application developed by Balouz and Buscaglia (Balouz et al., 2025). Based on this analysis, chromosomes were classified into three groups: core (more than 60% of genes labeled as core), disruptive (more than 60% labeled as disruptive), and mixed (neither category exceeding 60%) (**Table 2**). The chromosomal landscape resulting from this classification is shown in **Figure 3A**. The graphic clearly demonstrates this correlation (**Figure 3A**), identifying nine core chromosomes (example in **Figure 3B**), twelve disruptive chromosomes (example in **Figure 3C**), and eleven mixed chromosomes. When we applied the same analysis to Dm25 chromosomes, they displayed a nearly identical pattern (**Figures 3B** and **3C**, boxed regions; **Supplementary Figure 5**). The corresponding chromosomes in both strains maintained the same classifications: core chromosomes remained core, disruptive chromosomes remained disruptive, and mixed chromosomes remained mixed (**Supplementary Figure 5**). These consistent compartmental profiles across strains further support a high level of structural conservation in chromosome organization.

**Figure 3.**
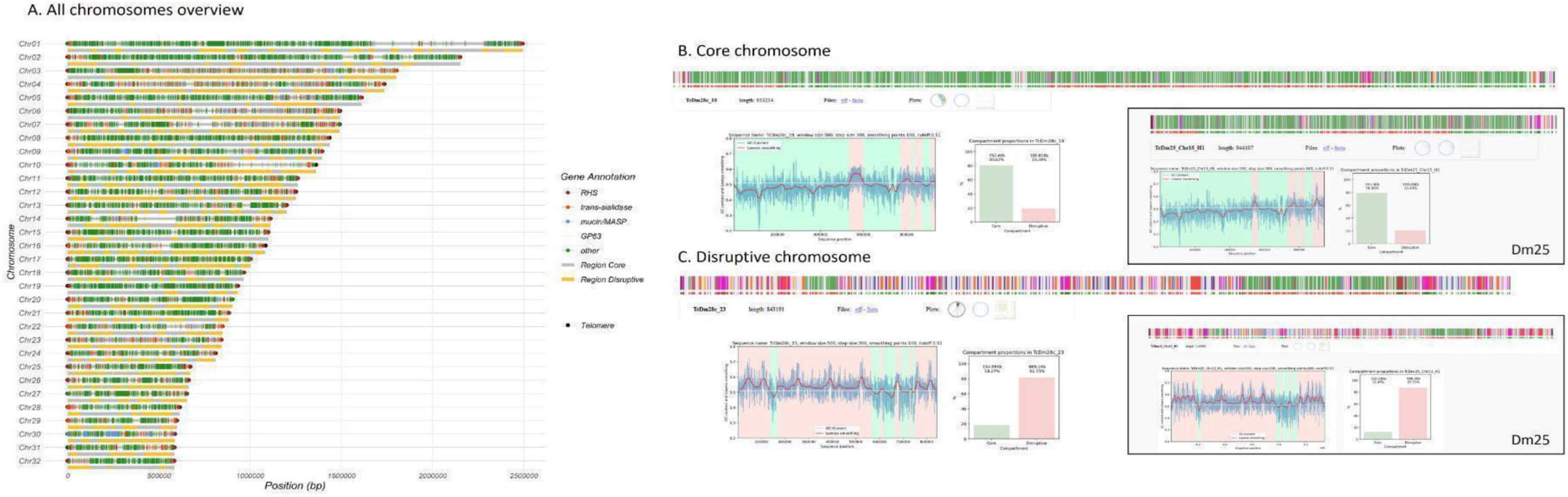
Architecture conservation. **A.** Representation of the 32 *T. cruzi* chromosomes. Below each chromosome, core and disruptive regions are shown in gray and orange, respectively. Telomeres are represented as black dots, and the first post-telomeric gene is indicated as dots or lines within chromosomes, color-coded according to the legend. **B.** Representation of a chromosome predominantly classified as Core. The plot below displays the GC content and categorization into core and disruptive regions, as determined by the GCScanner program (Balouz and Buscaglia, 2025). The graph on the right displays the percentage of each region type. The inset shows the same profile for the homologous chromosome of the Dm25 strain. **C.** Like **B**, displaying the profile of a chromosome classified as predominantly disruptive.

Overall, the finding that both the number of chromosomes and their compartmental architecture are conserved across *T. cruzi* strains challenges the current paradigm that genomic plasticity dynamically alters chromosome size and organization. Our results strongly indicate that the presence of 32 chromosomes and their organization into compartmental “blocks” are deeply conserved biological features of the *T. cruzi* species. Specifically, the distribution pattern of core and disruptive compartments shows remarkable conservation across strains, further supporting that the preservation of structural genomic organization is a fundamental property of *T. cruzi* genome architecture.

### 6- Subtelomeric Regions Define a Distinct Genomic Compartment in *T. cruzi*

All 64 chromosomal termini were confirmed to contain the expected hexameric repeat (CCCTAA/TTAGGG), followed by the 189 bp junction (Ramirez, 2020; **Figure 4A**). To examine the properties of the subtelomeres of *T. cruzi*, we employed a stepwise approach. First, we evaluated gene density in the first 50 kb adjacent to each telomere. The overall analysis of all chromosomes showed a significant decrease in gene density at the chromosomal ends, providing an initial rough boundary of the putative subtelomeric regions (**Figure 4B**). This decrease in gene density is observed in core, disruptive, and mixed chromosomes. Next, we analyzed their gene composition using sliding windows of five genes across the entire chromosomes and compared putative subtelomeres with internal regions. A primary observation was that all the subtelomeres comprise a unique DGC oriented toward the chromosomal ends (arrows in **Figure 4A**). Second, RHS is the most frequent gene adjacent to the 189 bp junction, appearing in over 80% of cases (**Figure 4C**). Third, RHS, TS, and DGF-1 are the most frequent protein-coding genes in the subtelomeres. Although core genes may be present, they do not consistently recur across subtelomeres (**Figure 4D and Supplementary Figure 6**). These results allowed us to define the subtelomeric boundary as the +1 nucleotide of the first conserved gene within the subtelomeric DGC—i.e., the gene located directly next to the internal (non-subtelomeric) region.

**Figure 4.**
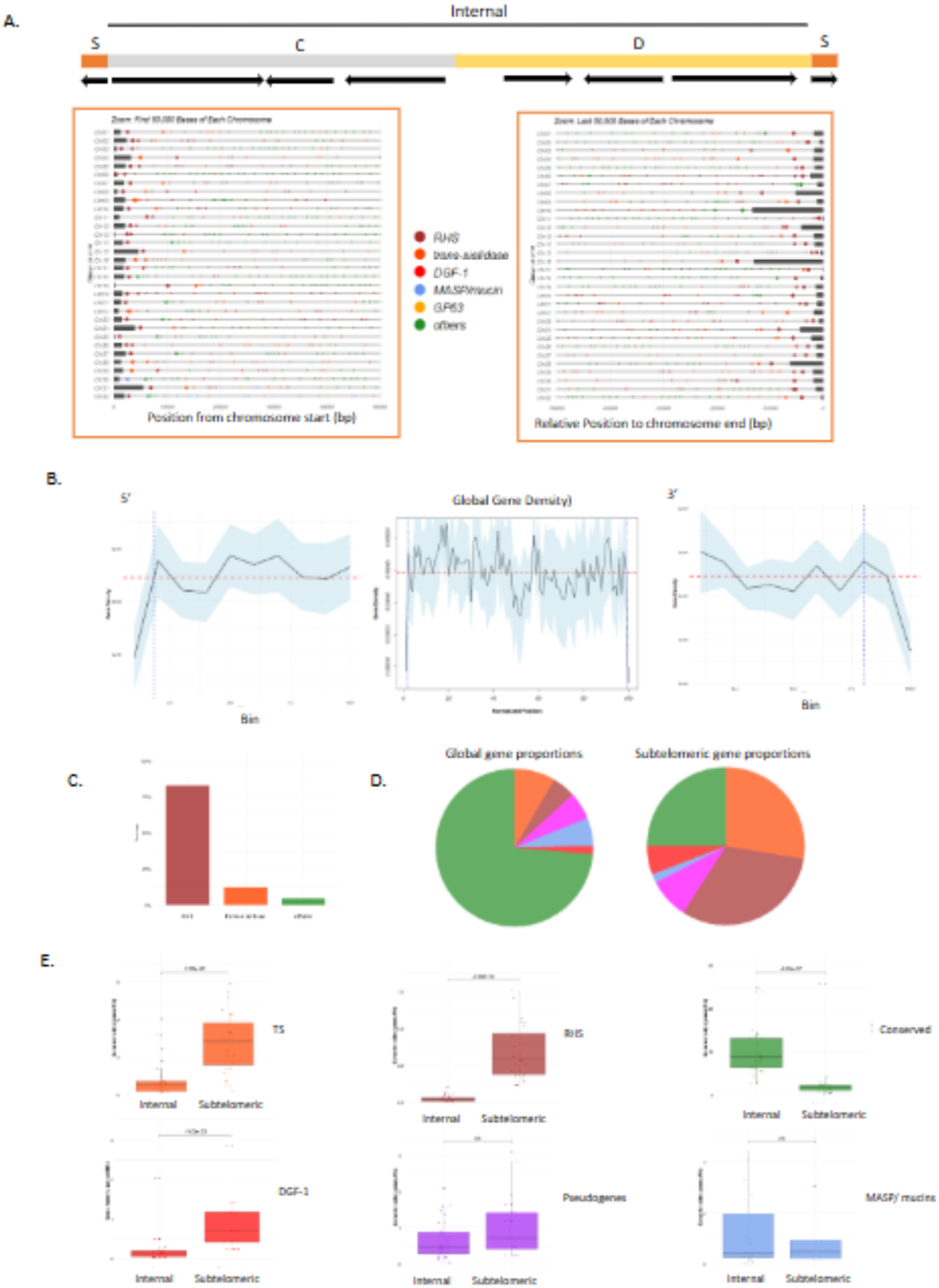
Telomeres and subtelomeres in *T. cruzi*. **A.** Representation of chromosomes with a zoom-in on the first and last 50 Kb. Dots indicate genes, and dark gray rectangles represent telomeric sequences. **B.** Gene density across chromosomes. Data aggregated from all 32 chromosomes, with lengths normalized into 100 bins. Gene density is shown for the first 10 bins (Crossing point= bin 1.83, bin length: 10.70-45.61Kb)(left plot), the entire chromosome (center plot), and the last 10 bins (Crossing point= bin 98, bin length: 5.85-24.92 Kb)(right plot). The red dotted line represents the mean gene density, while the blue line marks the crossing points at the beginning and end. **C.** Proportion of RHS, TS, mucin/MASP, or conserved genes detected as first gene post-telomere sequence. **D.** Proportions of genes in the subtelomeric region (right) and whole chromosomes (left). **E.** Boxplot representing global gene density (normalized by Mb) comparing internal and subtelomeric regions for Tran-sialidases, RHS, Conserved, DGF-1, pseudogenes and MASP/mucins.

We then compared the number of genes per megabase between subtelomeric and internal regions, confirming that TS, RHS, and DGF-1 are overrepresented in the subtelomeres, with statistical significance across all chromosomes (**Figure 4E**).This pattern persisted regardless of their classification as core, disruptive, or mixed (**Supplementary** Figure 7A). The same trend was observed for pseudogenes of TS, RHS, and DGF-1 (**Supplementary** Figure 7B).

To assess whether subtelomeres are a preferred environment for pseudogenization, we examined the distribution of genes and pseudogenes within the three gene families (**Table 3**). For the TS family, we found an uneven distribution: only 28.7% of its functional genes are located in subtelomeric regions, whereas 42.6% of its pseudogenes are located there, suggesting an accumulation or preferential retention of TS pseudogenes in these regions (**Table 3**). For the RHS and DGF-1 families, gene and pseudogene distributions were consistent, indicating no bias toward pseudogenization in either (**Table 3**). Finally, we observed an overrepresentation of L1Tc and SIRE retroelements in the subtelomeres (**Supplementary** Figure 8).

**Table 3.** TS, RHS and DGF-1 gene counts and proportions in subtelomeric and internal regions.

**Tabla 3.** TS, RHS and DGF-1 gene counts and proportions in subtelomeric and internal regions.
|  |  |  |  |  |  |
| --- | --- | --- | --- | --- | --- |
|  | **Trans-sialidasas** |  |  |  |  |
|  | **Internal** |  |  |  |  |
| **Genes and Pseudogenes** | **Core** | **Disruptive** | **Internal total** | **Subtelomeric** | **Total** |
| Number | 25 | 573 | 598 | 326 | 924 |
| Percentage | 2,71 | 62,01 | 64,72 | 35,28 | 100,00 |
| **Genes** |  |  |  |  |  |
| Number | 15 | 332 | 347 | 140 | 487 |
| Percentage | 3,08 | 68,17 | 71,25 | 28,75 | 100,00 |
| **Pseudogenes** |  |  |  |  |  |
| Number | 10 | 241 | 251 | 186 | 437 |
| Percentage | 2,29 | 55,15 | 57,44 | 42,56 | 100,00 |
|  | **RHS** |  |  |  |  |
|  | **Internal** |  |  |  |  |
| **Genes and Pseudogenes** | **Core** | **Disruptive** | **Internal total** | **Subtelomeric** | **Total** |
| Number | 30 | 159 | 189 | 446 | 635 |
| Percentage | 4,72 | 25,04 | 29,76 | 70,24 | 100,00 |
| **Genes** |  |  |  |  |  |
| Number | 28 | 108 | 136 | 293 | 429 |
| Percentage | 6,53 | 25,17 | 31,70 | 68,30 | 100,00 |
| **Pseudogenes** |  |  |  |  |  |
| Number | 2 | 51 | 53 | 153 | 206 |
| Percentage | 0,97 | 24,76 | 25,73 | 74,27 | 100,00 |
|  | **DGF-1** |  |  |  |  |
|  | **Internal** |  |  |  |  |
| **Genes and Pseudogenes** | **Core** | **Disruptive** | **Internal total** | **Subtelomeric** | **Total** |
| Number | 33 | 88 | 121 | 90 | 221 |
| Percentage | 14,93 | 39,82 | 54,75 | 40,72 | 100,00 |
| **Genes** |  |  |  |  |  |
| Number | 16 | 55 | 71 | 55 | 126 |
| Percentage | 12,70 | 43,65 | 56,35 | 43,65 | 100,00 |
| **Pseudogenes** |  |  |  |  |  |
| Number | 17 | 33 | 50 | 35 | 85 |
| Percentage | 20,00 | 38,82 | 58,82 | 41,18 | 100,00 |

### 7- Organization and Expression of the Subtelomeric Gene Families TS, RHS, and DGF1

The high representation of the TS genes in both the disruptive and subtelomeric compartments raises the question of whether the same TS variants are found in both regions. It is worth noting that the most detailed classification of TS genes into groups I to VIII was established over a decade ago (Freitas et al., 2011); however, the sequences used then may no longer reflect the current completeness, given advancements in long-read sequencing technologies. Therefore, we reanalyzed and categorized the TS family based on sequence alignments and similarity-based clustering. This revealed four main clades—called groups 1 to 4—each containing well-defined subclades (**Figure 5A**). Generally, these four clades correspond to the previously defined groups I to VIII, with some exceptions: Clade 1 includes groups I, III, and IV. Clade 2 consists of groups VII and VIII. Clade 3 corresponds to group II, which is further divided into at least two subclades. Clade 4 comprises groups V and VI, forming a single clade with five subclades (labeled a–e in **Figure 5A**), which contain members of both groups within each subclade. Next, we investigated whether there was a preference for TS genes to be located in subtelomeric or internal chromosomal regions. We found that Clades 1 and 3 mainly consist of subtelomeric TS genes (85% and 59% respectively) (**Figure 5A**, **B**, and **C**). In contrast, members of Clade 4 are primarily found in internal regions (95%). Clade 2 has an intermediate distribution (**Figure 5B** and **C**). To determine if TS gene expression varies across the T.cruzi life cycle, we mapped RNA-seq data from trypomastigotes, amastigotes, and epimastigotes onto the genome. We measured TS gene expression by TPM counts, and in all stages, subtelomeric TS genes are expressed at significantly higher levels than internal TS genes. Overall, their expression is highest in trypomastigotes, followed by amastigotes and epimastigotes (**Figure 5D**).

**Figure 5.**
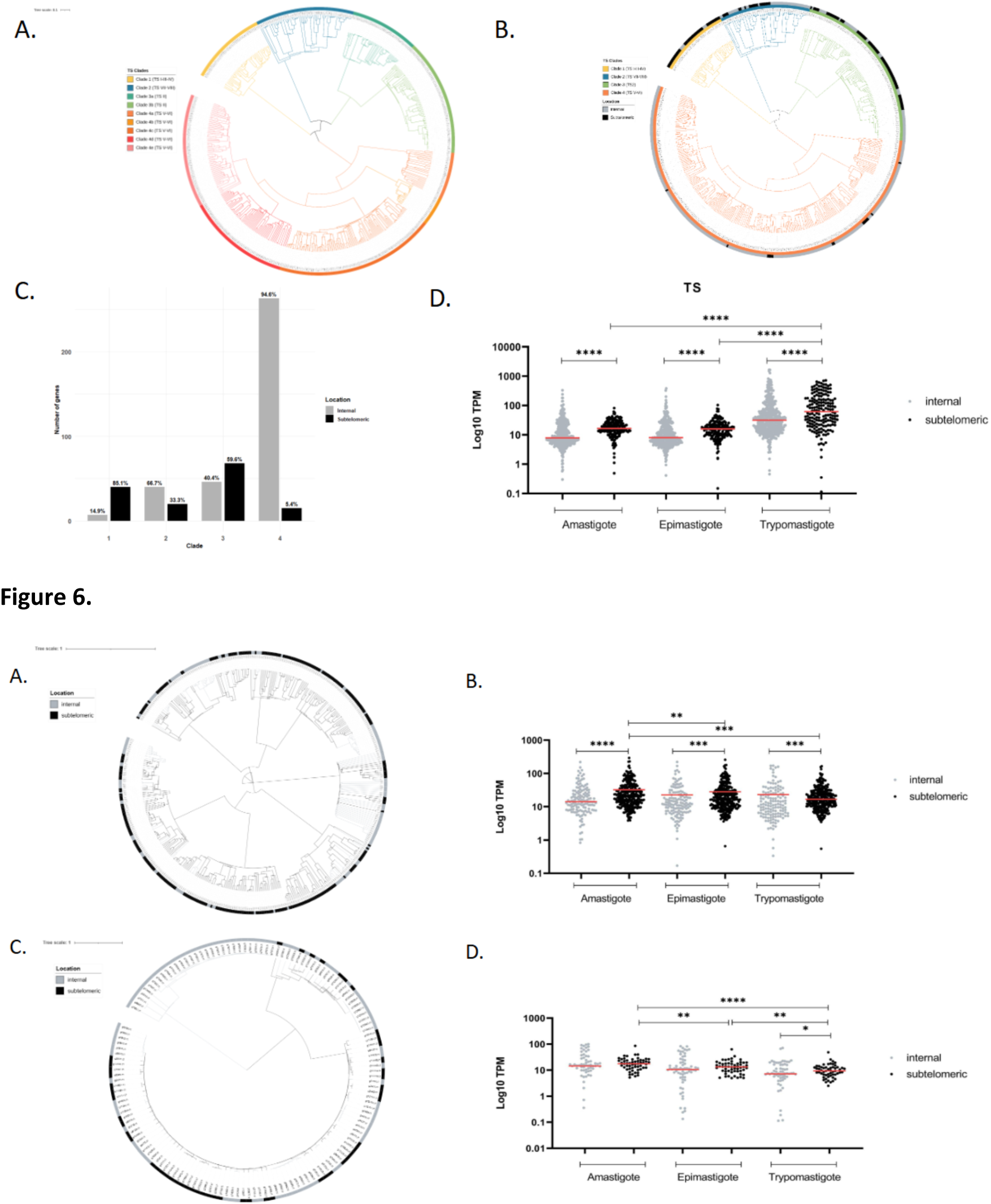
Guide tree and expression of Trans-sialidases. Sequences were aligned using Clustal Omega, and the tree was generated by the Neighbor-Joining method based on pairwise sequence distances and visualized using iTOL. **A.** Tree colored by clade and subgroup classification. **B.** Tree showing four clades and outer circle indicating location (subtelomeric or internal). **C.** Barplot showing the number of TS genes by clade and location. **D.** Expression levels (TPM) by genomic location in epimastigotes, amastigotes, and trypomastigotes. Statistical differences were assessed using the Mann–Whitney test. Asterisks indicate significance levels: p < 0.05 (*), p < 0.01 (*\*\*), and p < 0.001 (***).

Sequence-based clustering was also performed for RHS and DGF-1 genes. Unlike TS genes, they do not show a pattern of compartment-specific distribution (**Figure 6A** and **C**). The dendrograms reveal no tendency for subtelomeric genes to cluster within specific clades, indicating no correlation between sequence similarity and chromosomal location. Expression analysis shows that both gene families are mainly active during the replicative stages—epimastigotes and amastigotes—regardless of their chromosomal position (**Figure 6B** and **D**).

**Figure 6.** Guide tree and expression of RHS and DGF-1 proteins. **A.** RHS guided tree. Sequences were aligned using Clustal Omega based on pairwise sequence distances, and the tree generated by the Neighbor-joining method was visualized with iTOL. Subtelomeric or internal genomic location is indicated for each sequence (outer circle). **B.** Expression levels (tpm) by location in epimastigotes, amastigotes, and trypomastigotes were plotted for RHS genes. Statistical differences were assessed using the Mann–Whitney test. Asterisks indicate significance levels: p < 0.05 (*), p < 0.01 (**), and p < 0.001 (***). **C.** Guided tree as **A** for DGF-1 proteins. **D.** Same expression analysis shown in B, for DGF-1 genes.

These findings support the concept that subtelomeres are a distinct genomic compartment of *T. cruzi*. While the disruptive compartment constitutes a genome expansion mainly expressed in the trypomastigote stage, the subtelomeric region contains genes that are primarily expressed during either reproductive (RHS and DGF-1) or non-reproductive (TS) stages.

## Discussion

Twenty years ago, the draft genomes of *T. cruzi*, *L. major*, and *T. brucei* (referred to as TriTryps) were published, along with a comparative genomic analysis (Berriman et al., 2005; El-Sayed et al., 2005a, 2005b; Iven et al., 2005). At that time, only the *T. cruzi* genome could not be assembled at chromosomal resolution due to its highly repetitive nature, which caused extensive fragmentation. As a result, its chromosome number remained unknown. Additionally, chromatin does not condense sufficiently during mitosis to allow for the visualization of individual chromosomes, which hampers cytogenetic approaches. Therefore, pulsed-field gel electrophoresis (PFGE) became the standard method for estimating the karyotype in *T. cruzi*. Rigorous PFGE studies showed that the parasite has between 20 and 40 chromosomes, with sizes ranging from approximately 0.46 to 3.27 Mb across different lineages (Henriksson et al., 1990, 1996; Santos et al., 1997; Vargas et al., 2004; Henriksson et al., 1995; Galindo et al., 2007; Souza et al., 2011). However, although these studies produced distinguishable bands that appeared to be chromosomes, stoichiometric resolution was not always achieved because of overlapping bands, which made it difficult to determine the exact chromosome number. The emergence of long-read sequencing technologies, such as Nanopore and PacBio, has significantly advanced genome characterization and organization. However, while some studies referred to scaffolds assembled from contigs as chromosomes (Wang et al., 2021), these designations were overstated; it was only with the introduction of PacBio HiFi sequencing that complete chromosomes were finally recovered (Sanchez et al., 2024). Nonetheless, a fully complete chromosomal complement has yet to be achieved.

In this study, we successfully achieved the T2T assembly of the *T. cruzi* genome, resulting in a comprehensive characterization of its karyotype, which consists of 32 linear chromosomes, ranging in size from 0.58 to 2.49 Mb. Each chromosome contains canonical telomeric repeats of variable lengths and the conserved 189-bp junction sequence at both ends (Chiurillo et al., 1999; Kim et al., 2005; Freitas et al., 1999), confirming its chromosomal identity. Notably, PFGE analysis of the Dm28c strain previously reported chromosome sizes ranging from 0.57 to 2.50 Mb (Souza et al., 2011), which closely matches our findings and highlights the accuracy of earlier studies. Importantly, this karyotype is conserved even in genetically and geographically distant strains, such as Dm25. Each Dm28c chromosome maps to a syntenic homolog in Dm25, allowing for the reconstruction of the Dm25 karyotype and enabling detailed comparative genomics studies. Chromosome sizes are largely conserved, with most differences due to the absence of telomeric ends in Dm25. To evaluate the conservation of chromosomal architecture, we compared the GC content profiles of both karyotypes, revealing a high degree of similarity, with nearly identical GC content along each chromosome. However, it is essential to note that GC content and compartmental organization are not equivalent. For example, while DGF-1 genes have a high average GC content (∼63%), their presence is not limited to the disruptive compartment—they can also be located in subtelomeric regions or within the core. Nonetheless, GC profiling proved to be a helpful proxy for broadly delineating genomic compartments and identifying structural conservation across strains (Berná et al., 2018). We also observed remarkable conservation of ploidy among additional, genetically and geographically diverse strains. All studied strains are diploid, except for chromosome 16, which is consistently tetrasomic in all analyzed strains. Strikingly, the tetrasomic chromosome in *T. cruzi* corresponds to chromosome 31 in *L. major*, which is also tetrasomic. Finally, extensive synteny was observed between the core genomes of *T. cruzi* and *L. major*, highlighting a conserved chromosomal architecture among trypanosomatids.

Resolving the complete *T. cruzi* karyotype establishes a reference framework for a standardized chromosomal nomenclature (chromosomes 1 to 32) that serves as a basis for future genomic, transcriptomic, and proteomic studies. This facilitates the identification of structural variations associated with pathogenicity, virulence, host adaptation, and drug resistance, ultimately aiding in the development of more effective strategies to combat Chagas disease. Additionally, the remarkable conservation of chromosomal architecture across strains challenges the prevailing notion that *T. cruzi* exhibits extensive genome plasticity driven by widespread rearrangements. Instead, the discovery of a well-organized genome and conserved chromosomal structure suggests a paradigm shift: large-scale chromosomal rearrangements are not the primary drivers of plasticity. The presence of a conserved genome structure across strains strongly indicates that evolutionary constraints support a model in which plasticity occurs within a stable and regulated genomic framework. This means that *T. cruzi* generates diversity not through random, large-scale rearrangements but through localized and controlled mechanisms such as recombination, duplication, and small-scale rearrangements. This model aligns with our previous findings on the conservation of three-dimensional chromatin organization (Díaz-Viraqué et al., 2023). The genome is organized into chromatin folding domains (CFDs), which are closely associated with gene expression and are conserved across strains and life cycle stages. Furthermore, our 3D studies revealed that, at the 3D level, the *T. cruzi* genome is partitioned into two 3D compartments, C and D, corresponding to the core and disruptive genomic regions, respectively. While predominantly intrachromosomal interactions characterize compartment C, compartment D shows extensive interchromosomal contacts —a conformation that favors discrete sites for recombination and rearrangement, thereby aiding gene family diversification and antigenic variability without threatening chromosomal integrity. These observations argue against a model of stochastic genome organization and instead support a structured, compartmentalized form of plasticity. We propose that chromatin organization represents the key link between the highly ordered chromosomal structure and the genomic variability observed in *T. cruzi*.

The observed synteny across chromosomes from different *T. cruzi* strains is completely lost at the chromosomal ends, as shown in the CIRCOS plots, highlighting the variability of these terminal regions. This prompted us to define and analyze the gene organization of all 64 subtelomeric regions. These regions exhibit a sharp decrease in gene density, a strong enrichment of retroelements, and a complete loss of synteny between strains. Each subtelomere contains a unique directional gene cluster (DGC) oriented toward the telomere, along with a characteristic buildup of the multigene families RHS, TS, and DGF-1. In contrast, mucin and MASP genes are nearly absent. The specific subtelomeres previously reported and analyzed through cloning and sequencing strategies have been shown to be enriched in these genes (Chiurillo et al., 1999, 2002; Freitas et al., 1999; Kim et al., 2005). RHS genes appear to play a crucial role in subtelomeric architecture and function, as most are located in these regions and, in approximately 80% of cases, lie immediately adjacent to the 189-bp telomeric junction. Conversely, DGF-1 genes are more widely distributed across the genome, found in both disruptive and subtelomeric regions, as well as 20% in core regions, suggesting a broader function not limited to subtelomeres. Regarding TS genes, a general analysis suggests they are relatively evenly spread across genomic compartments. However, a closer look shows that this is not entirely accurate. The trans-sialidase family, initially divided into eight groups (I-VIII), needs reevaluation based on the now-available full-length gene sequences. Our analysis reveals that groups I, III, and IV form a single, well-supported clade, which we now refer to as TS group 1. Within this clade, three distinct subclades are identifiable. Notably, these subclades do not directly correspond to the previously described groups I, III, and IV. Similar findings were seen for the other main clades 2, 3, and 4, emphasizing the need for a revised classification based on phylogenetic relationships derived from complete TS gene sequences from different strains. Most genes from the three subtelomeric gene families are transcriptionally active. Unlike the disruptive compartment, which is predominantly enriched in genes required during the trypomastigote stage, subtelomeres harbor genes expressed either in replicative stages (RHS and DGF-1) or in non-replicative stages (TS).

Finally, the association of RHS and DGF-1 genes emerges as a hallmark of subtelomeric regions, while the co-occurrence of MASP and mucin genes characterizes the disruptive compartment. TS genes, in contrast, are spread across both compartments, with specific clades showing a tendency to be enriched in one or the other. Notably, the exact functions of these multigene families remain largely unknown, even for trans-sialidases, whose name is misleading: only members of group I have the enzymatic activity that defines the family.

In summary, these findings converge on a model in which *T. cruzi*’s genome is organized into three functionally distinct compartments—core, disruptive, and subtelomeric—each playing different roles in the parasite’s adaptability and evolution. The core compartment provides a conserved backbone necessary for parasite survival and is maintained across trypanosomatids; the disruptive compartment contains specific gene expansions of genes overexpressed in the infective and non-replicative trypomastigote stage; and the subtelomeric compartment, now clearly characterized, functions as a hotspot for gene diversification and pseudogenization, acting as a reservoir for antigenic diversity and genomic change. This chromosomal and compartmental organization offers a strong framework for understanding how *T. cruzi* maintains genome integrity while enabling high levels of functional plasticity. The reference-quality genome assembly and annotation provided here lay the foundation for future research on *T. cruzi* genome evolution, antigenic diversity, and host–pathogen interactions. Besides delivering a structural and functional genome map, this study allows for detailed analysis of the genomic, epigenetic, and regulatory features that influence the balance between conservation and variability, and ultimately support the parasite’s ability to adapt, persist, and interact with its host.

## Data availability

The raw data (PacBio HiFi and Illumina reads) and genome were deposited in NCBI under Accession number PRJNA1173111. The genome and annotation can also be accessed using our webpage: www.cruzi.pasteur.uy/.

## Materials and Methods

### Genome Visualization and Web Interface

As described in Results, to allow data visualization and usability, we utilized that model again with minor modifications. The new Dm28cT2T genome assembly and annotation are accessible through https://cruzi.pasteur.uy/. As in the previous version, the direction of the DGC can be found underneath each represented chromosome. From each chromosome, the circos and Yass plots can be displayed, and the fasta and gff files can be downloaded (**Supplementary** Figure 1). As before, the additional functionalities were preserved, such as viewing gene annotations, retrieving nucleotide and amino acid sequences, and conducting various searches by annotation or keywords.

The genome is available at NCBI (NCBI: PRJNA1173111, Genome accession number: JBIQOJ000000000).

### DNA extraction and Sequencing

Cryopreserved epimastigotes of *T. cruzi* Dm28c_2018 (Berná et al., 2018), were grown in liver infusion tryptose (LIT) medium supplemented with 10 % fetal bovine serum at 28° C; total DNA was extracted using the Quick DNA Universal kit and Quick DNA HMW MagBeads Kit (Zymo Research, USA), quantified with Qubit (Invitrogen, USA). Five µg of each extraction were pooled and sent to the Macrogen service to perform PacBio Hi-Fi sequencing. 301,047 reads were obtained (Read N50=10688), representing 2,926,558,665 bases (73X coverage for an estimated 40 Mb genome) with an average read quality of Q29. Raw reads were deposited in SRA (SRR31351846). Illumina sequencing was performed in Macrogen service. 54,374,830 paired end reads (101 bp) were obtained (274X coverage). Raw illumina data was deposited in SRA under SRR33678280 accession number.

### Genome Assembly

The default HiFiAsm pipeline (Cheng et al., 2021) (hifiasm -o Prefix -t 32 Reads.fq.gz) generated an initial assembly of 51 contigs and two haplotypes with 80 and 86 contigs, respectively. In this first step, 25 telomere-to-telomere chromosomes were recovered (18 of them were also T2T reconstructed in previous *T. cruzi* Dm25 HiFi genome assembly (Wang et al., 2021; Sanchez et al., 2024). Contigs with telomeres at 5’ or 3’ ends were manually assembled using previously assembled T2T chromosomes from the Dm25 strain as a reference. Terminal telomeric signals (CCCTAA/TTAGGG) were detected in all the chromosomes assembled, indicating T2T assembled chromosomes. Chromosomes were sorted and numbered in order of decreasing length and designated as chromosomes 1 to 32. Their sizes range from 2.5 Mbp (TcDm28c_01) to 0.6 Mbp (TcDm28c_32). Consistent with previous reports, the maxicircle (TcDm28c_aMaxicircle) has a length of 50,477 bp (bases) and belongs to the type “a” and was recovered in one contig. Finally, 47 contigs of the first HiFiAsm output were used to reconstruct the 32 complete telomere-to-telomere chromosomes.

### Genome Annotation

To annotate the coding sequences, we first trained Augustus (Stanke and Morgenstern, 2005; Stanke et al., 2006) using T. cruzi (previous annotation of the Dm28c genome), to produce the initial annotation GFF file. We then corrected the start and stop codons of each CDS utilizing the getorf method as previously described (Rice et al., 2000). All CDS were retrieved and annotated using MMseqs (Mirdita et al., 2019) against the UniProt database (uniref_100). The coding sequences were categorized into three possible outcomes: no similarity to the UniProt database, similarity to the UniProt database, or truncated/multiple possible annotations (**Supplementary** Figure 1A). Busco (Manni et al., 2021) was used to evaluate the completeness of the gene space against the Euglenozoa database, indicating the quality of the assembly (**Supplementary** Figure 1B). The complete pipeline is summarized in **Supplementary** Figure 1A.

The web page featuring the new genome and its annotation was constructed based on a previous publication by our group (Berná et al., 2018) and is available at https://www.cruzi.pasteur.uy.

Tandem repeats and satellite sequences were annotated with the Tandem Repeat Finder (TRF) algorithm (Benson, 1999). Satellite regions were retrieved from the TRF output by filtering the repeat lengths between 190 and 200 bp. Additionally, we tested our annotation by comparing it with the algorithm developed by Dean et al. (2025) for classifying MASP genes, obtaining equivalent results.

### Ploidy analysis

For ploidy analysis, Illumina reads (SRR3676268, SRR3676272, and SRR33678280) were mapped against the Dm28c T2T genome assembly using minimap2 (Li, H., 2018). Samtools depth (Danecek et al. 2021) was used to calculate depth using sliding windows. Finally, Rscript (see next) was used to plot data.

### Computational analysis

All scripts used to produce data and figures were available on github (www.github.com/Gon1976/TcruziGenome). R scripts were used to estimate gene density along the chromosomes. Gene density was computed using the countOverlaps function from the GenomicRanges package in R (Lawrence et al., 2013). This analysis was performed for all chromosomes. Circos (Krzywinski et al., 2009) was used to create Figure 2A, using data from minimap2 comparison with Dm28c and Dm25 genomes. SyMap (Soderlund et al., 2011) was used for synteny mapping, and circos in Figure 2B, C, D, and E. The Circlize package (Gu et al., 2014) in R was used to create Figure 1, including information on GC content (calculated using the BioString package (Lawrence et al., 2013) and gene density. ggplot package in R was used to create gene representation in the subtelomeric regions using gff data.

Transcript expression was quantified from RNA-seq data using Salmon v1.5.1 (Patro et al., 2019). Each transcript is expressed in transcripts per million (TPM) units. Replicate consistency was evaluated using Pearson correlation and principal component analysis (PCA).

### TS, RHS, and DGF-1 analysis

For the analysis of TS coding genes, we first retrieved the TS sequences classified into eight groups as defined previously (Freitas 2011). From the initial dataset of 505 sequences, those shorter than 1300 nucleotides and lacking the VTV motif were excluded. The remaining sequences were aligned and guide trees were generated using Clustal Omega and visualized in iTOL to identify clusters of similar TS sequences. Based on these analyses, the sequences were regrouped into TS1, TS2, TS3, TS4, TS5-6, and TS7-8, and clustered using CD-HIT at 85% identity.

TS sequences from the Dm28c strain were manually curated by removing those lacking the characteristic TcS (VTV) motif of this family. A guide tree was then constructed with these sequences alongside TS sequences from CL Brener, allowing classification of Dm28c TSs according to the previously defined groups.

For RHS and DGF-1, sequences were retrieved from gff and assigned as internal or subtelomeric based on their coordinates. The sequences were aligned, and a guide tree was constructed using Clustal Omega, as previously described for TS, and visualized in iTOL.

TPM values were retrieved for all genes from expression data and plotted using R.

## Acknowledgments

We thank Matías Rodriguez (Institute of Bioinformatics, Faculty of Medicine, University of Münster, Münster, Germany) for facilitating the code for the webpage and for his significant assistance with the annotation scripts; L. Berna (I. Pasteur Montevideo, Uruguay) for providing the initial version of the script used to detect telomeres; A. C. Dean, V. Balouz, and C. Buscaglia (Universidad Nacional de San Martín, Argentina) for sharing the GCScanner algorithm and the MASP annotation pipeline before publication; J.L. Ramirez (Instituto de Estudios Avanzados-IDEA and Universidad Central de Venezuela) for valuable discussions and suggestions regarding chromosomes, telomeres, and subtelomeres; and all members of Laboratorio de Interacciones Hospedero-Patógeno — UBM (Institut Pasteur de Montevideo) for their helpful comments and engaging discussions. This work was supported by the Research Council United Kingdom Grand Challenges Research Fund (GCRF) ‘A Global Network for Neglected Tropical Diseases’ MR/P027989/1 (CR); Agencia Nacional de Investigación e Innovación (ANII, Uruguay) DCIALA/ 2011/023–502, ‘Contrato de apoyo a las políticas de innovación y cohesión territorial’; Fondo para la Convergencia Estructural del Mercado Común del Sur (FOCEM) 03/1 (C.R.); Pasteur Network ACIP 2024 n°1846 / Chromacruzi (FDV); and Agencia Nacional de Investigación e Innovación (ANII, Uruguay), FCE_3_2022_1_172653 (FDV). GG, FDV, MLC, CS, and CR are members of the Sistema Nacional de Investigadores (SNI-ANII, Uruguay) and the “Programa de Desarrollo de Ciencias Básicas” (PEDECIBA, Uruguay).

