## Supplementary material for "*Trypanosoma cruzi* has 32 Chromosomes: A Telomere-to-Telomere Assembly Defines Its Karyotype": SupFigure1_Web.pptx

### Slide 1
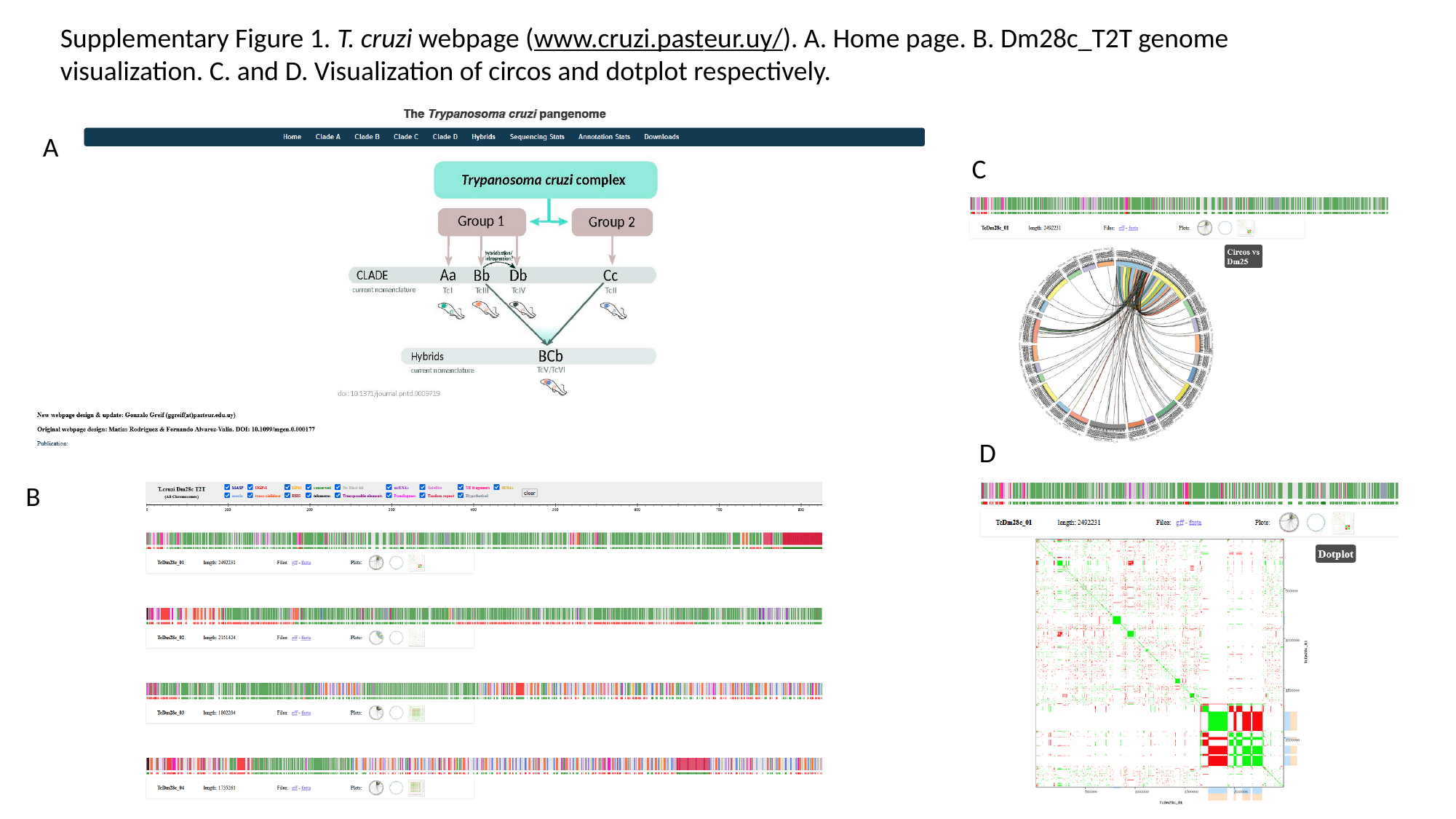

Supplementary Figure 1. T. cruzi webpage (www.cruzi.pasteur.uy/). A. Home page. B. Dm28c_T2T genome visualization. C. and D. Visualization of circos and dotplot respectively.
A
C
D
B
