## Supplementary material for "*Trypanosoma cruzi* has 32 Chromosomes: A Telomere-to-Telomere Assembly Defines Its Karyotype": SupFigure2_AnotationPipelineAndBusco.pptx

### Slide 1
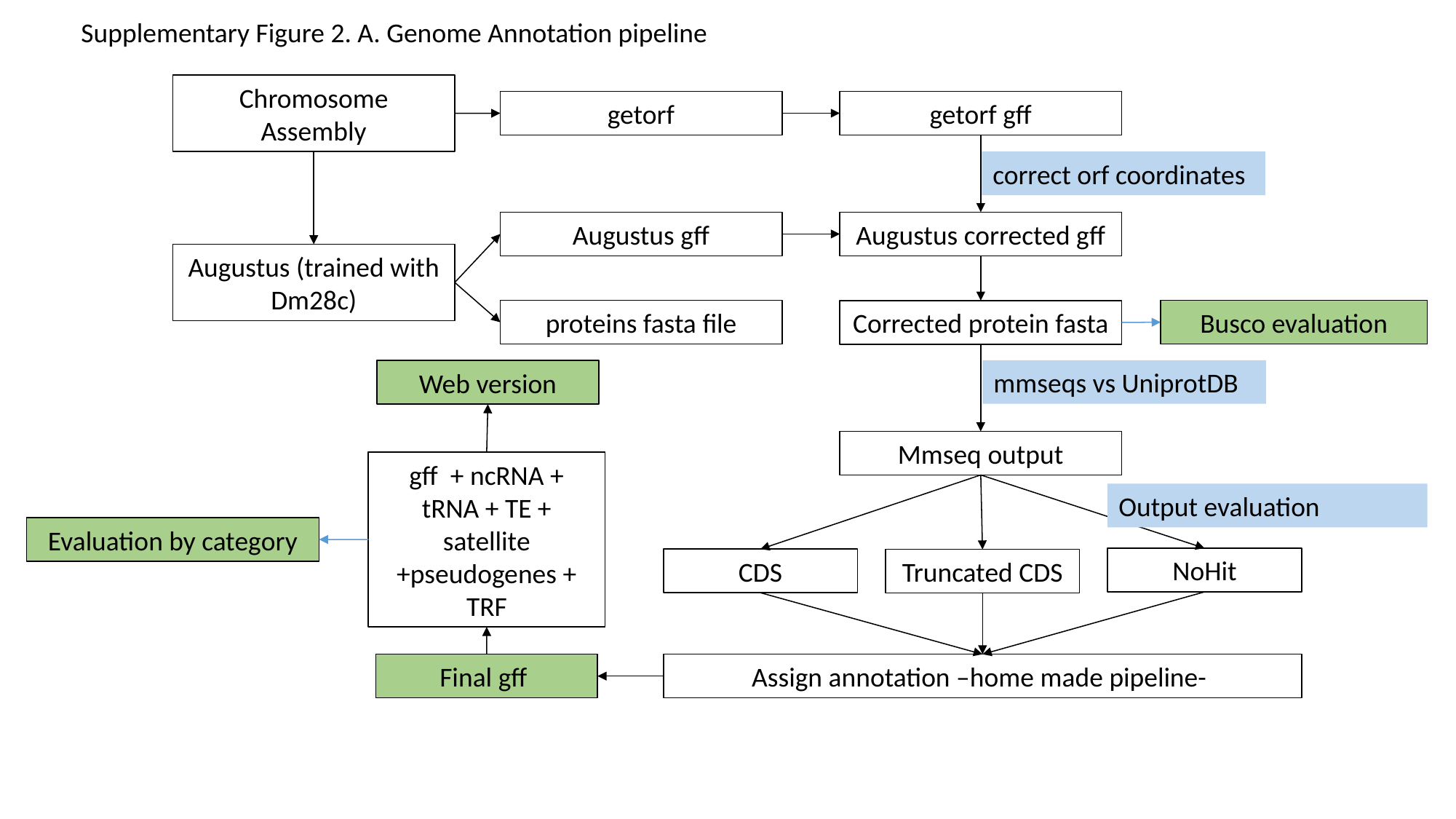

Supplementary Figure 2. A. Genome Annotation pipeline
Chromosome Assembly
getorf
getorf gff
correct orf coordinates
Augustus gff
Augustus corrected gff
Augustus (trained with Dm28c)
proteins fasta file
Busco evaluation
Corrected protein fasta
mmseqs vs UniprotDB
Web version
Mmseq output
gff + ncRNA + tRNA + TE + satellite +pseudogenes + TRF
Output evaluation
Evaluation by category
NoHit
CDS
Truncated CDS
Final gff
Assign annotation –home made pipeline-

### Slide 2
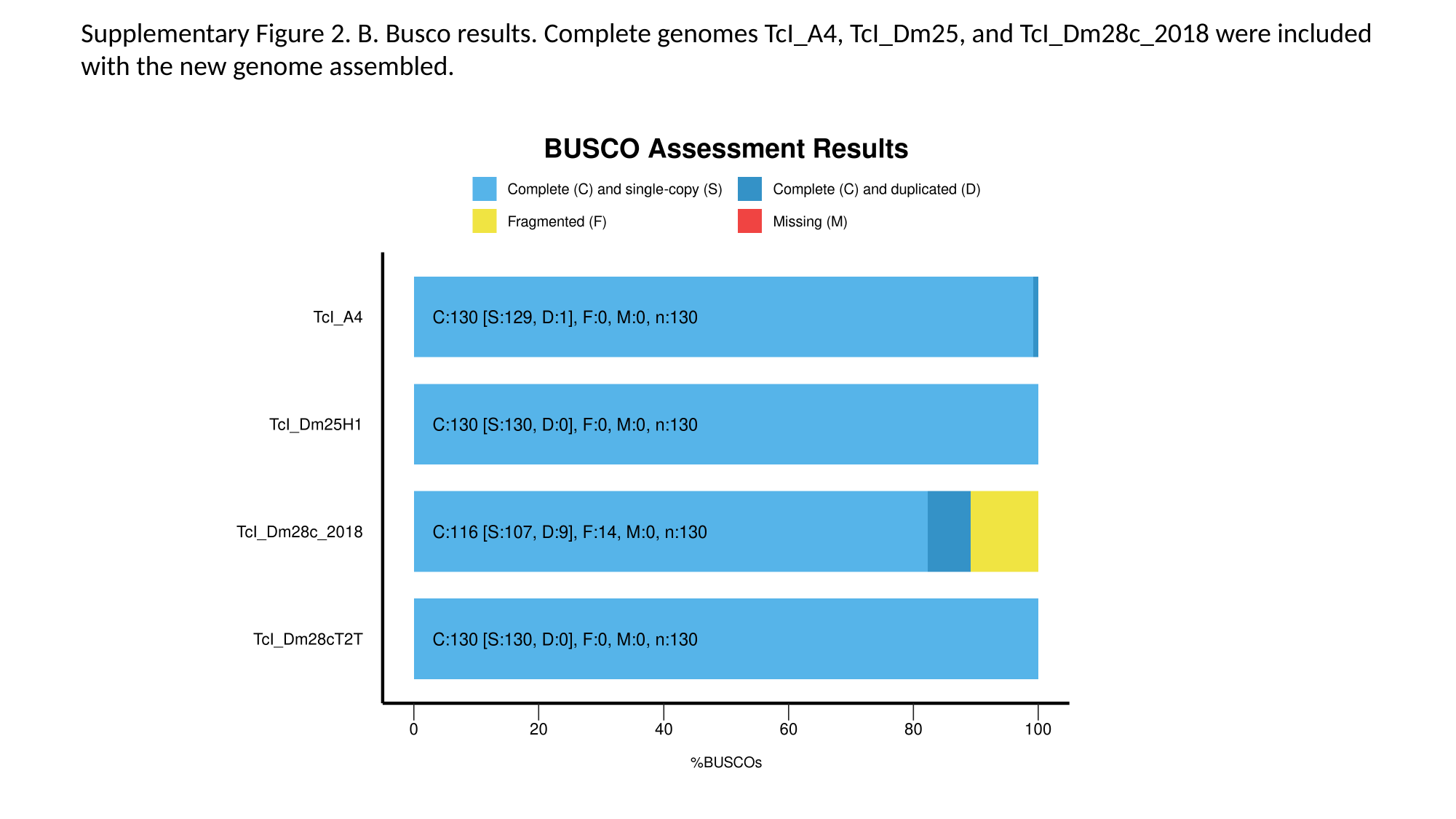

Supplementary Figure 2. B. Busco results. Complete genomes TcI_A4, TcI_Dm25, and TcI_Dm28c_2018 were included with the new genome assembled.
