## Supplementary material for "*Trypanosoma cruzi* has 32 Chromosomes: A Telomere-to-Telomere Assembly Defines Its Karyotype": SupFigure3_L1Tc_27May25.pptx

### Slide 1
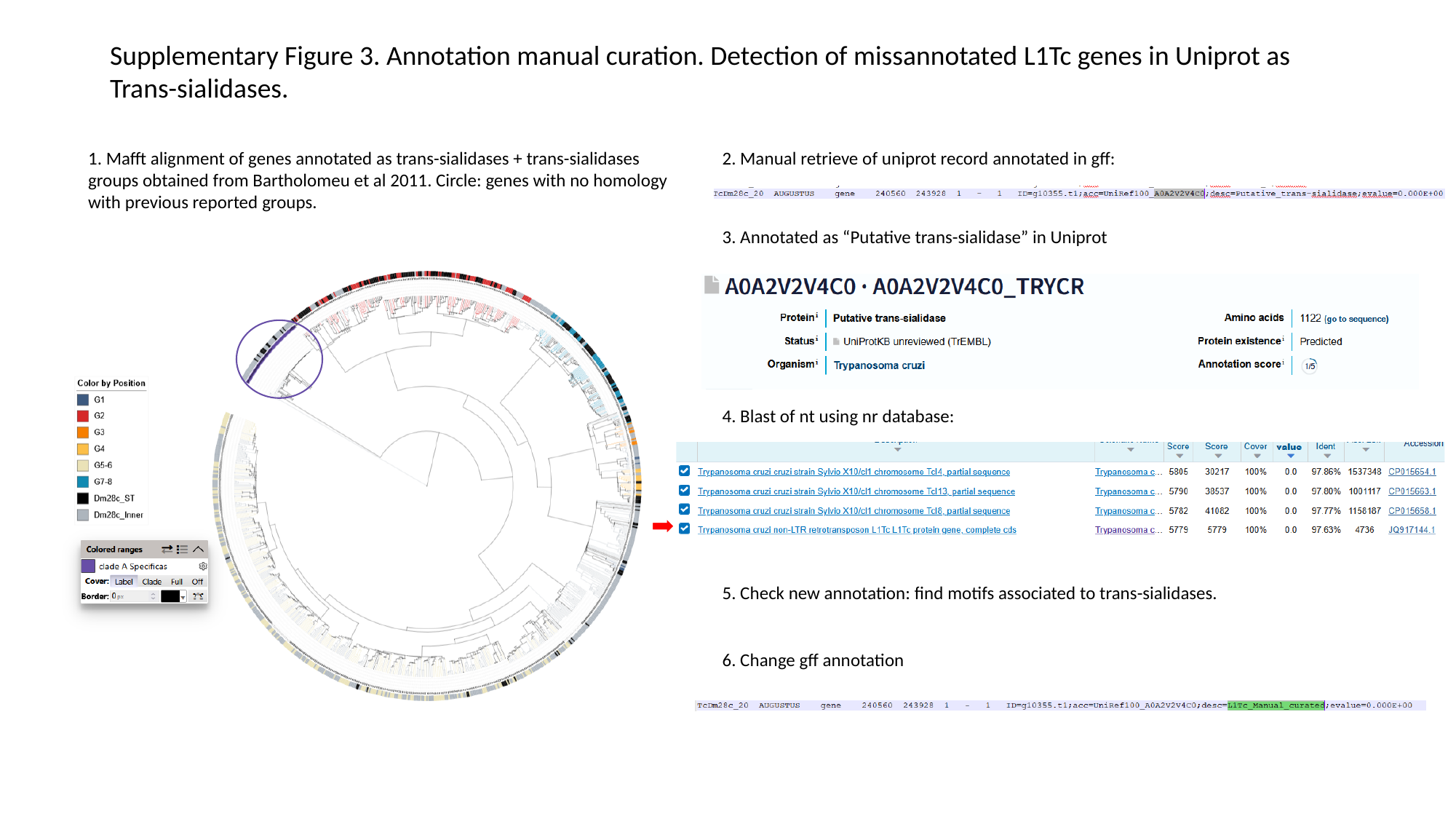

Supplementary Figure 3. Annotation manual curation. Detection of missannotated L1Tc genes in Uniprot as Trans-sialidases.
2. Manual retrieve of uniprot record annotated in gff:
1. Mafft alignment of genes annotated as trans-sialidases + trans-sialidases groups obtained from Bartholomeu et al 2011. Circle: genes with no homology with previous reported groups.
3. Annotated as “Putative trans-sialidase” in Uniprot
4. Blast of nt using nr database:
5. Check new annotation: find motifs associated to trans-sialidases.
6. Change gff annotation
