## Supplementary material for "*Trypanosoma cruzi* has 32 Chromosomes: A Telomere-to-Telomere Assembly Defines Its Karyotype": SupFigure4_Circos_AllChromosomesDm28cvsDm25_13Mayo25.pptx

### Slide 1
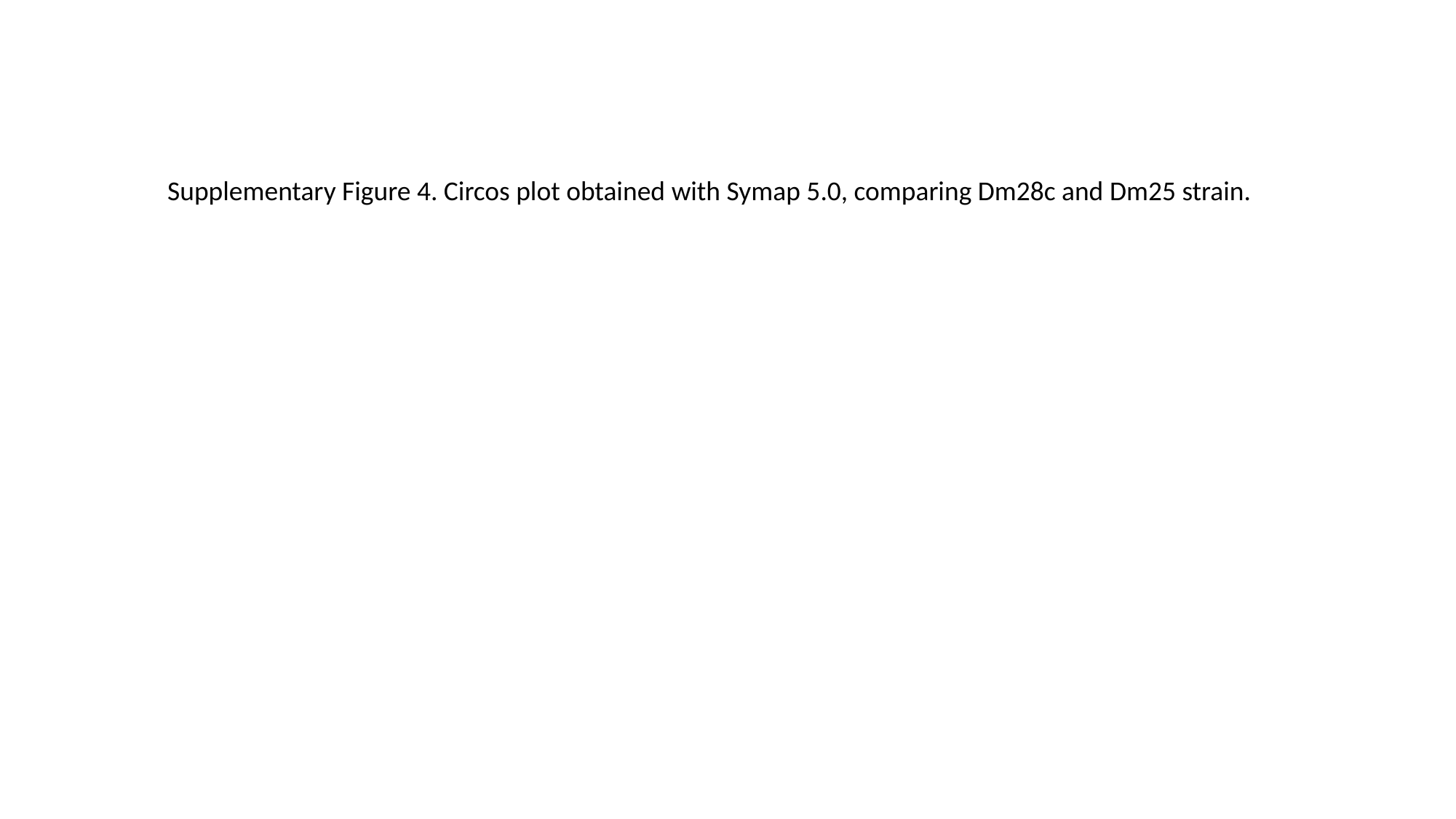

Supplementary Figure 4. Circos plot obtained with Symap 5.0, comparing Dm28c and Dm25 strain.

### Slide 2
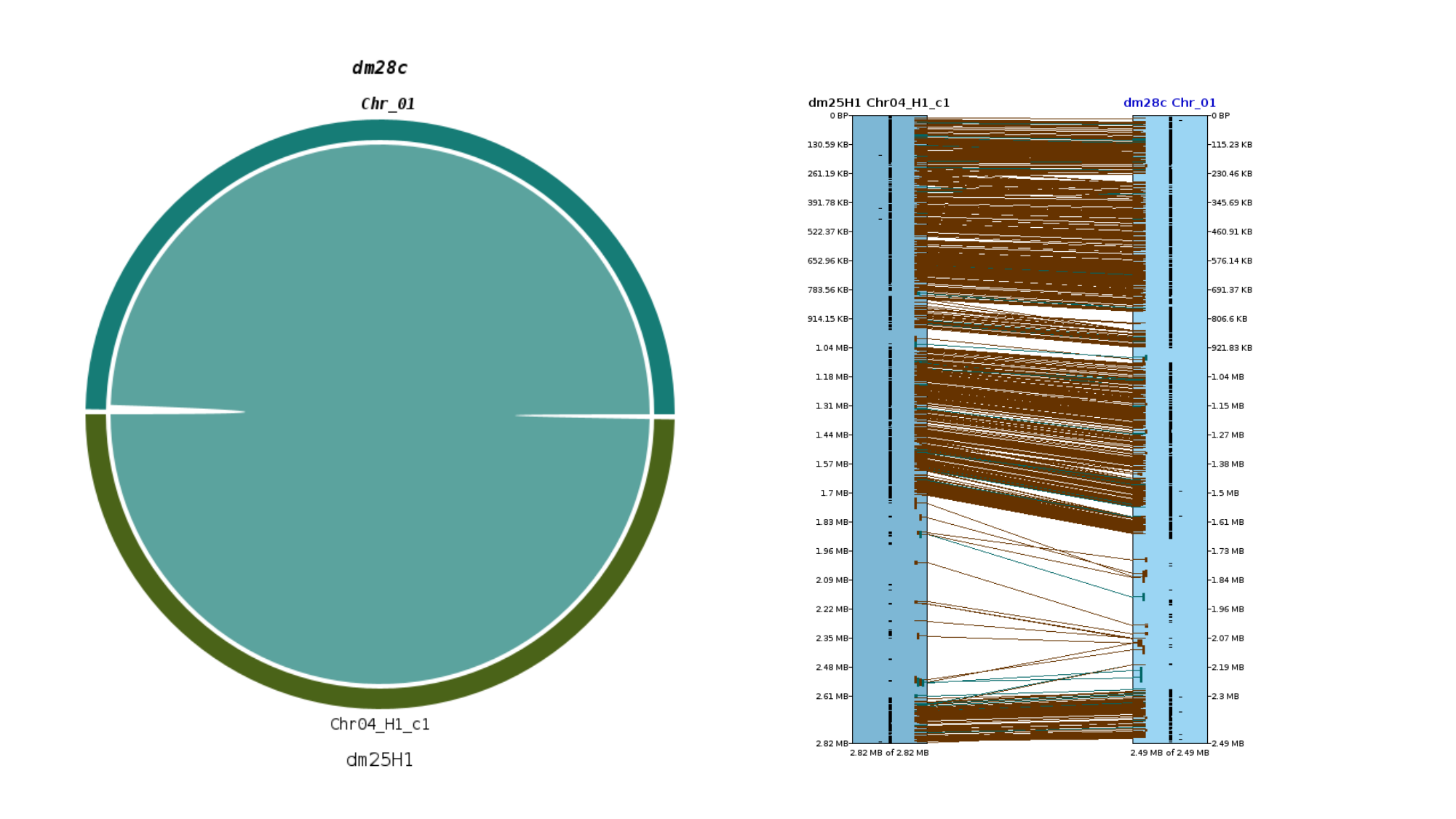

### Slide 3
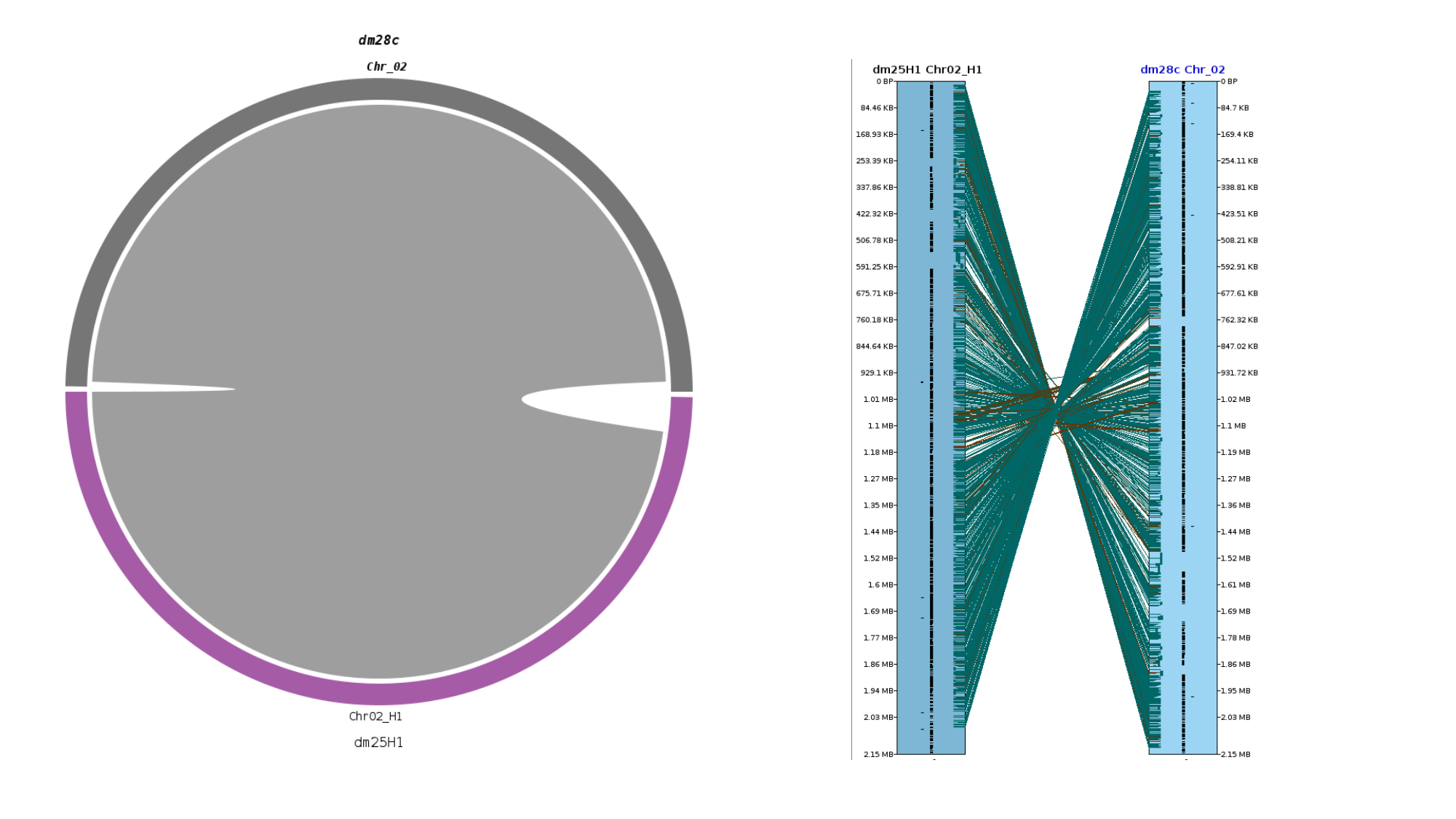

### Slide 4
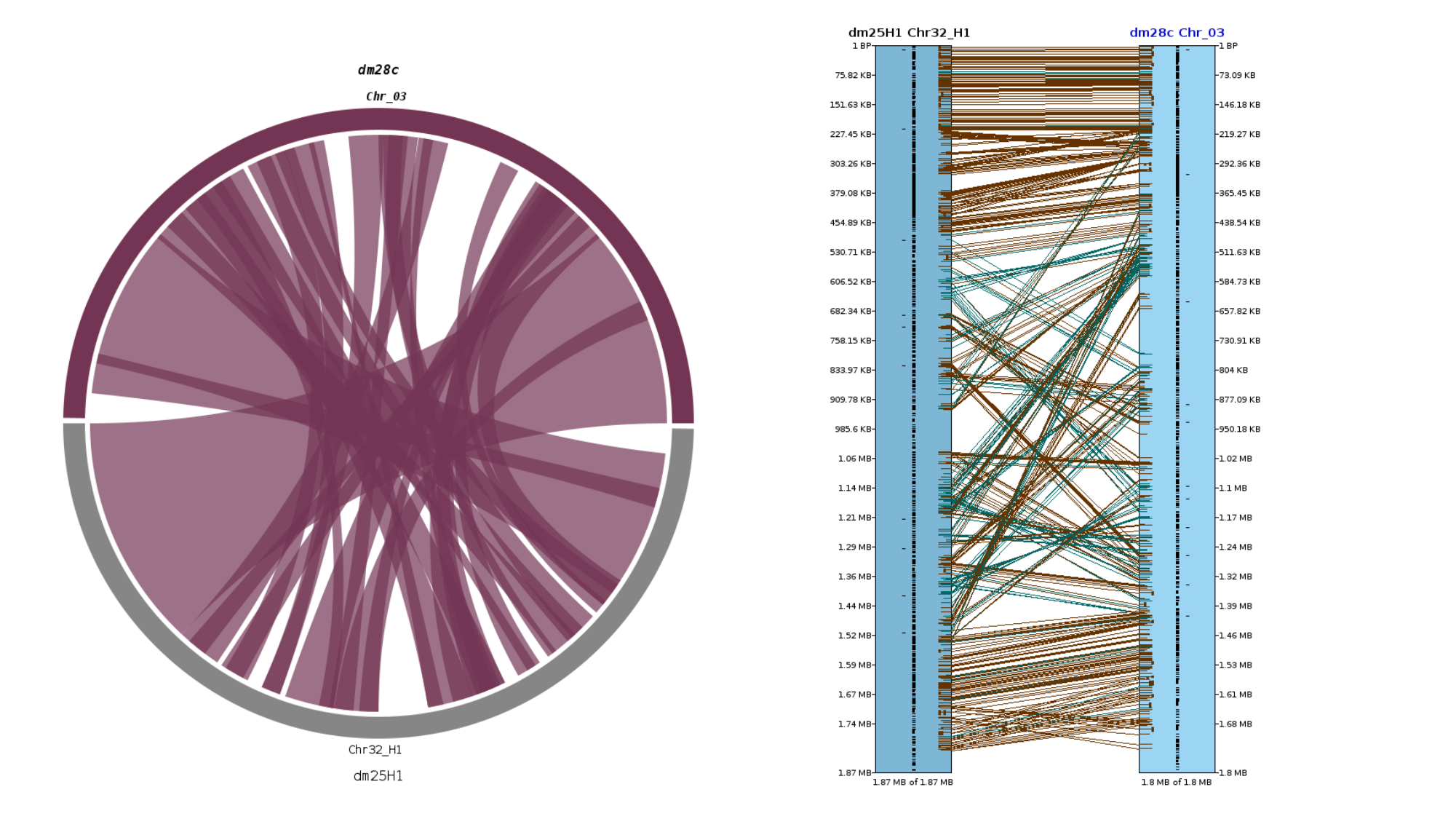

### Slide 5
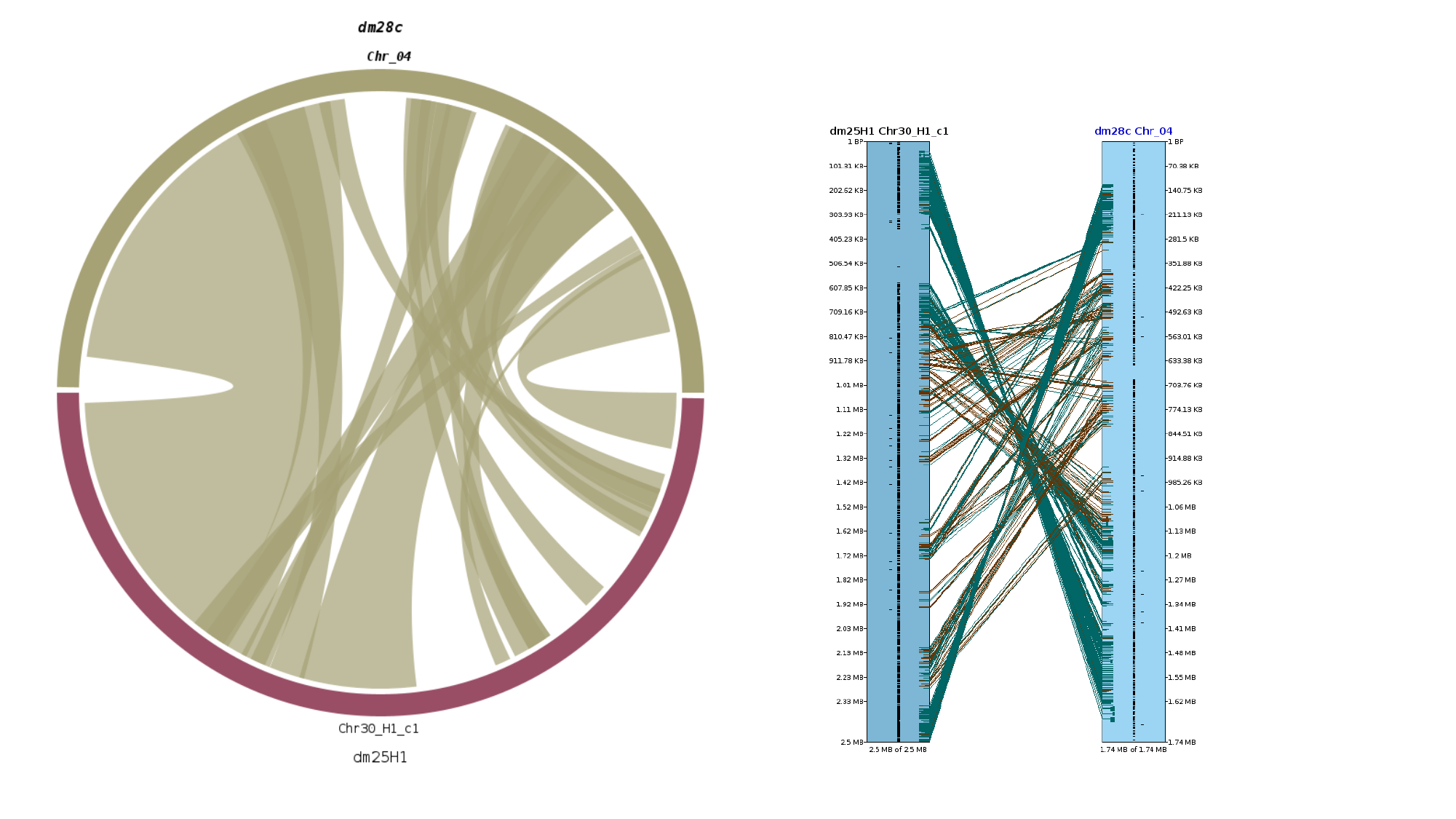

### Slide 6
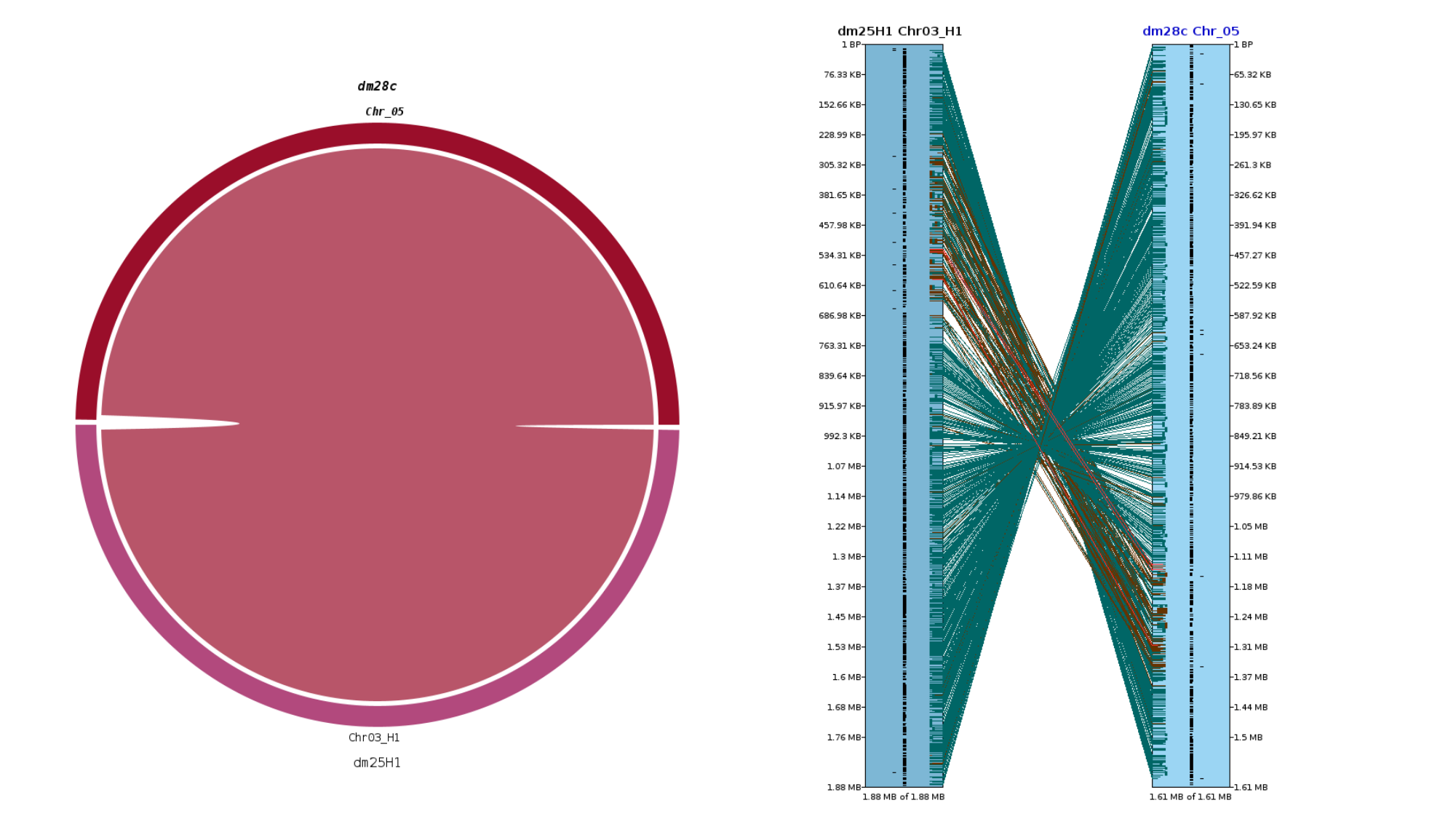

### Slide 7
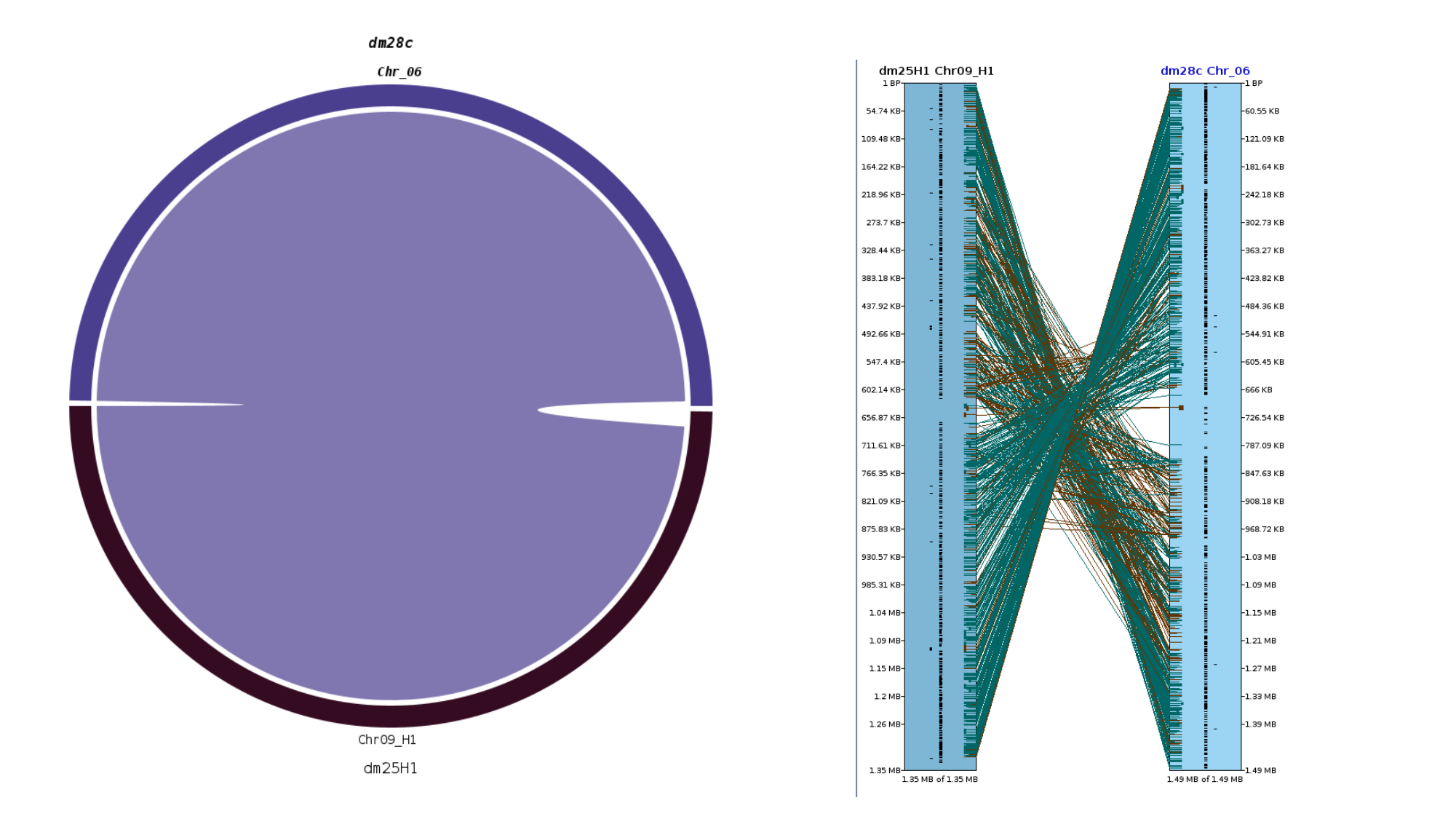

### Slide 8
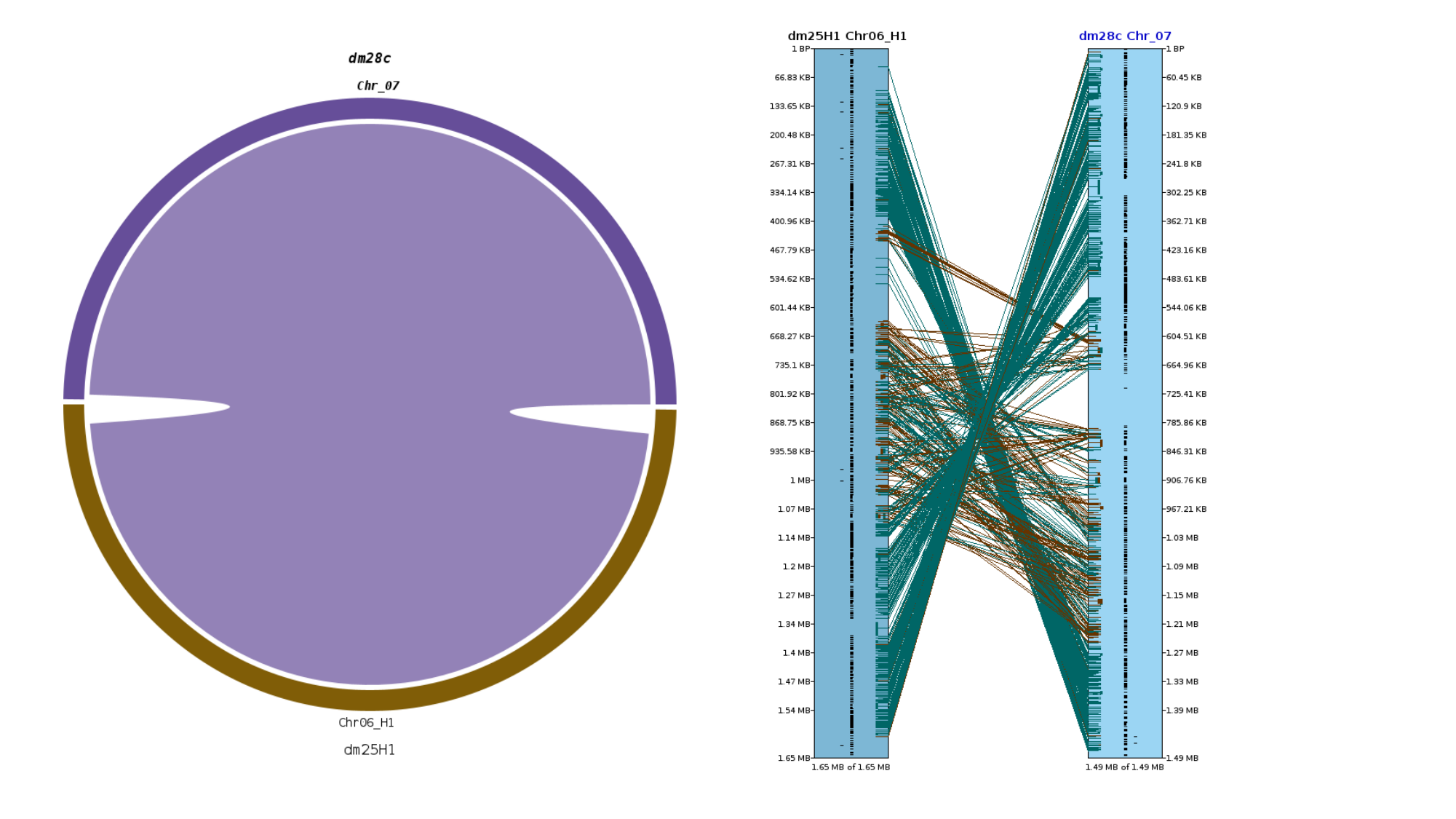

### Slide 9
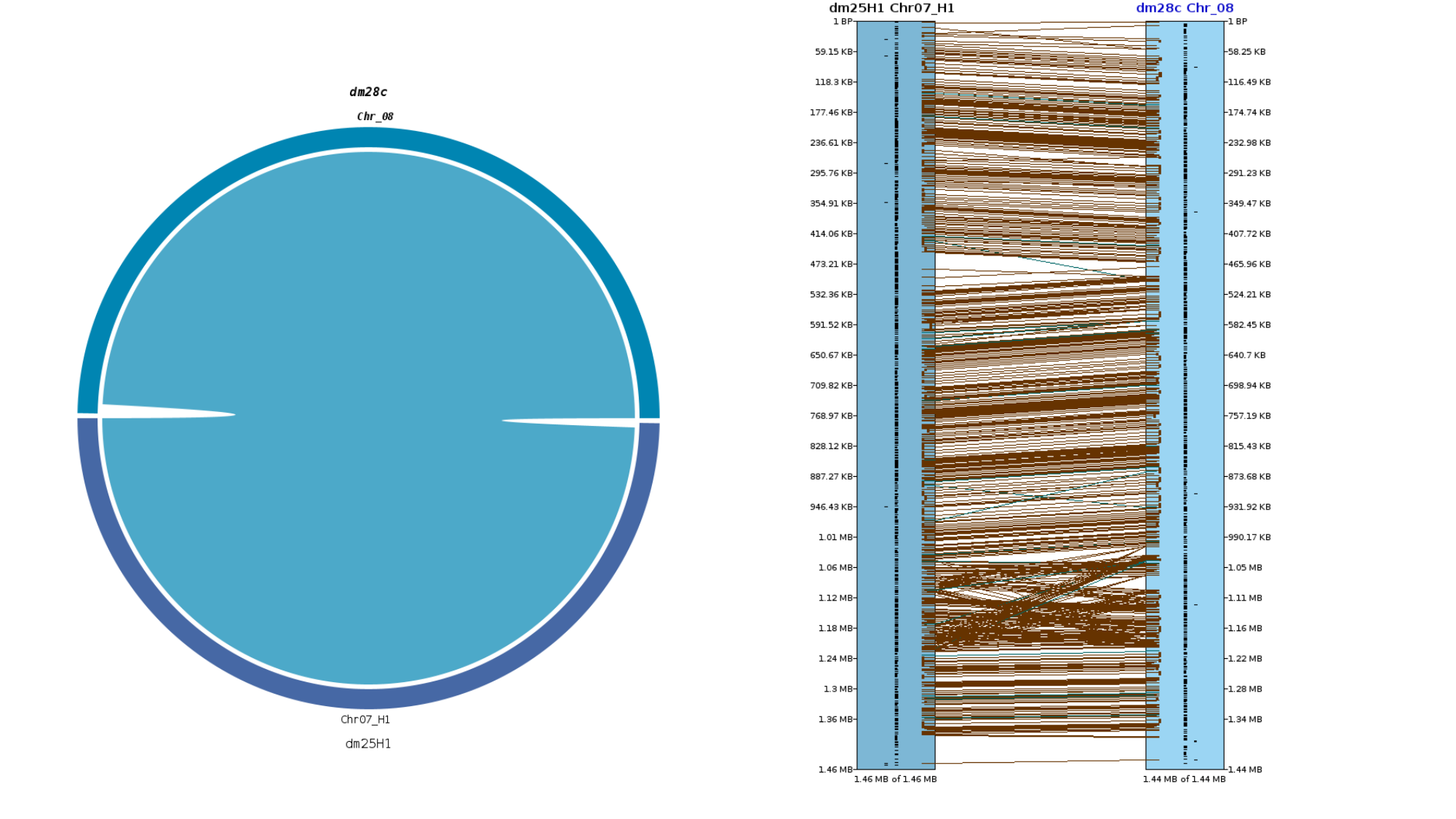

### Slide 10
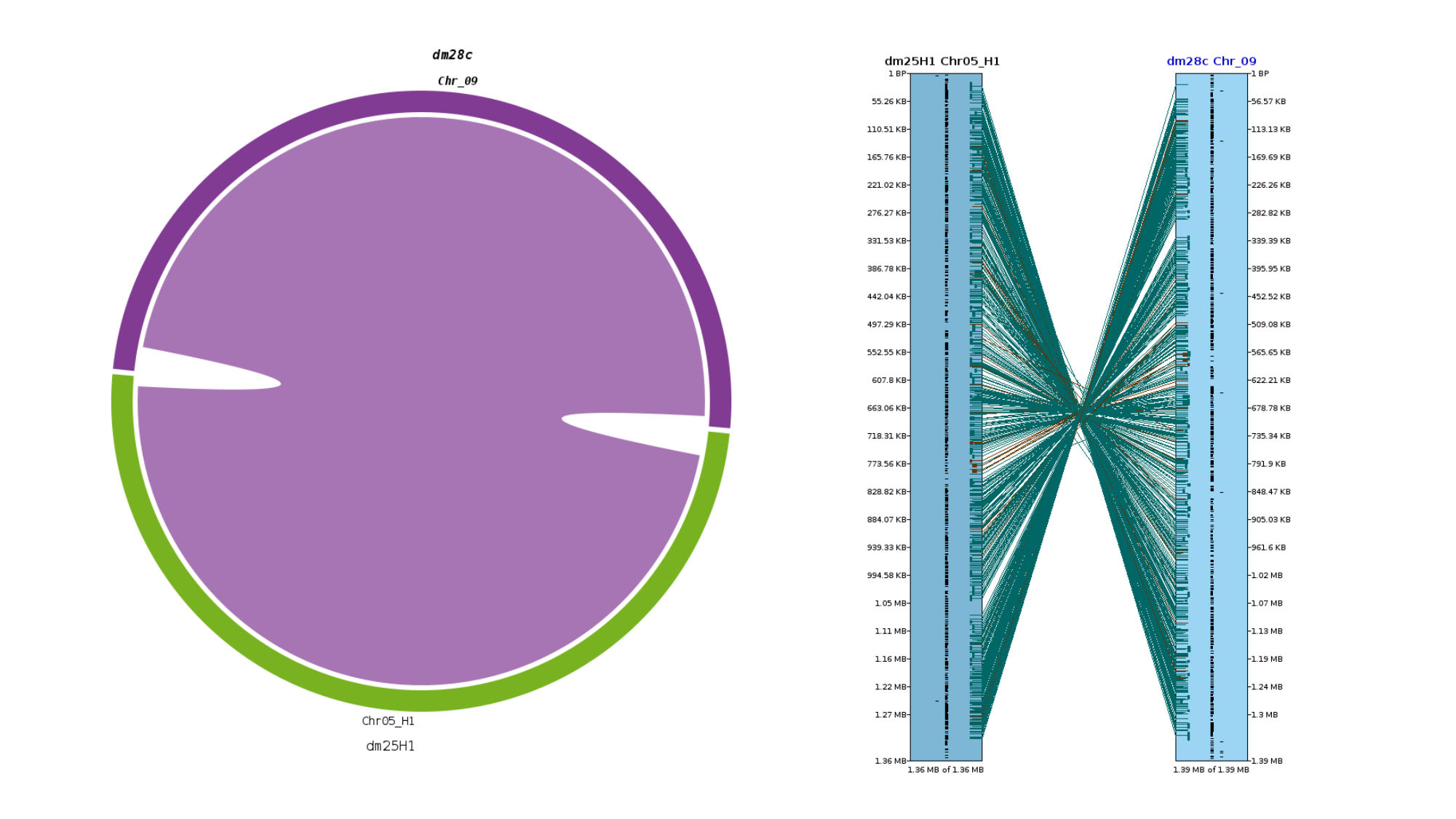

### Slide 11
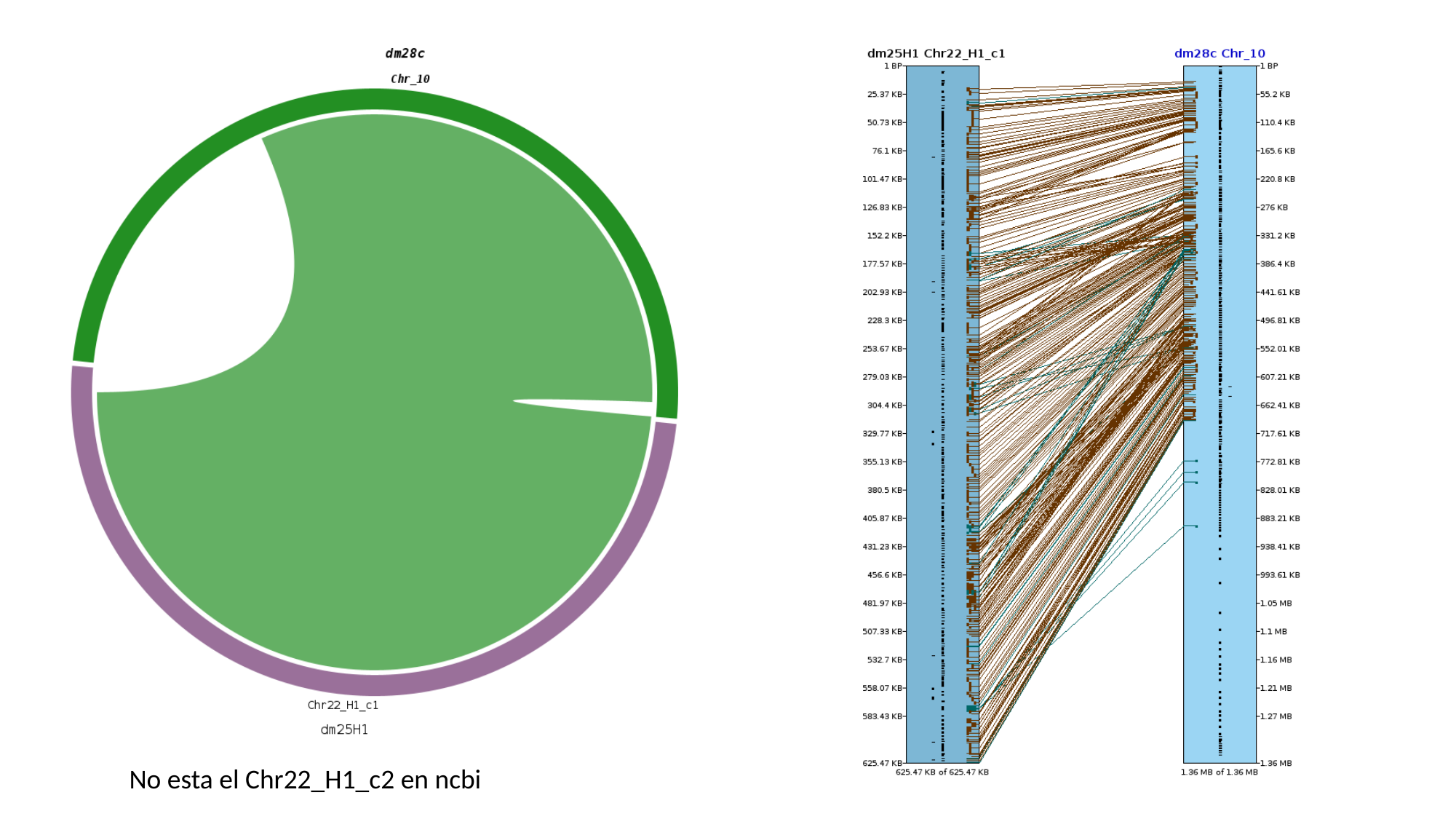

No esta el Chr22_H1_c2 en ncbi

### Slide 12
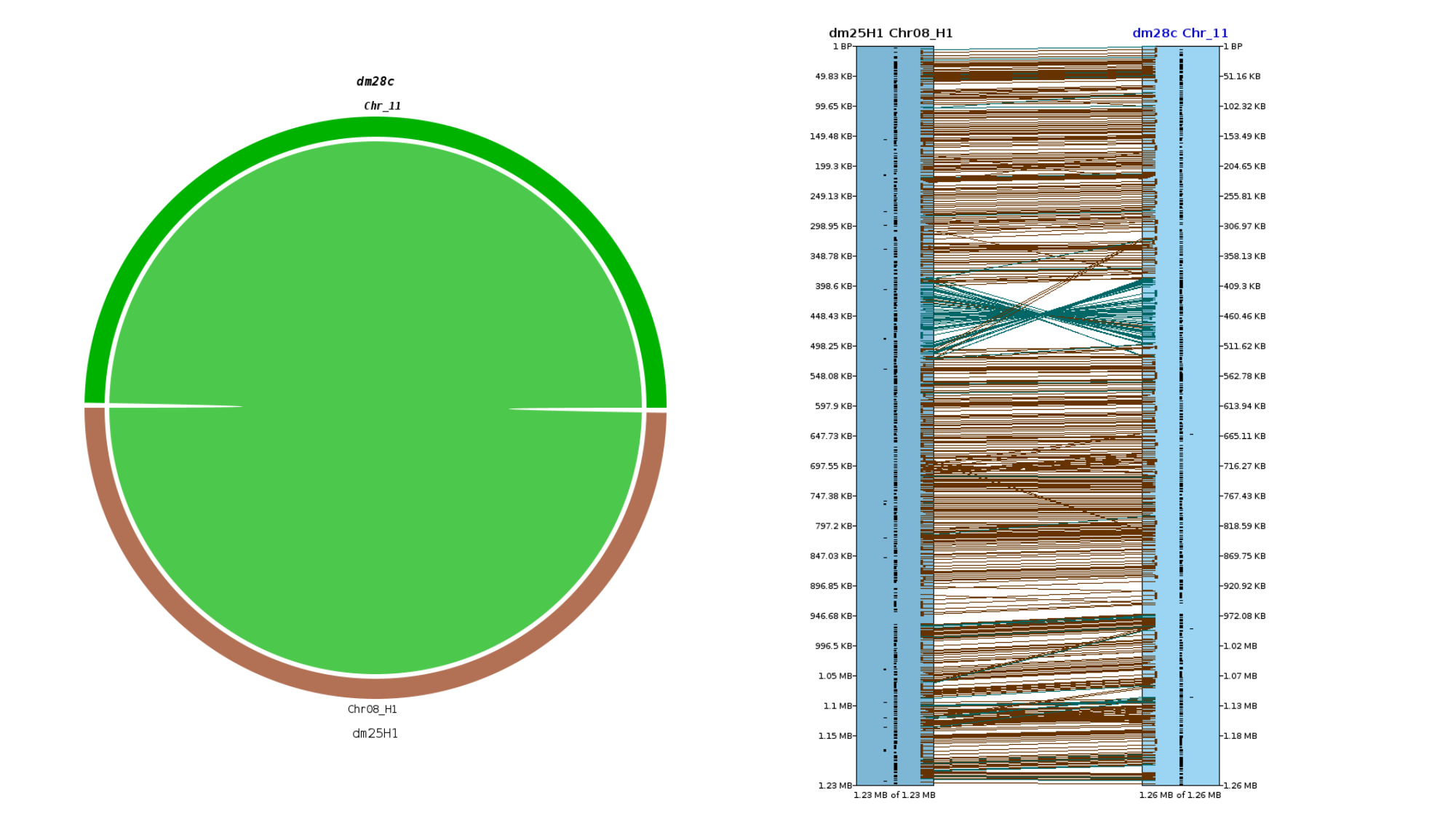

### Slide 13
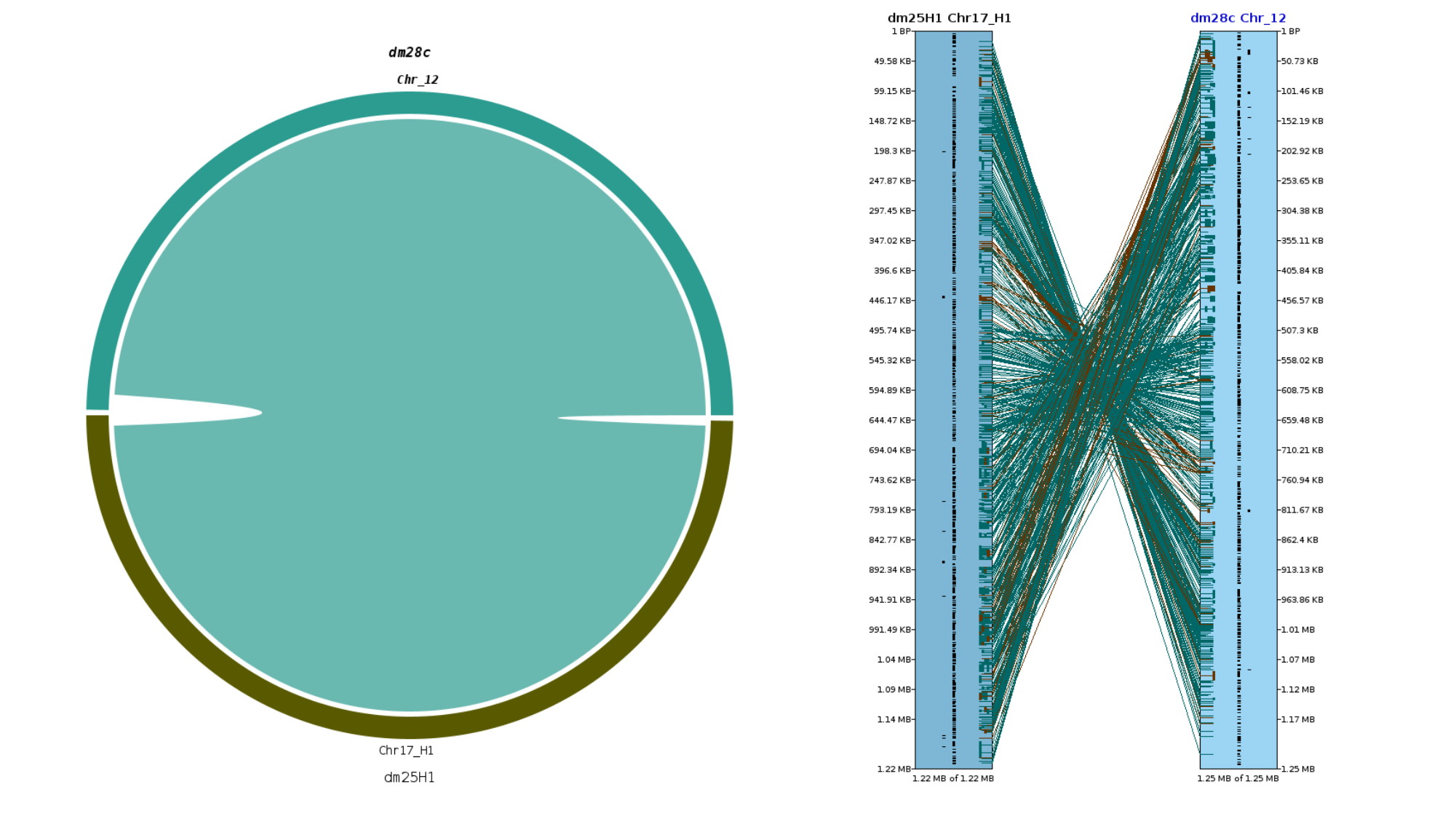

### Slide 14
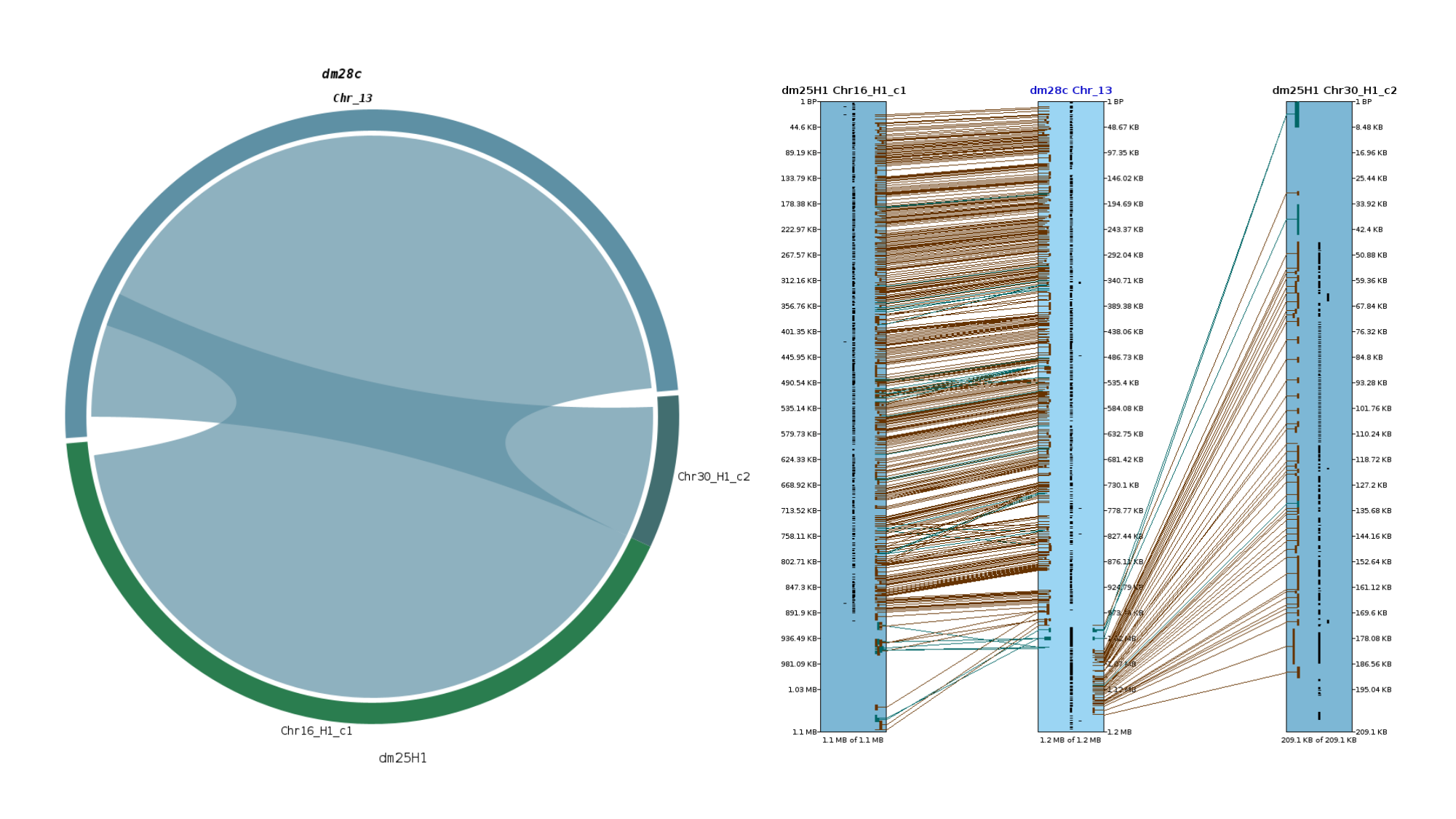

### Slide 15
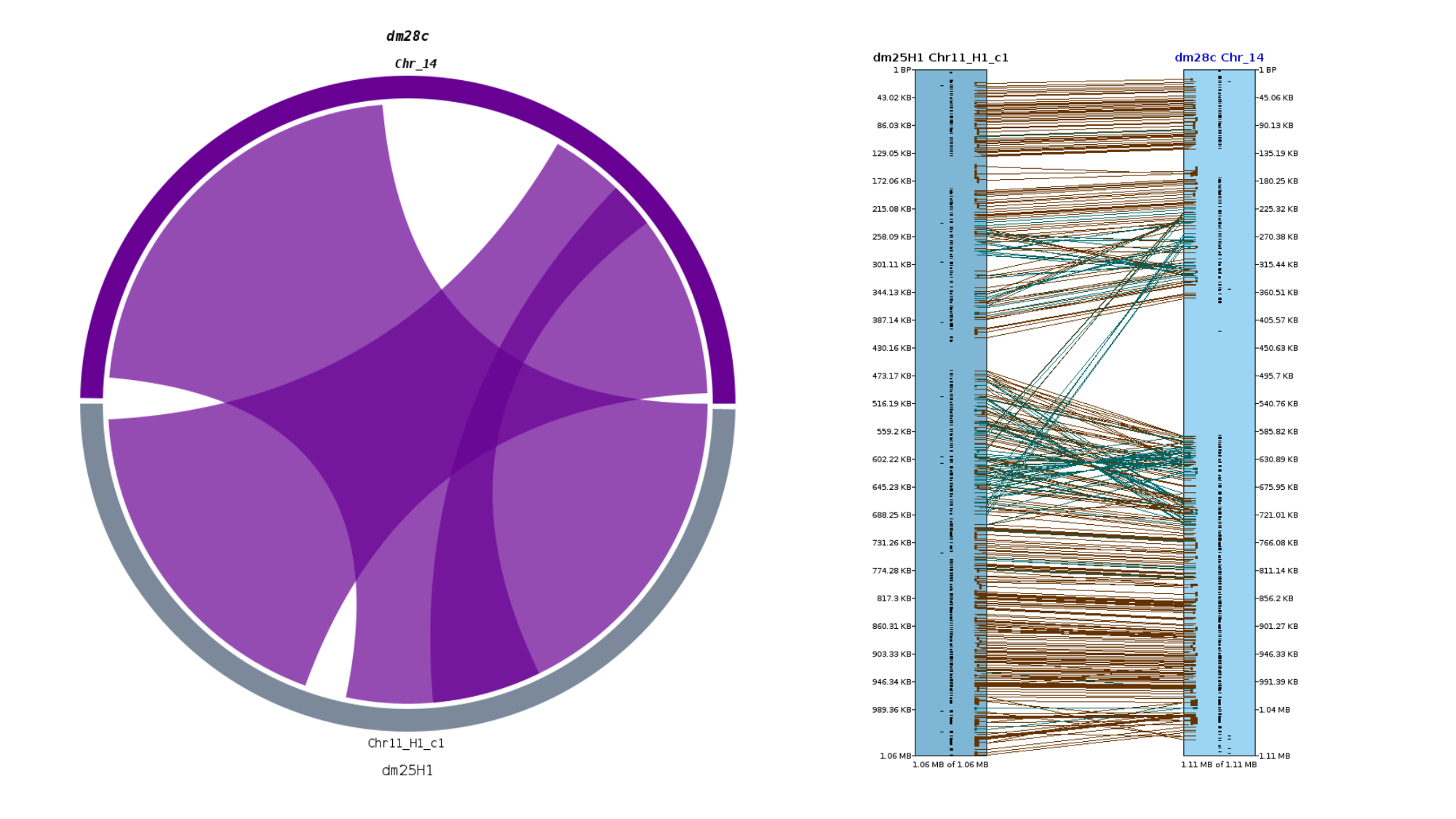

### Slide 16
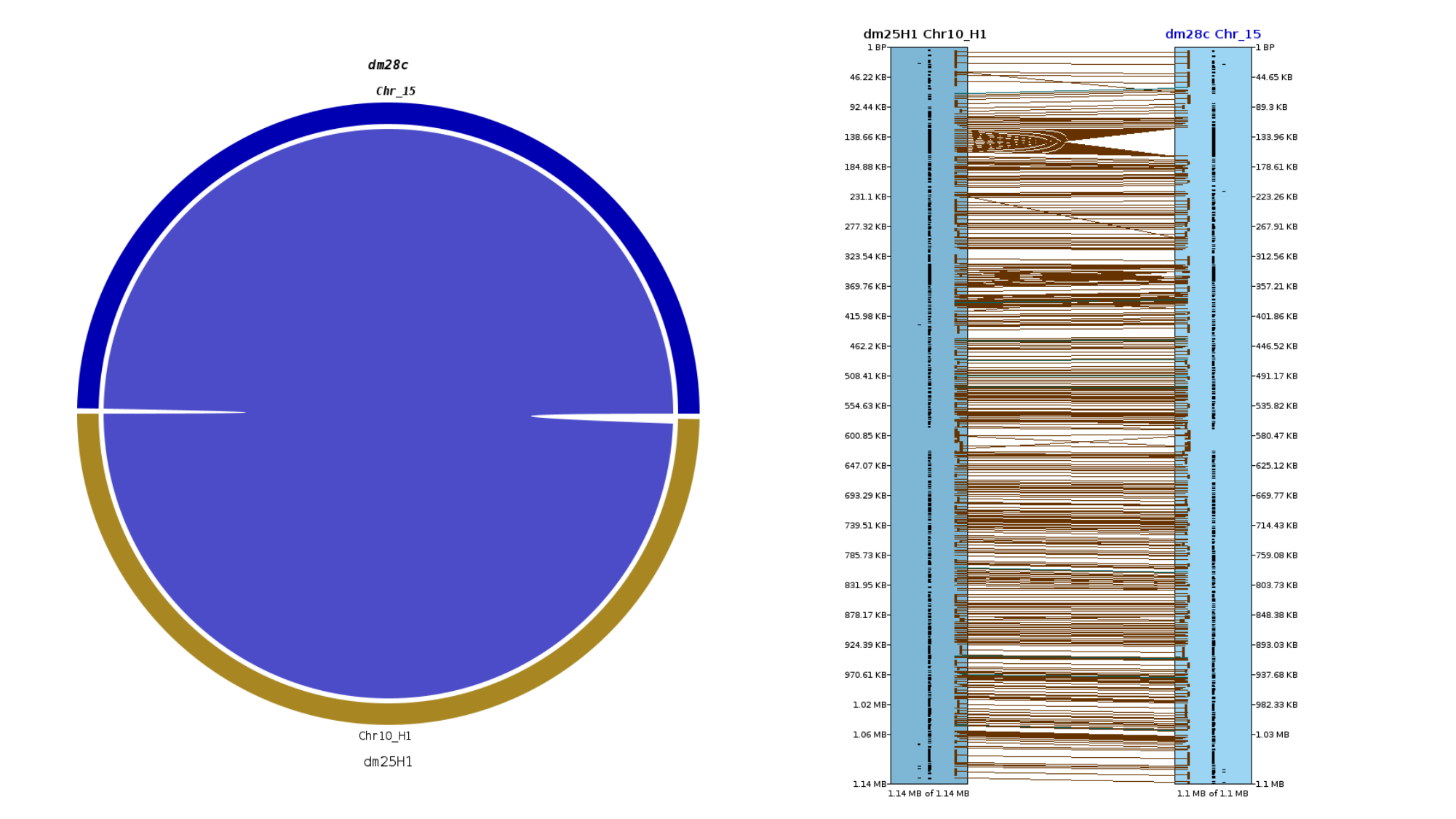

### Slide 17
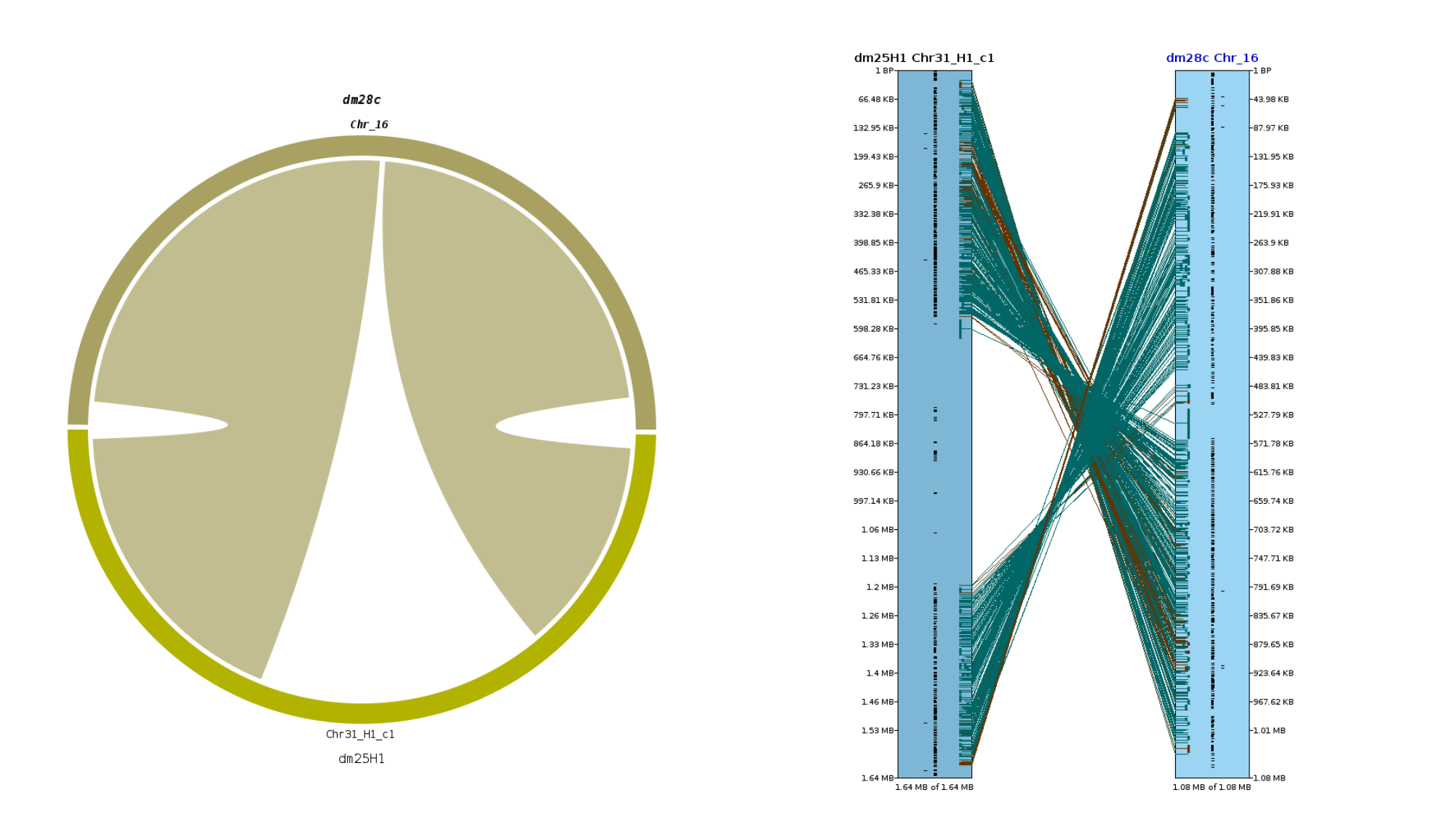

### Slide 18
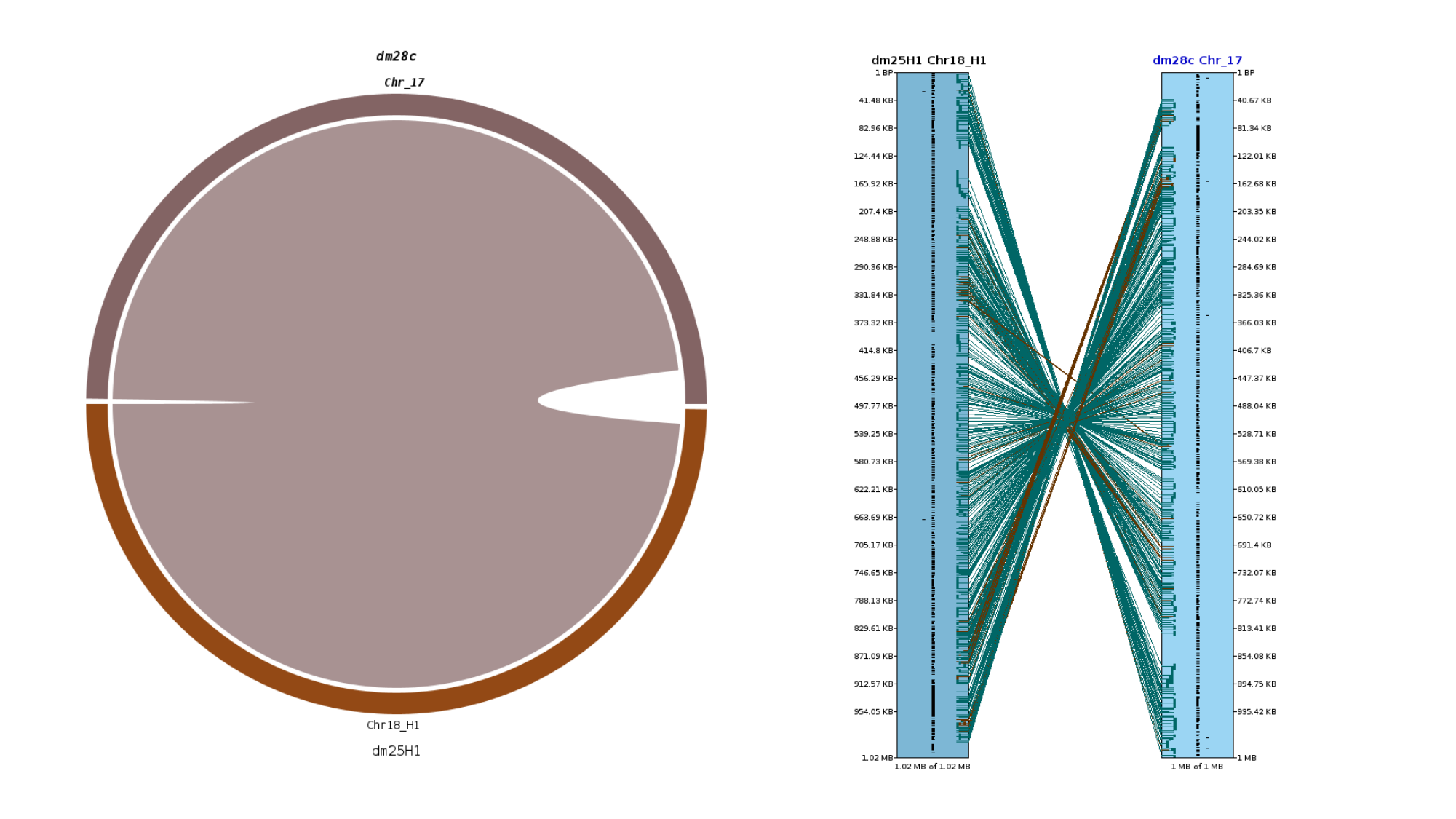

### Slide 19
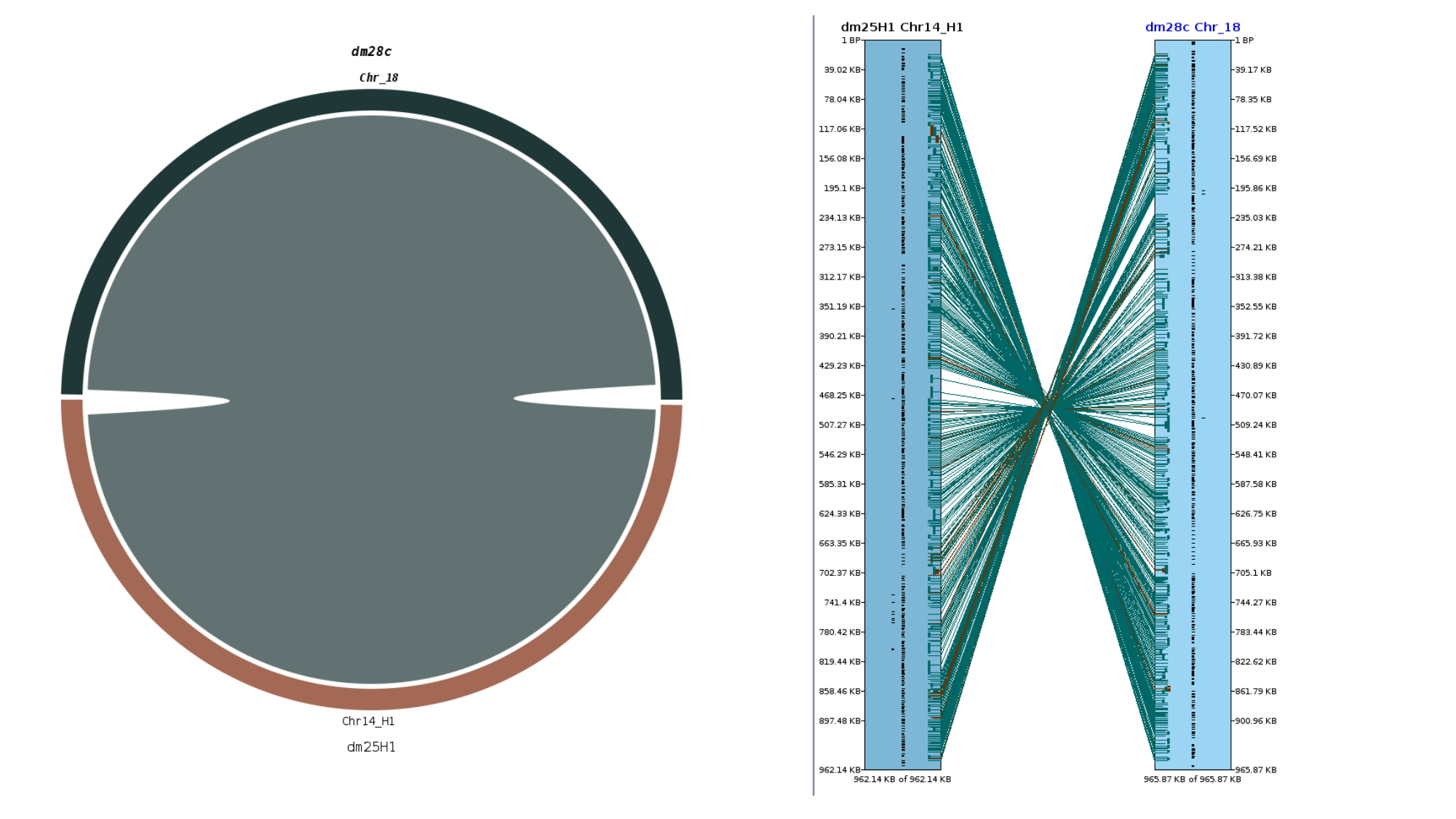

### Slide 20
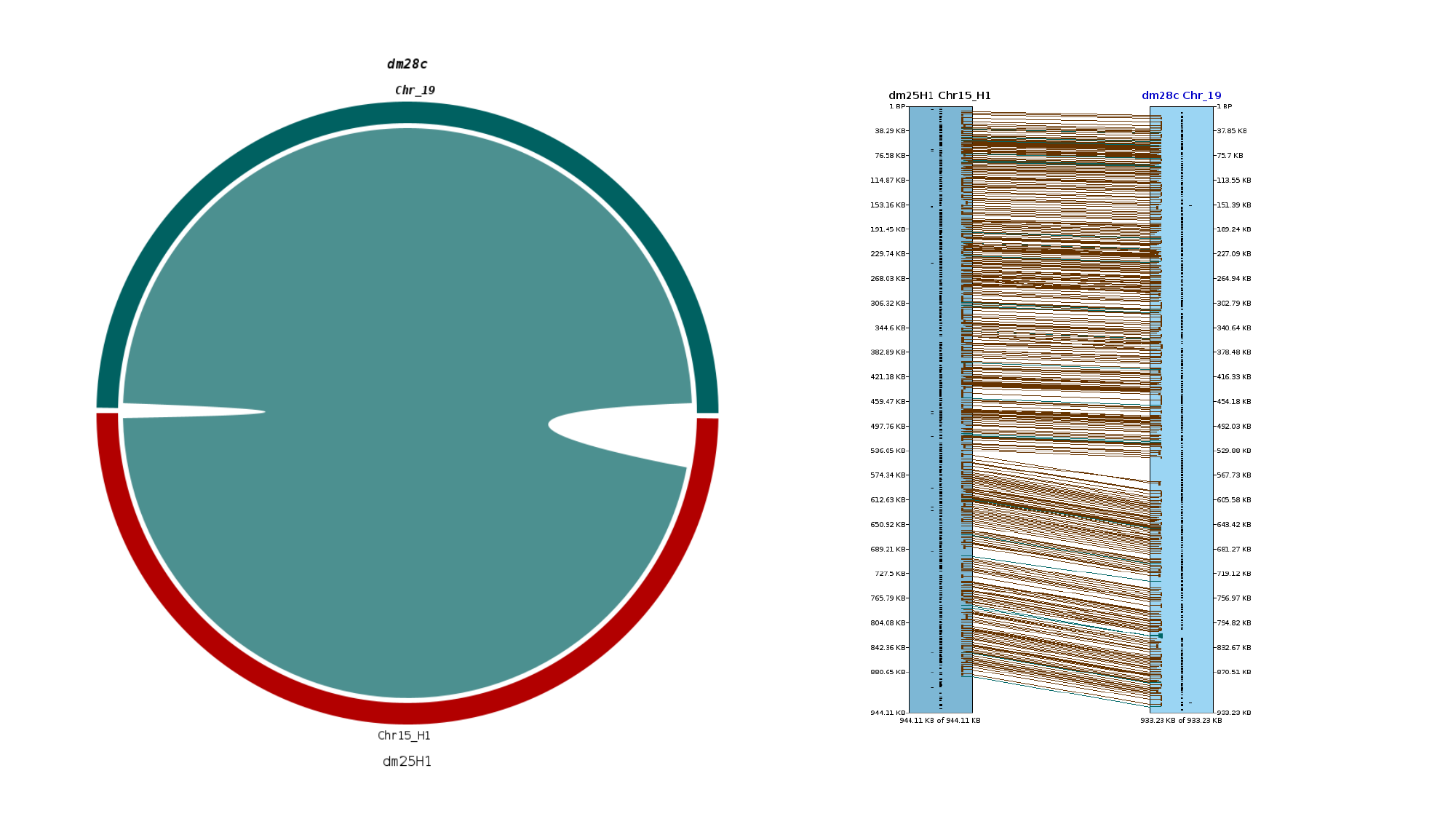

### Slide 21
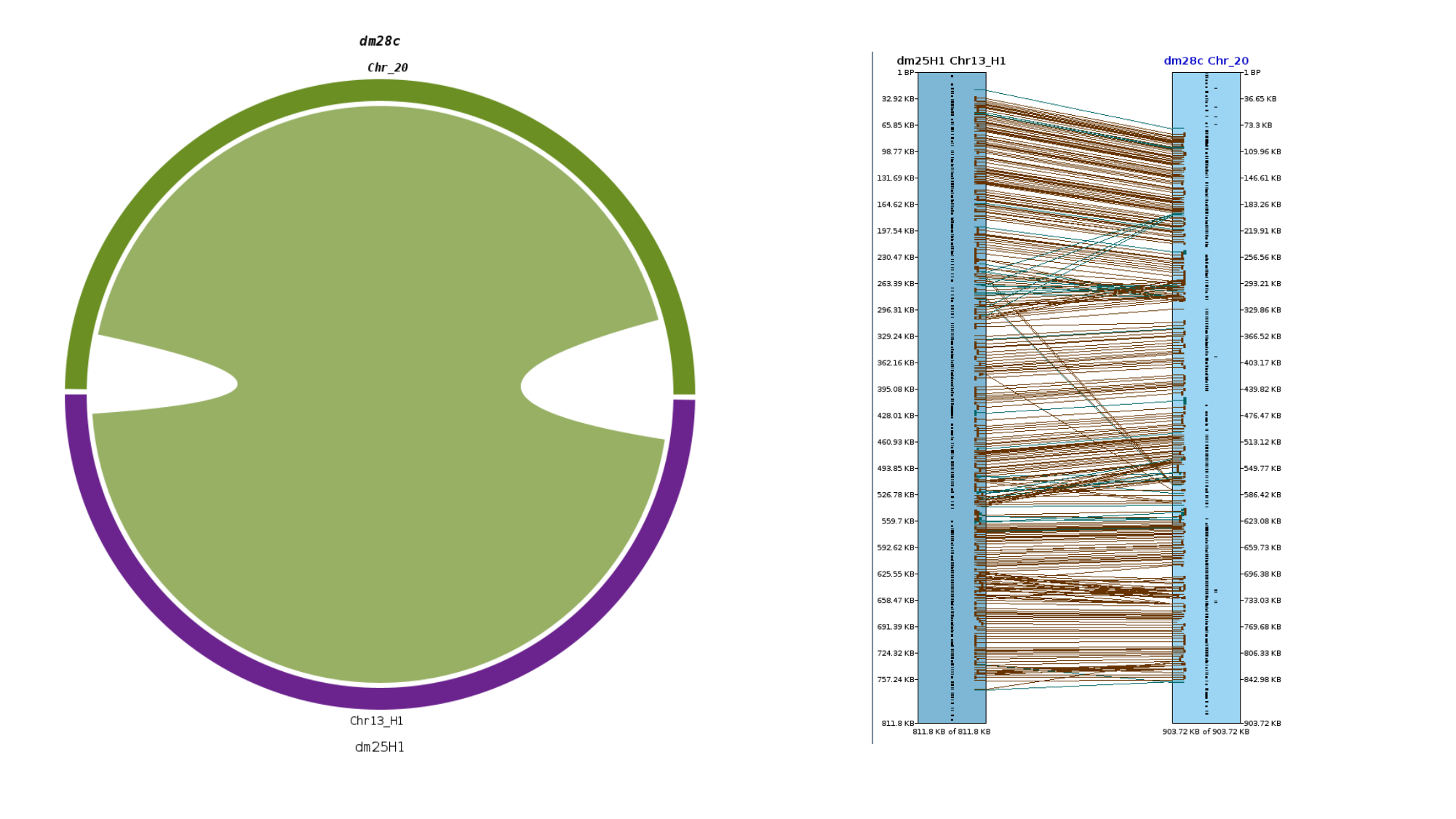

### Slide 22
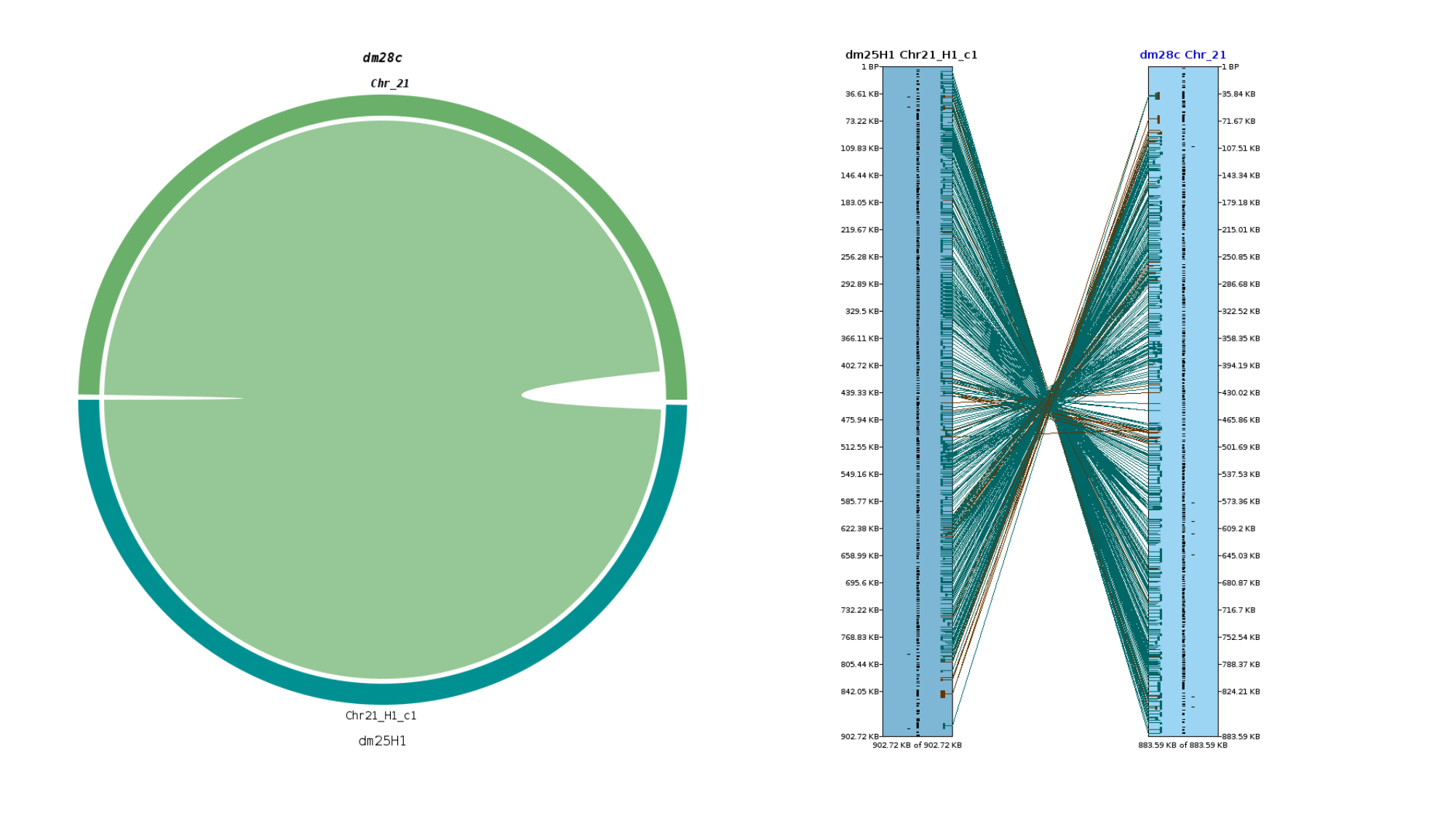

### Slide 23
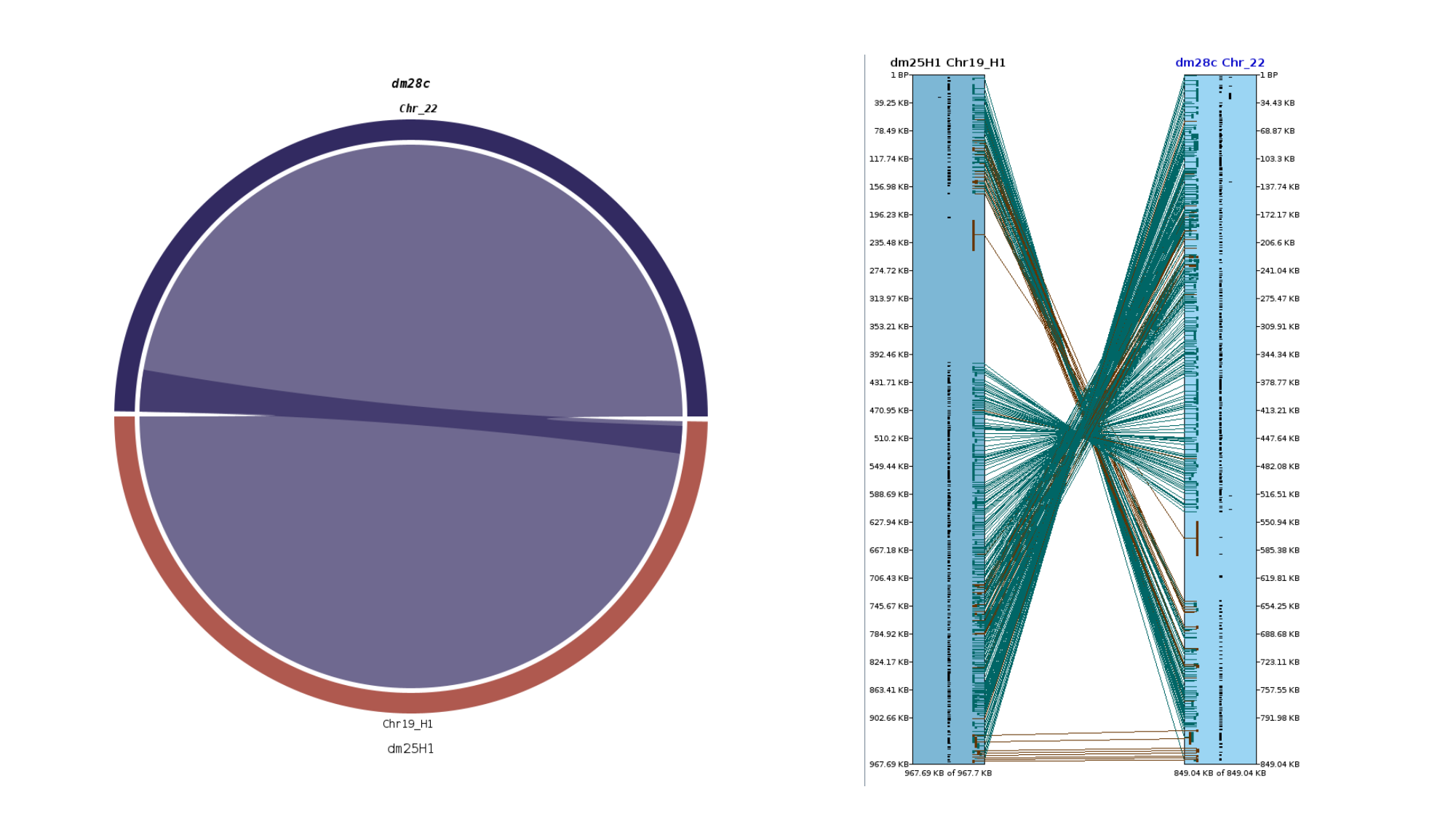

### Slide 24
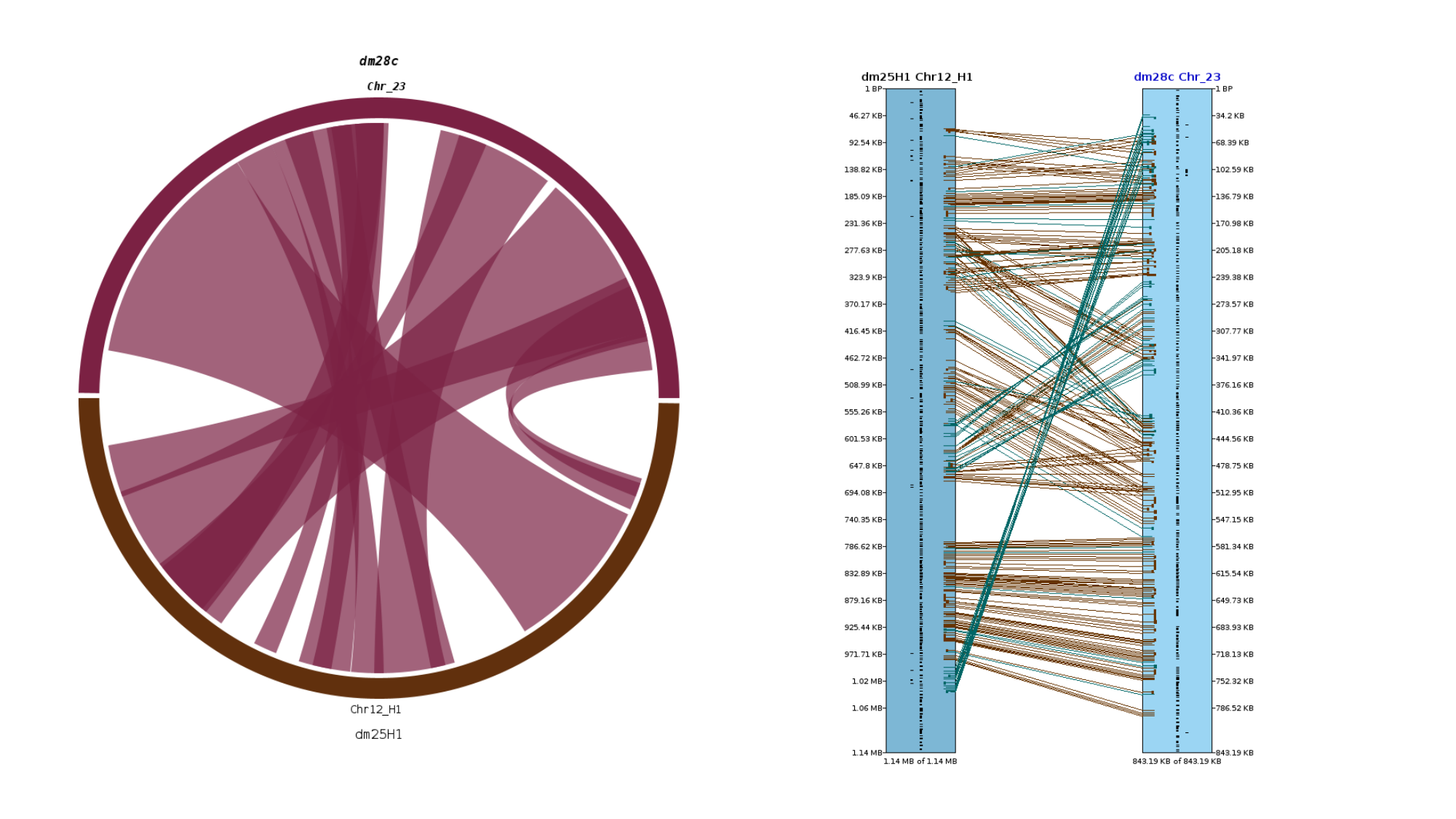

### Slide 25
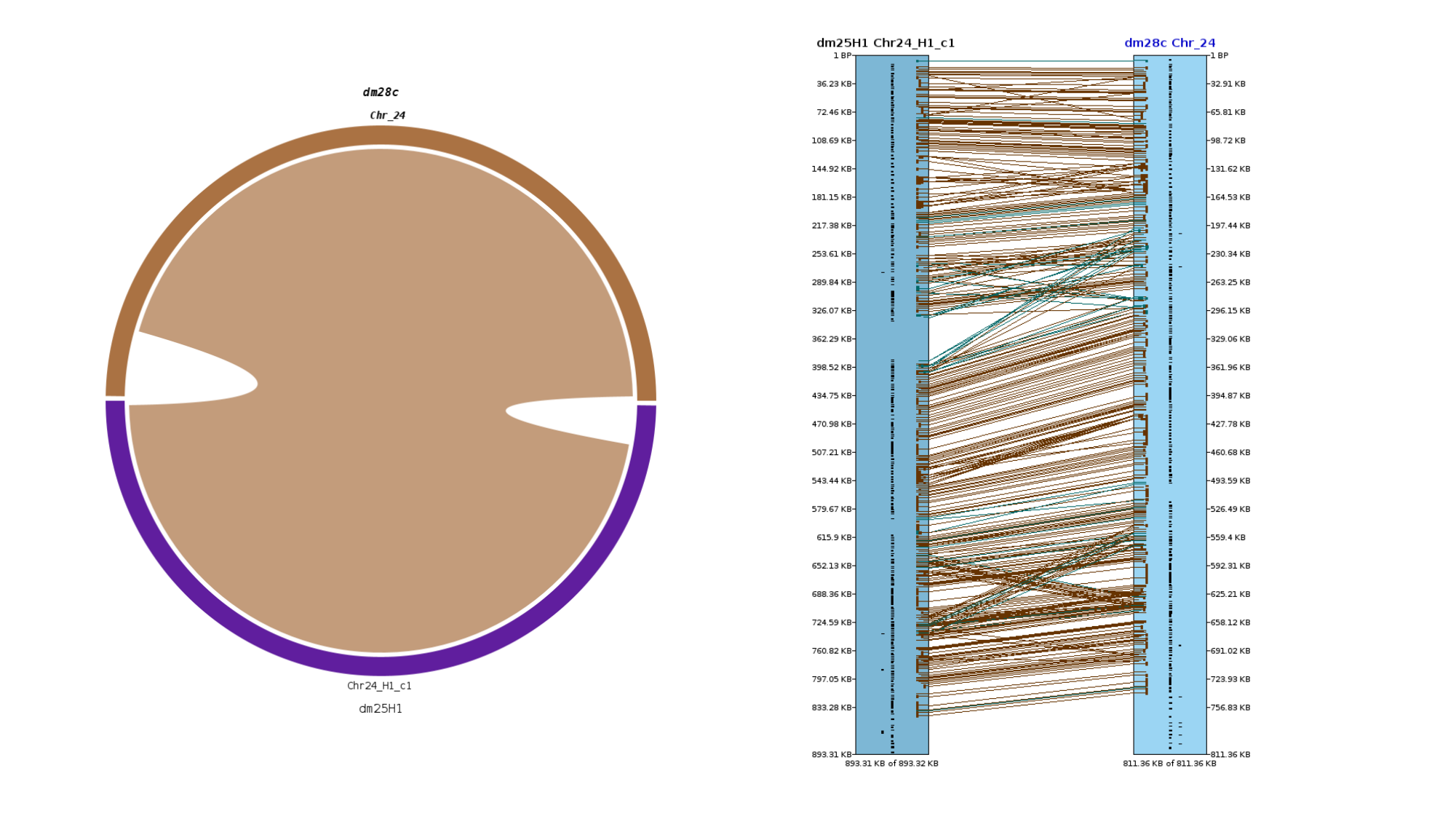

### Slide 26
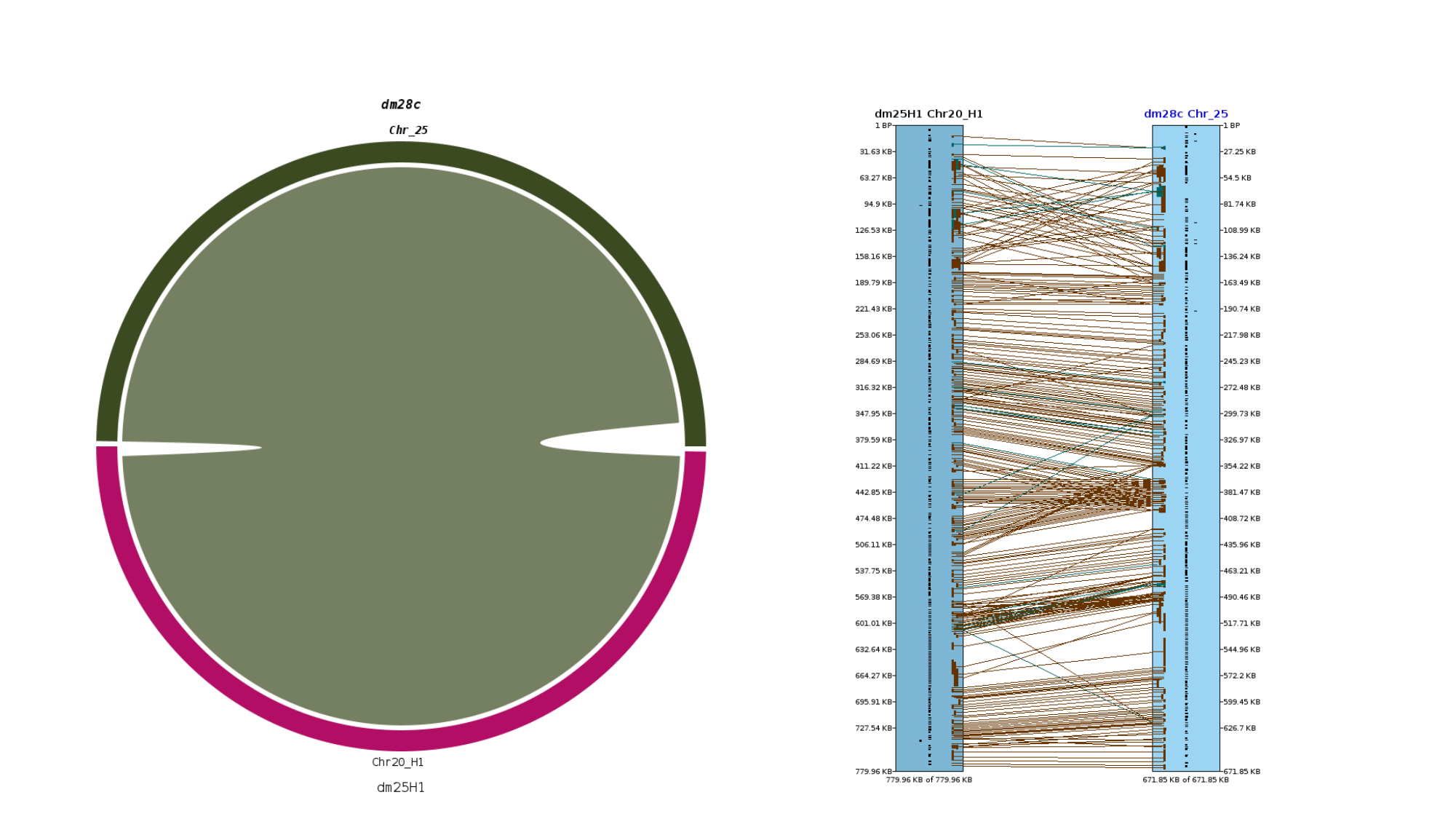

### Slide 27
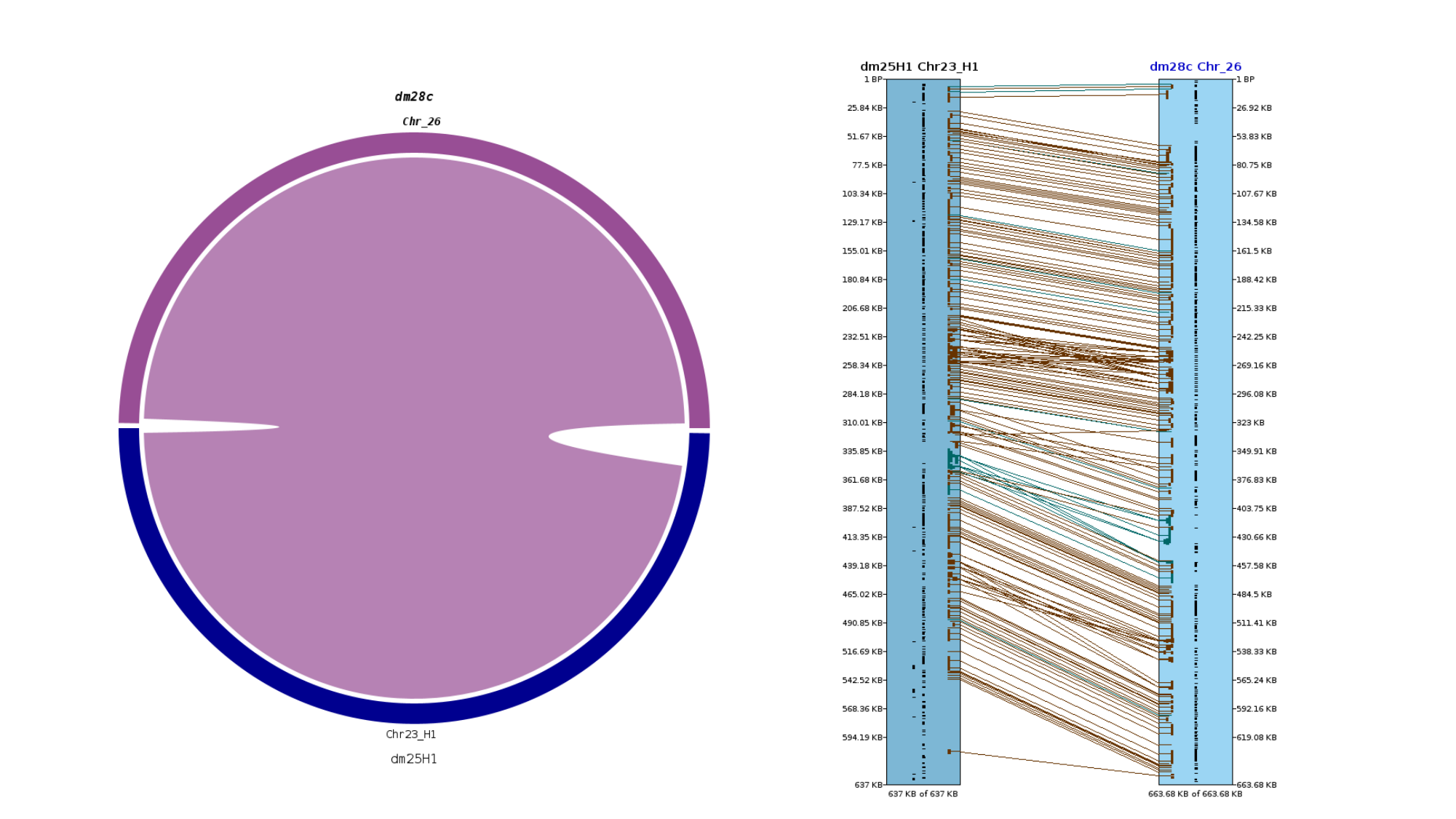

### Slide 28
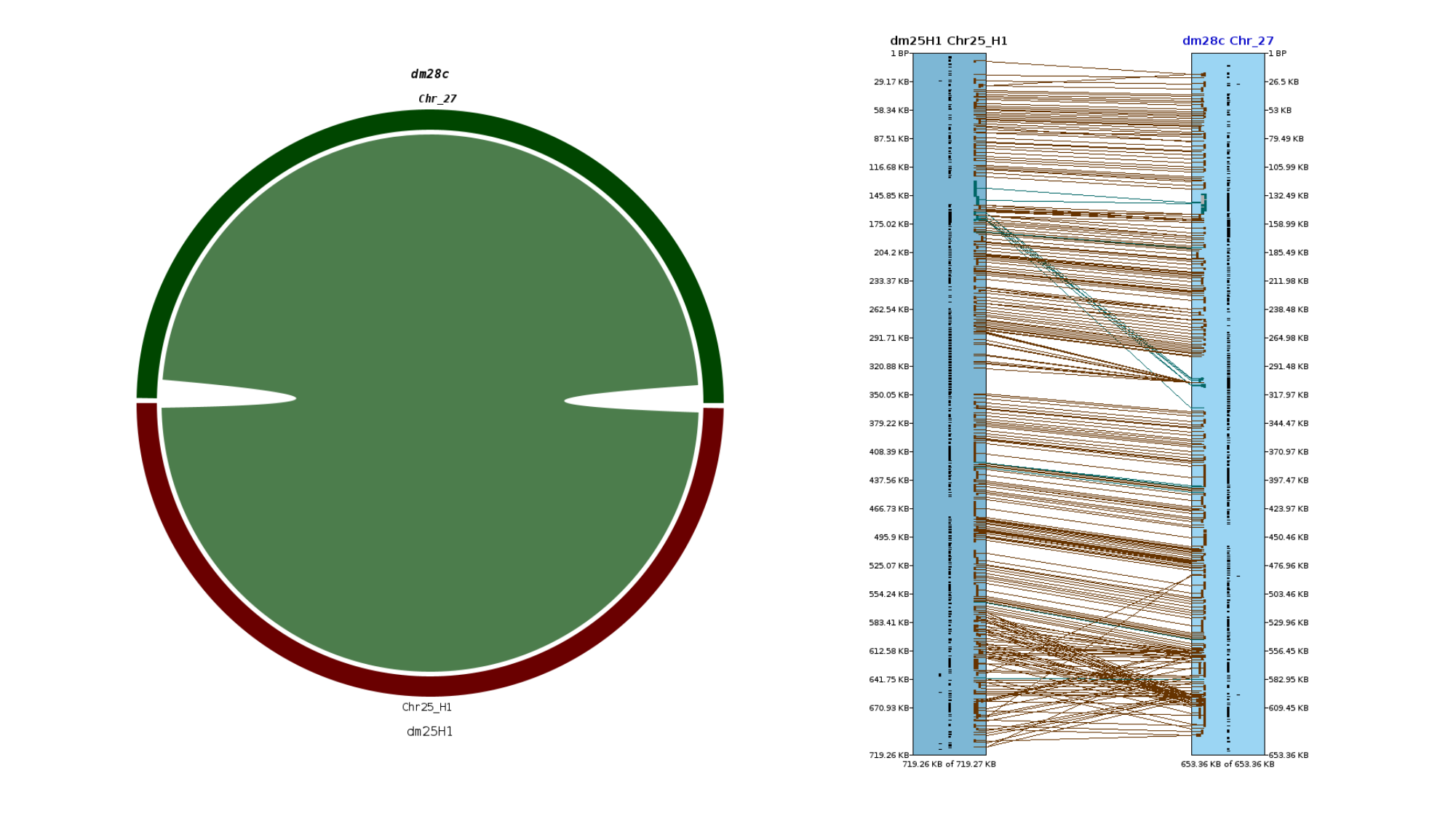

### Slide 29
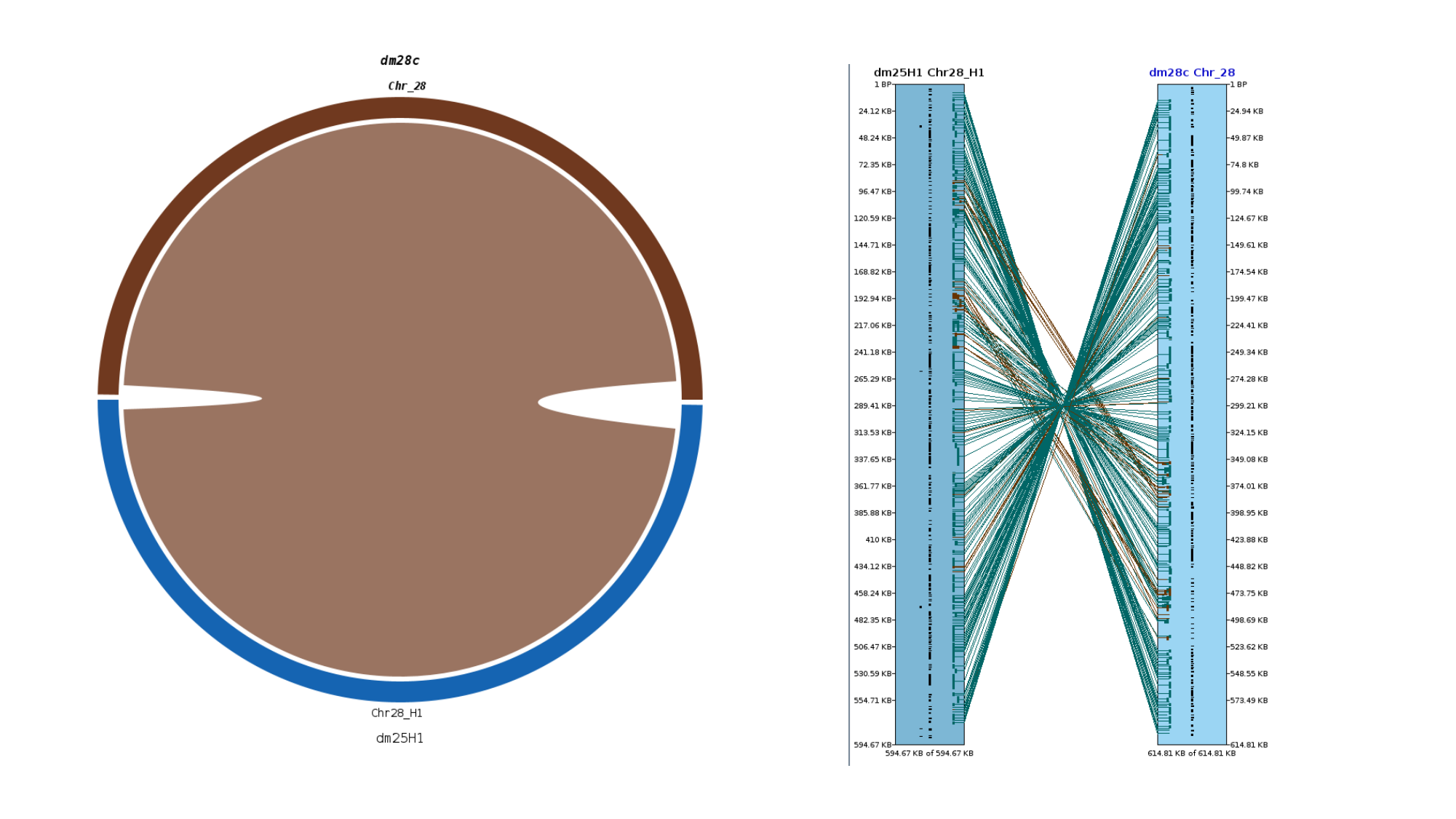

### Slide 30
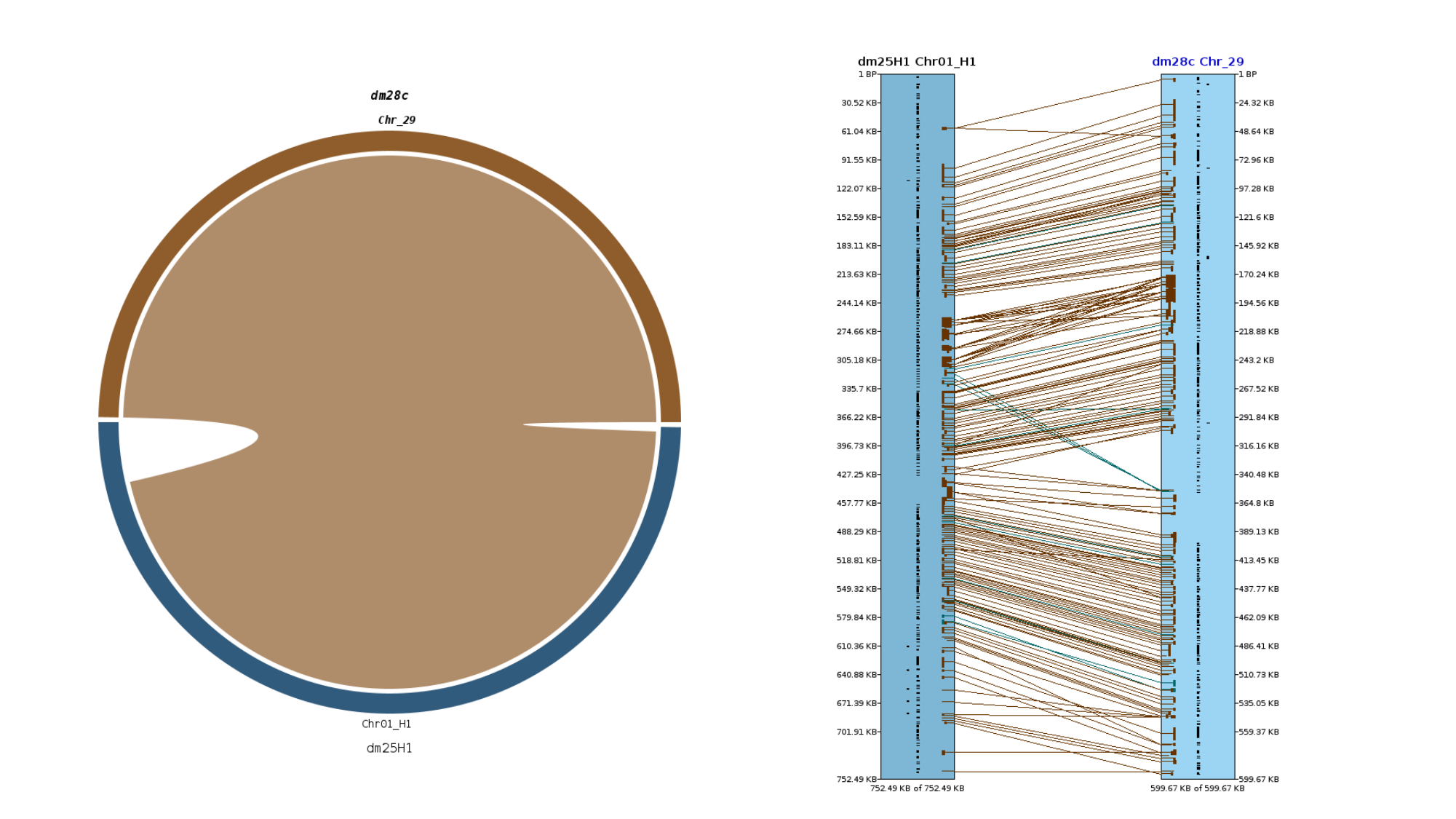

### Slide 31
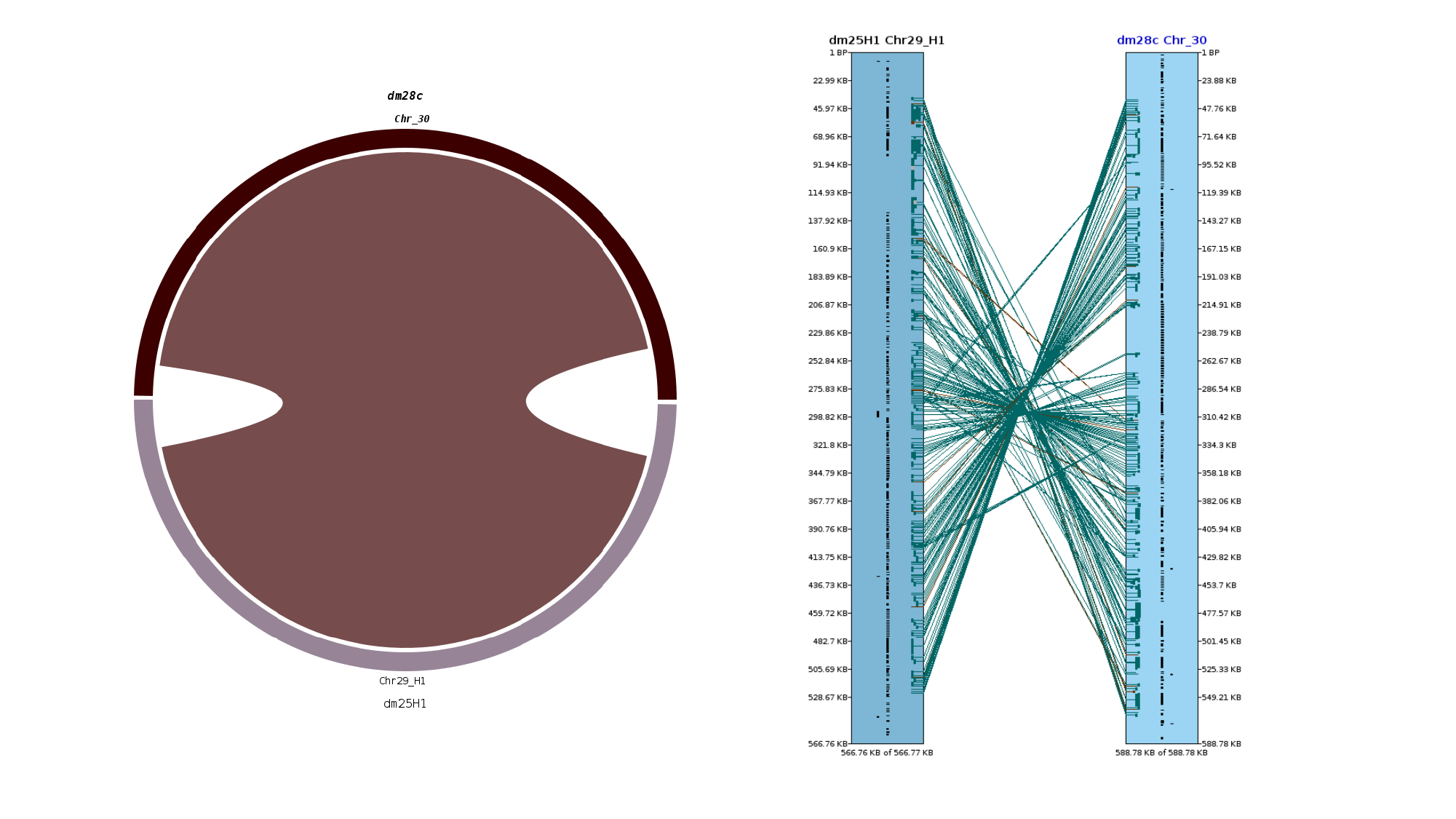

### Slide 32
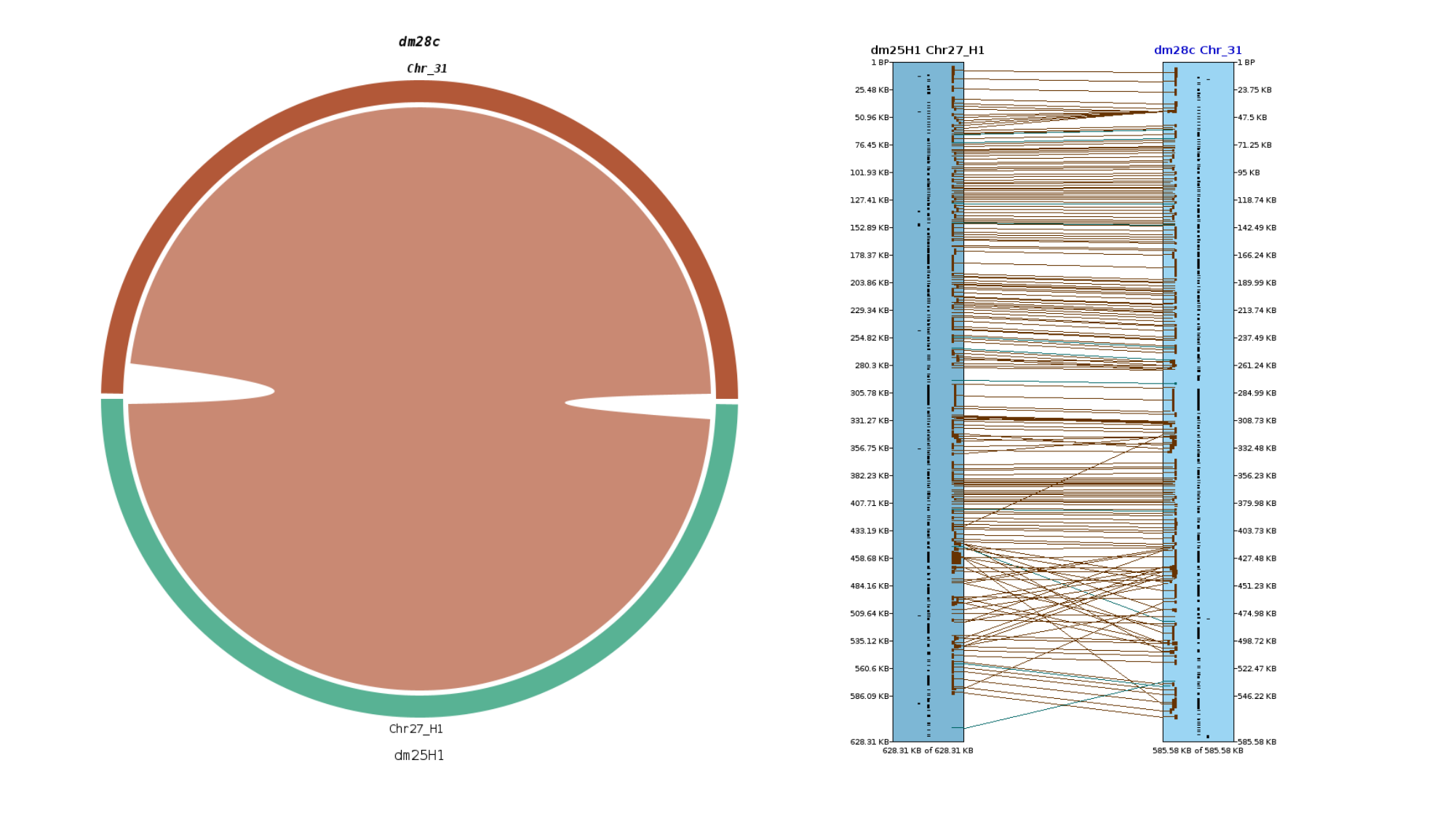

### Slide 33
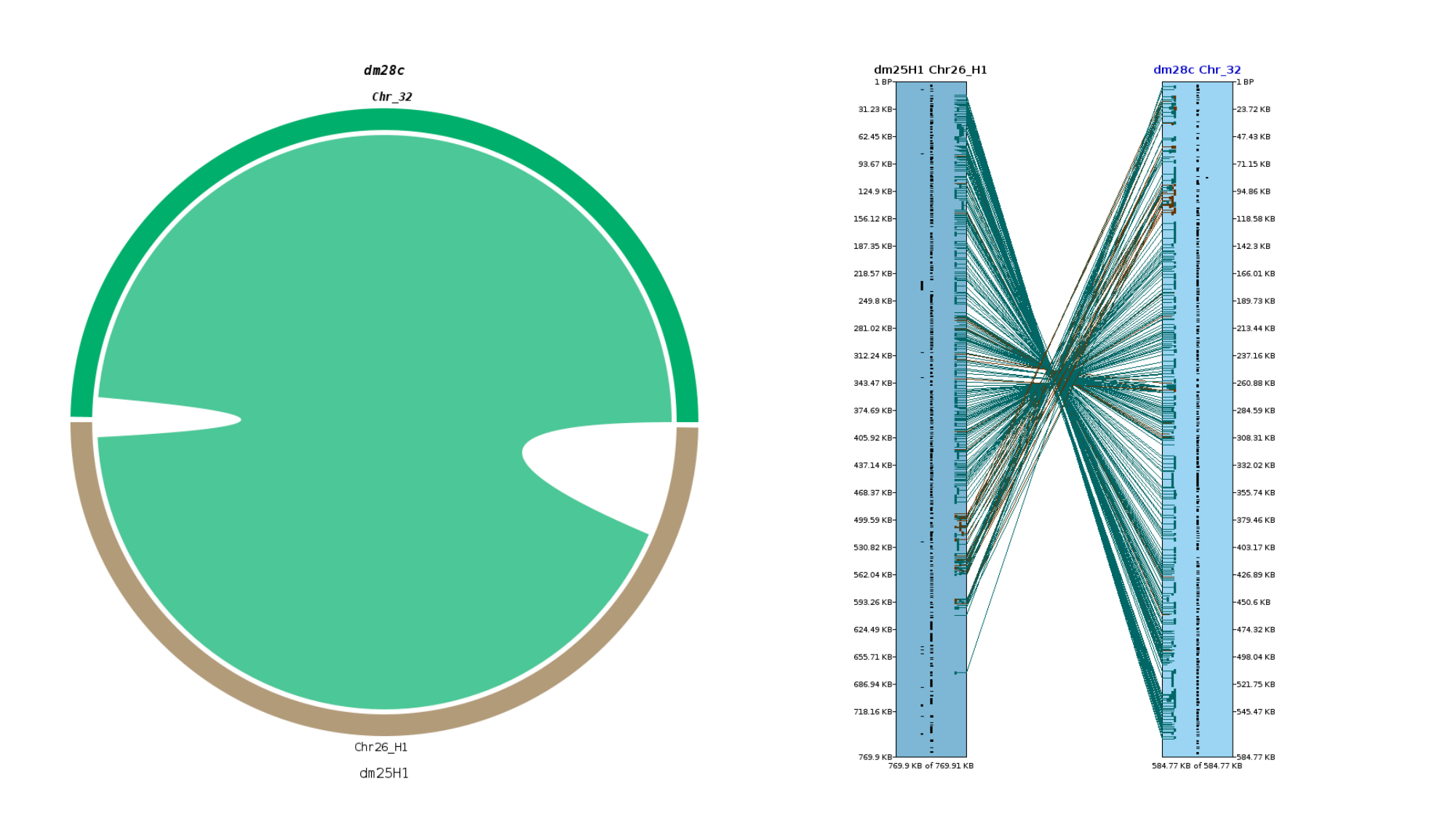
