## Supplementary material for "*Trypanosoma cruzi* has 32 Chromosomes: A Telomere-to-Telomere Assembly Defines Its Karyotype": SupFigure5_ChromosomesStructure.pptx

### Slide 1
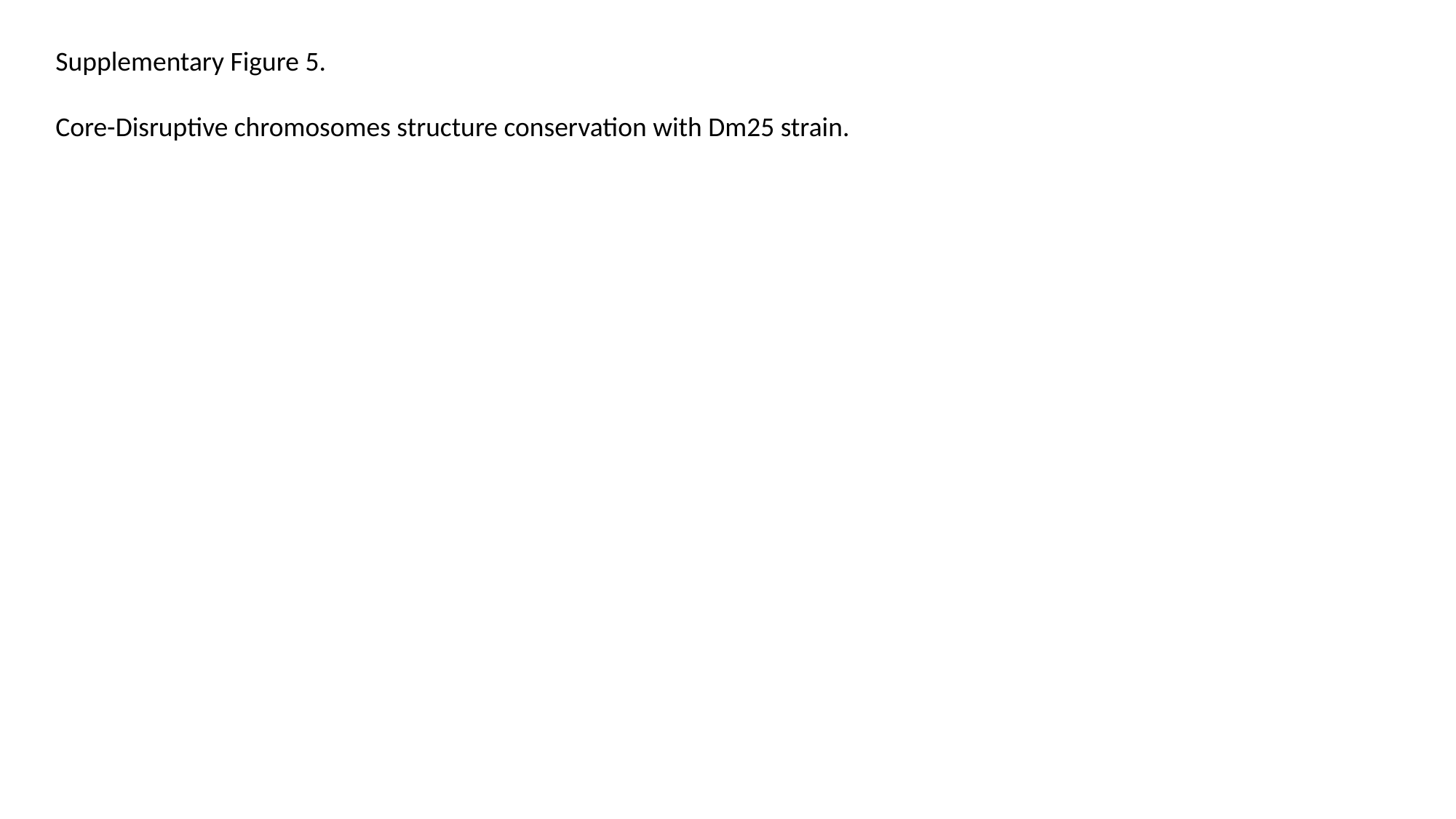

Supplementary Figure 5.
Core-Disruptive chromosomes structure conservation with Dm25 strain.

### Slide 2
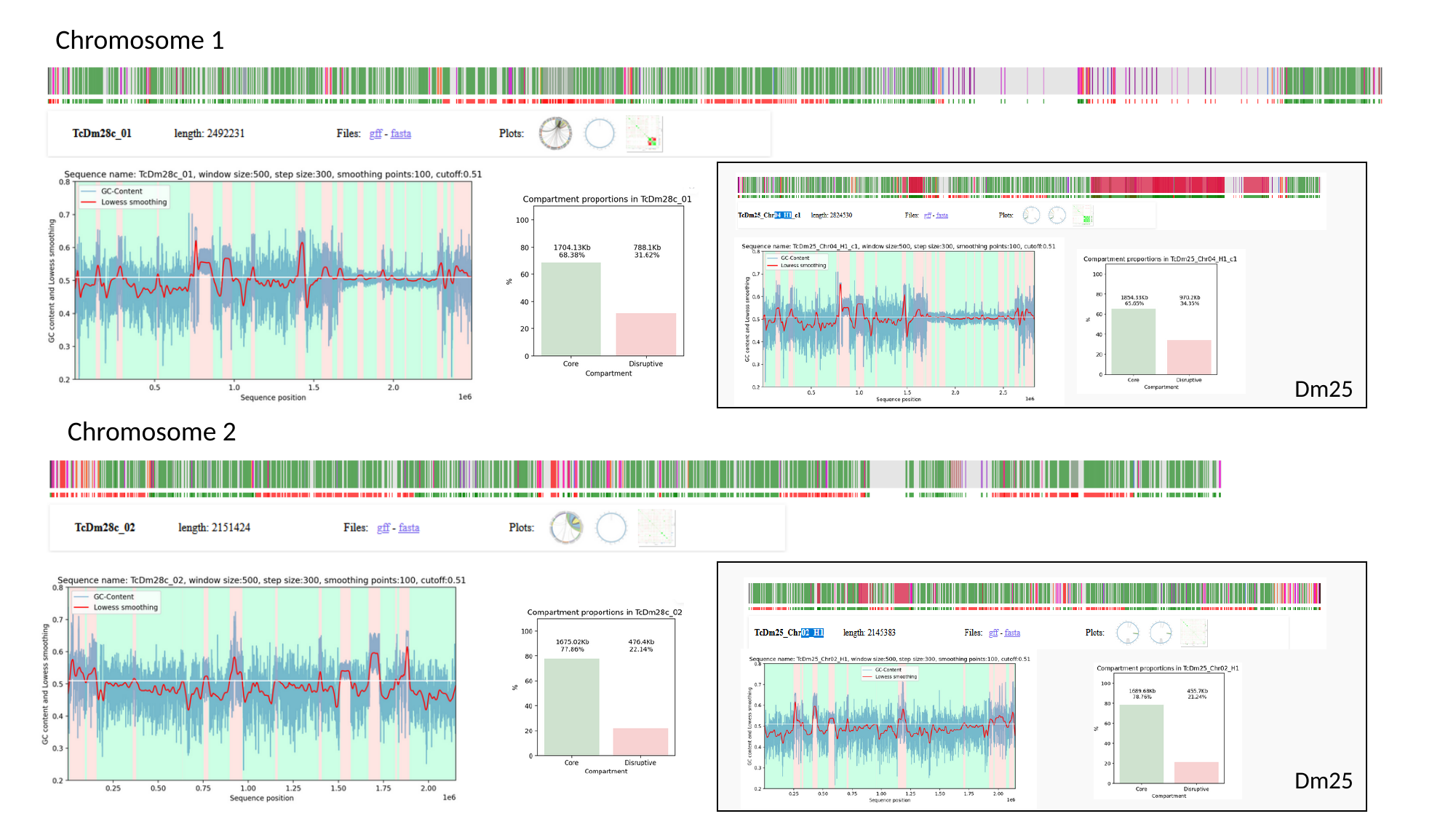

Chromosome 1
Dm25
Chromosome 2
Dm25

### Slide 3
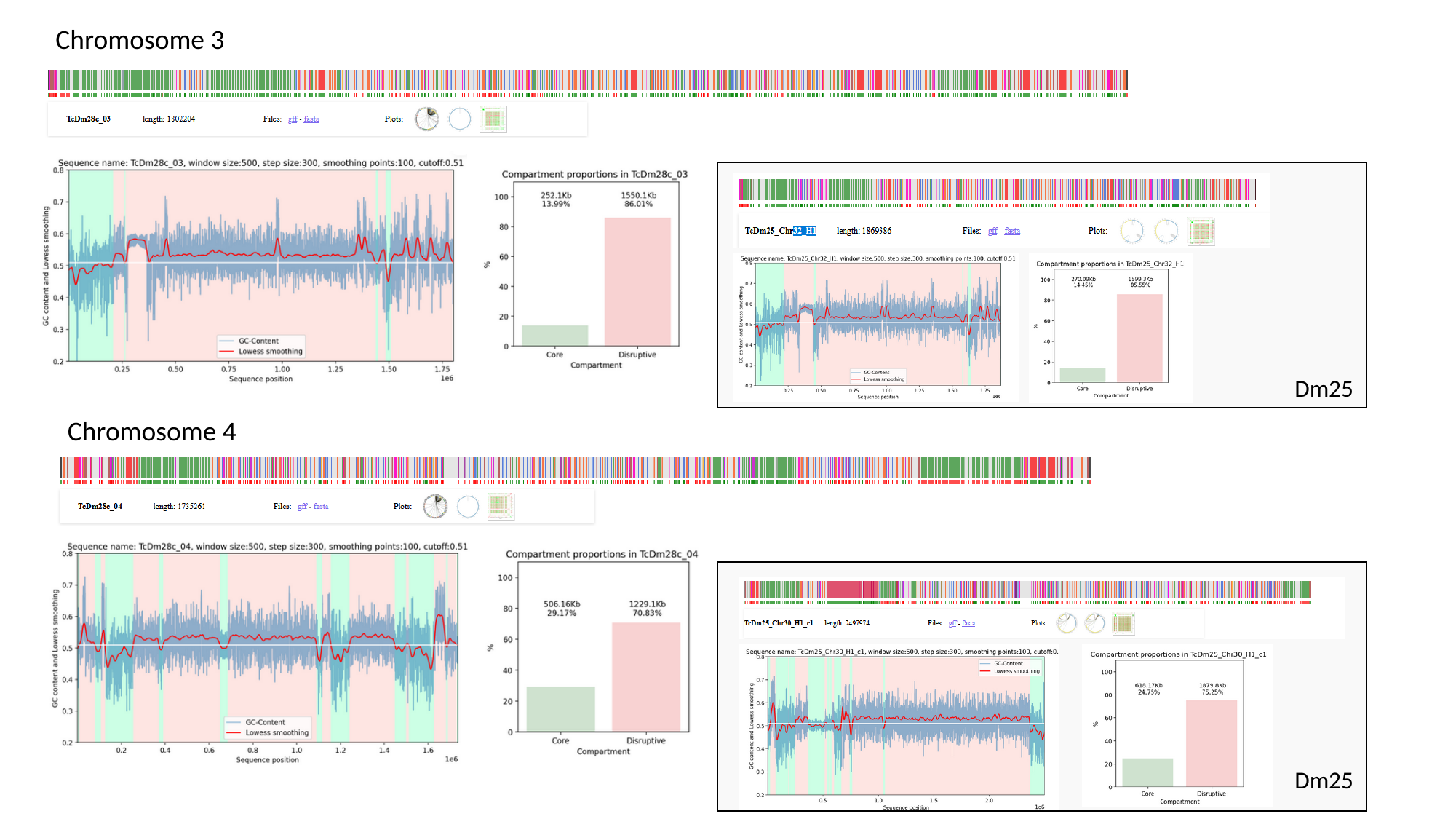

Chromosome 3
Dm25
Chromosome 4
Dm25

### Slide 4
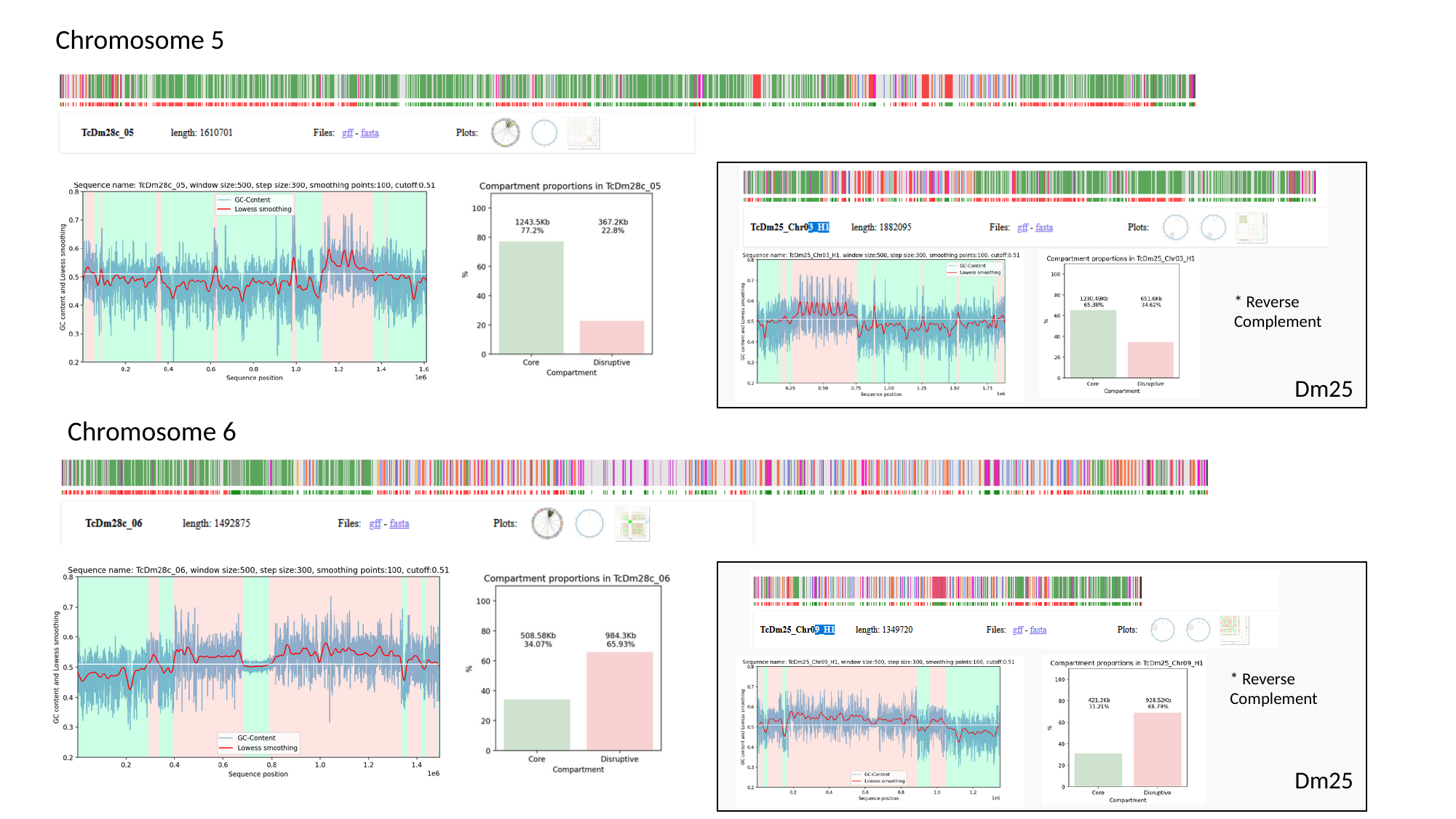

Chromosome 5
* Reverse Complement
Dm25
Chromosome 6
* Reverse Complement
Dm25

### Slide 5
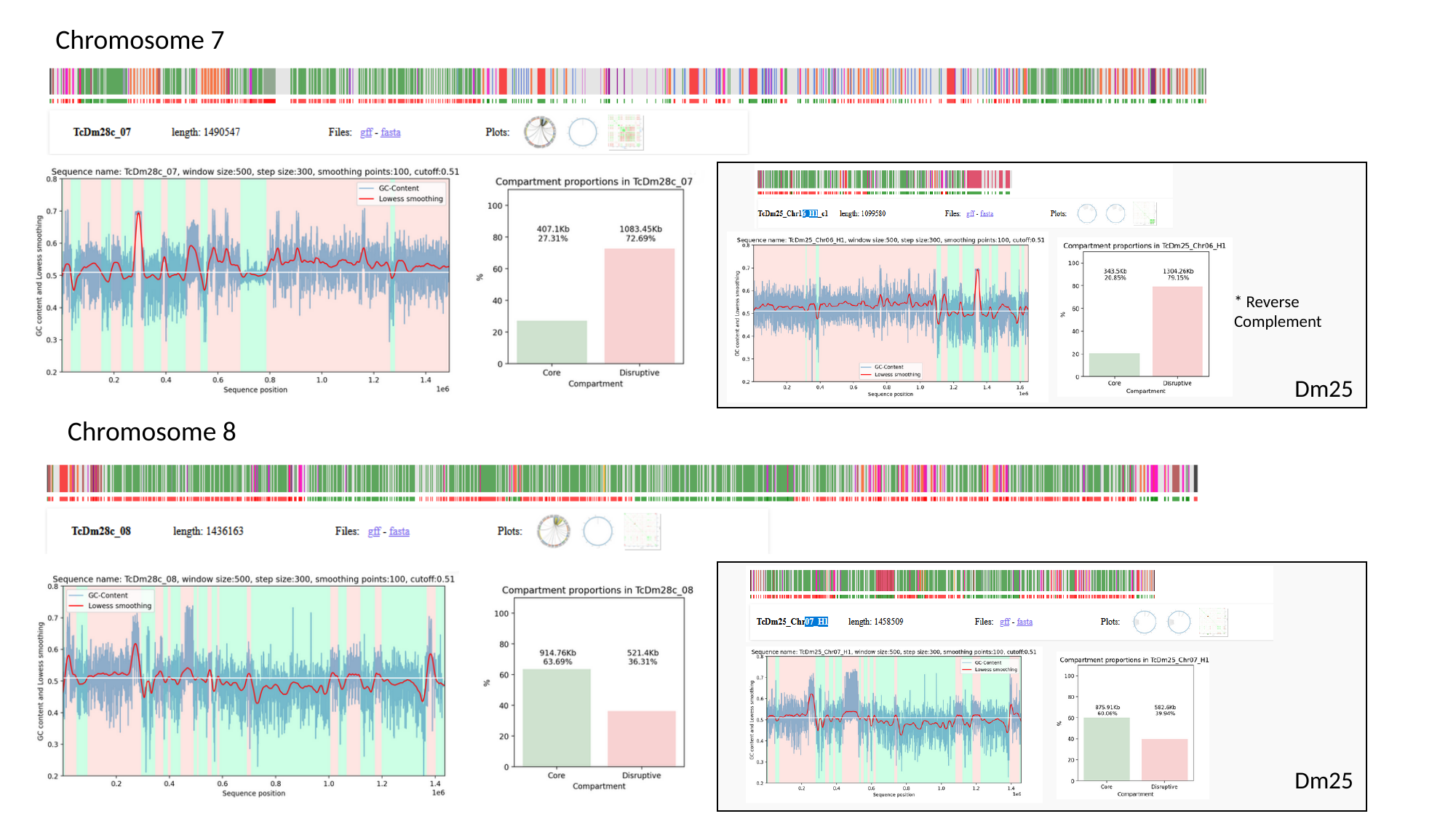

Chromosome 7
* Reverse Complement
Dm25
Chromosome 8
Dm25

### Slide 6
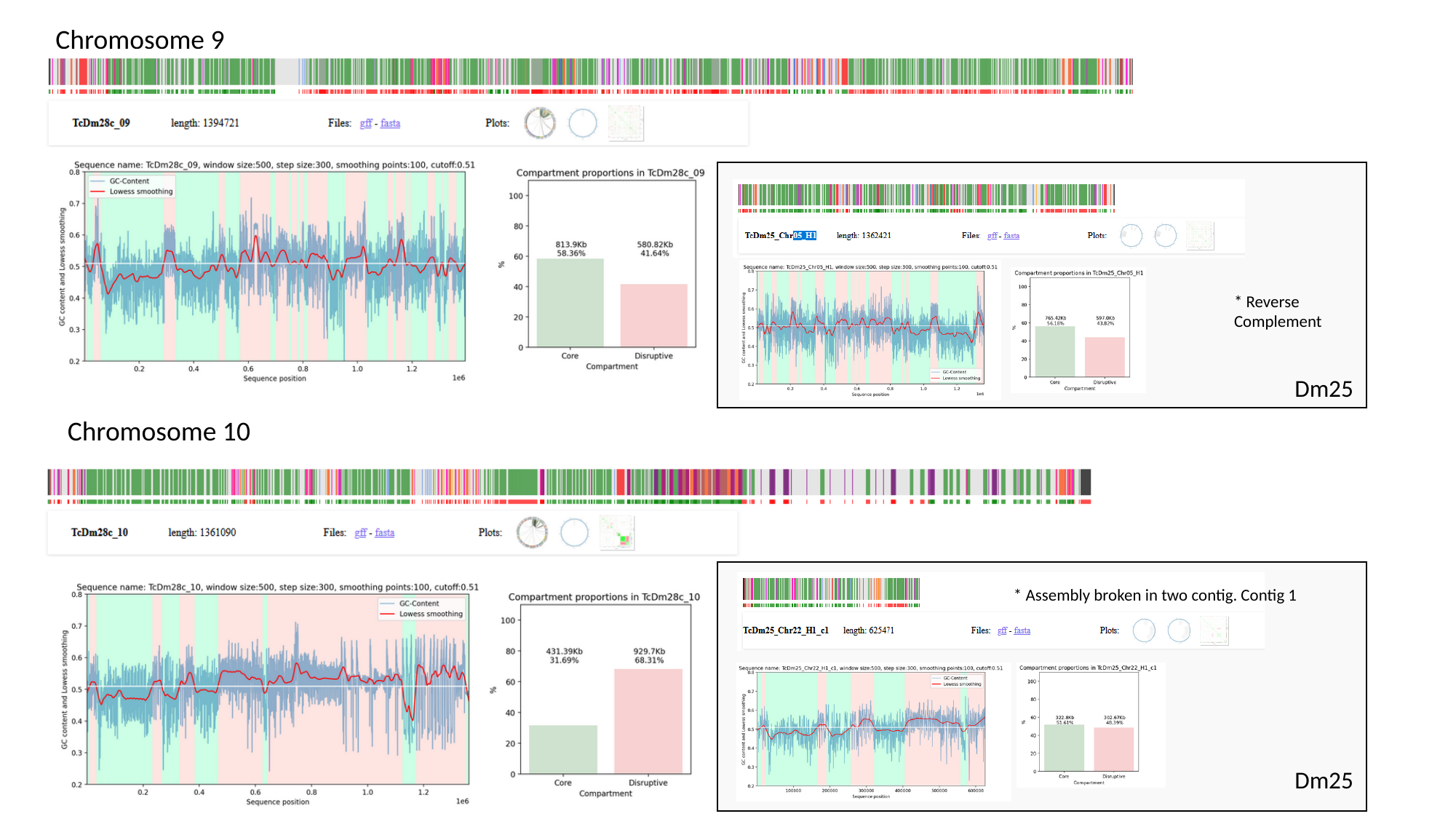

Chromosome 9
* Reverse Complement
Dm25
Chromosome 10
* Assembly broken in two contig. Contig 1
Dm25

### Slide 7
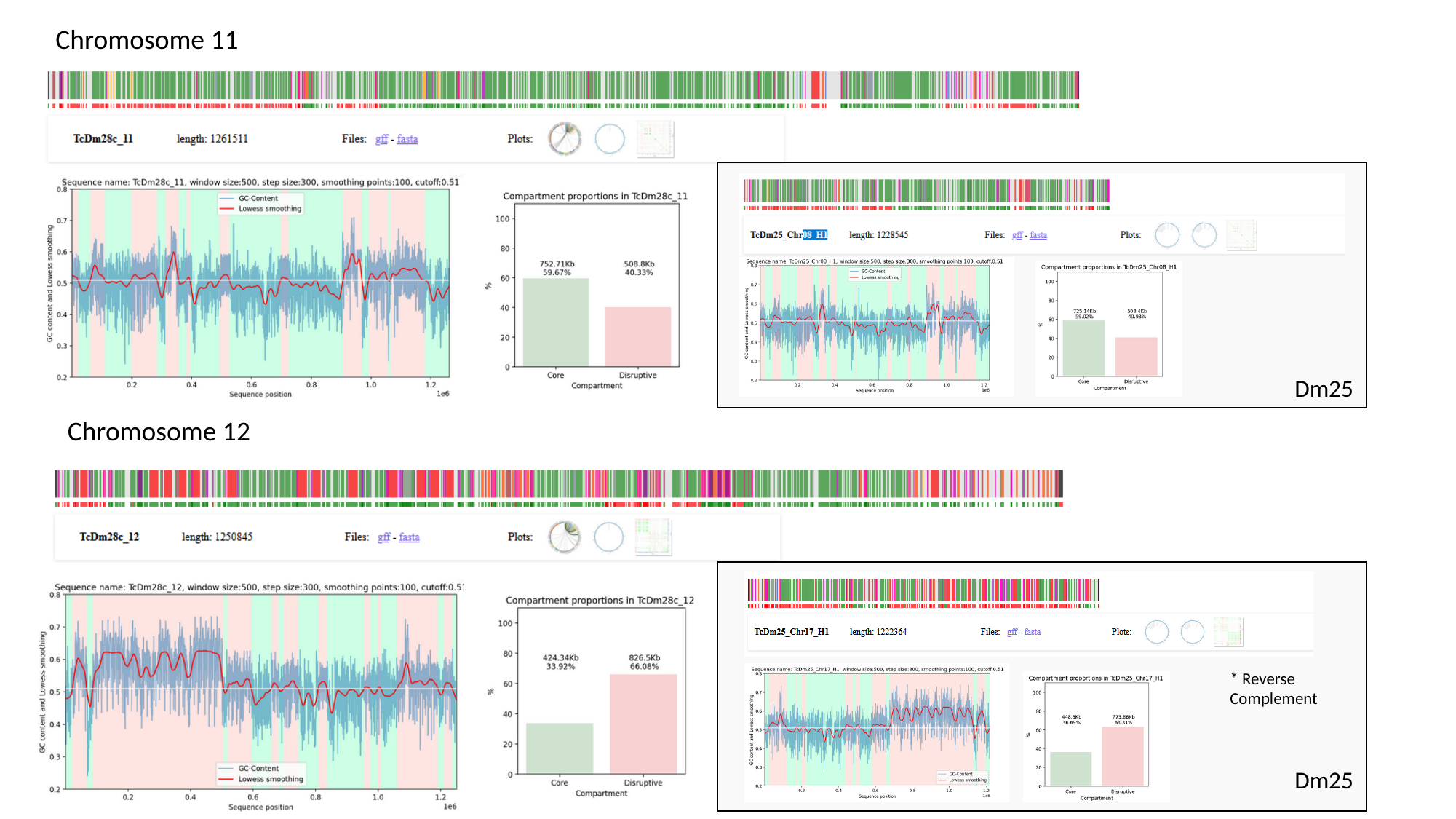

Chromosome 11
Dm25
Chromosome 12
* Reverse Complement
Dm25

### Slide 8
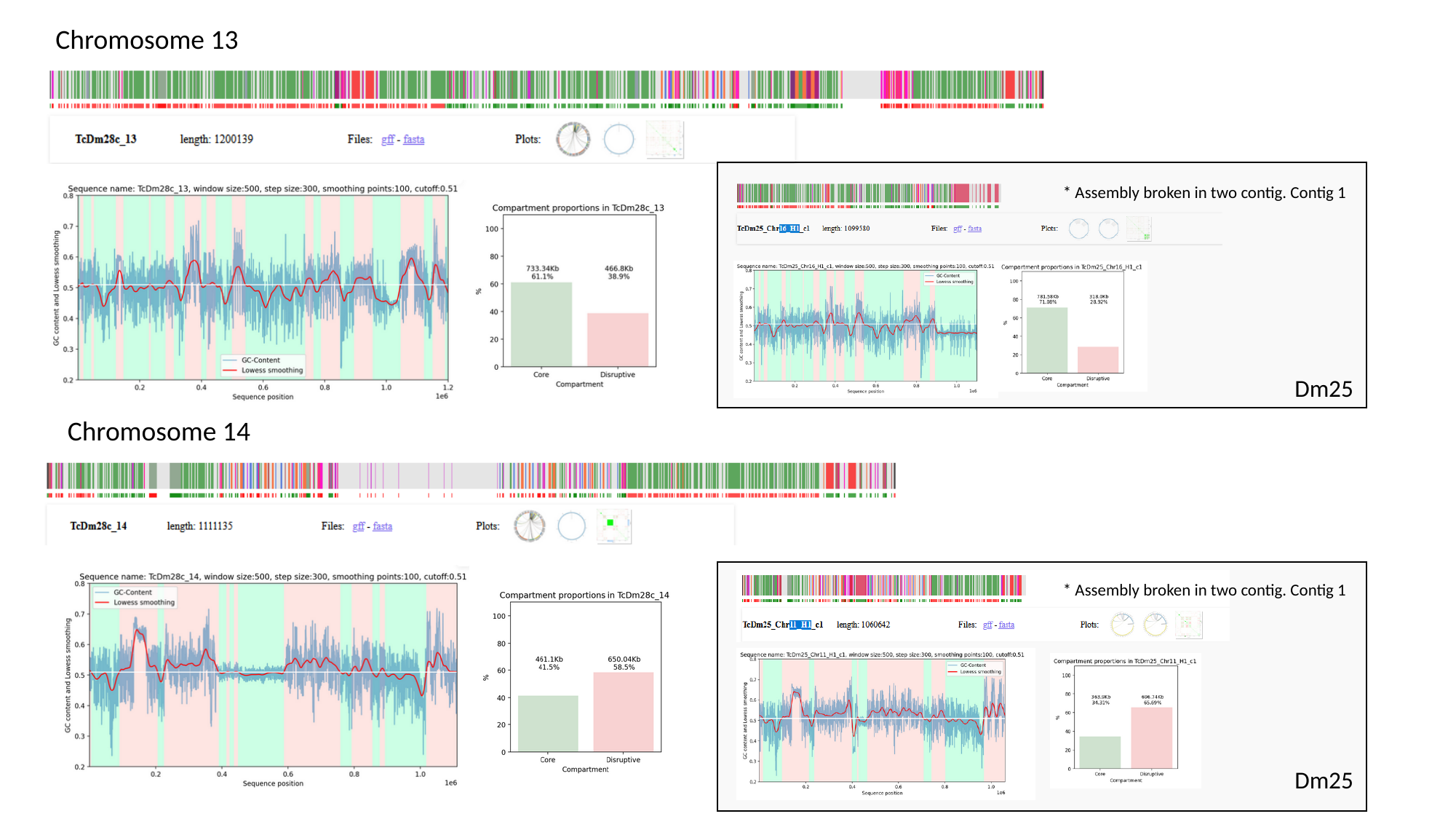

Chromosome 13
* Assembly broken in two contig. Contig 1
Dm25
Chromosome 14
* Assembly broken in two contig. Contig 1
Dm25

### Slide 9
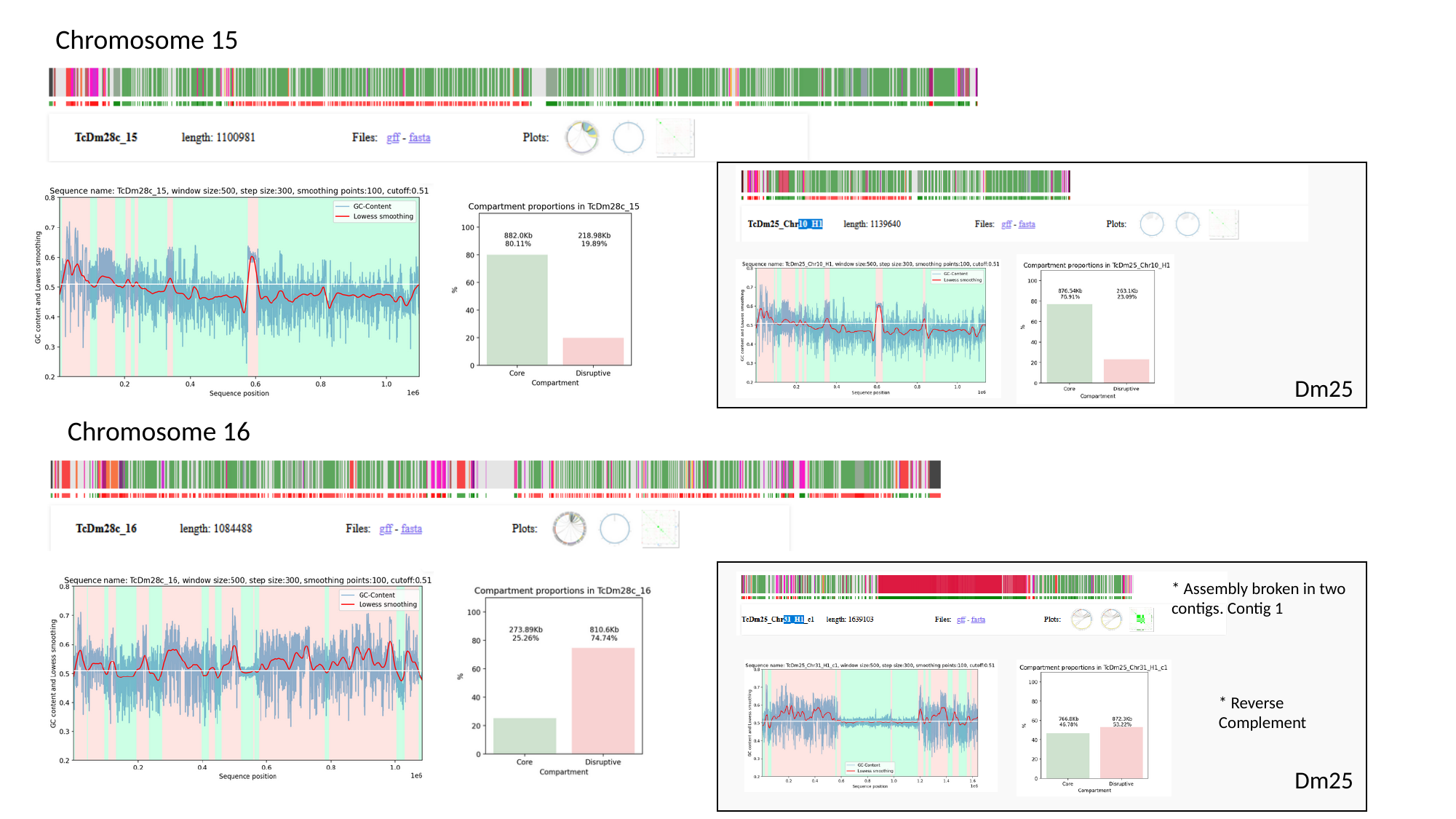

Chromosome 15
Dm25
Chromosome 16
* Assembly broken in two
contigs. Contig 1
* Reverse Complement
Dm25

### Slide 10
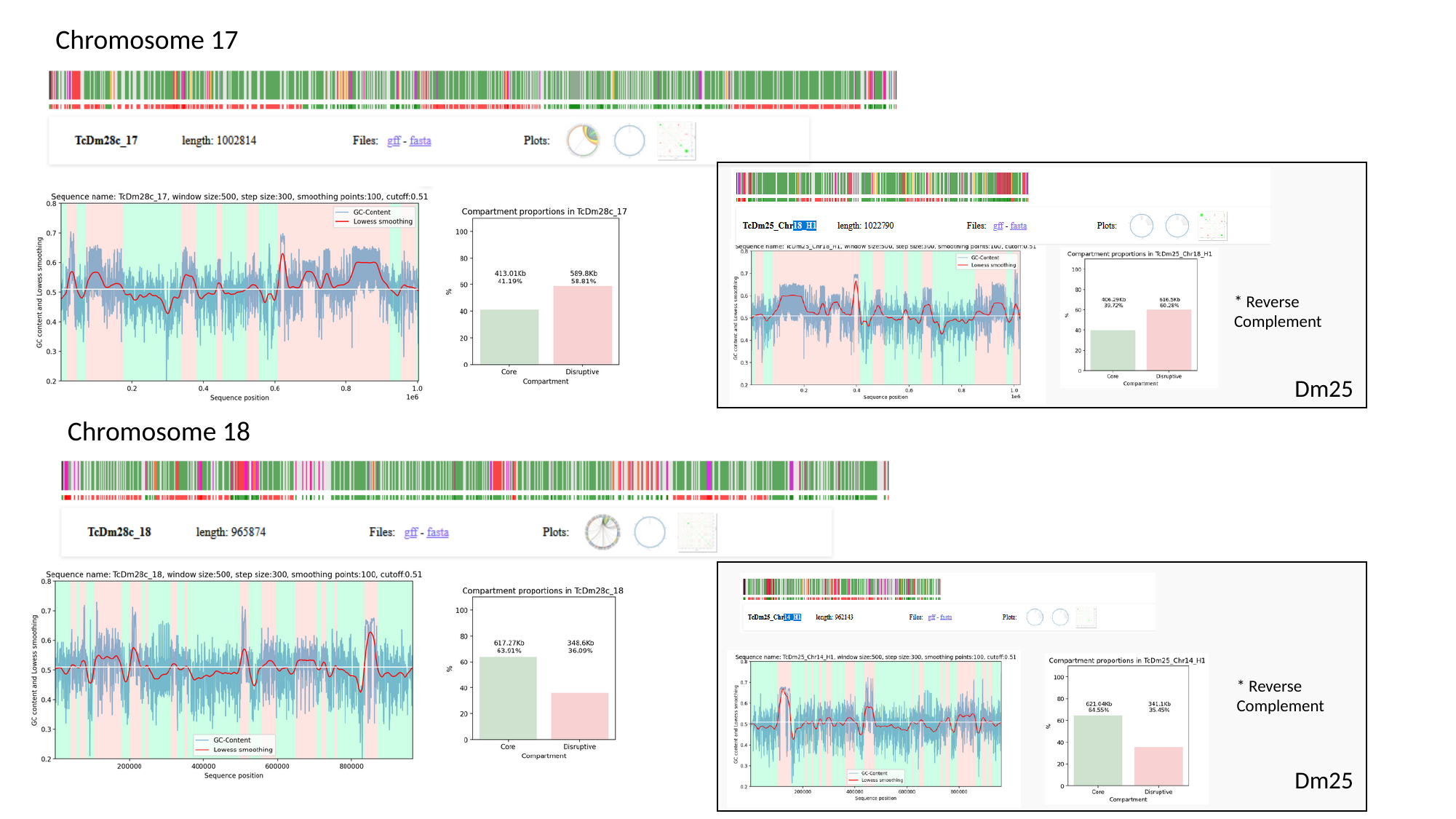

Chromosome 17
* Reverse Complement
Dm25
Chromosome 18
* Reverse Complement
Dm25

### Slide 11
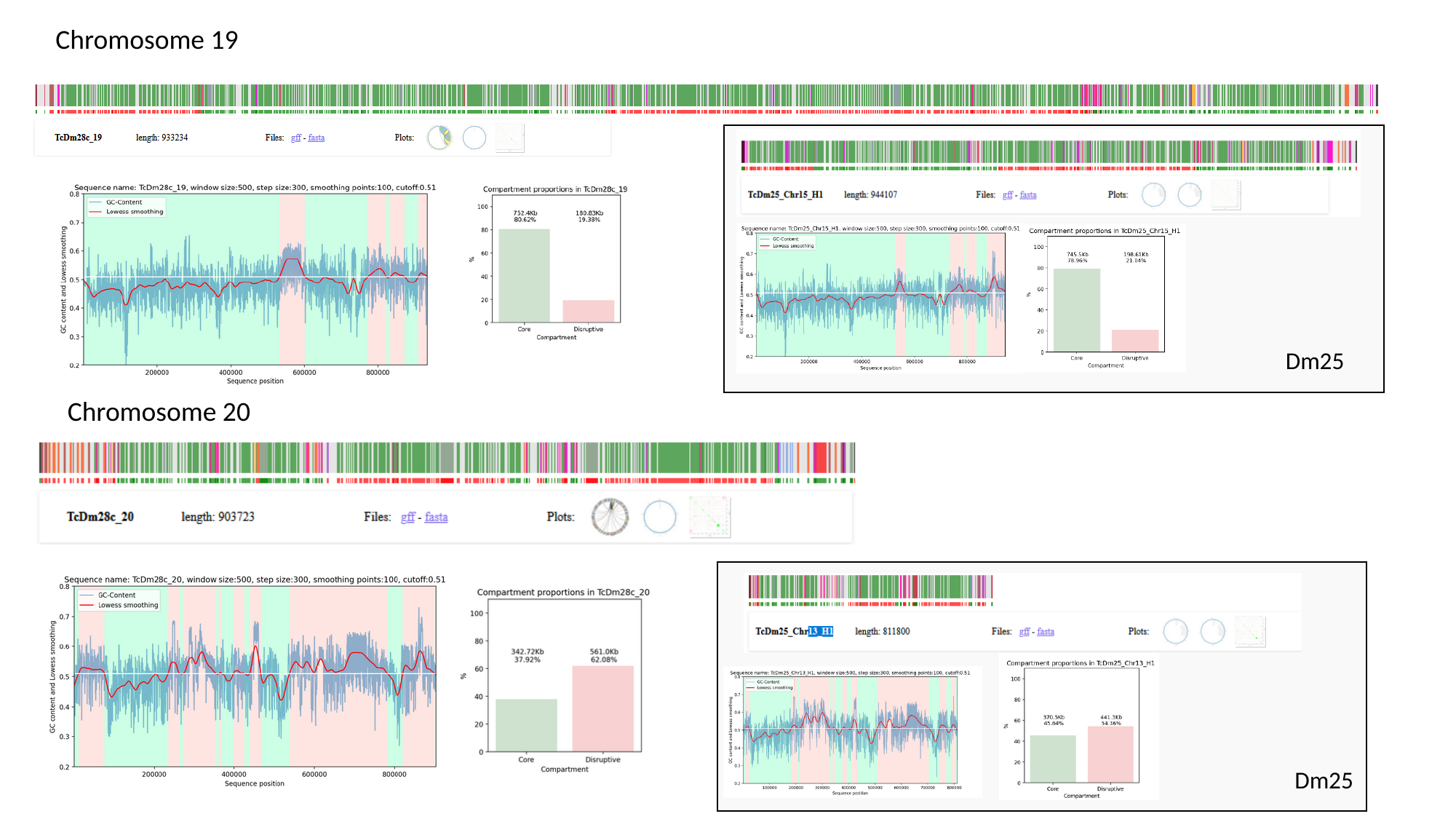

Chromosome 19
Dm25
Chromosome 20
Dm25

### Slide 12
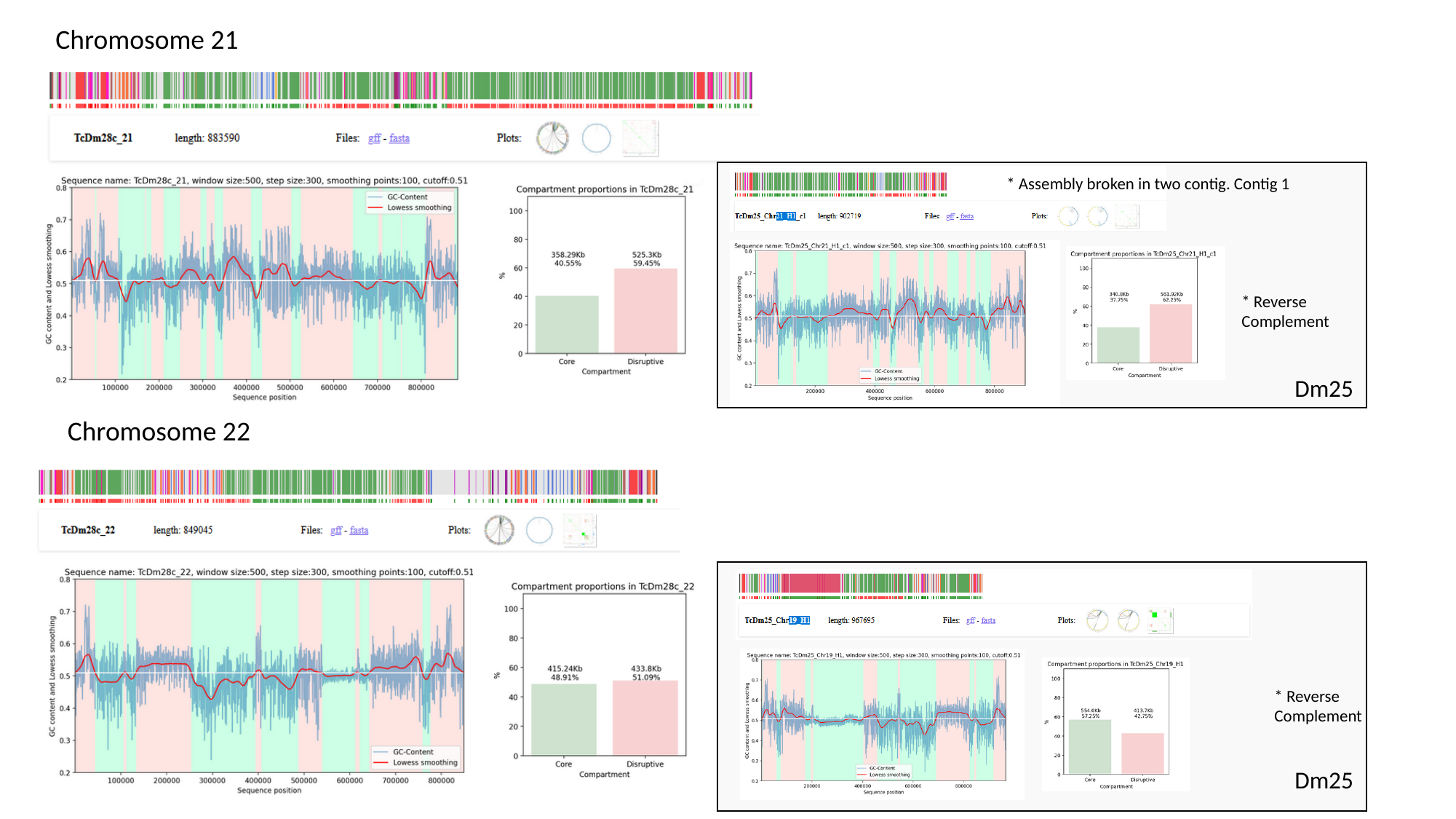

Chromosome 21
* Assembly broken in two contig. Contig 1
* Reverse Complement
Dm25
Chromosome 22
* Reverse Complement
Dm25

### Slide 13
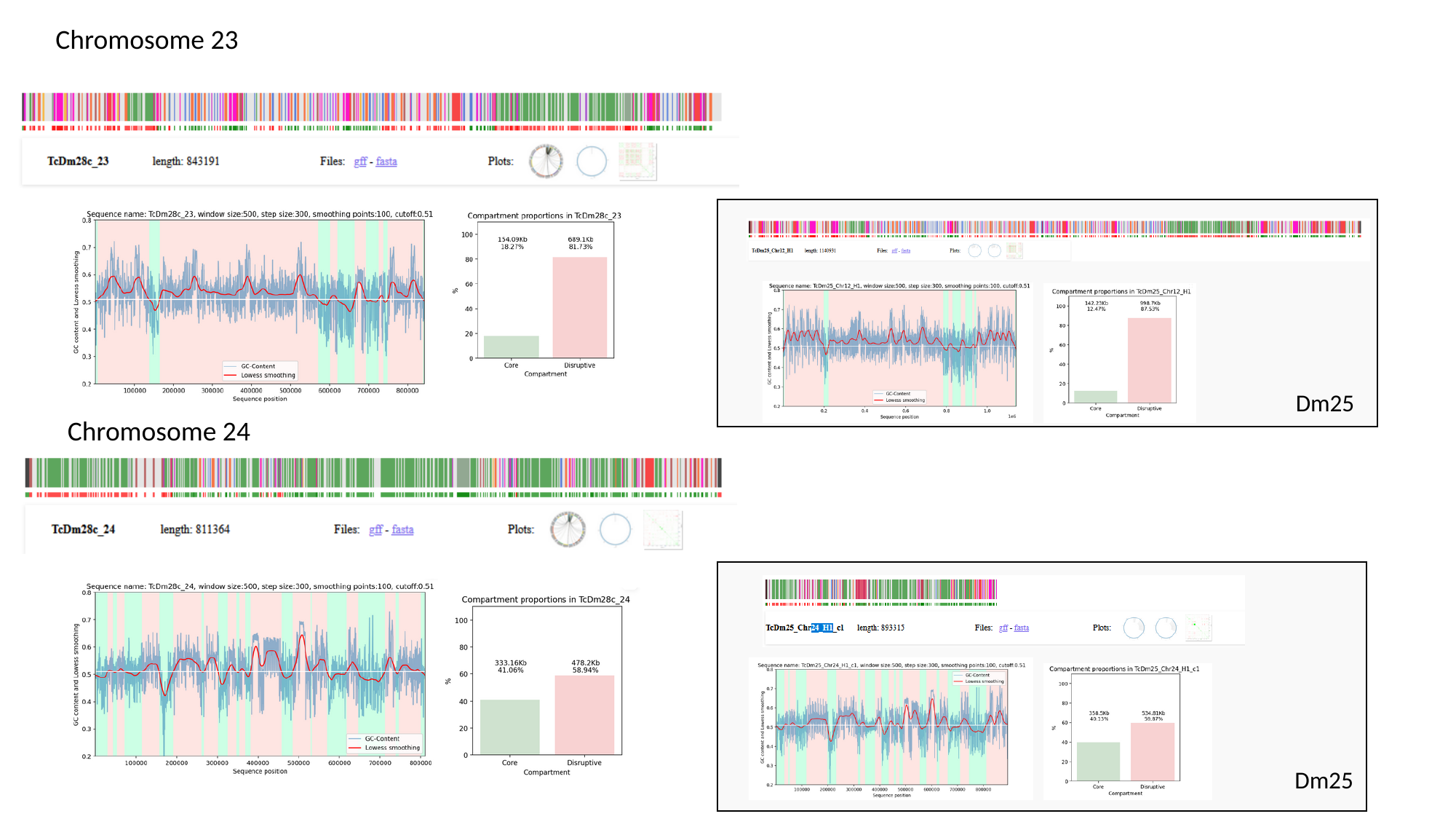

Chromosome 23
Dm25
Chromosome 24
Dm25

### Slide 14
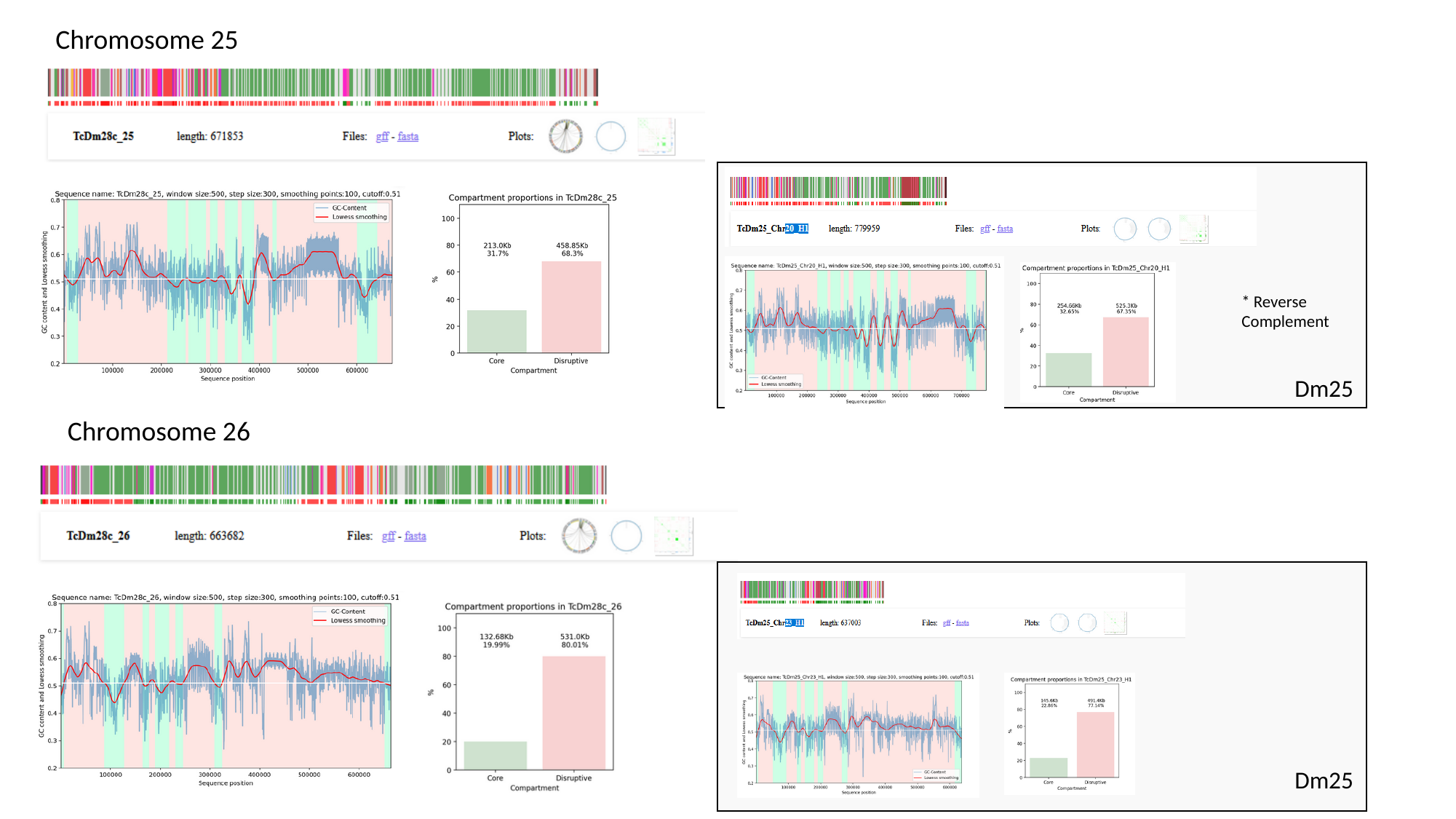

Chromosome 25
* Reverse Complement
Dm25
Chromosome 26
Dm25

### Slide 15
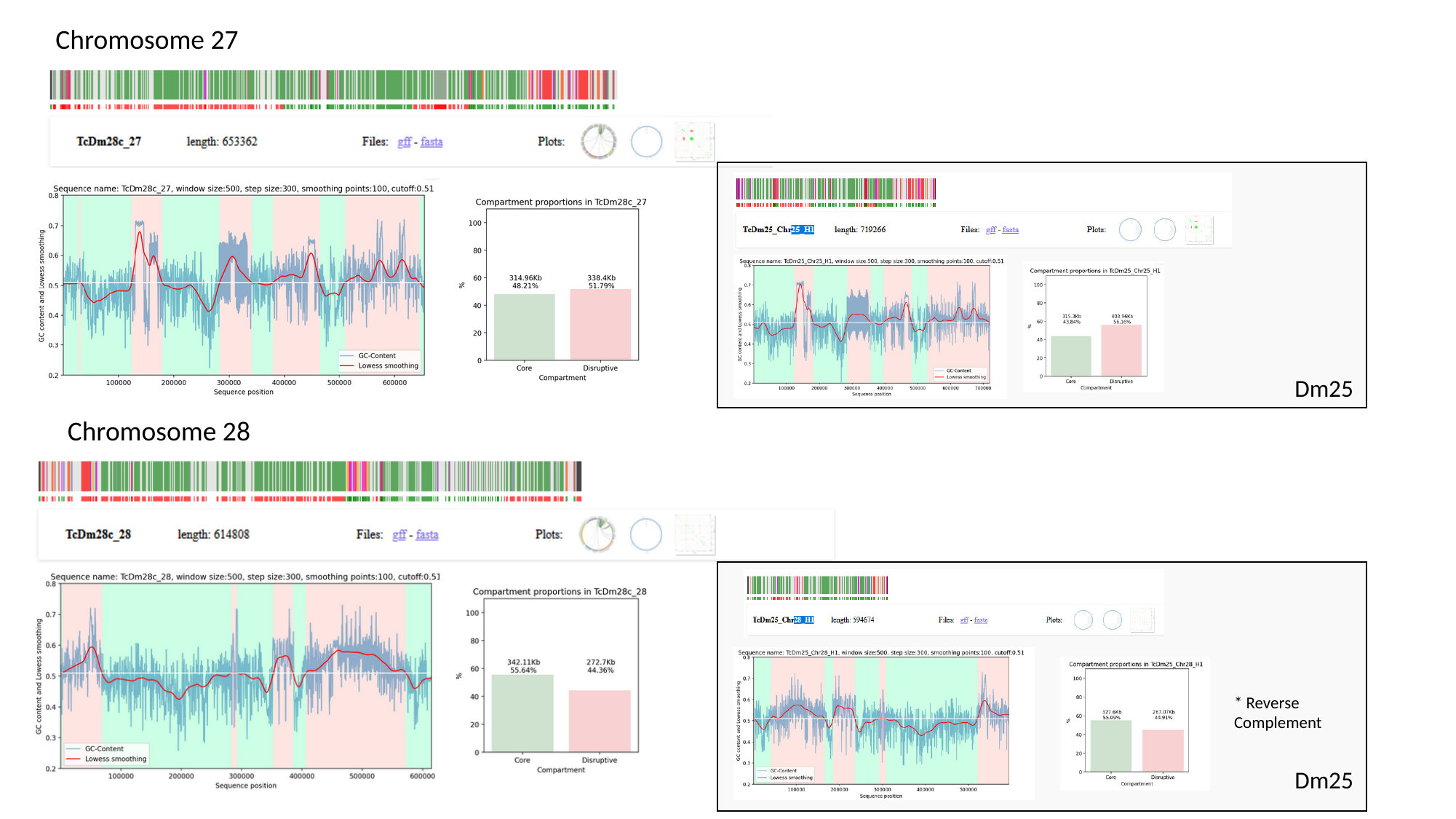

Chromosome 27
Dm25
Chromosome 28
* Reverse Complement
Dm25

### Slide 16
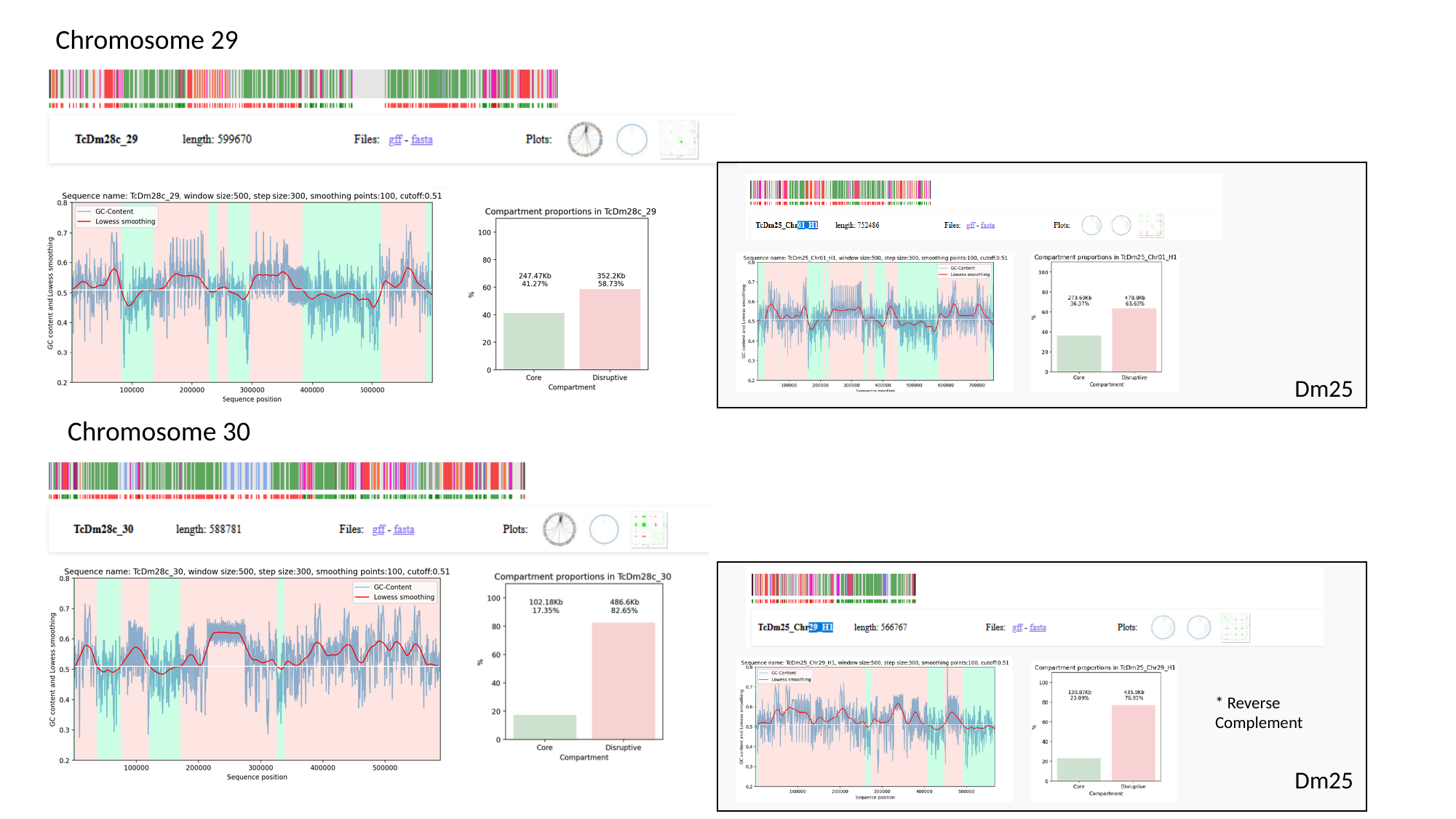

Chromosome 29
Dm25
Chromosome 30
* Reverse Complement
Dm25

### Slide 17
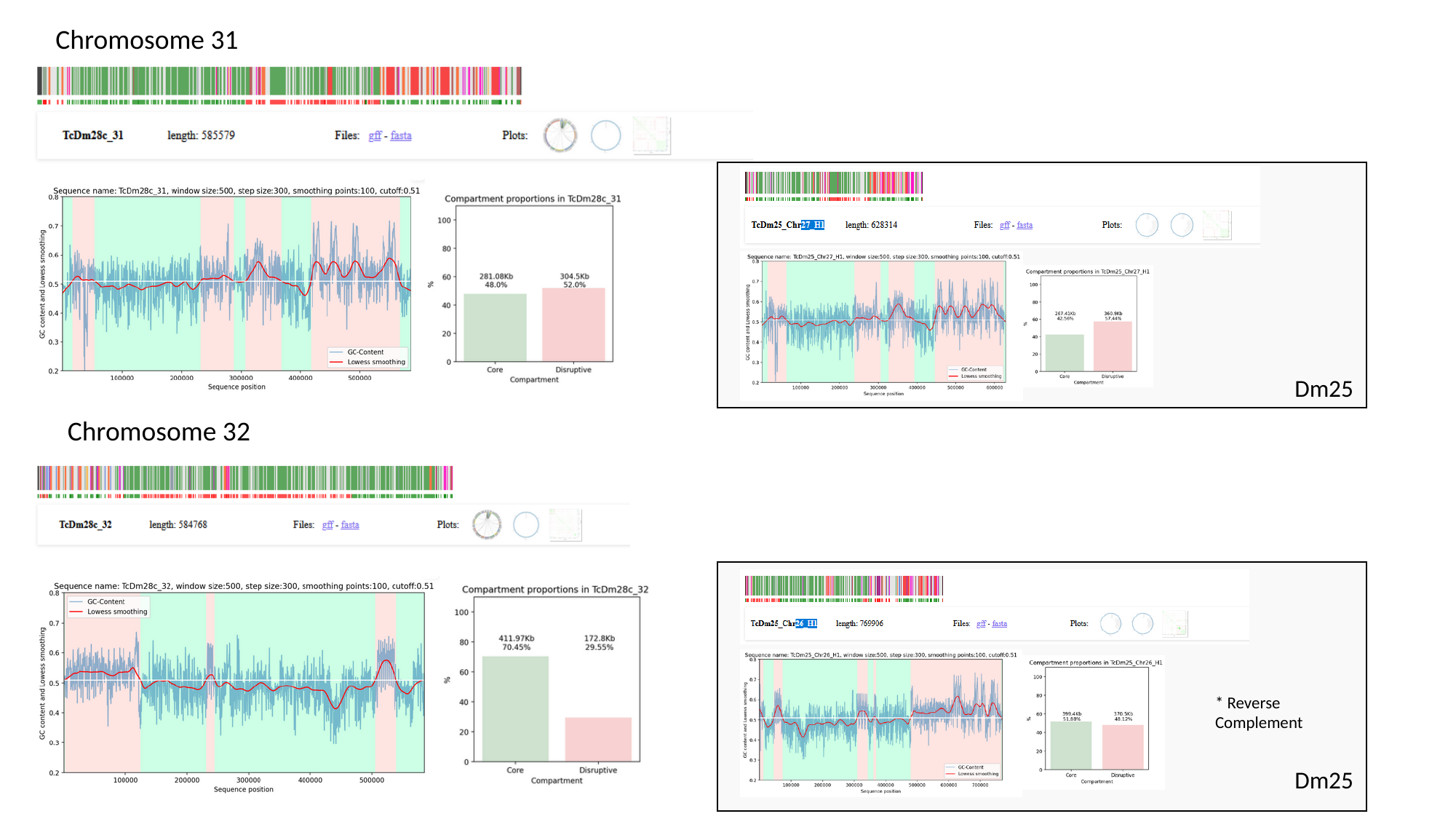

Chromosome 31
Dm25
Chromosome 32
* Reverse Complement
Dm25
