## Supplementary material for "*Trypanosoma cruzi* has 32 Chromosomes: A Telomere-to-Telomere Assembly Defines Its Karyotype": SupFigure6_GeneProportions.pdf

Supplementary Figure 6.

Gene proportions (Conserved, MASP/mucin, trans-sialidase and RHS) in 5 gene Windows across all chromosomes.

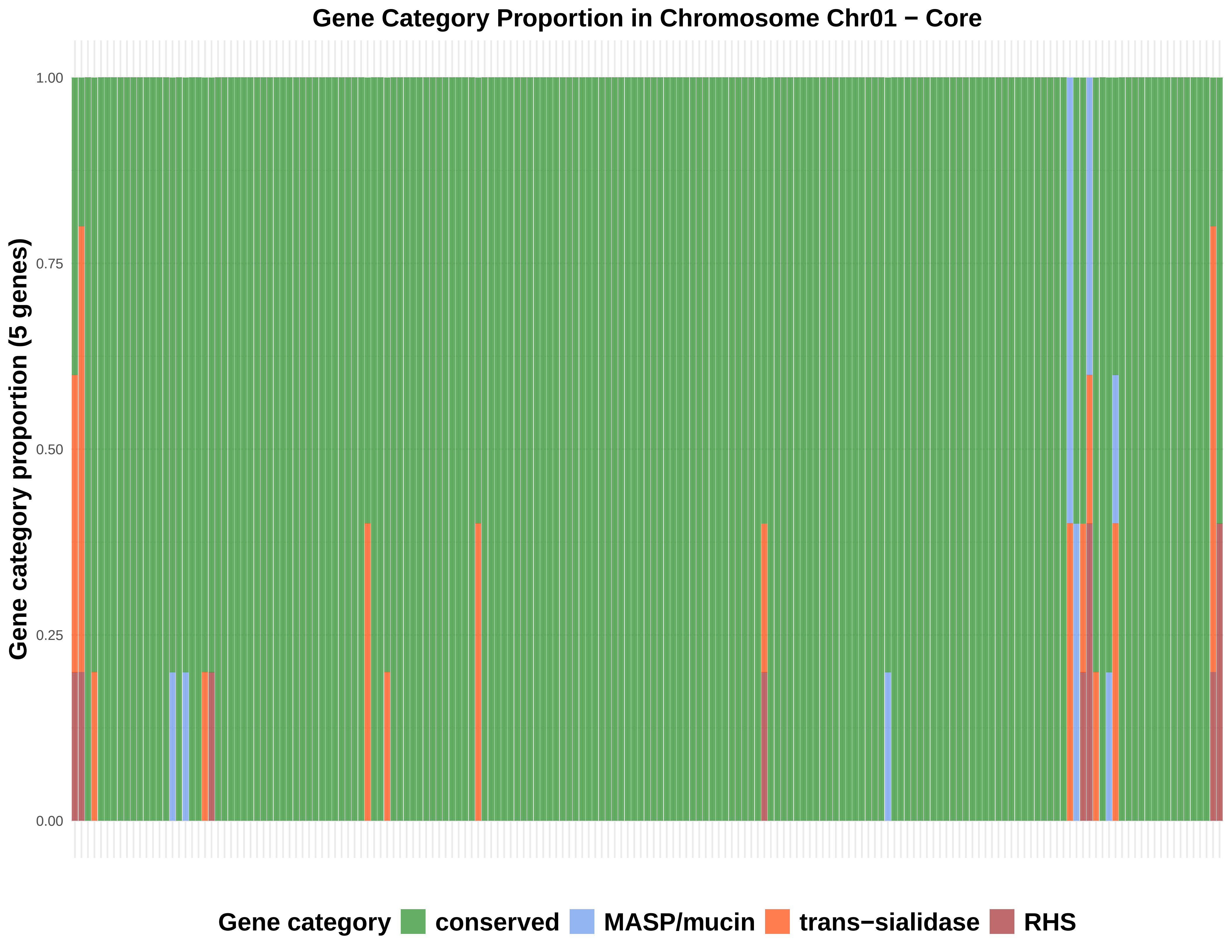

Gene Category Proportion in Chromosome Chr01 − Core
Gene category proportion (5 genes)
1.00
0.75
0.50
0.25
0.00
Gene category
conserved
MASP/mucin
trans−sialidase
RHS

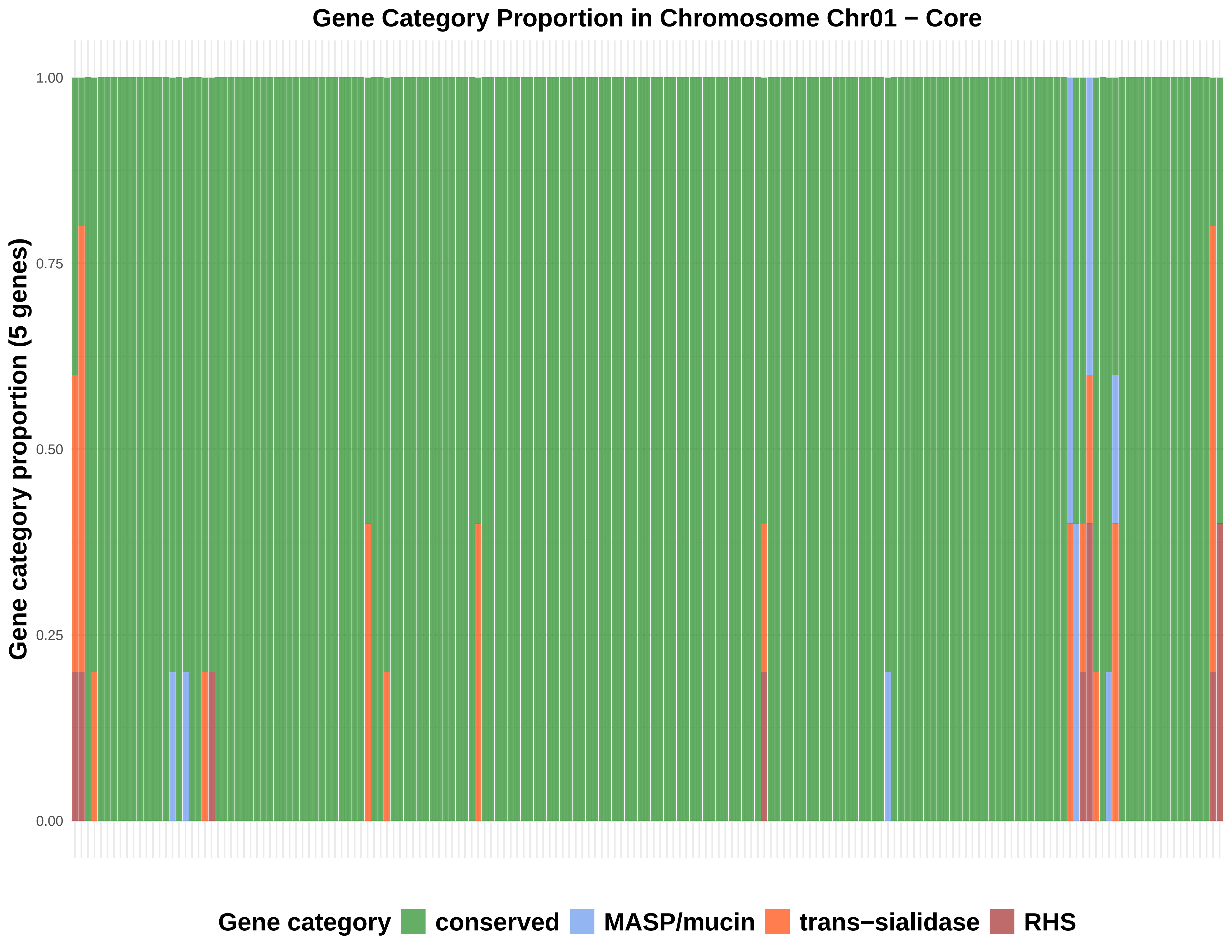
Gene Category Proportion in Chromosome Chr01 − Core
Gene category proportion (5 genes)
1.00
0.75
0.50
0.25
0.00
Gene category
conserved
MASP/mucin
trans−sialidase
RHS

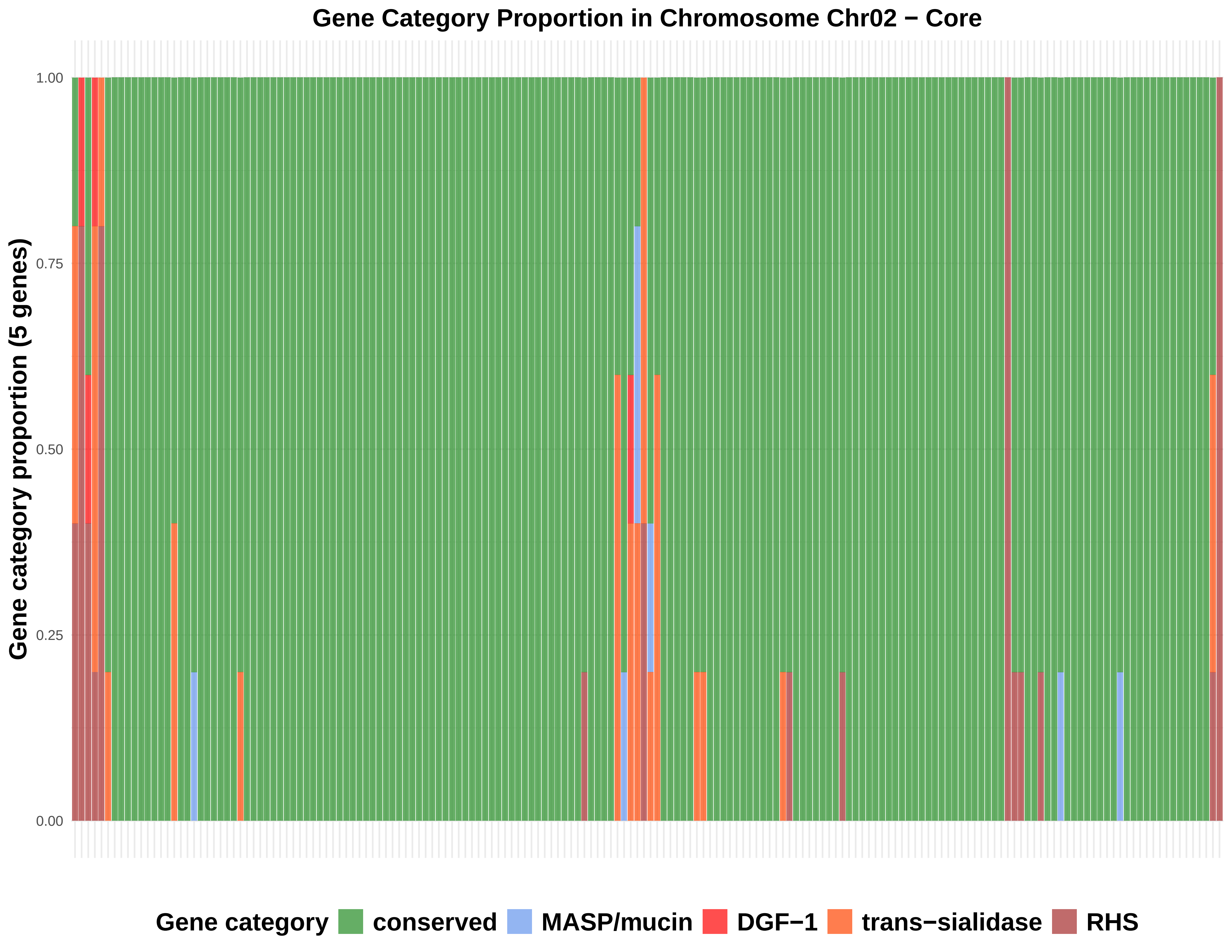

Gene Category Proportion in Chromosome Chr02 − Core
Gene category proportion (5 genes)
1.00
0.75
0.50
0.25
0.00
Gene category
conserved
MASP/mucin
DGF−1
trans−sialidase
RHS

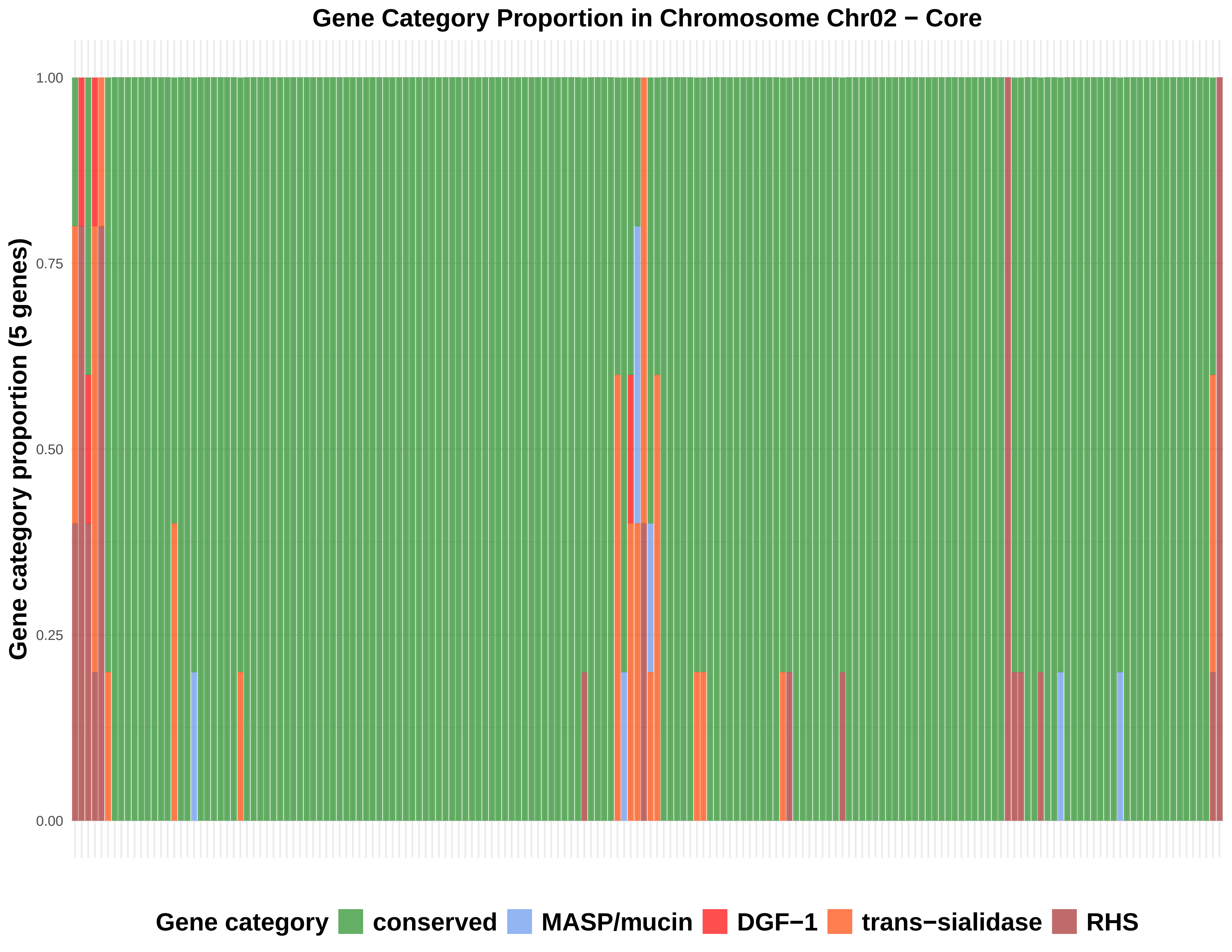
Gene Category Proportion in Chromosome Chr02 − Core
Gene category proportion (5 genes)
1.00
0.75
0.50
0.25
0.00
Gene category
conserved
MASP/mucin
DGF−1
trans−sialidase
RHS

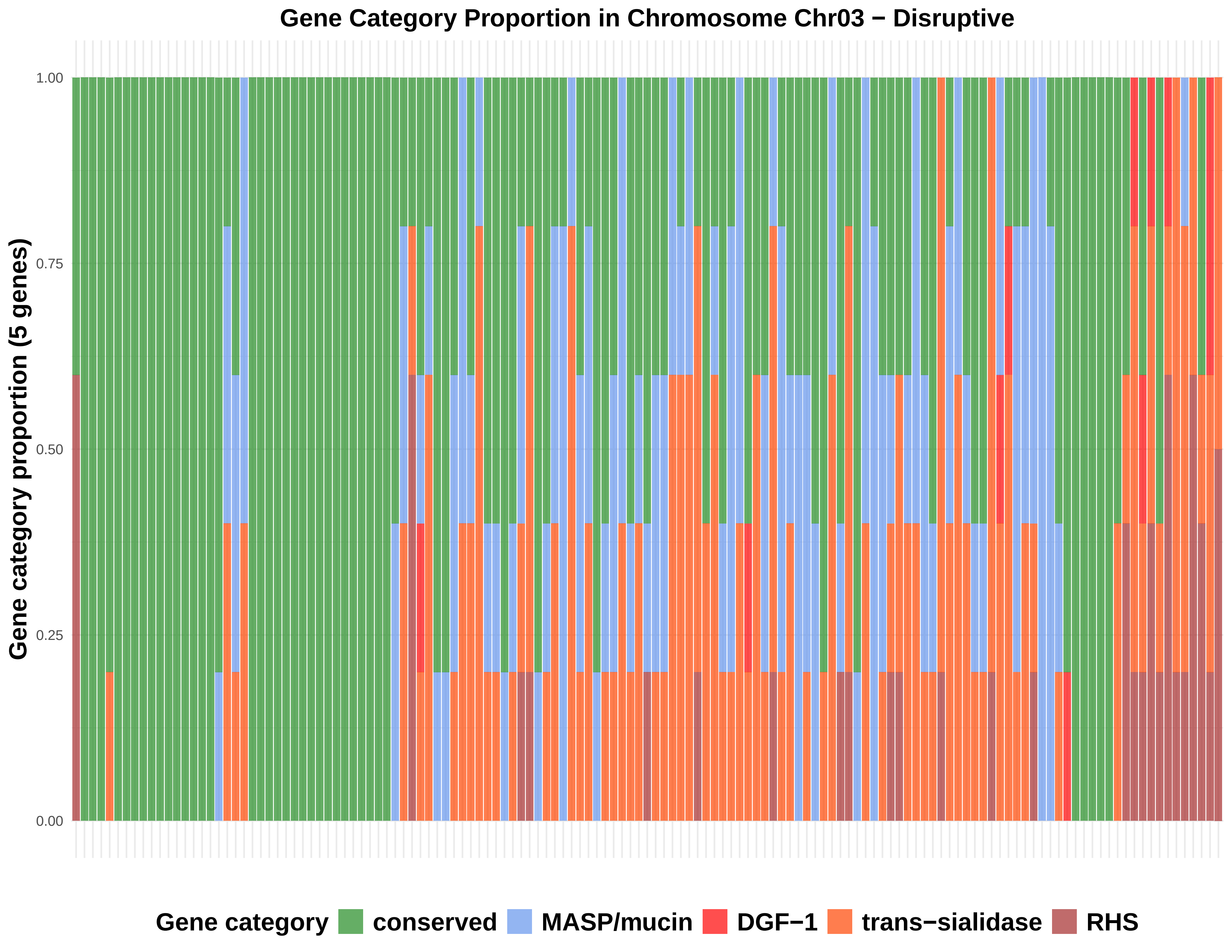

Gene Category Proportion in Chromosome Chr03 − Disruptive
Gene category proportion (5 genes)
1.00
0.75
0.50
0.25
0.00
Gene category
conserved
MASP/mucin
DGF−1
trans−sialidase
RHS

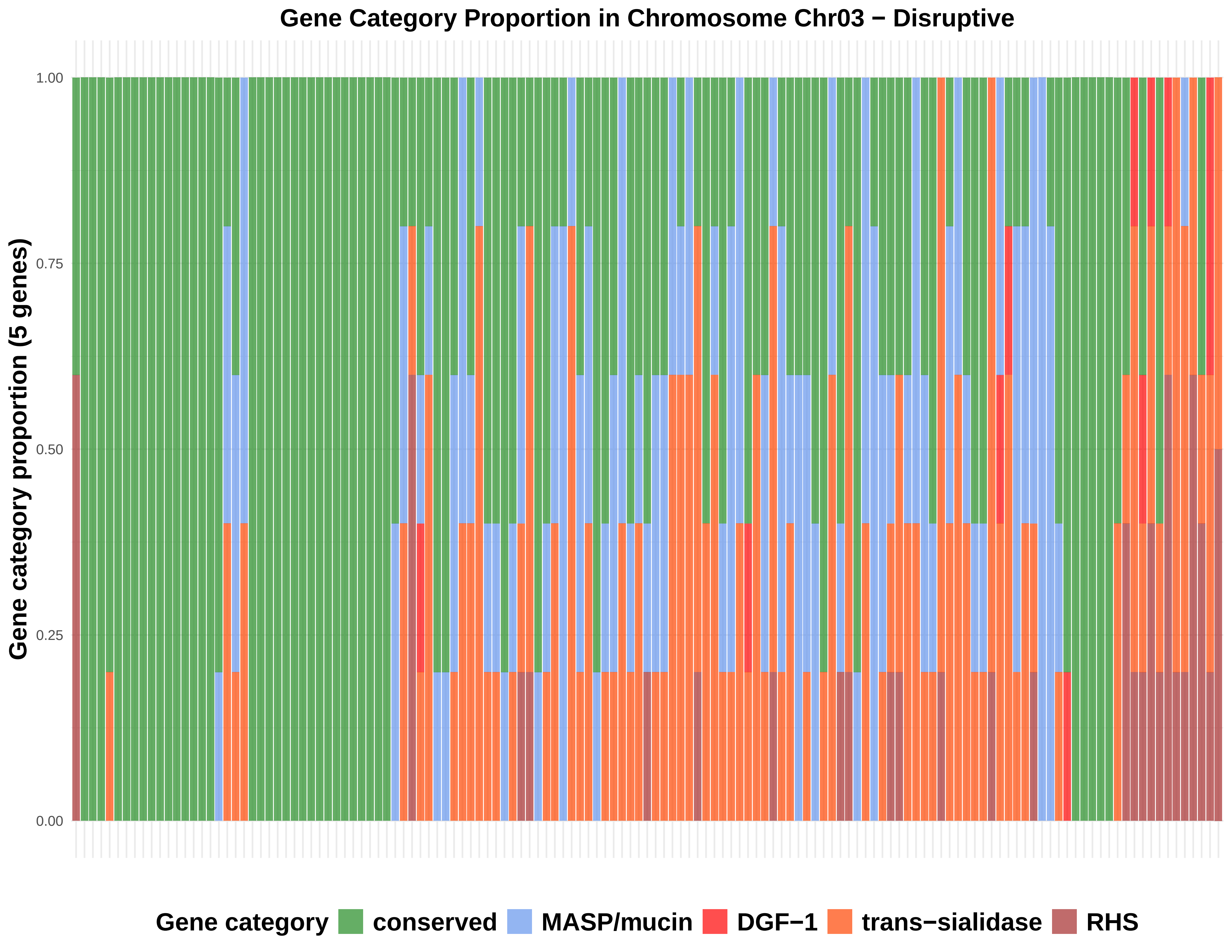
Gene Category Proportion in Chromosome Chr03 − Disruptive
Gene category proportion (5 genes)
1.00
0.75
0.50
0.25
0.00
Gene category
conserved
MASP/mucin
DGF−1
trans−sialidase
RHS

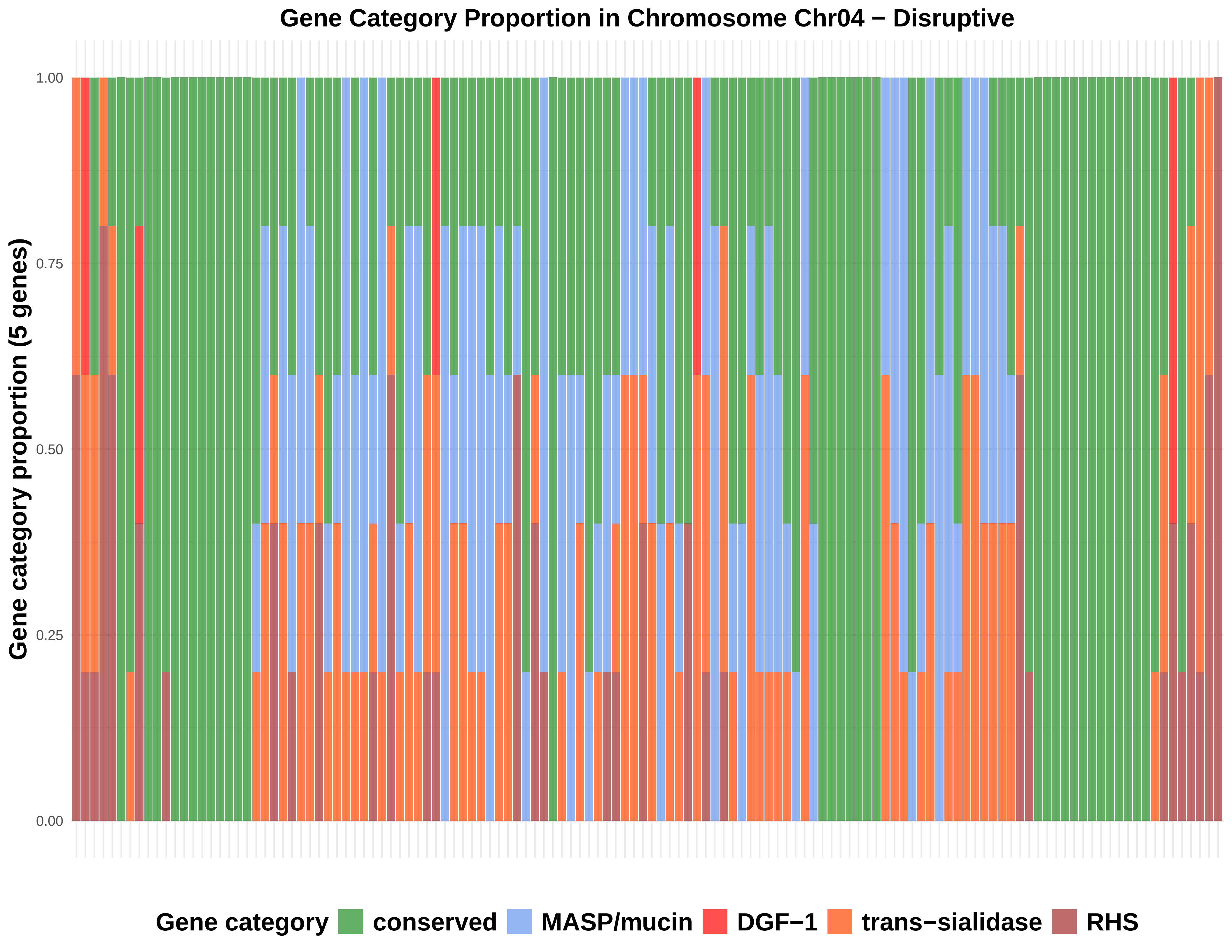

Gene Category Proportion in Chromosome Chr04 − Disruptive
Gene category proportion (5 genes)
1.00
0.75
0.50
0.25
0.00
Gene category
conserved
MASP/mucin
DGF−1
trans−sialidase
RHS

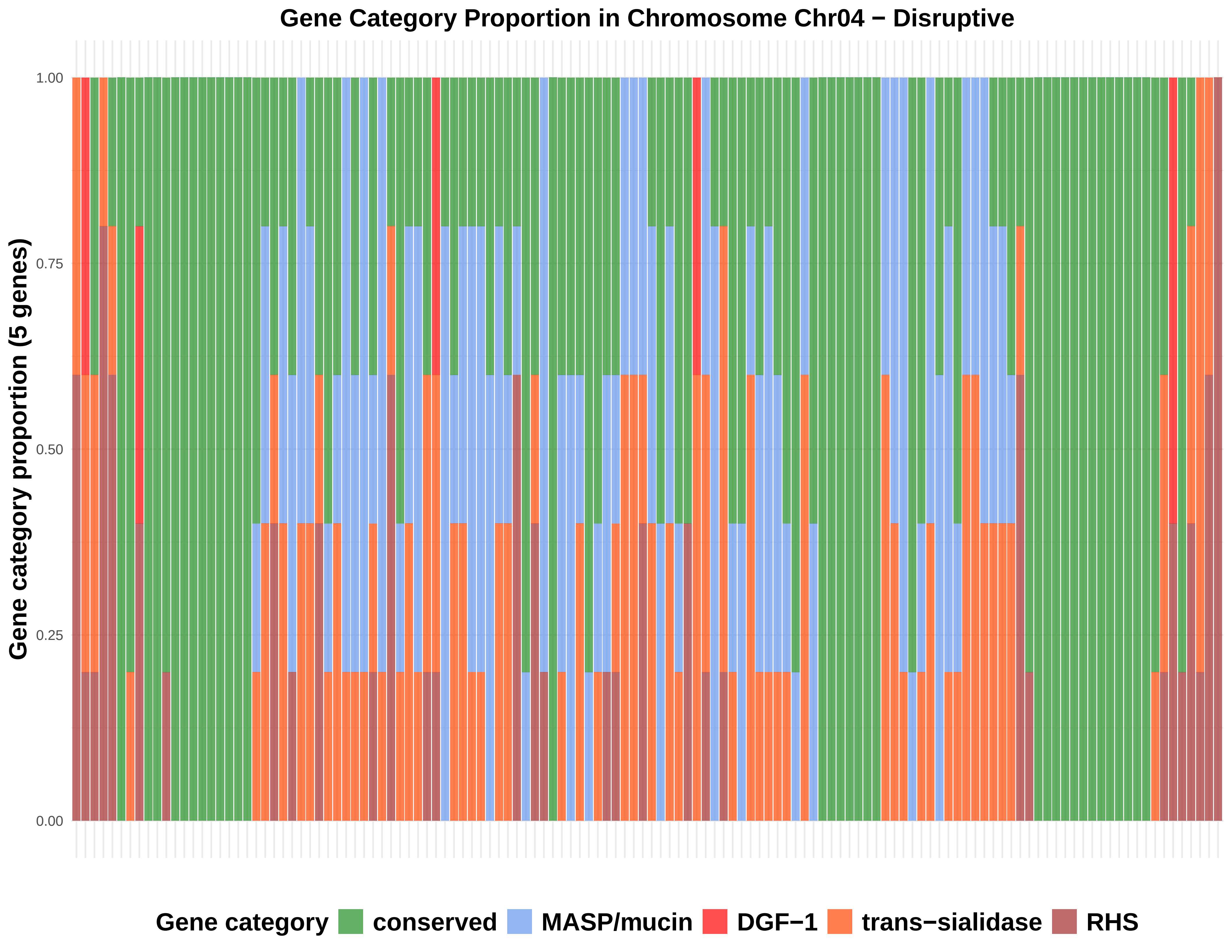
Gene Category Proportion in Chromosome Chr04 − Disruptive
Gene category proportion (5 genes)
1.00
0.75
0.50
0.25
0.00
Gene category
conserved
MASP/mucin
DGF−1
trans−sialidase
RHS

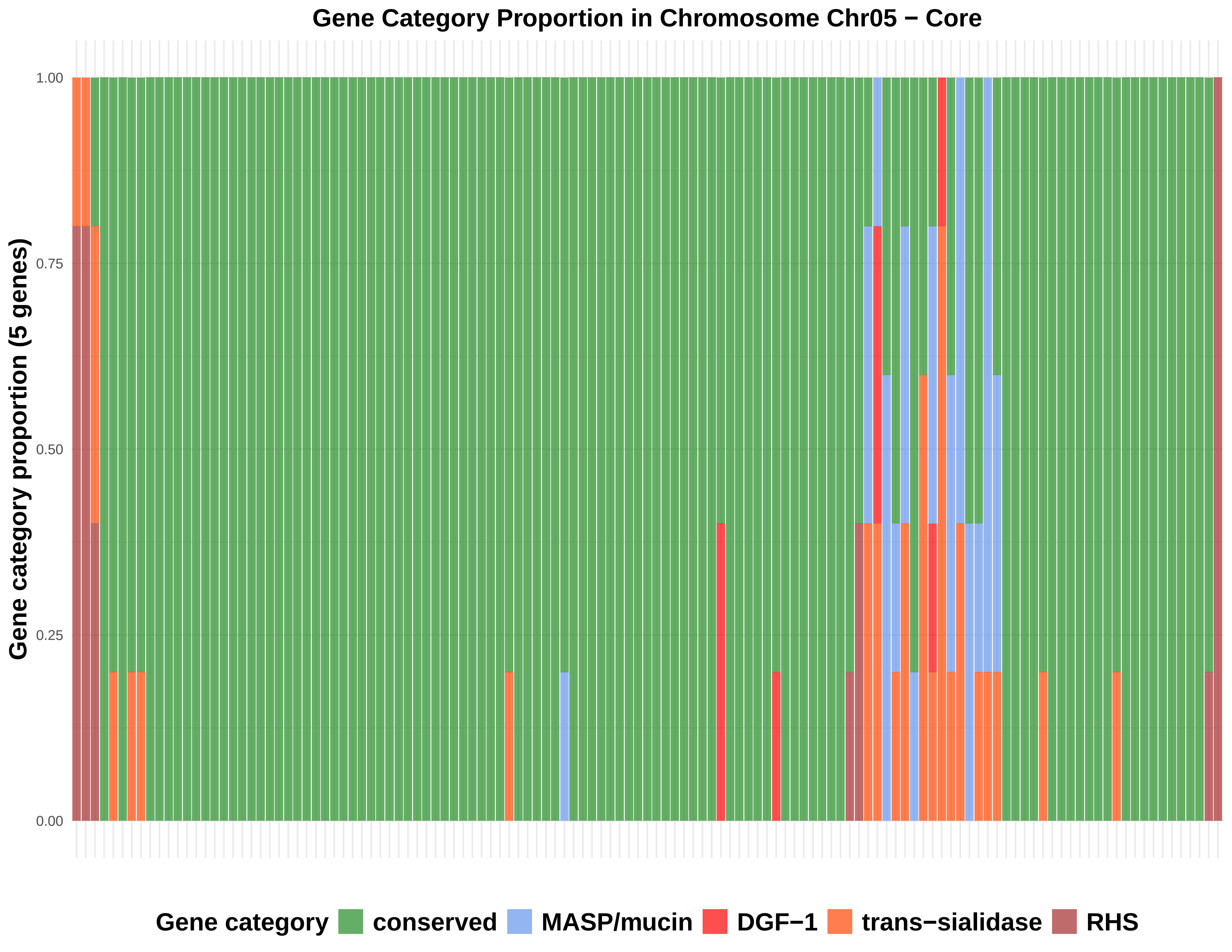
Gene Category Proportion in Chromosome Chr05 − Core
Gene category proportion (5 genes)
1.00
0.75
0.50
0.25
0.00
Gene category
conserved
MASP/mucin
DGF−1
trans−sialidase
RHS

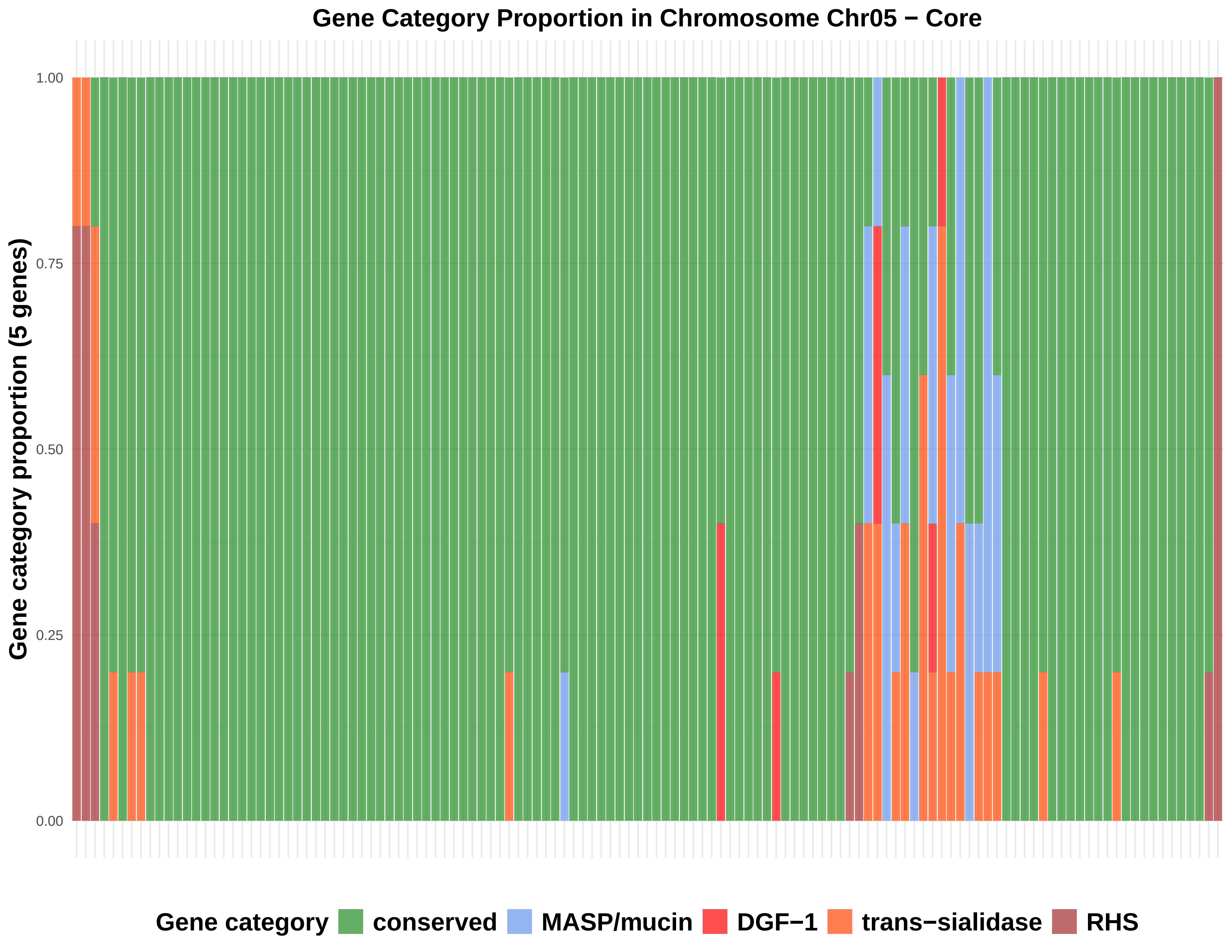

Gene Category Proportion in Chromosome Chr05 − Core
Gene category proportion (5 genes)
1.00
0.75
0.50
0.25
0.00
Gene category
conserved
MASP/mucin
DGF−1
trans−sialidase
RHS

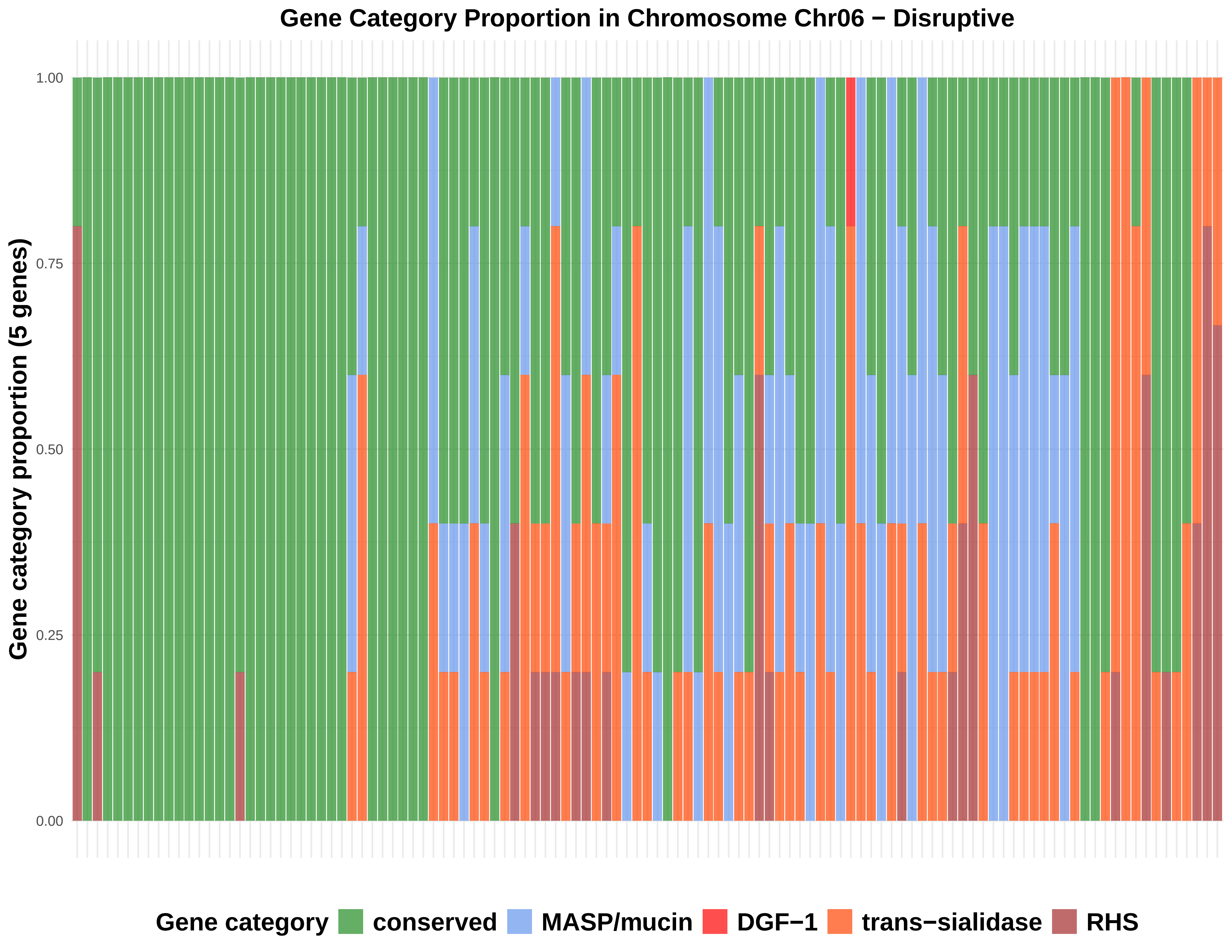

Gene Category Proportion in Chromosome Chr06 − Disruptive
Gene category proportion (5 genes)
1.00
0.75
0.50
0.25
0.00
Gene category
conserved
MASP/mucin
DGF−1
trans−sialidase
RHS

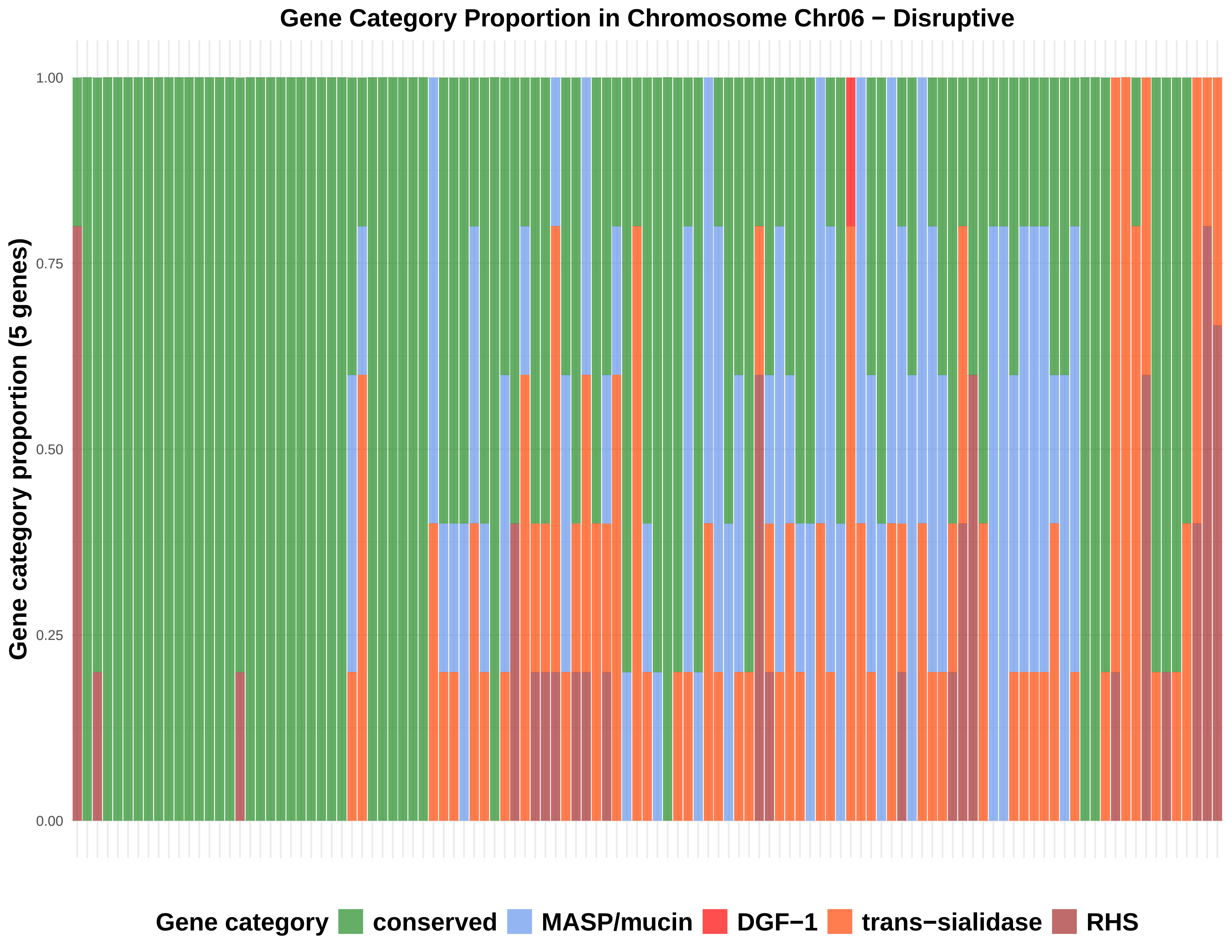

Gene Category Proportion in Chromosome Chr06 − Disruptive
Gene category proportion (5 genes)
1.00
0.75
0.50
0.25
0.00
Gene category
conserved
MASP/mucin
DGF−1
trans−sialidase
RHS

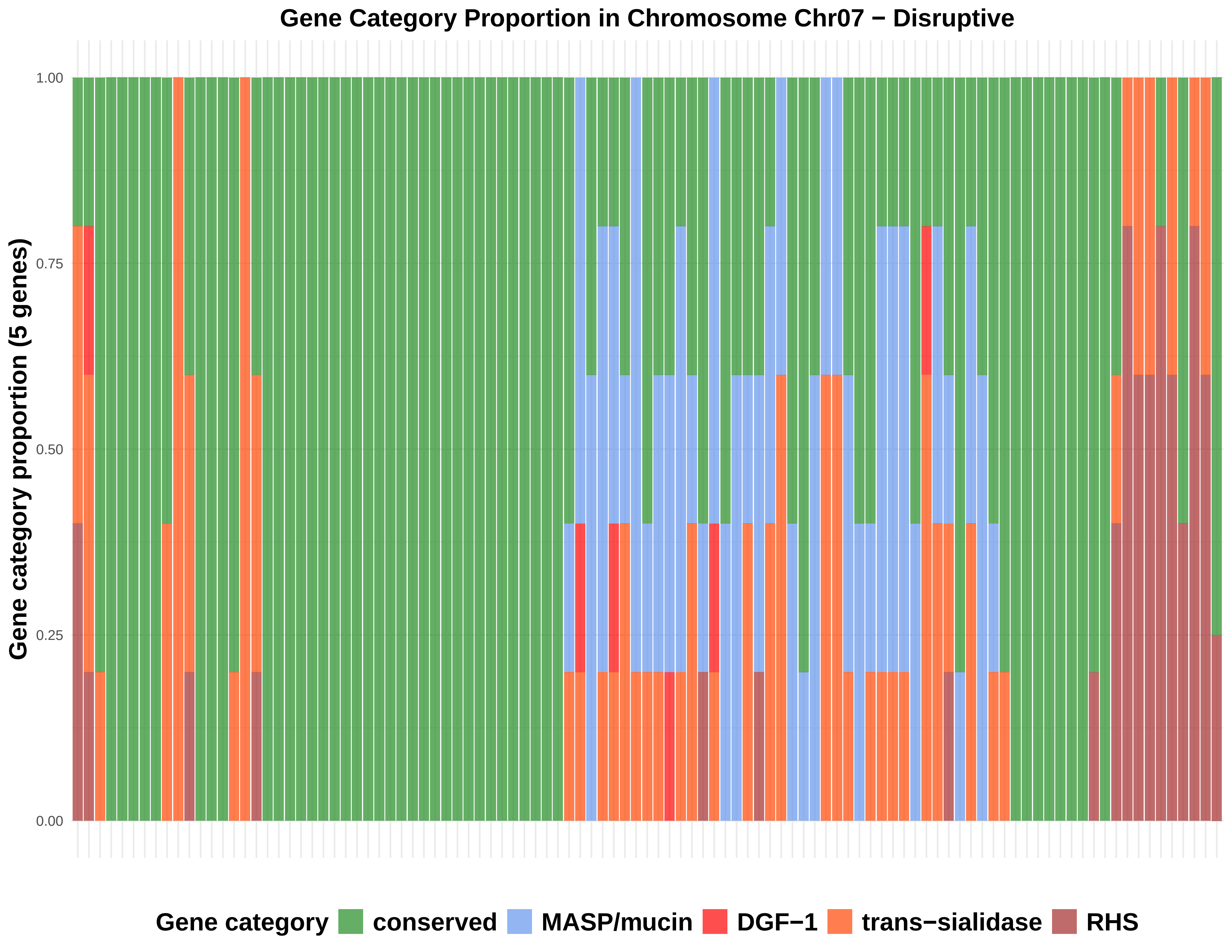
Gene Category Proportion in Chromosome Chr07 − Disruptive
Gene category proportion (5 genes)
1.00
0.75
0.50
0.25
0.00
Gene category
conserved
MASP/mucin
DGF−1
trans−sialidase
RHS

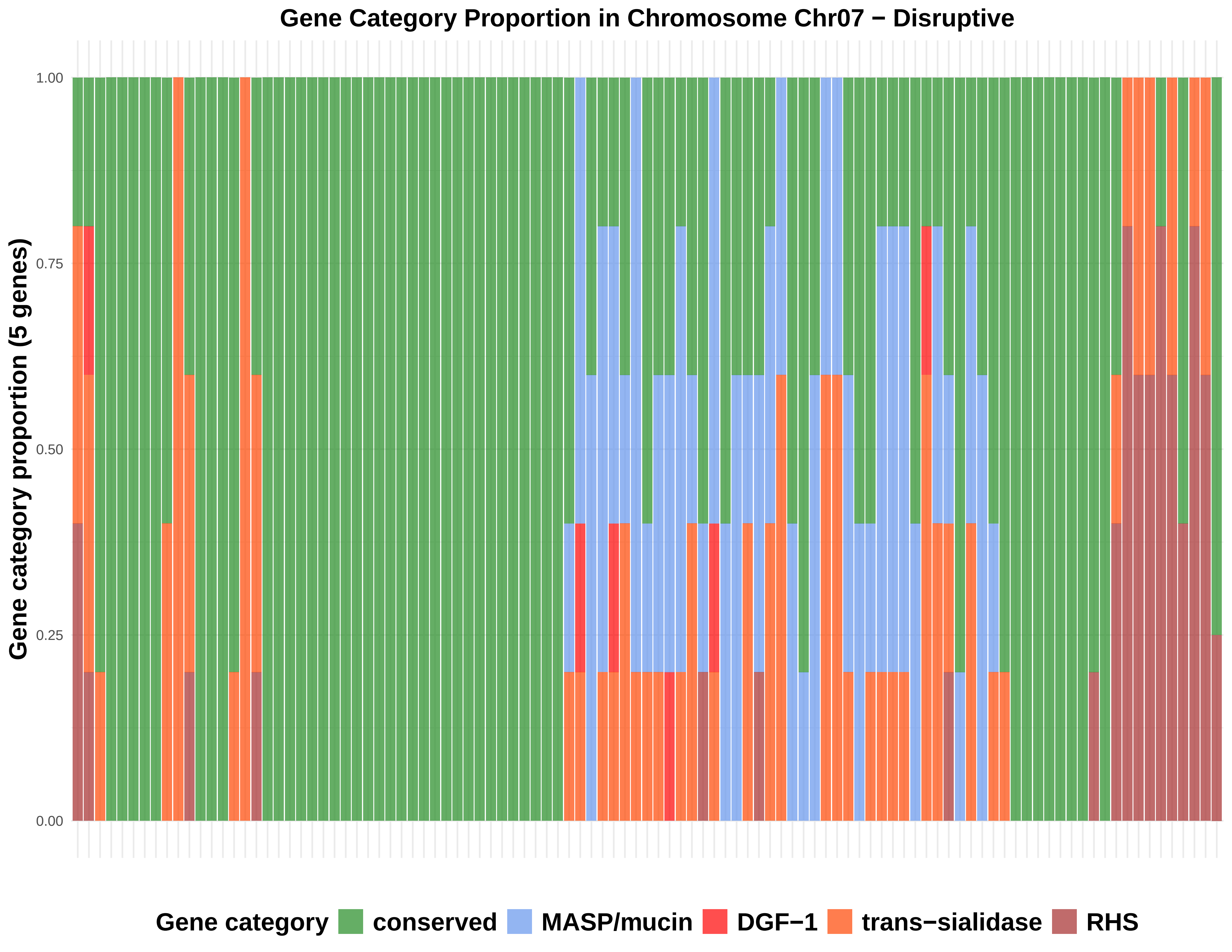

Gene Category Proportion in Chromosome Chr07 − Disruptive
Gene category proportion (5 genes)
1.00
0.75
0.50
0.25
0.00
Gene category
conserved
MASP/mucin
DGF−1
trans−sialidase
RHS

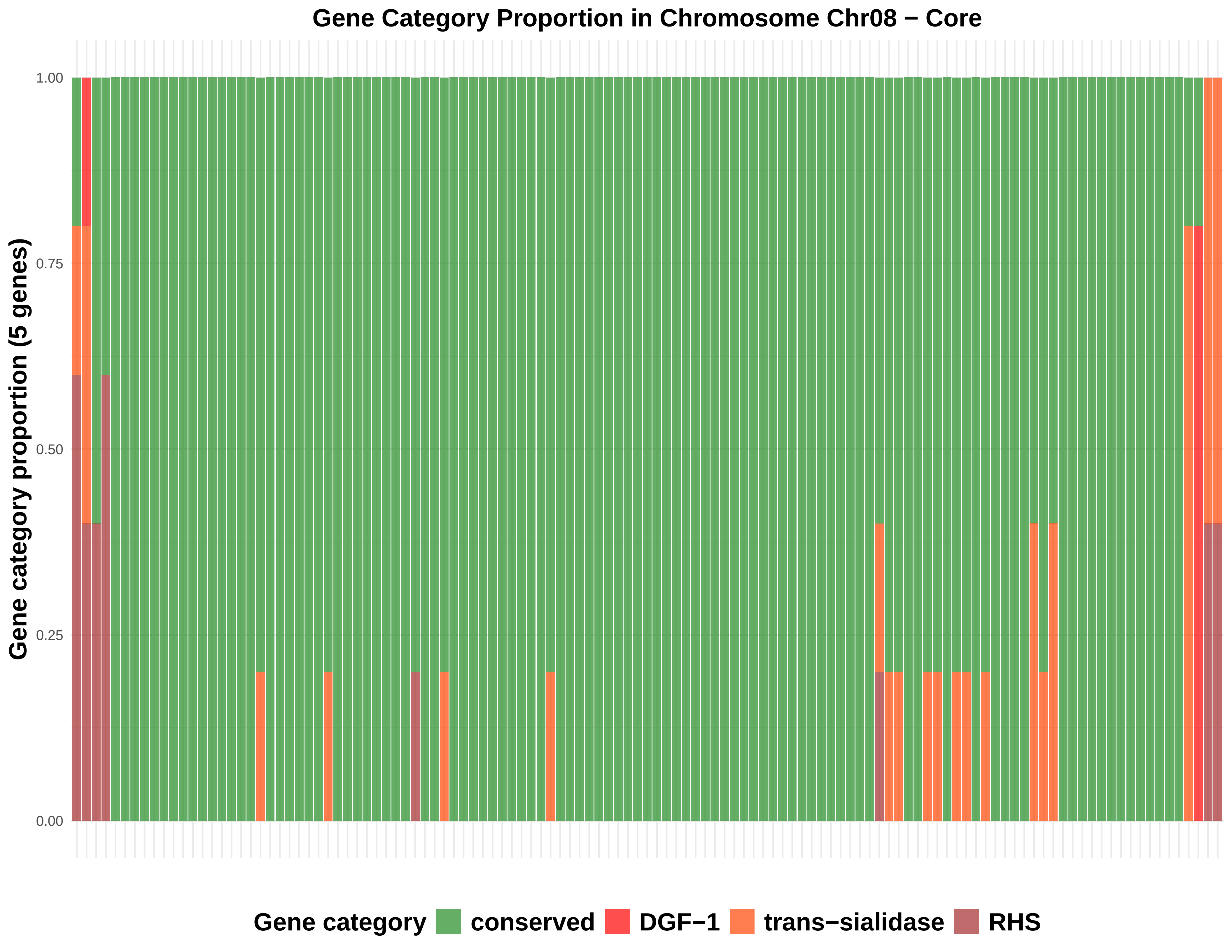
Gene Category Proportion in Chromosome Chr08 − Core
Gene category proportion (5 genes)
1.00
0.75
0.50
0.25
0.00
Gene category
conserved
DGF−1
trans−sialidase
RHS

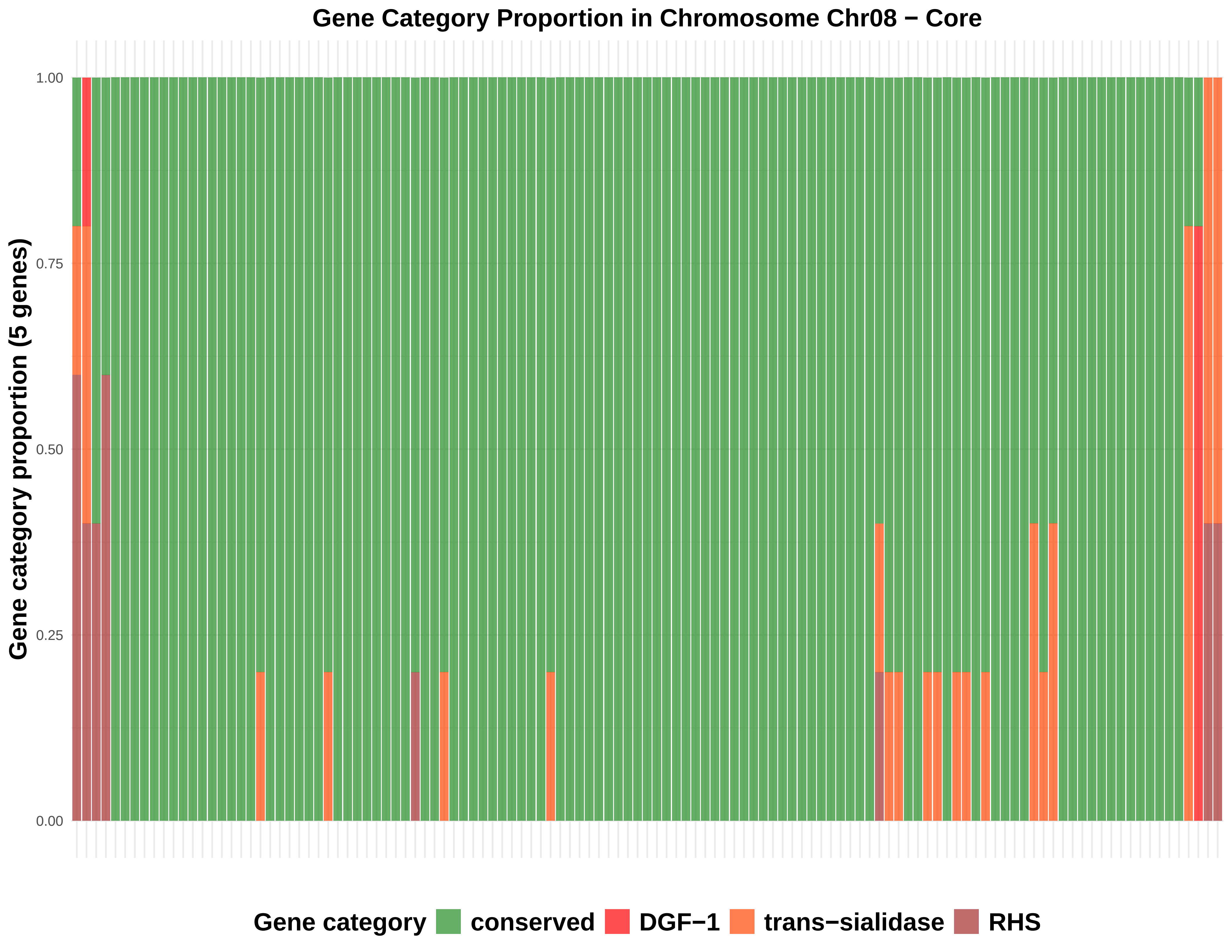

Gene Category Proportion in Chromosome Chr08 − Core
Gene category proportion (5 genes)
1.00
0.75
0.50
0.25
0.00
Gene category
conserved
DGF−1
trans−sialidase
RHS

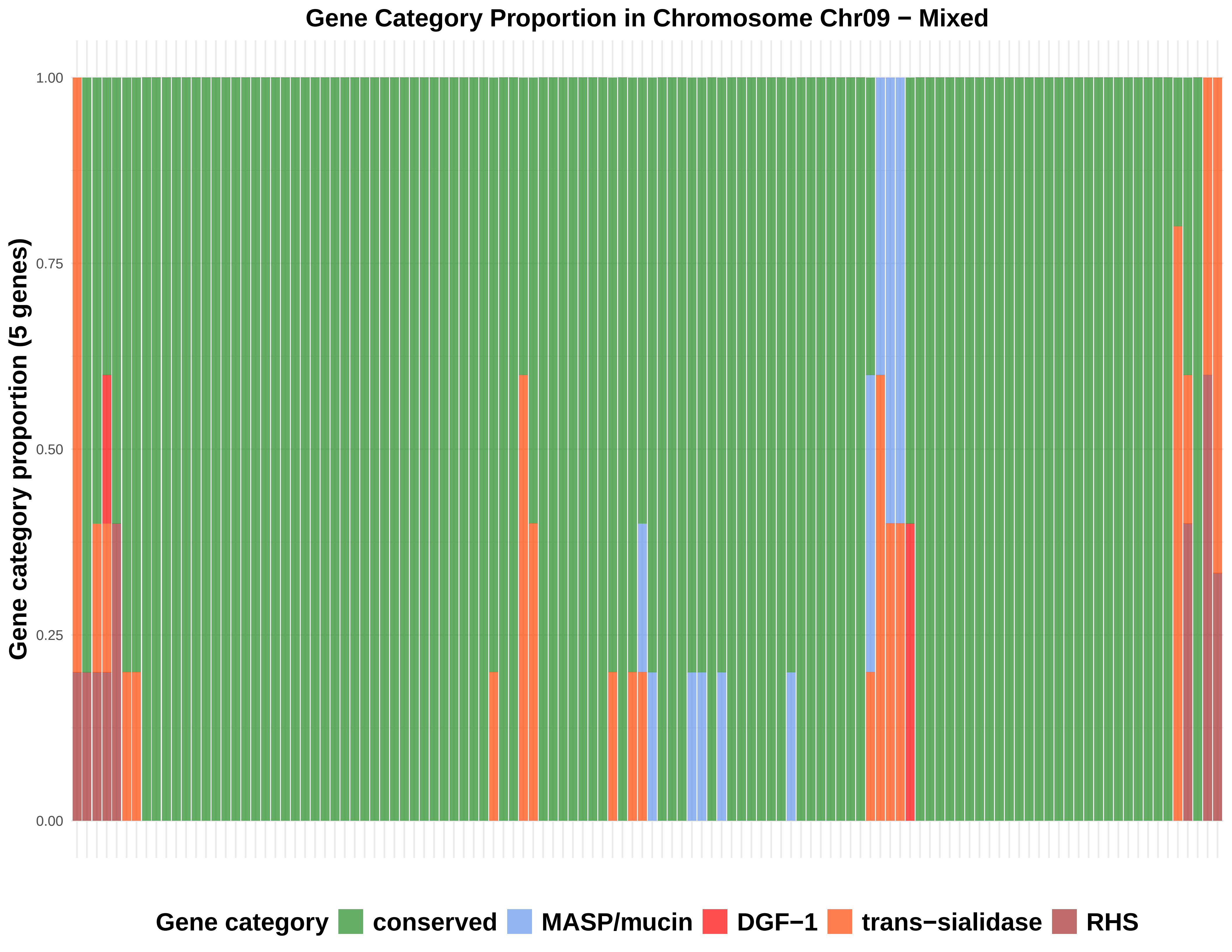

Gene Category Proportion in Chromosome Chr09 − Mixed
Gene category proportion (5 genes)
1.00
0.75
0.50
0.25
0.00
Gene category
conserved
MASP/mucin
DGF−1
trans−sialidase
RHS

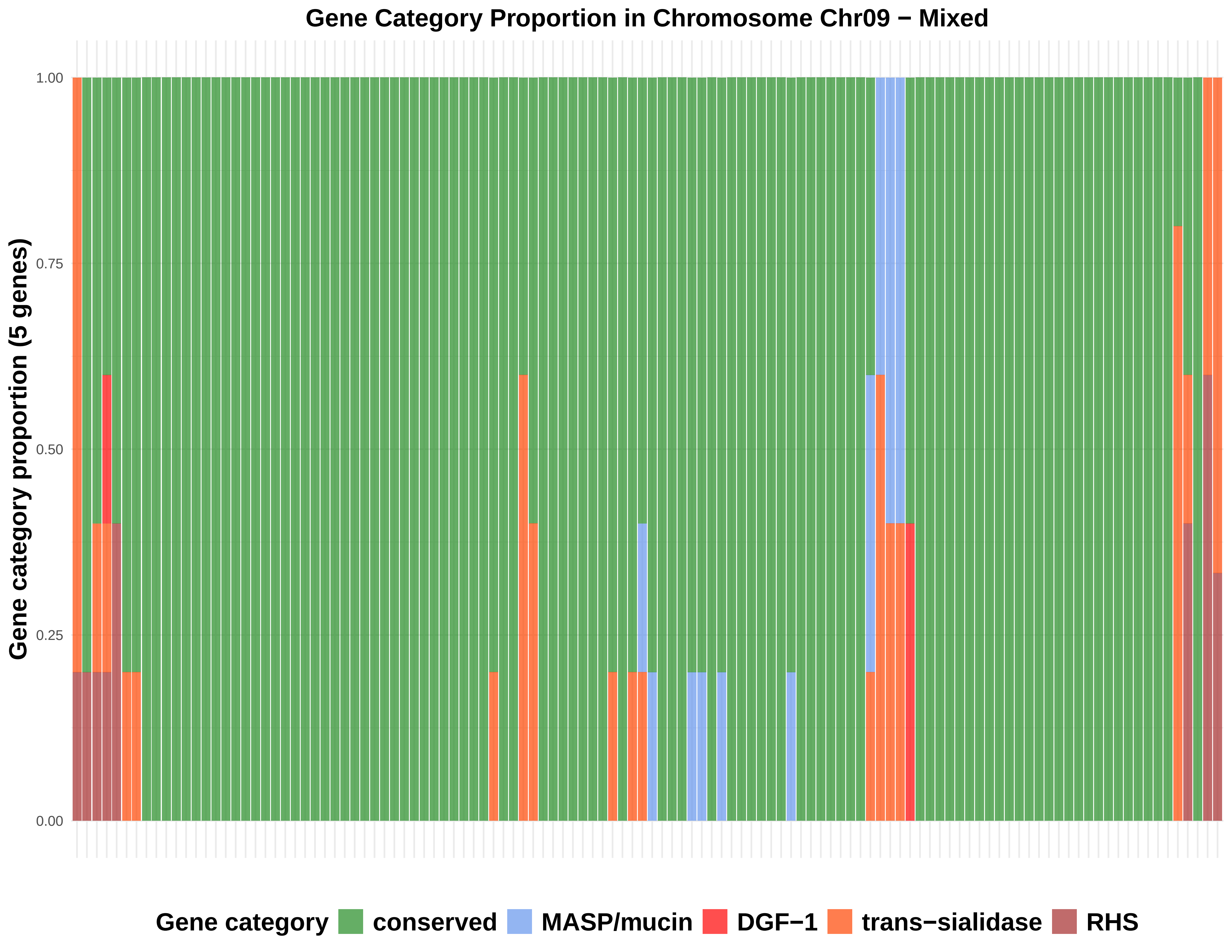

Gene Category Proportion in Chromosome Chr09 − Mixed
Gene category proportion (5 genes)
1.00
0.75
0.50
0.25
0.00
Gene category
conserved
MASP/mucin
DGF−1
trans−sialidase
RHS

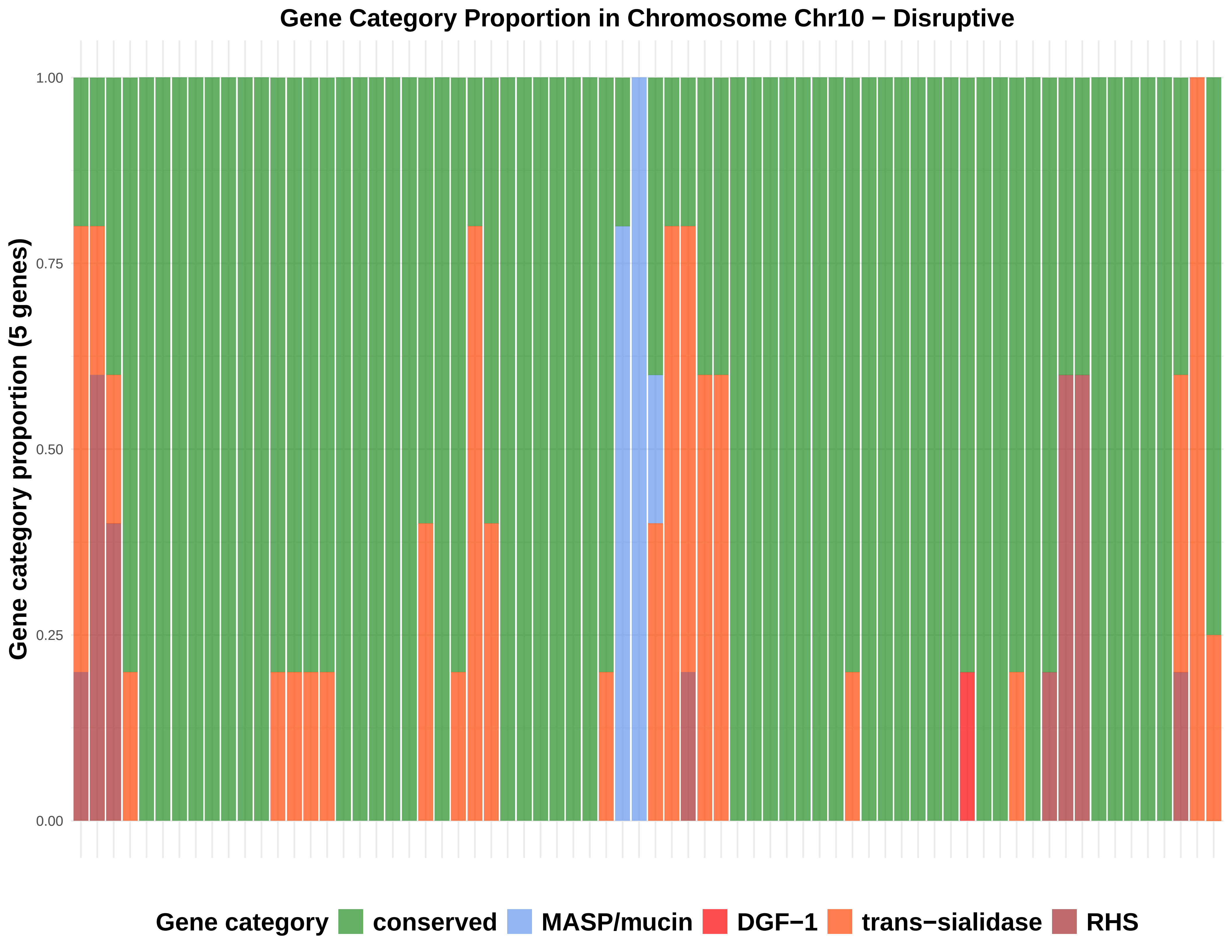

Gene Category Proportion in Chromosome Chr10 − Disruptive
Gene category proportion (5 genes)
1.00
0.75
0.50
0.25
0.00
Gene category
conserved
MASP/mucin
DGF−1
trans−sialidase
RHS

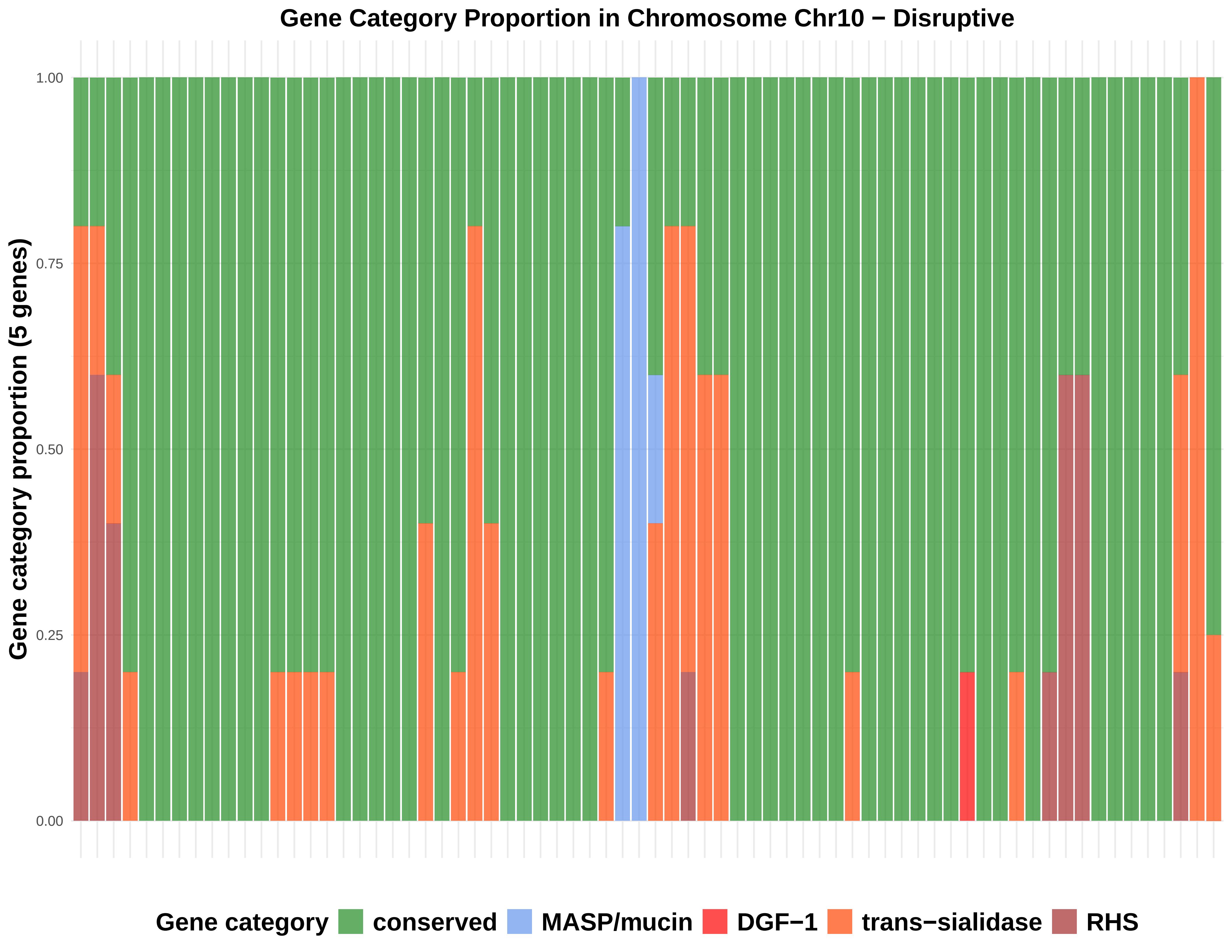
Gene Category Proportion in Chromosome Chr10 − Disruptive
Gene category proportion (5 genes)
1.00
0.75
0.50
0.25
0.00
Gene category
conserved
MASP/mucin
DGF−1
trans−sialidase
RHS

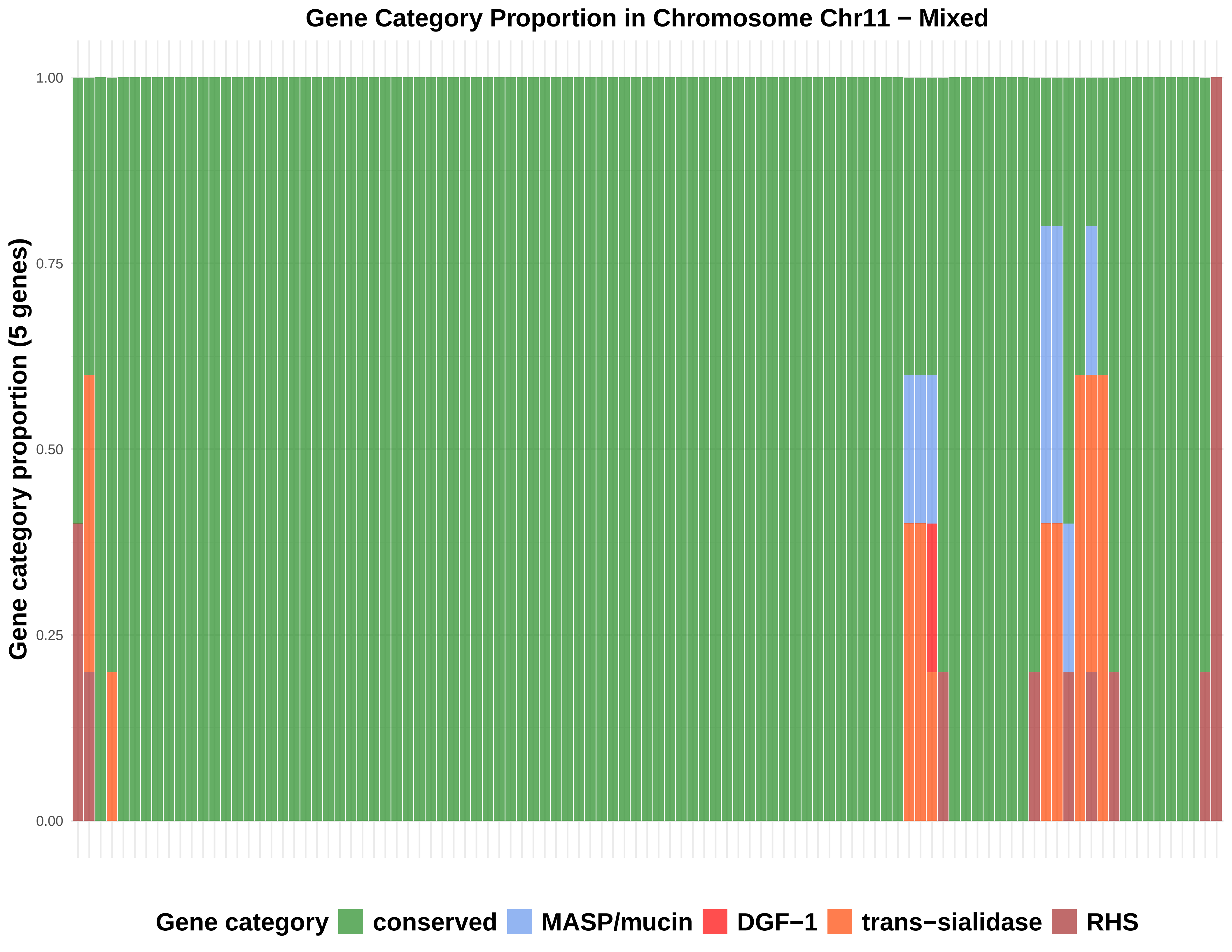
Gene Category Proportion in Chromosome Chr11 − Mixed
Gene category proportion (5 genes)
1.00
0.75
0.50
0.25
0.00
Gene category
conserved
MASP/mucin
DGF−1
trans−sialidase
RHS

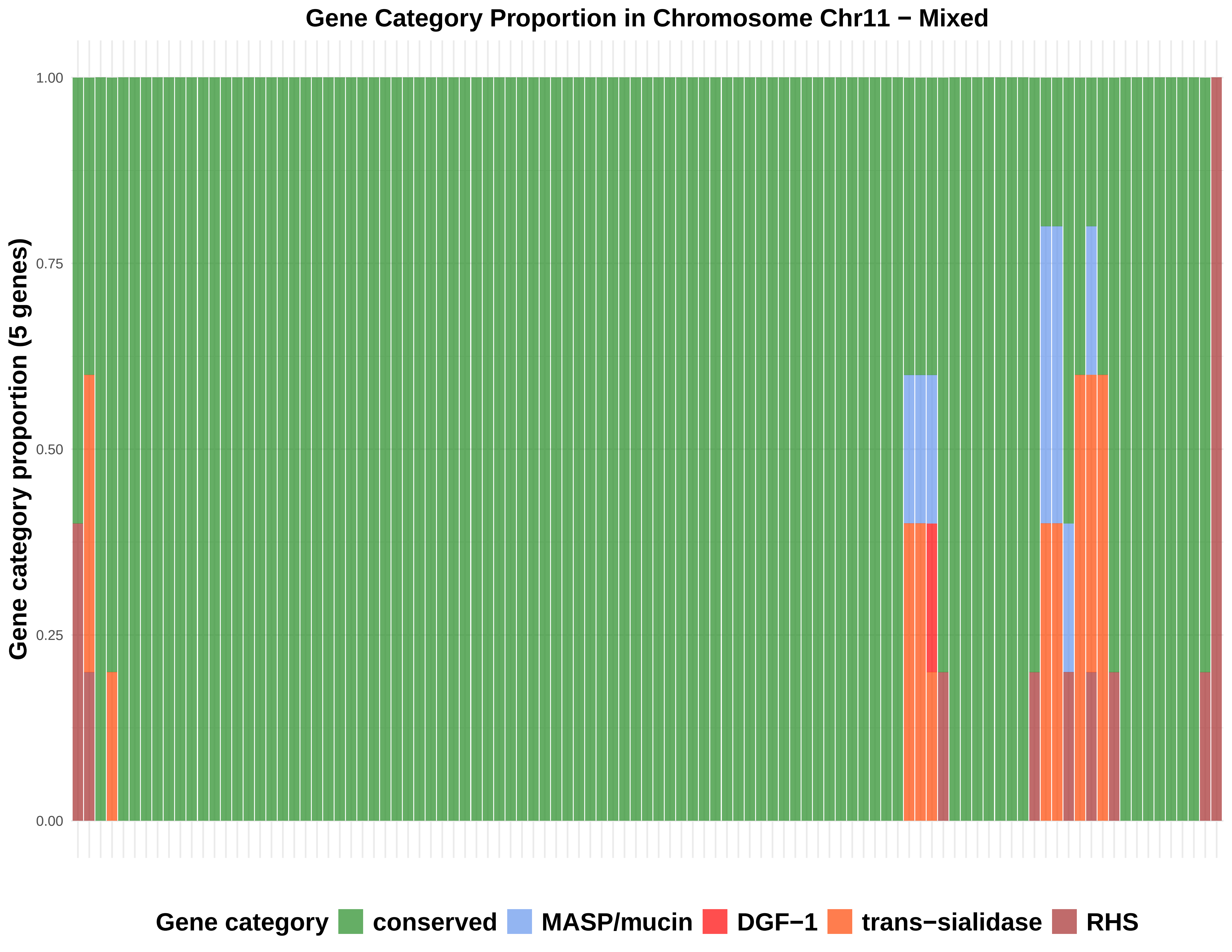
Gene Category Proportion in Chromosome Chr11 − Mixed
Gene category proportion (5 genes)
1.00
0.75
0.50
0.25
0.00
Gene category
conserved
MASP/mucin
DGF−1
trans−sialidase
RHS

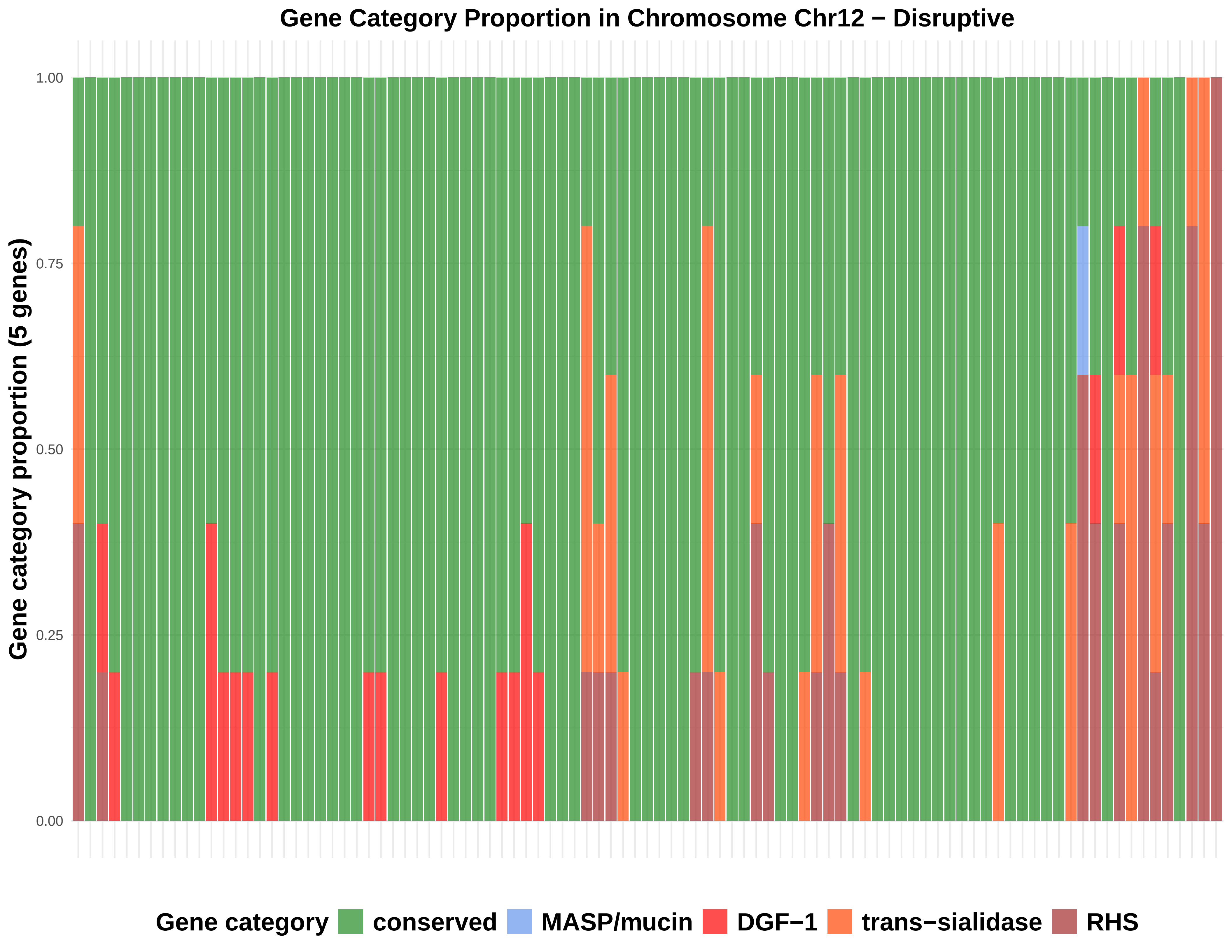
Gene Category Proportion in Chromosome Chr12 − Disruptive
Gene category proportion (5 genes)
1.00
0.75
0.50
0.25
0.00
Gene category
conserved
MASP/mucin
DGF−1
trans−sialidase
RHS

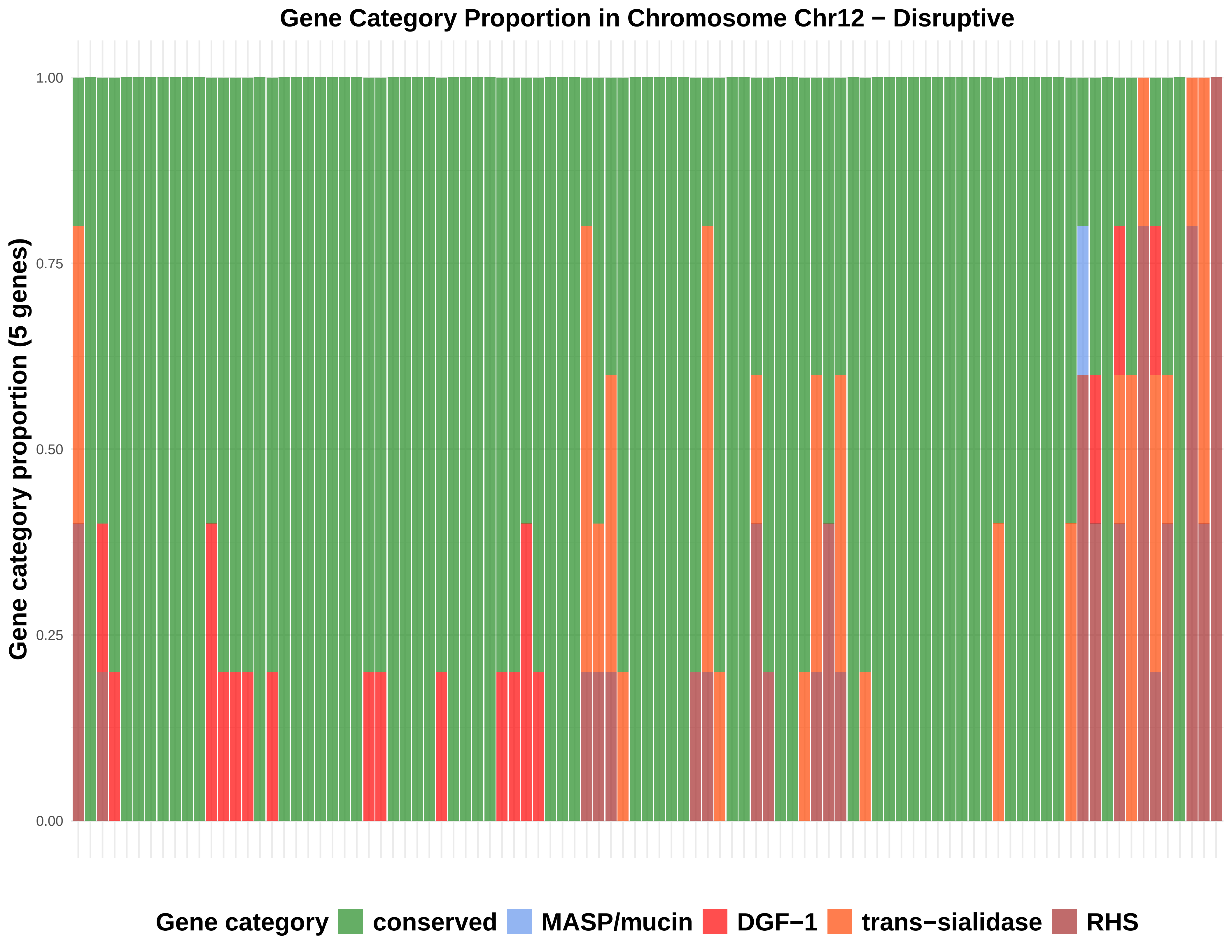

Gene Category Proportion in Chromosome Chr12 − Disruptive
Gene category proportion (5 genes)
1.00
0.75
0.50
0.25
0.00
Gene category
conserved
MASP/mucin
DGF−1
trans−sialidase
RHS

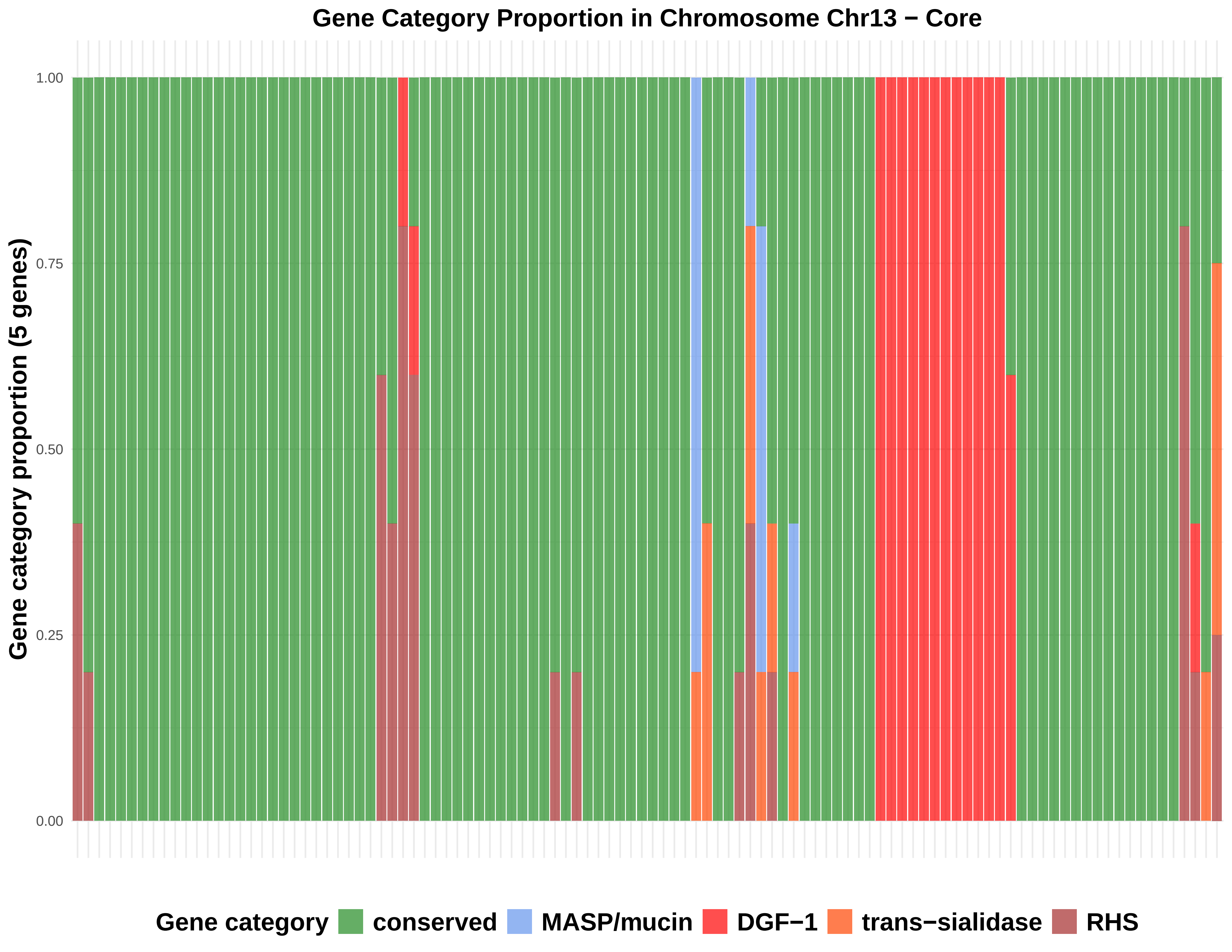

Gene Category Proportion in Chromosome Chr13 − Core
Gene category proportion (5 genes)
1.00
0.75
0.50
0.25
0.00
Gene category
conserved
MASP/mucin
DGF−1
trans−sialidase
RHS

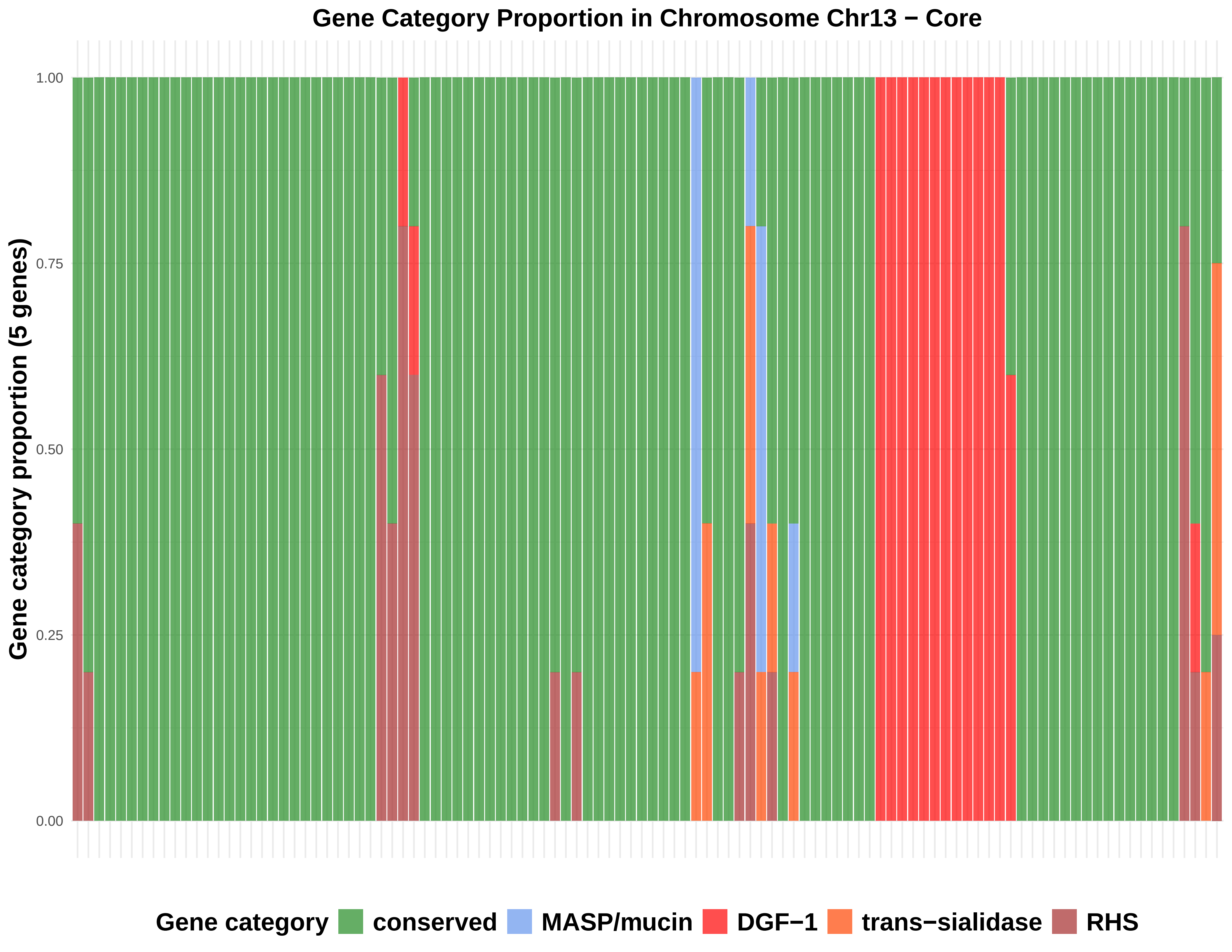

Gene Category Proportion in Chromosome Chr13 − Core
Gene category proportion (5 genes)
1.00
0.75
0.50
0.25
0.00
Gene category
conserved
MASP/mucin
DGF−1
trans−sialidase
RHS

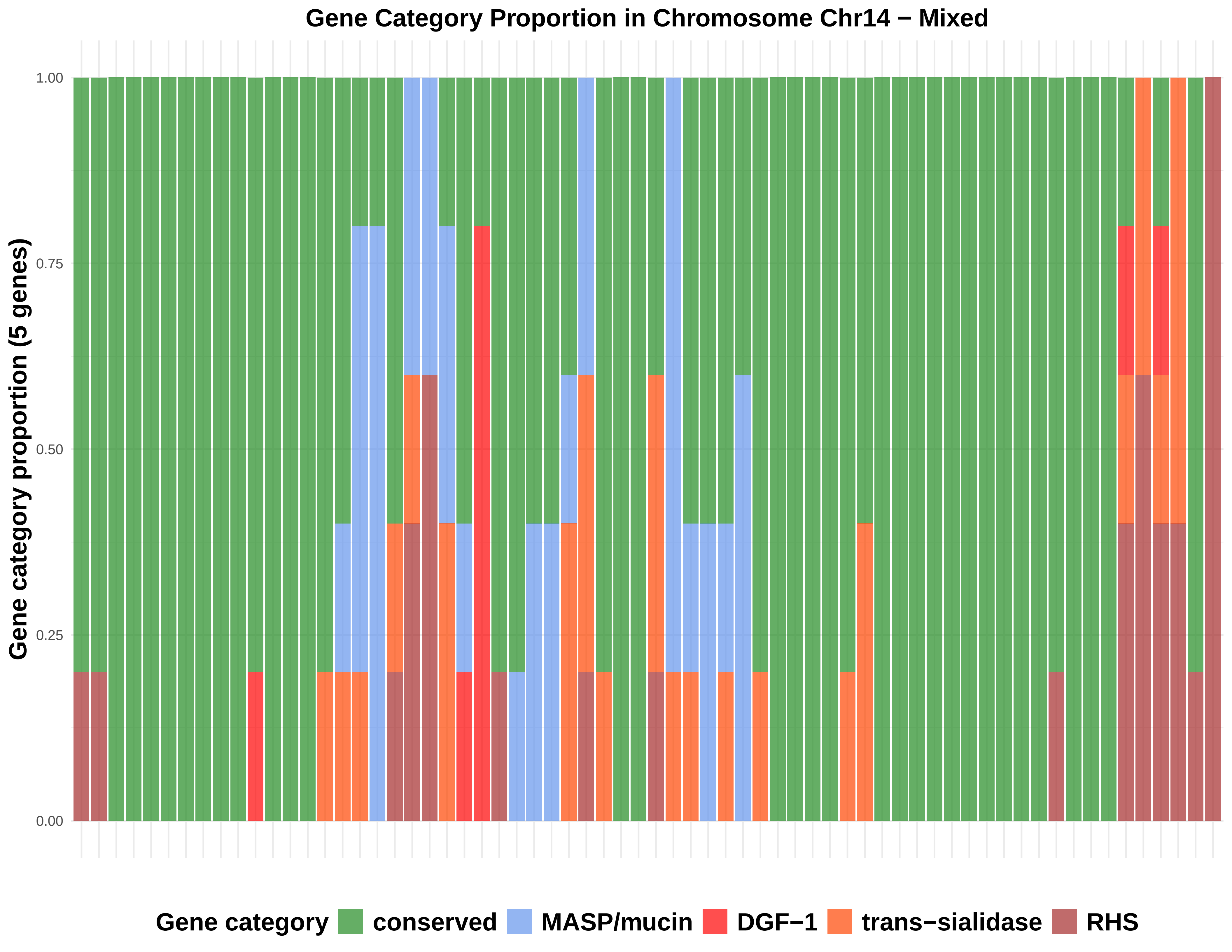

Gene Category Proportion in Chromosome Chr14 − Mixed
Gene category proportion (5 genes)
1.00
0.75
0.50
0.25
0.00
Gene category
conserved
MASP/mucin
DGF−1
trans−sialidase
RHS

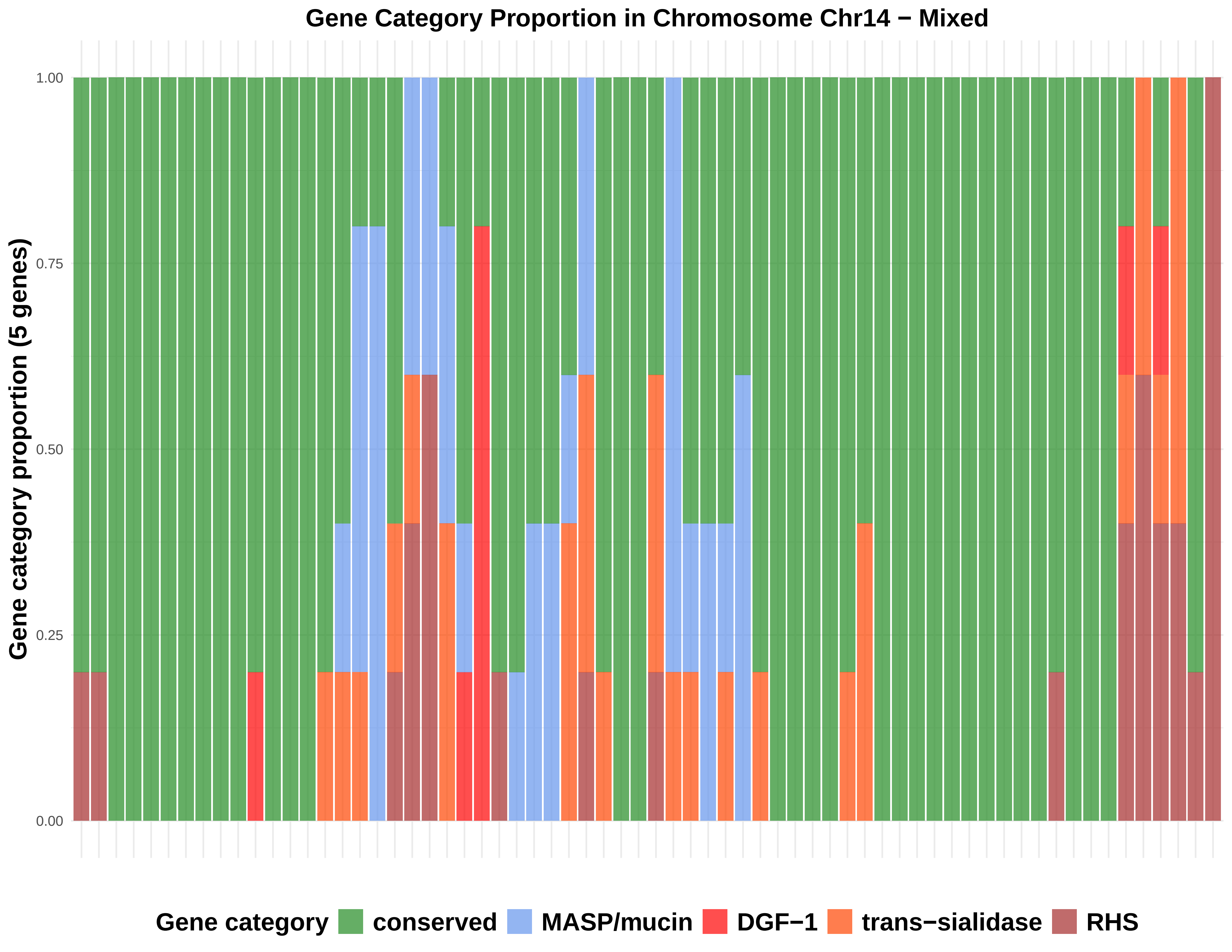
Gene Category Proportion in Chromosome Chr14 − Mixed
Gene category proportion (5 genes)
1.00
0.75
0.50
0.25
0.00
Gene category
conserved
MASP/mucin
DGF−1
trans−sialidase
RHS

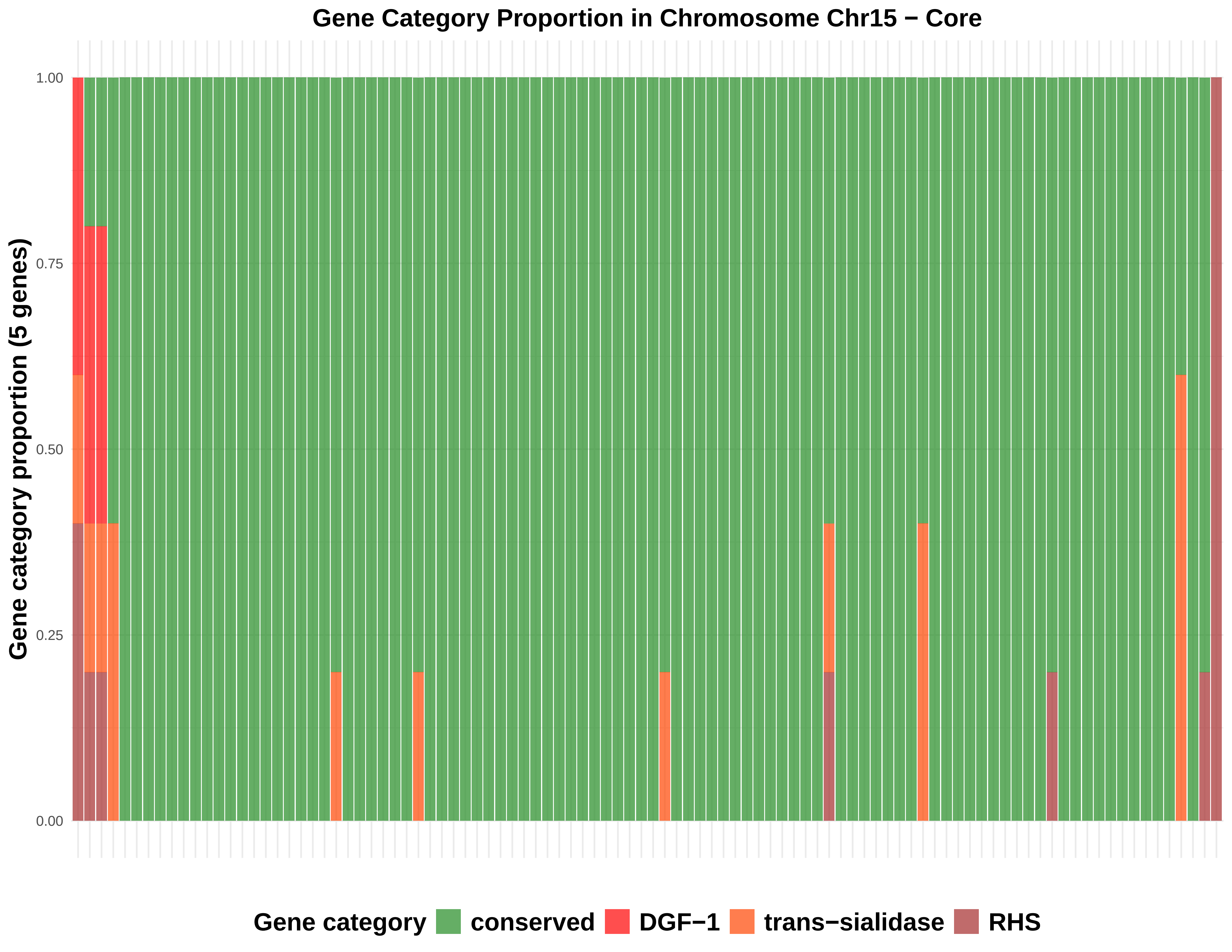
Gene Category Proportion in Chromosome Chr15 − Core
Gene category proportion (5 genes)
1.00
0.75
0.50
0.25
0.00
Gene category
conserved
DGF−1
trans−sialidase
RHS

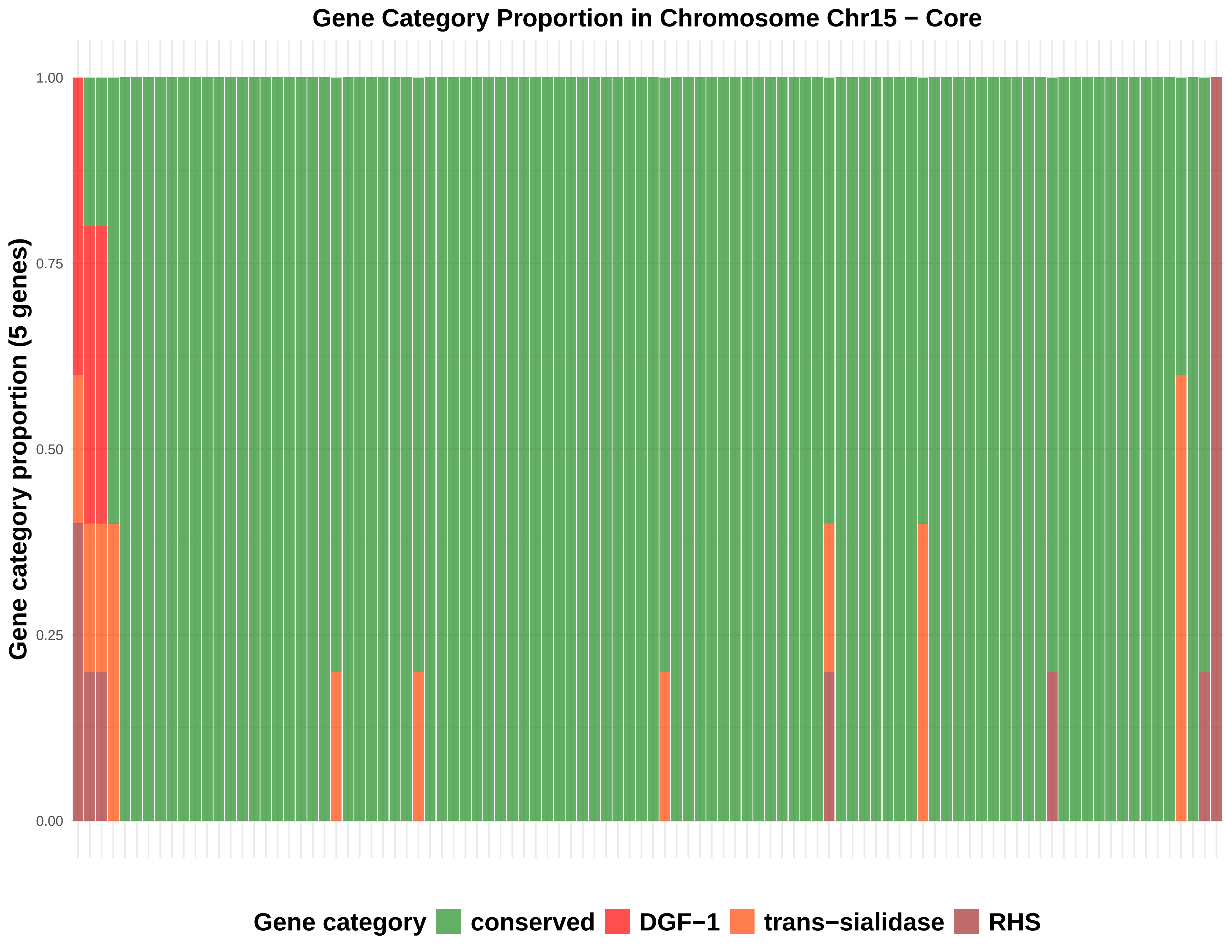

Gene Category Proportion in Chromosome Chr15 − Core
Gene category proportion (5 genes)
1.00
0.75
0.50
0.25
0.00
Gene category
conserved
DGF−1
trans−sialidase
RHS

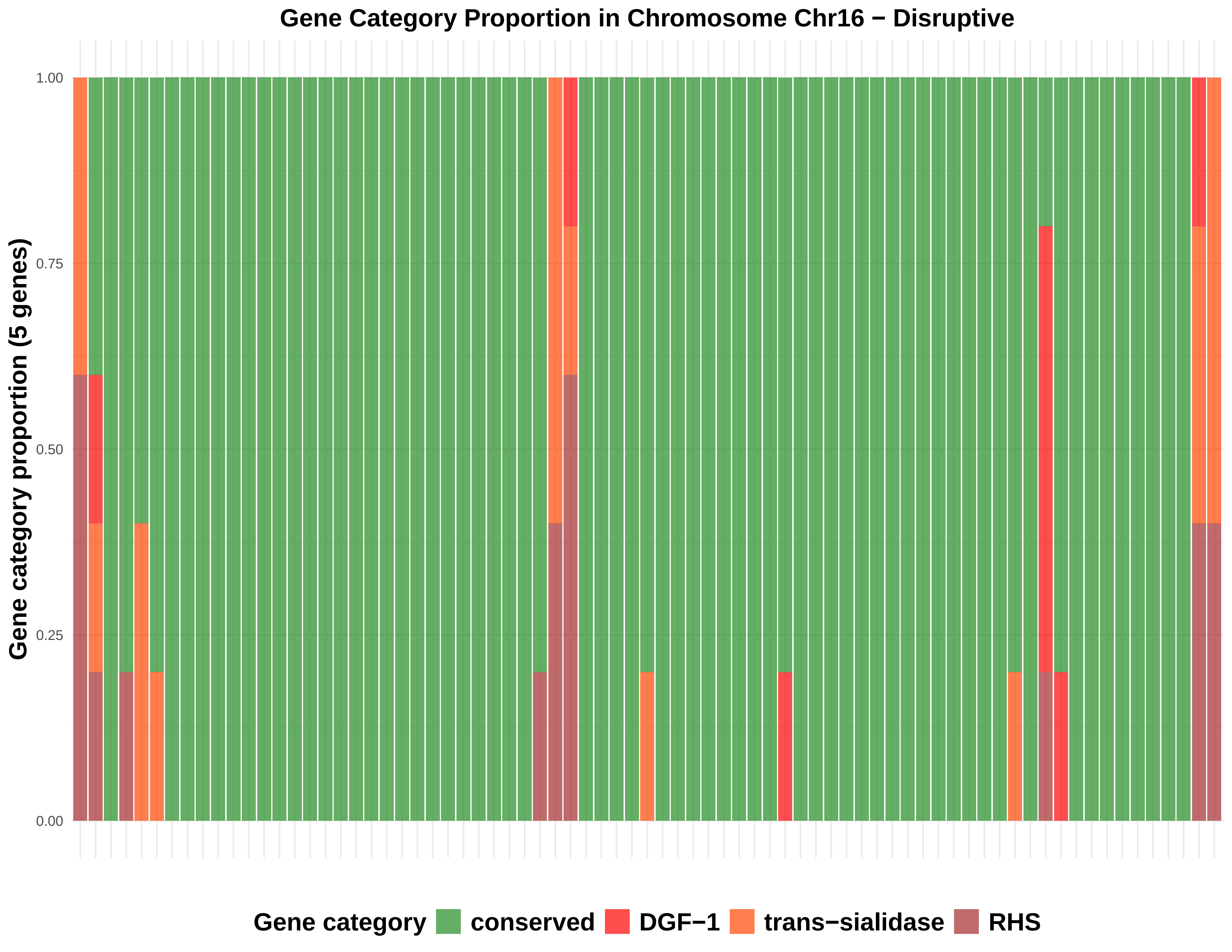

Gene Category Proportion in Chromosome Chr16 − Disruptive
Gene category proportion (5 genes)
1.00
0.75
0.50
0.25
0.00
Gene category
conserved
DGF−1
trans−sialidase
RHS

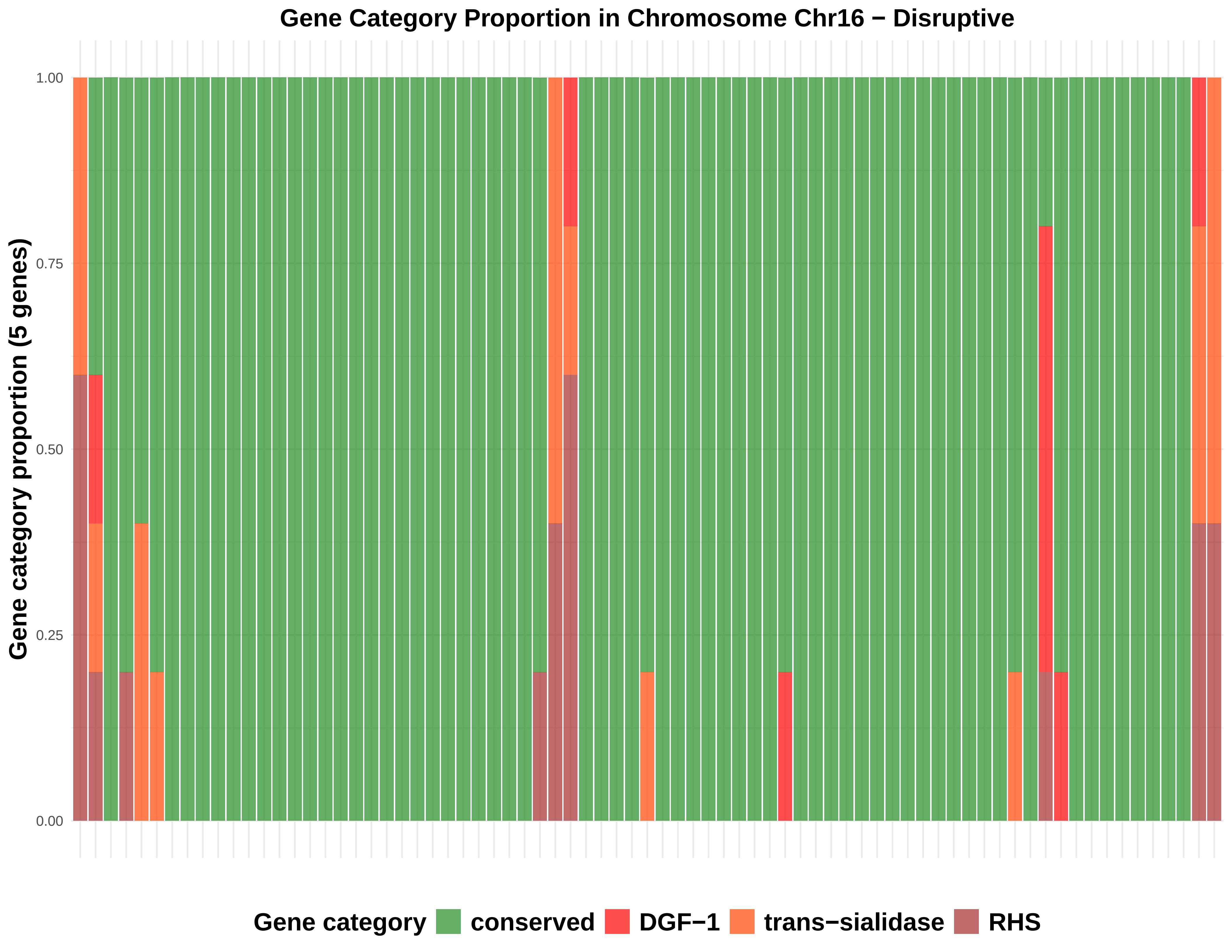
Gene Category Proportion in Chromosome Chr16 − Disruptive
Gene category proportion (5 genes)
1.00
0.75
0.50
0.25
0.00
Gene category
conserved
DGF−1
trans−sialidase
RHS

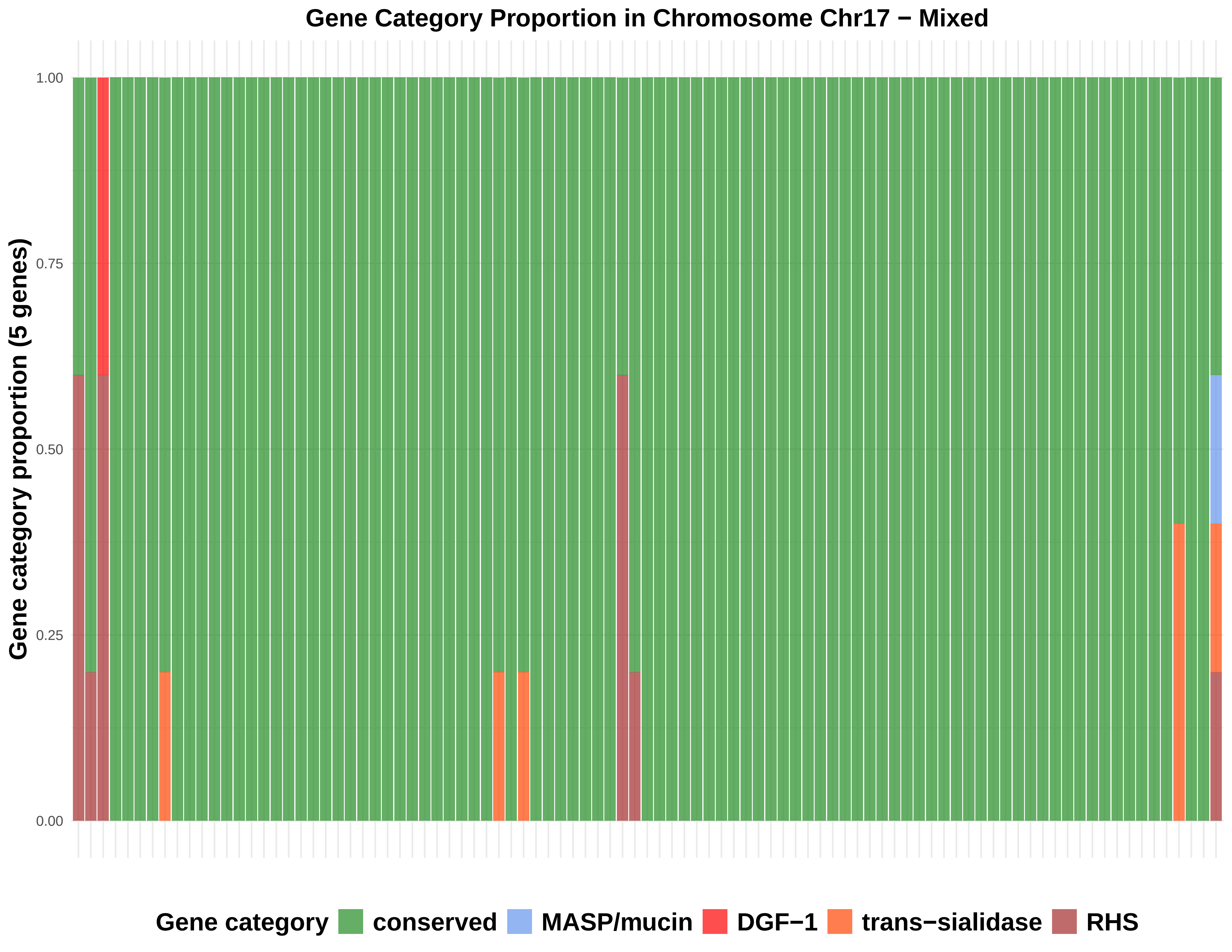
Gene Category Proportion in Chromosome Chr17 − Mixed
Gene category proportion (5 genes)
1.00
0.75
0.50
0.25
0.00
Gene category
conserved
MASP/mucin
DGF−1
trans−sialidase
RHS

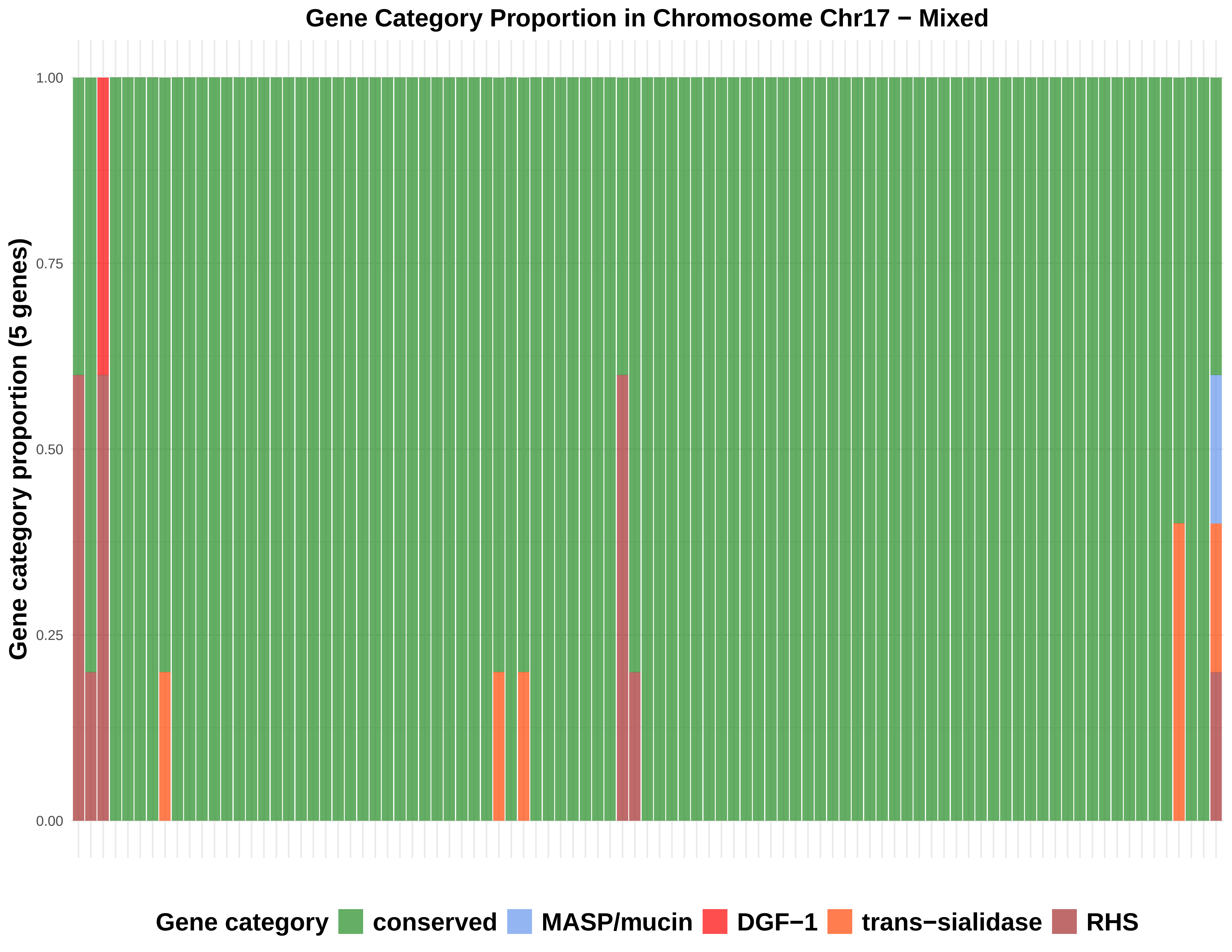

Gene Category Proportion in Chromosome Chr17 − Mixed
Gene category proportion (5 genes)
1.00
0.75
0.50
0.25
0.00
Gene category
conserved
MASP/mucin
DGF−1
trans−sialidase
RHS

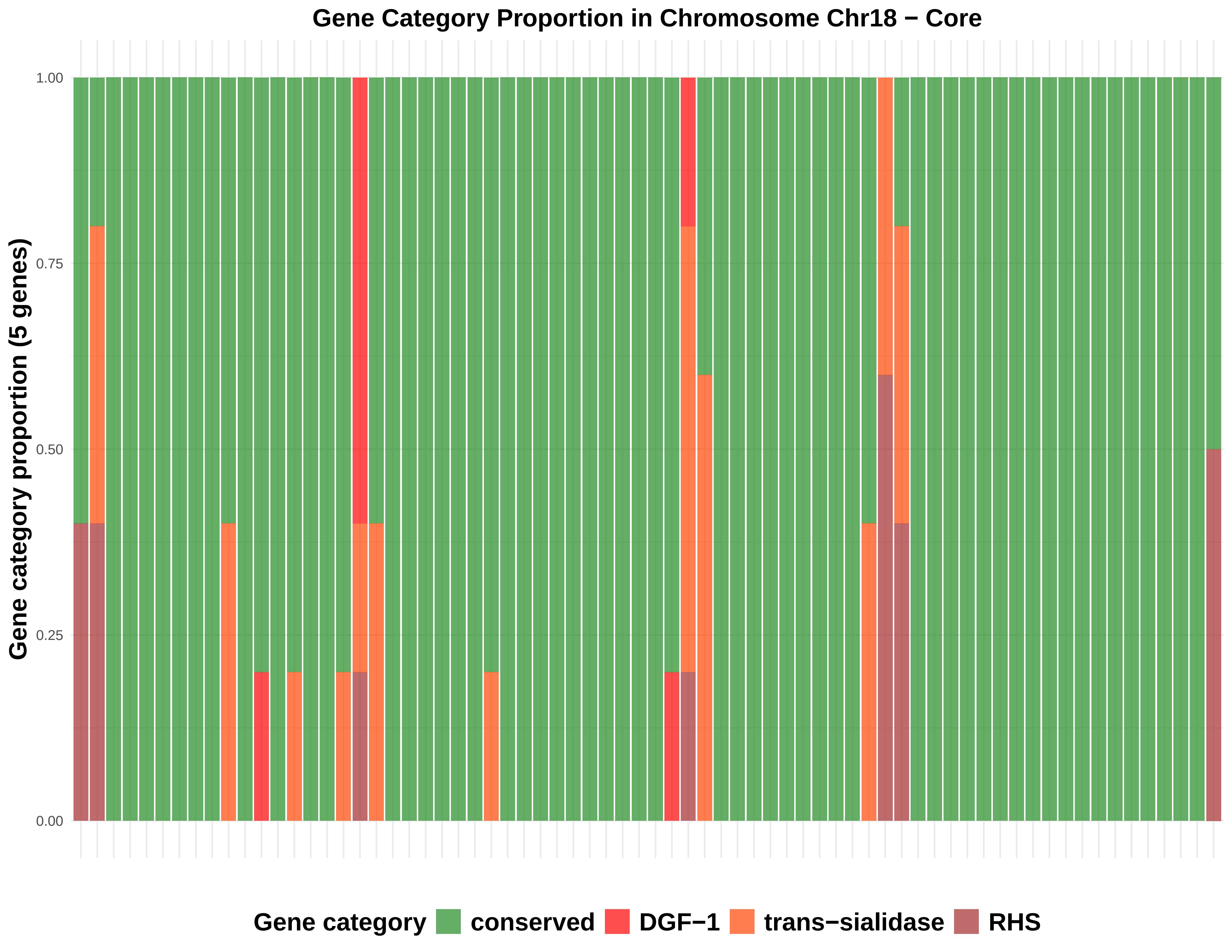
Gene Category Proportion in Chromosome Chr18 − Core
Gene category proportion (5 genes)
1.00
0.75
0.50
0.25
0.00
Gene category
conserved
DGF−1
trans−sialidase
RHS

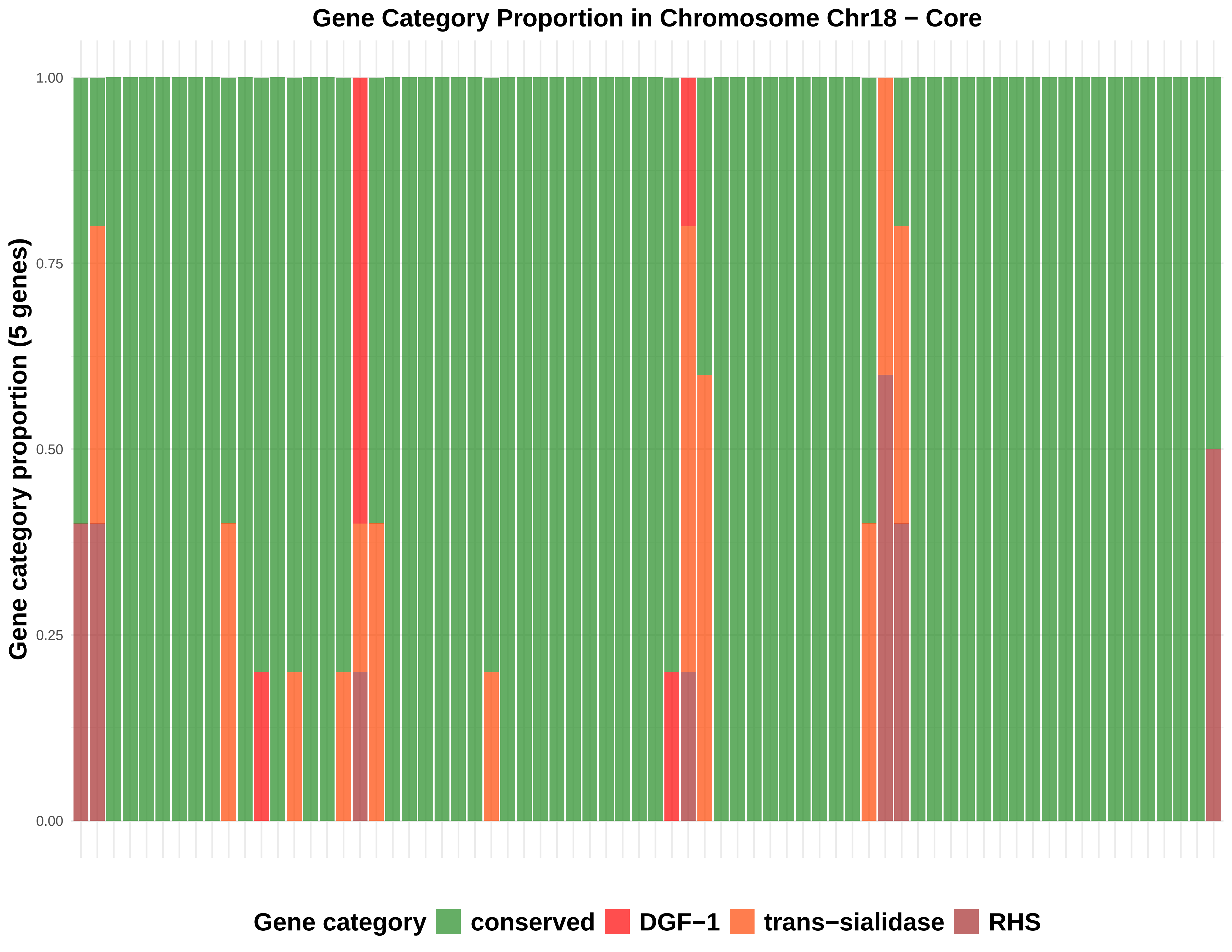
Gene Category Proportion in Chromosome Chr18 − Core
Gene category proportion (5 genes)
1.00
0.75
0.50
0.25
0.00
Gene category
conserved
DGF−1
trans−sialidase
RHS

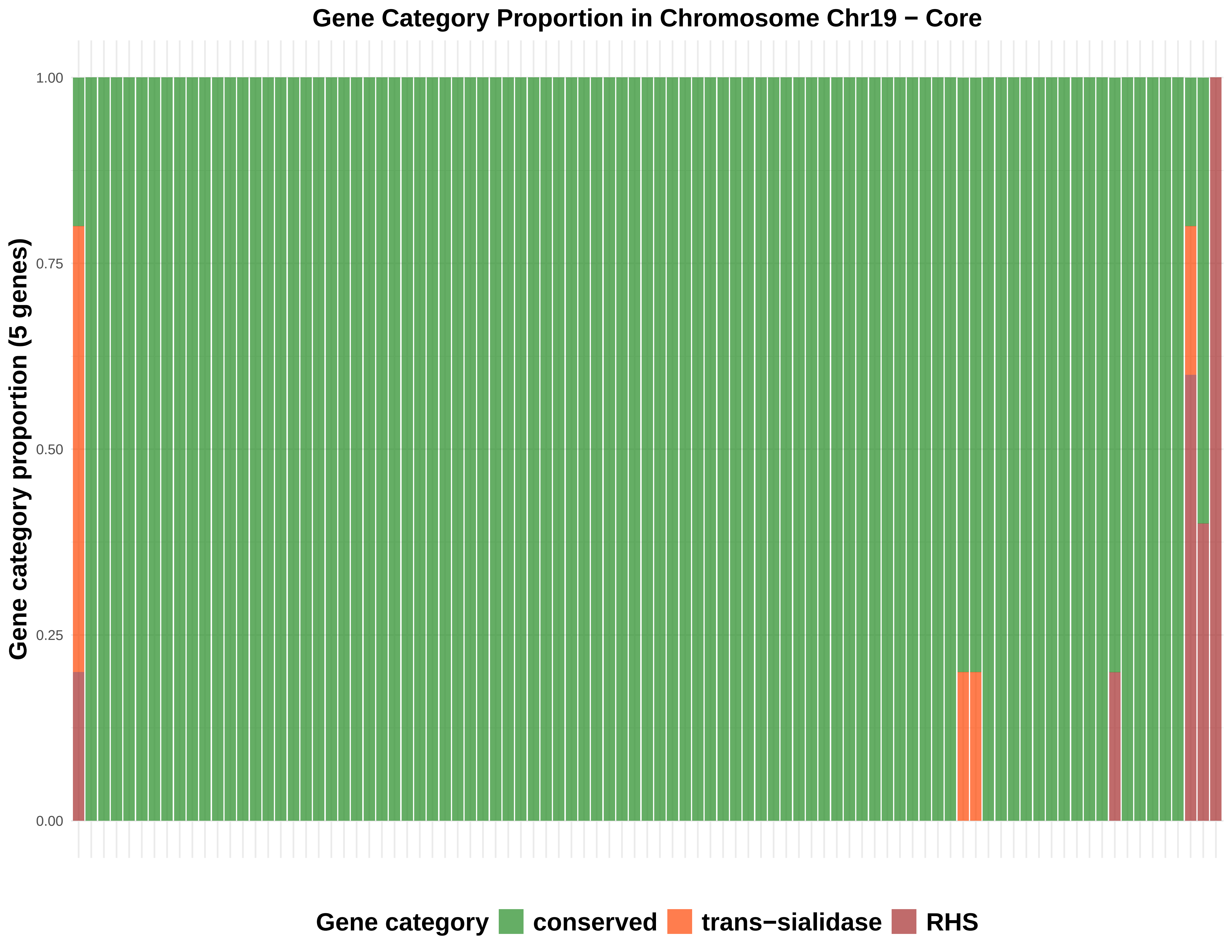

Gene Category Proportion in Chromosome Chr19 − Core
Gene category proportion (5 genes)
1.00
0.75
0.50
0.25
0.00
Gene category
conserved
trans−sialidase
RHS

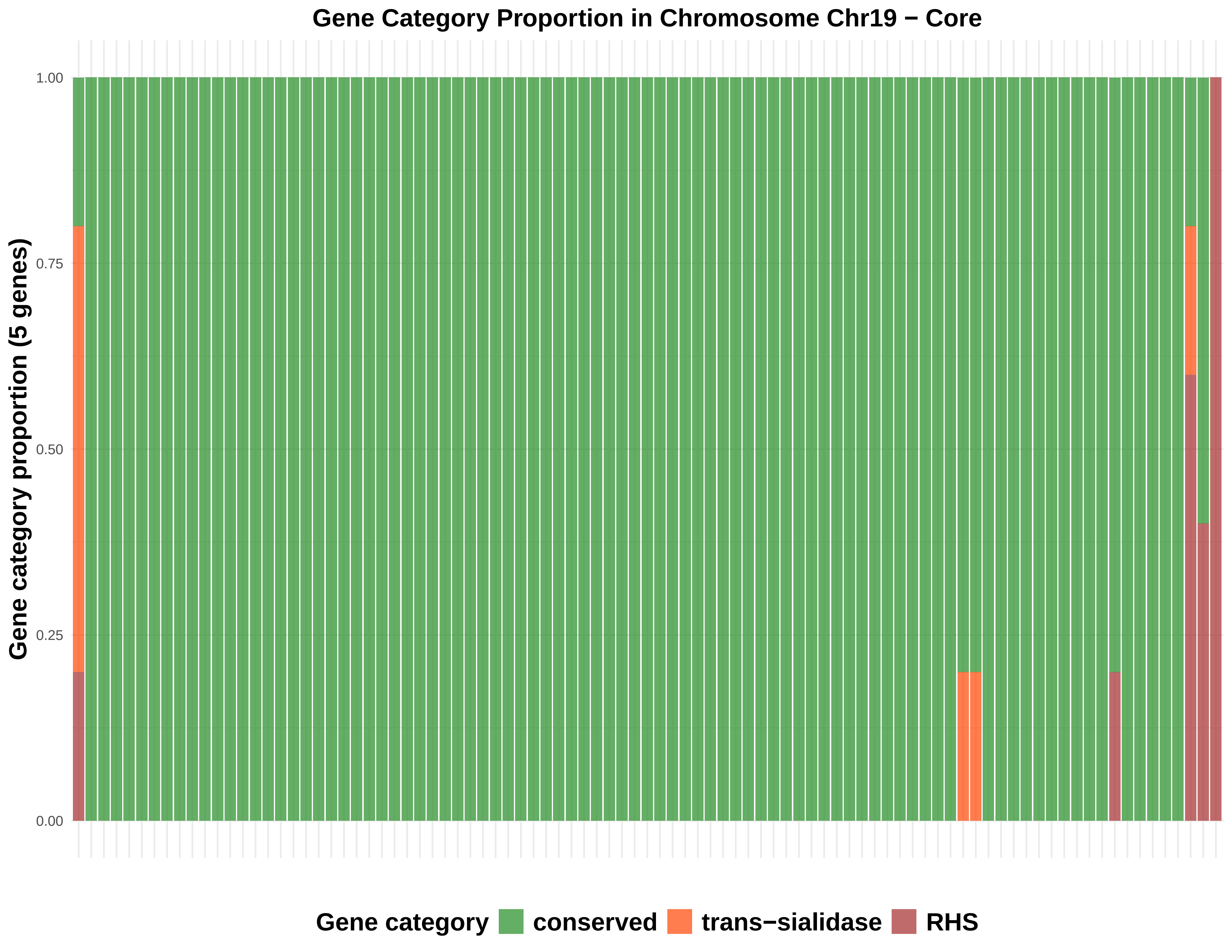
Gene Category Proportion in Chromosome Chr19 − Core
Gene category proportion (5 genes)
1.00
0.75
0.50
0.25
0.00
Gene category
conserved
trans−sialidase
RHS

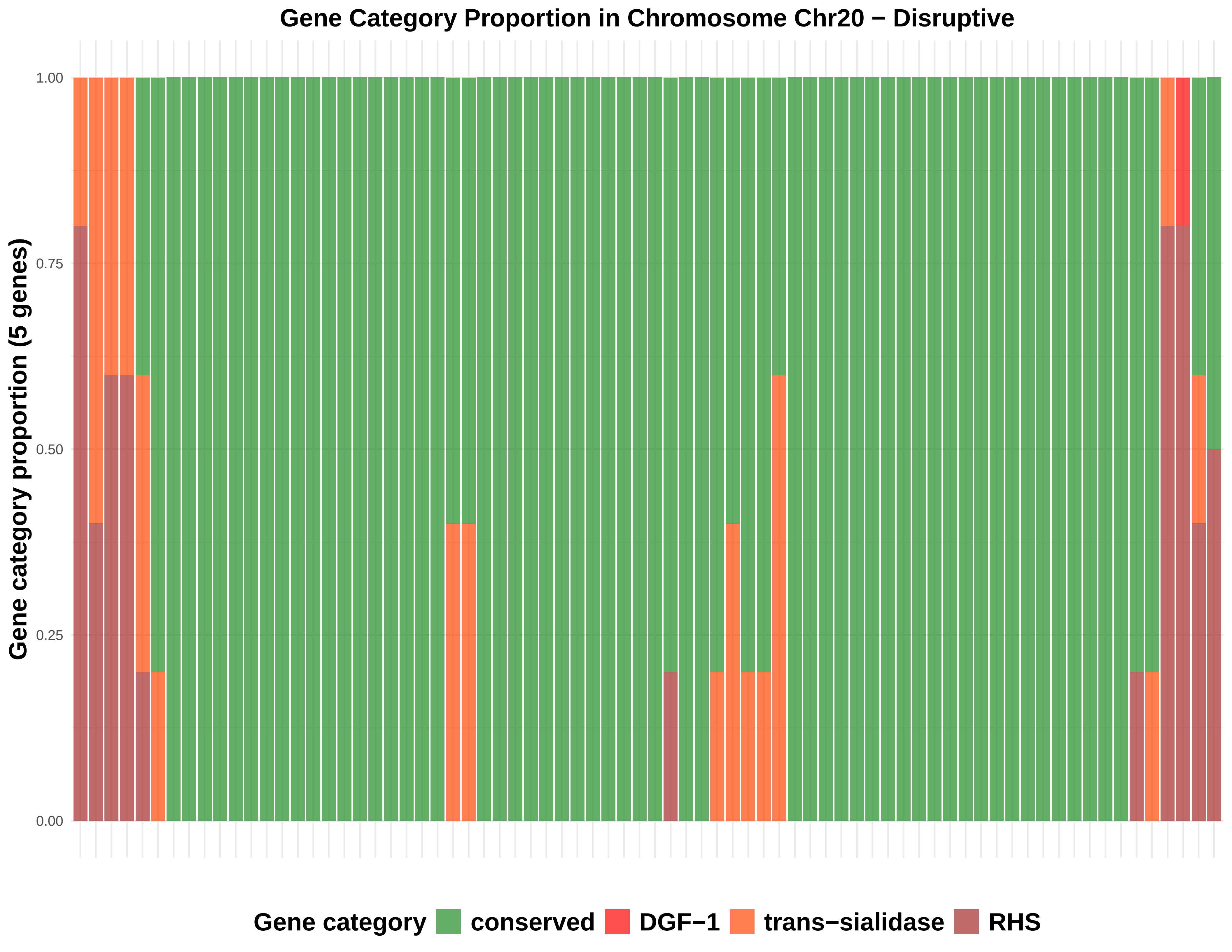
Gene Category Proportion in Chromosome Chr20 − Disruptive
Gene category proportion (5 genes)
1.00
0.75
0.50
0.25
0.00
Gene category
conserved
DGF−1
trans−sialidase
RHS

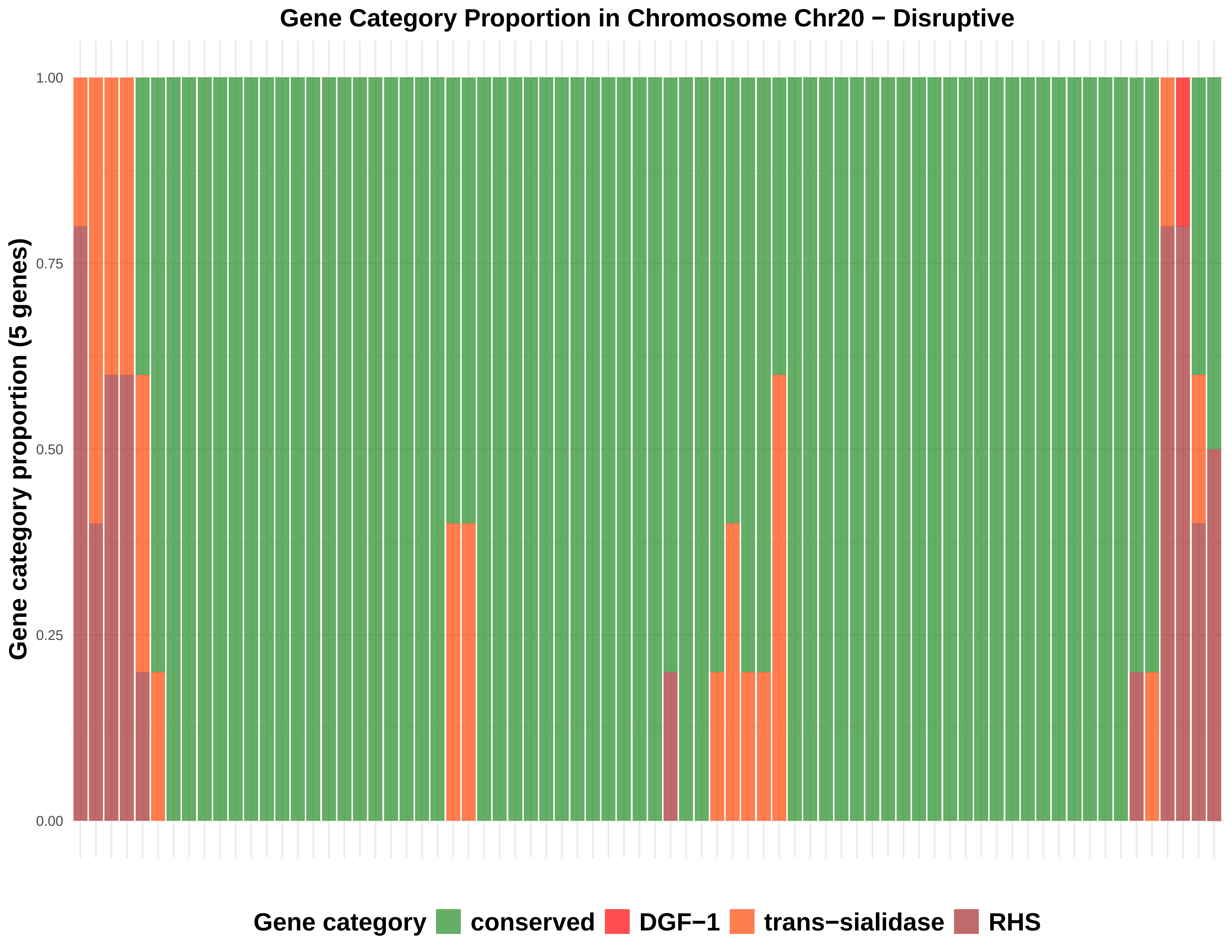

Gene Category Proportion in Chromosome Chr20 − Disruptive
Gene category proportion (5 genes)
1.00
0.75
0.50
0.25
0.00
Gene category
conserved
DGF−1
trans−sialidase
RHS

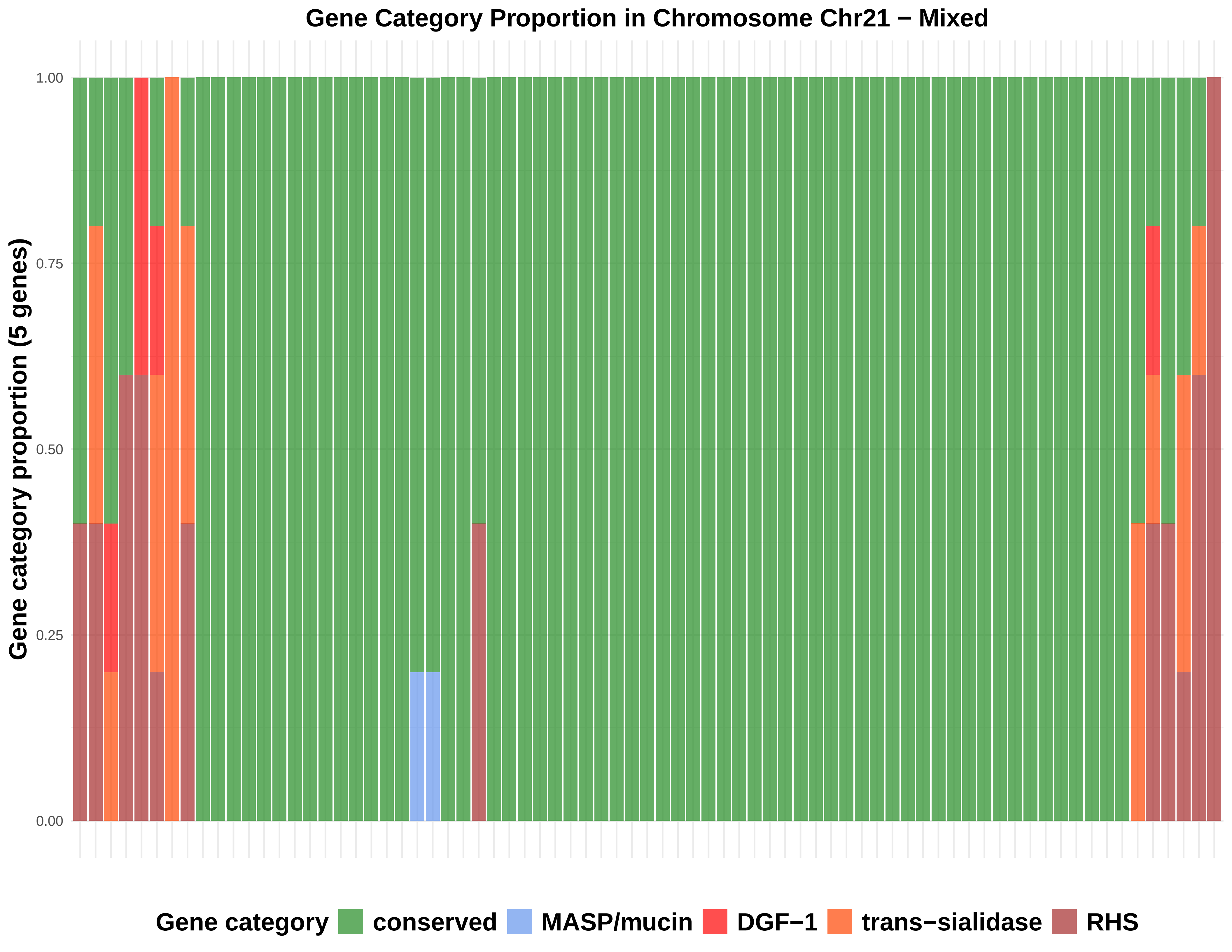
Gene Category Proportion in Chromosome Chr21 − Mixed
Gene category proportion (5 genes)
1.00
0.75
0.50
0.25
0.00
Gene category
conserved
MASP/mucin
DGF−1
trans−sialidase
RHS

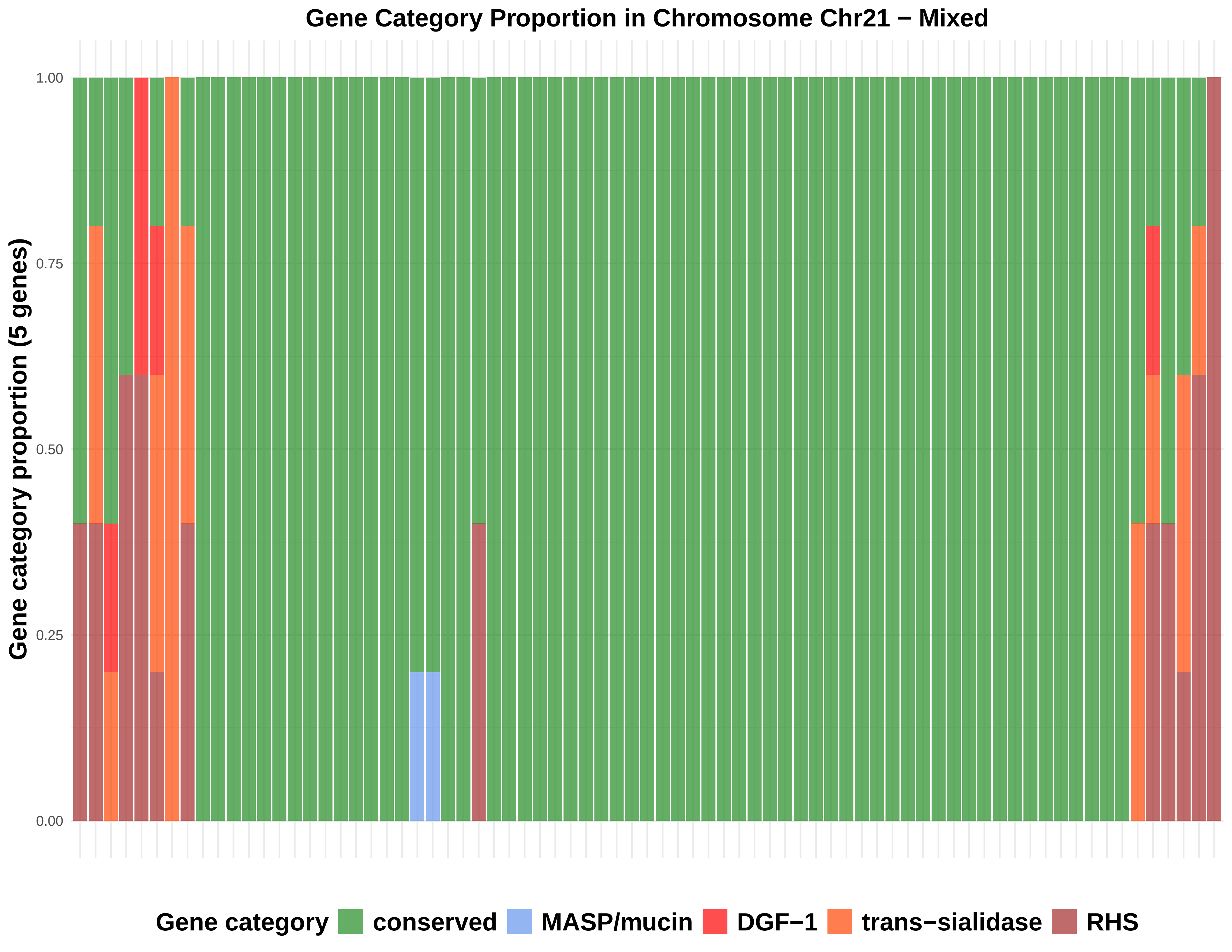
Gene Category Proportion in Chromosome Chr21 − Mixed
Gene category proportion (5 genes)
1.00
0.75
0.50
0.25
0.00
Gene category
conserved
MASP/mucin
DGF−1
trans−sialidase
RHS

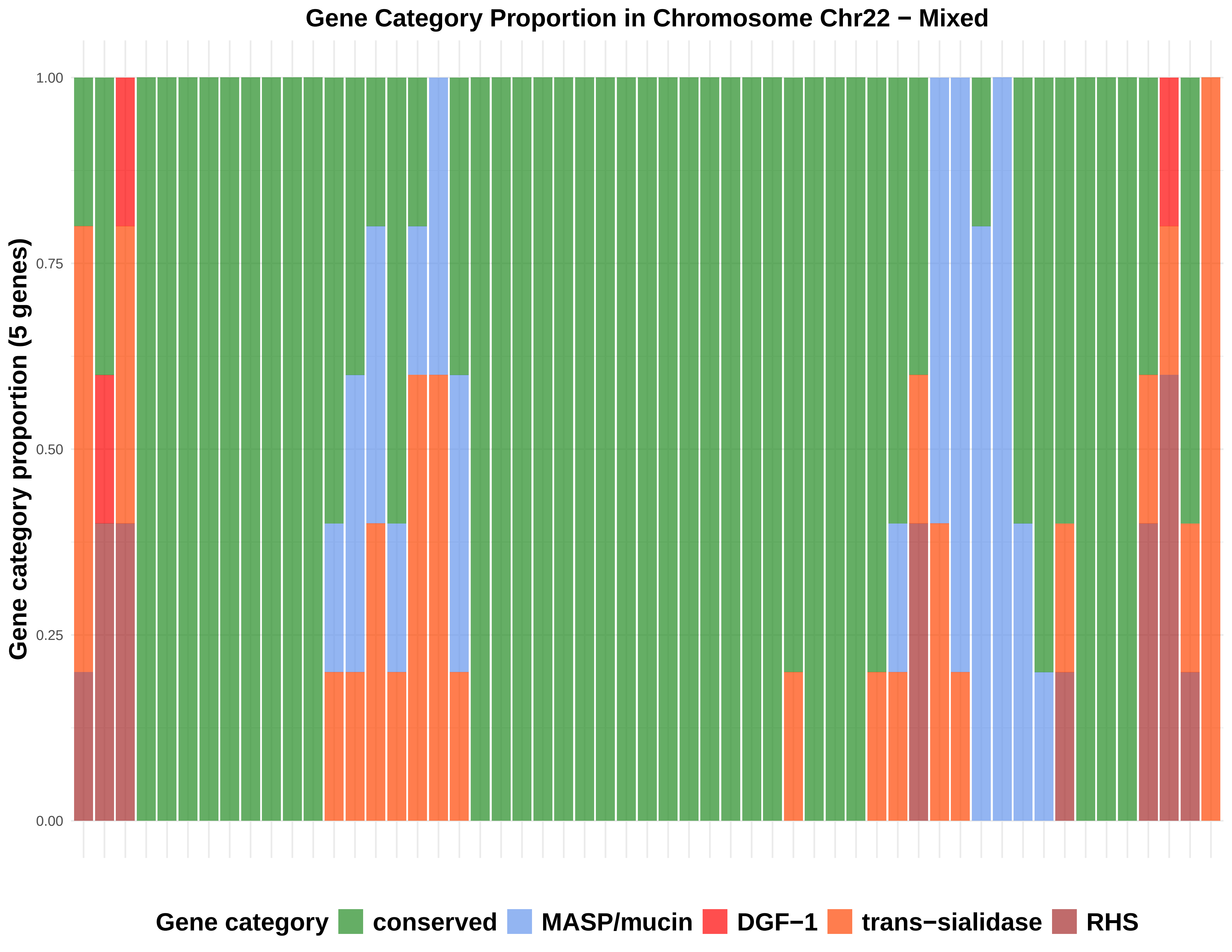
Gene Category Proportion in Chromosome Chr22 − Mixed
Gene category proportion (5 genes)
1.00
0.75
0.50
0.25
0.00
Gene category
conserved
MASP/mucin
DGF−1
trans−sialidase
RHS

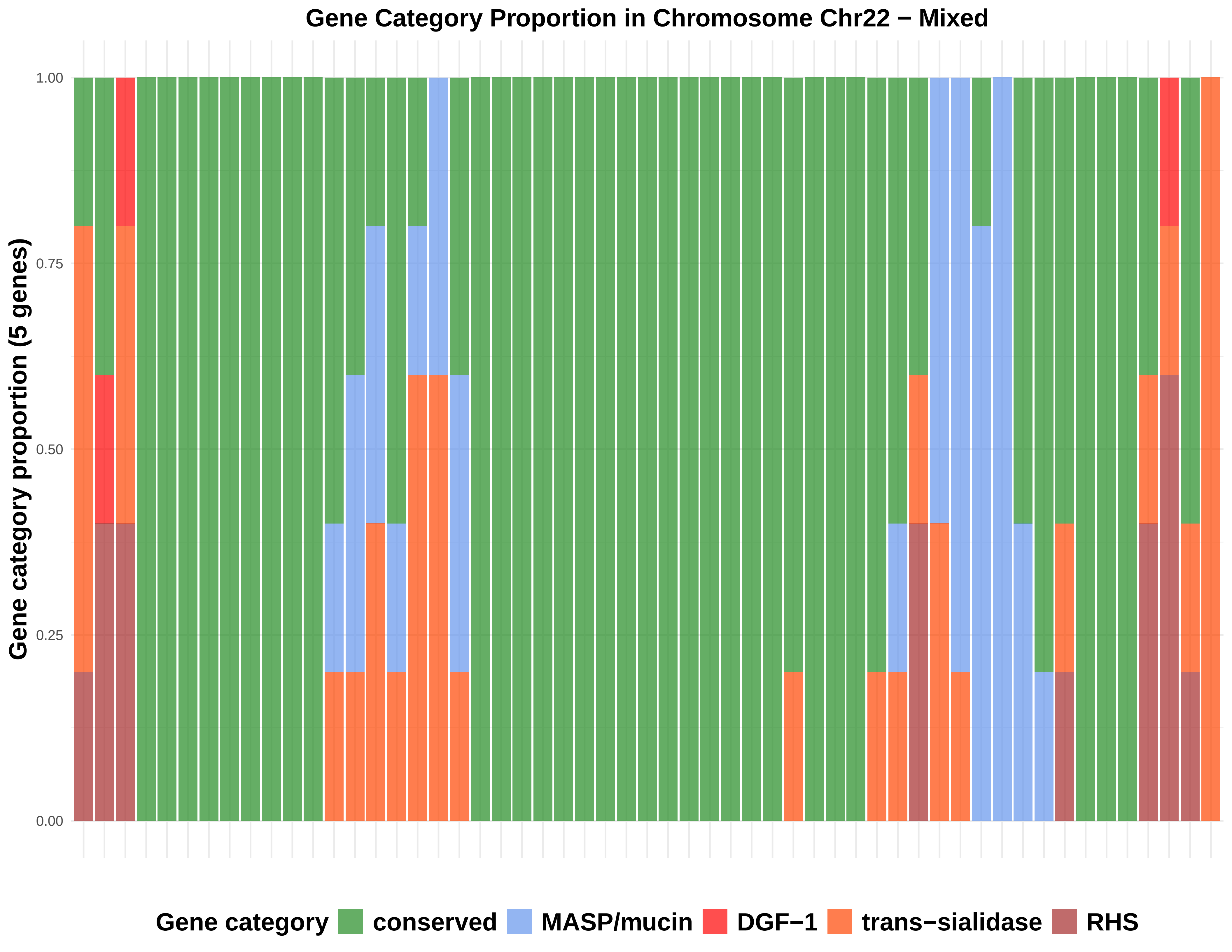
Gene Category Proportion in Chromosome Chr22 − Mixed
Gene category proportion (5 genes)
1.00
0.75
0.50
0.25
0.00
Gene category
conserved
MASP/mucin
DGF−1
trans−sialidase
RHS

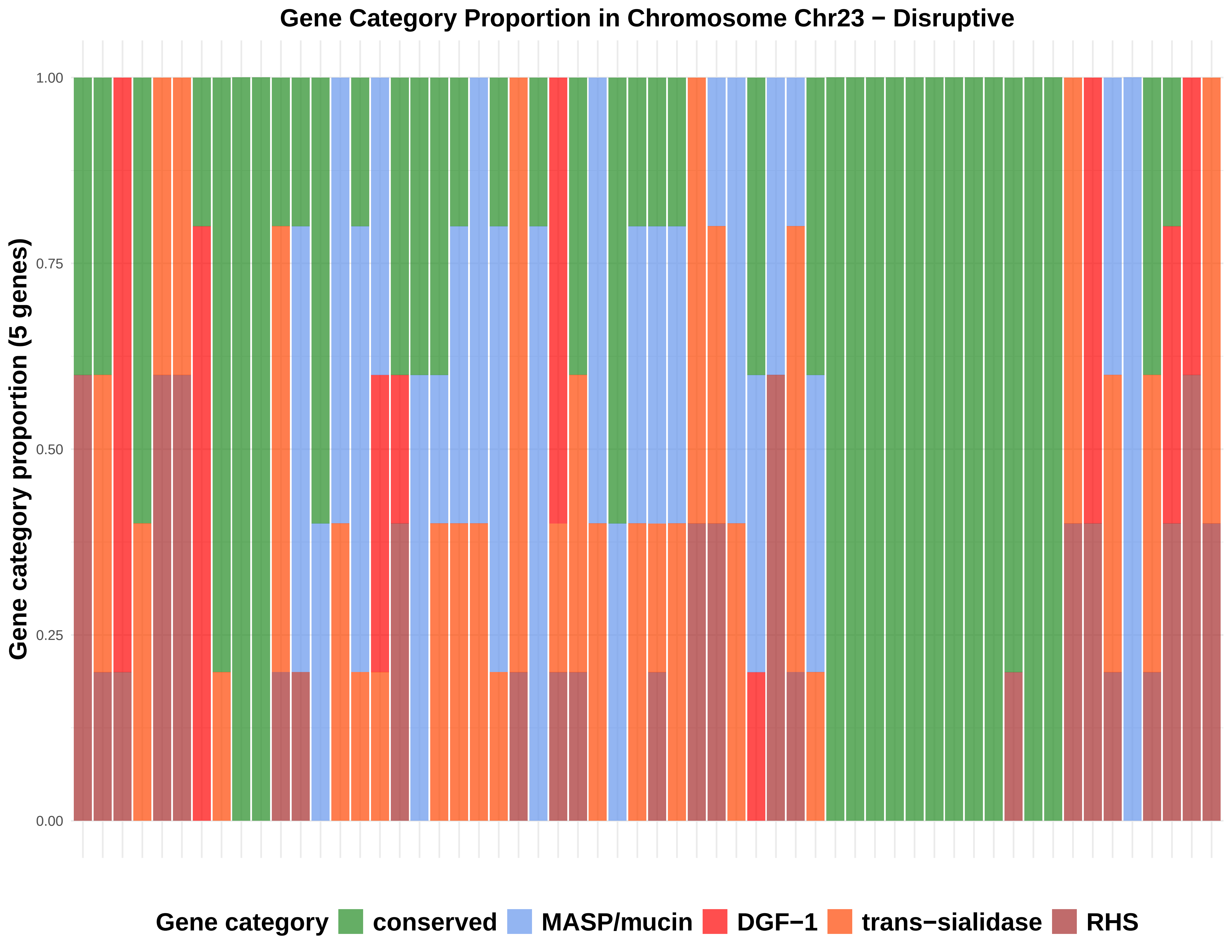

Gene Category Proportion in Chromosome Chr23 − Disruptive
Gene category proportion (5 genes)
1.00
0.75
0.50
0.25
0.00
Gene category
conserved
MASP/mucin
DGF−1
trans−sialidase
RHS

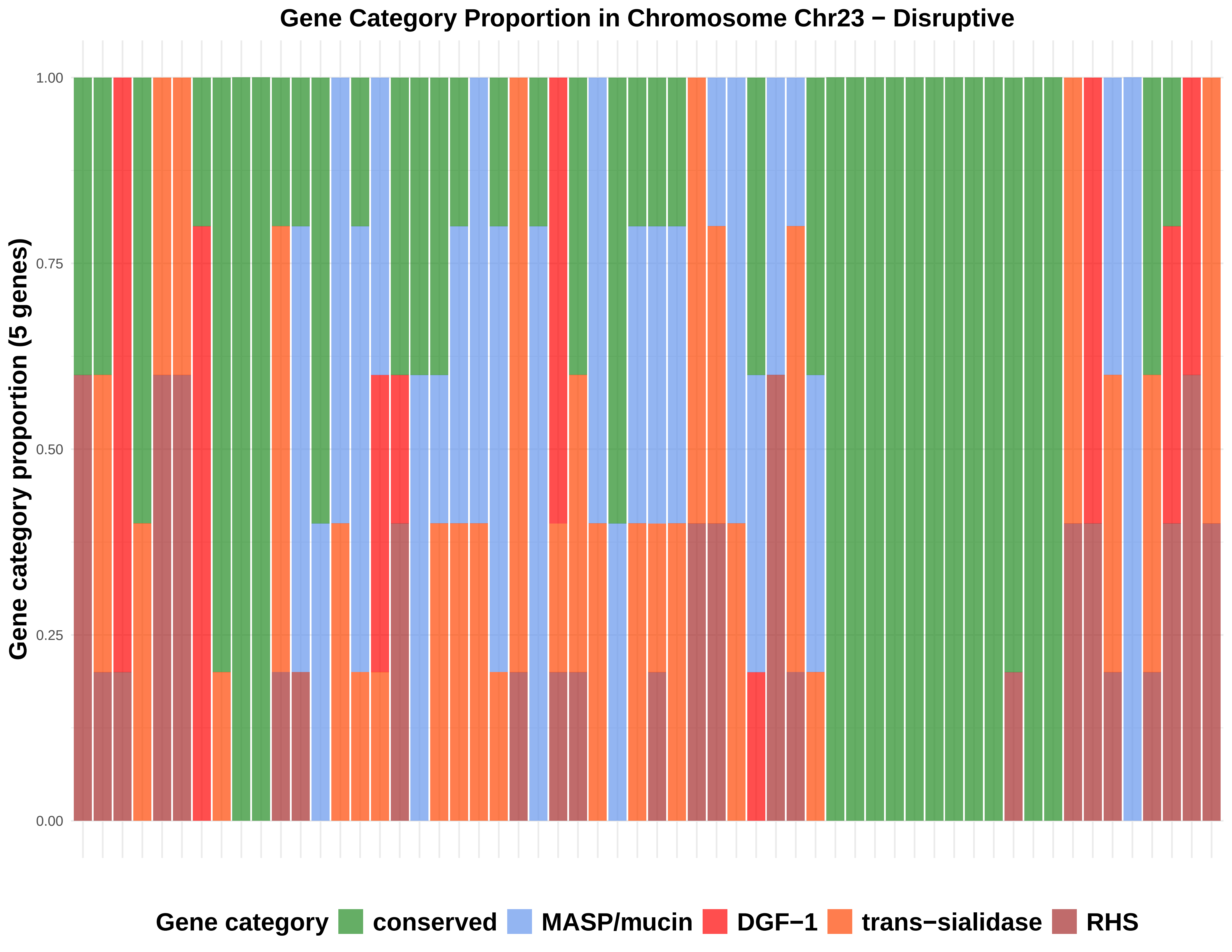

Gene Category Proportion in Chromosome Chr23 − Disruptive
Gene category proportion (5 genes)
1.00
0.75
0.50
0.25
0.00
Gene category
conserved
MASP/mucin
DGF−1
trans−sialidase
RHS

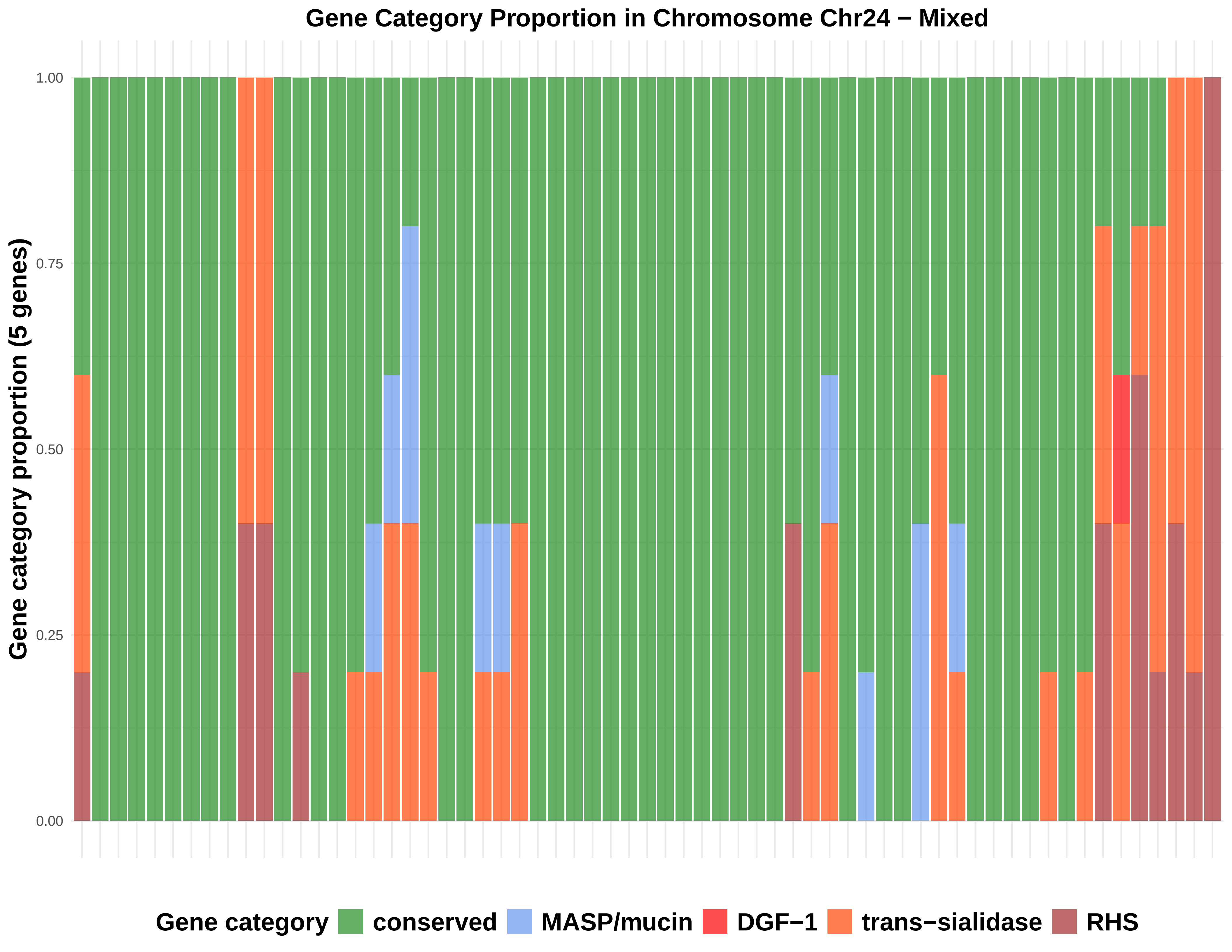
Gene Category Proportion in Chromosome Chr24 − Mixed
Gene category proportion (5 genes)
1.00
0.75
0.50
0.25
0.00
Gene category
conserved
MASP/mucin
DGF−1
trans−sialidase
RHS

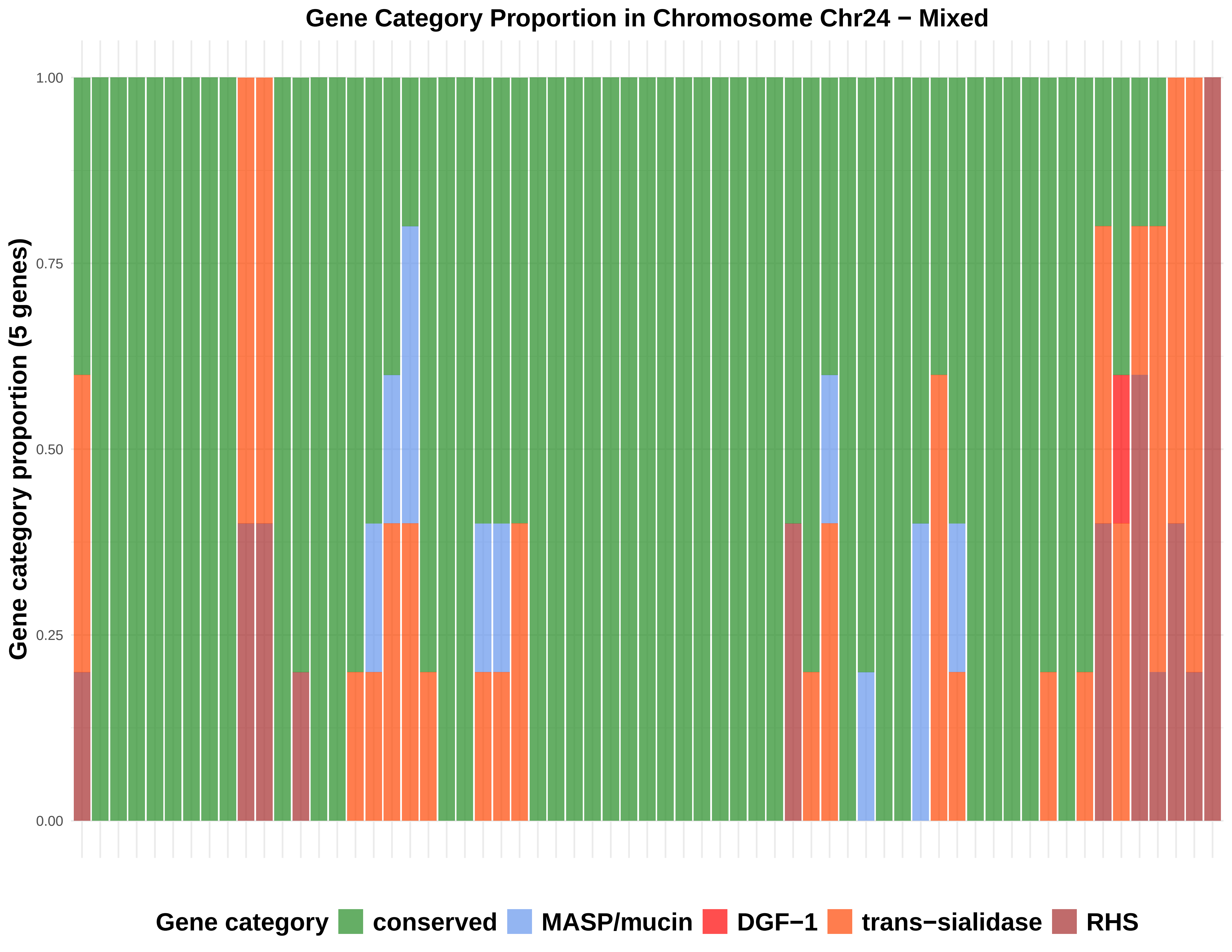

Gene Category Proportion in Chromosome Chr24 − Mixed
Gene category proportion (5 genes)
1.00
0.75
0.50
0.25
0.00
Gene category
conserved
MASP/mucin
DGF−1
trans−sialidase
RHS

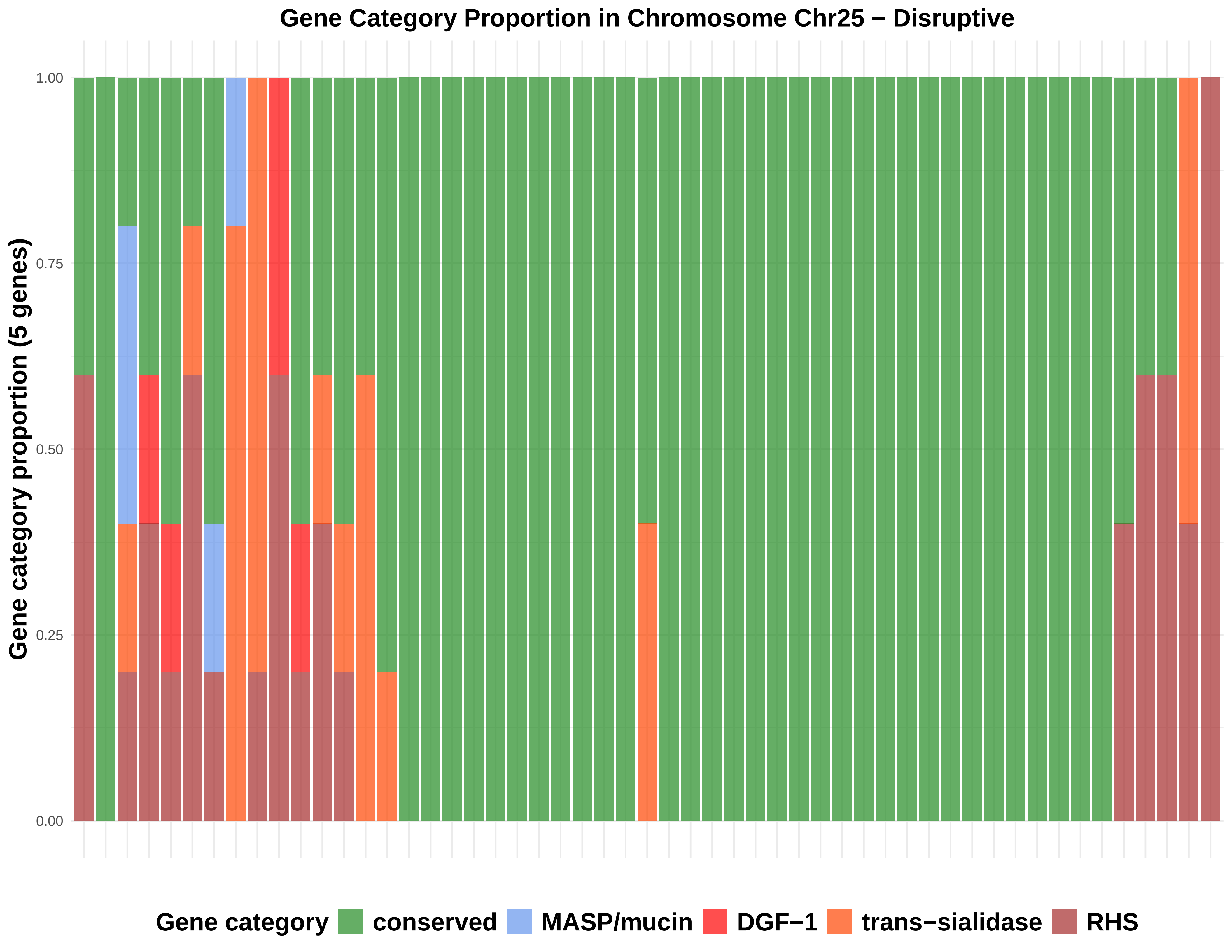

Gene Category Proportion in Chromosome Chr25 − Disruptive
Gene category proportion (5 genes)
1.00
0.75
0.50
0.25
0.00
Gene category
conserved
MASP/mucin
DGF−1
trans−sialidase
RHS

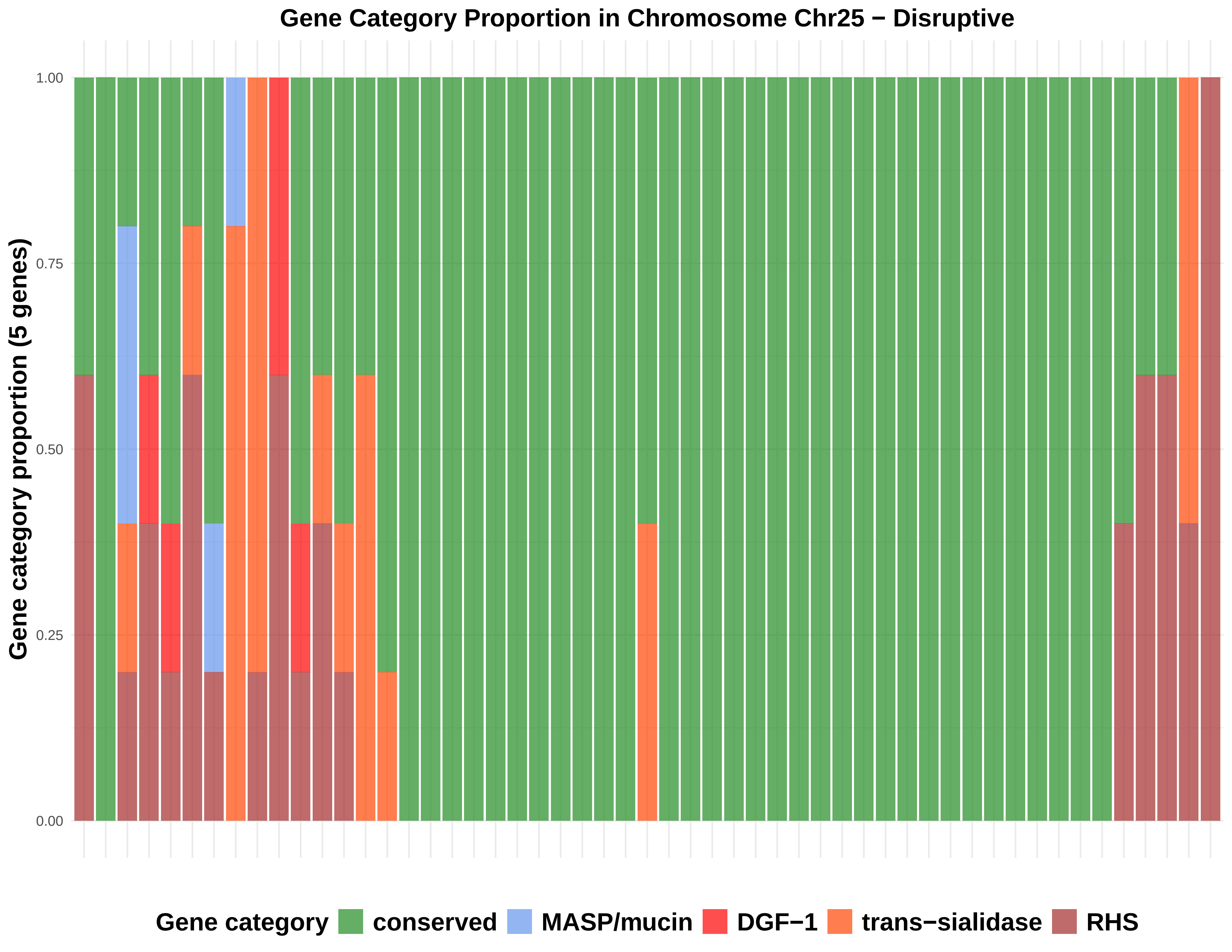

Gene Category Proportion in Chromosome Chr25 − Disruptive
Gene category proportion (5 genes)
1.00
0.75
0.50
0.25
0.00
Gene category
conserved
MASP/mucin
DGF−1
trans−sialidase
RHS

### Gene Category Proportion in Chromosome Chr26 − Disruptive

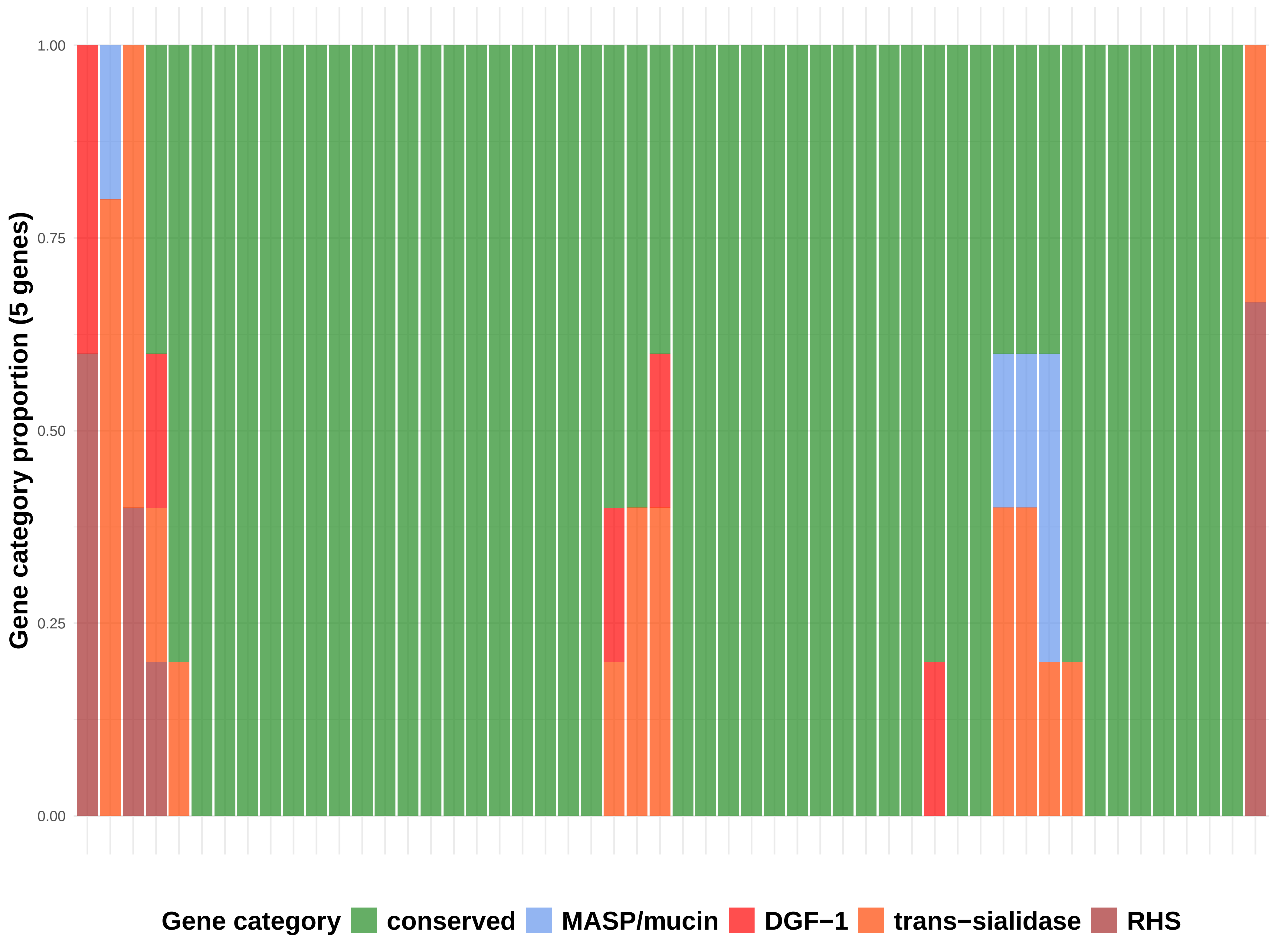

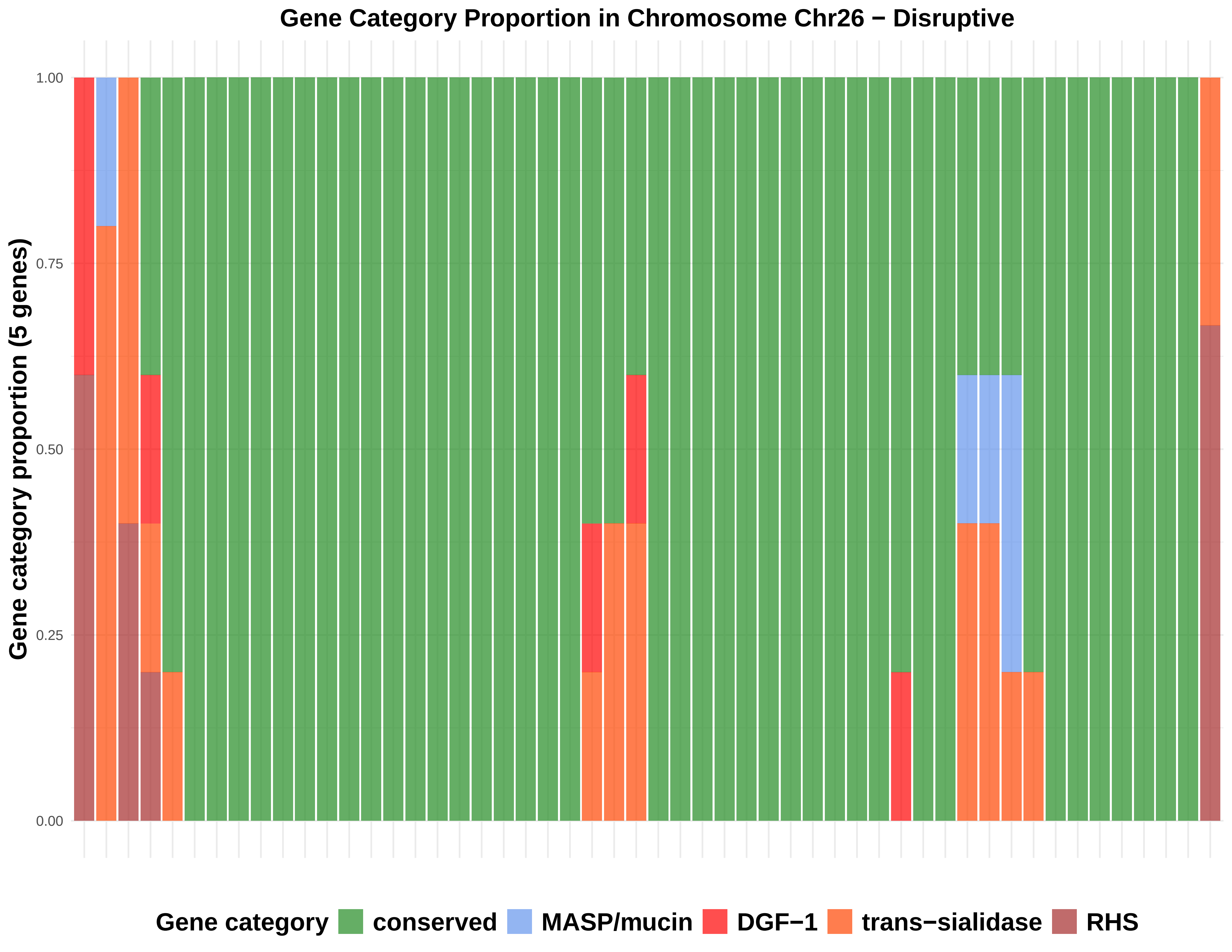

Gene Category Proportion in Chromosome Chr26 − Disruptive
Gene category proportion (5 genes)
1.00
0.75
0.50
0.25
0.00
Gene category
conserved
MASP/mucin
DGF−1
trans−sialidase
RHS

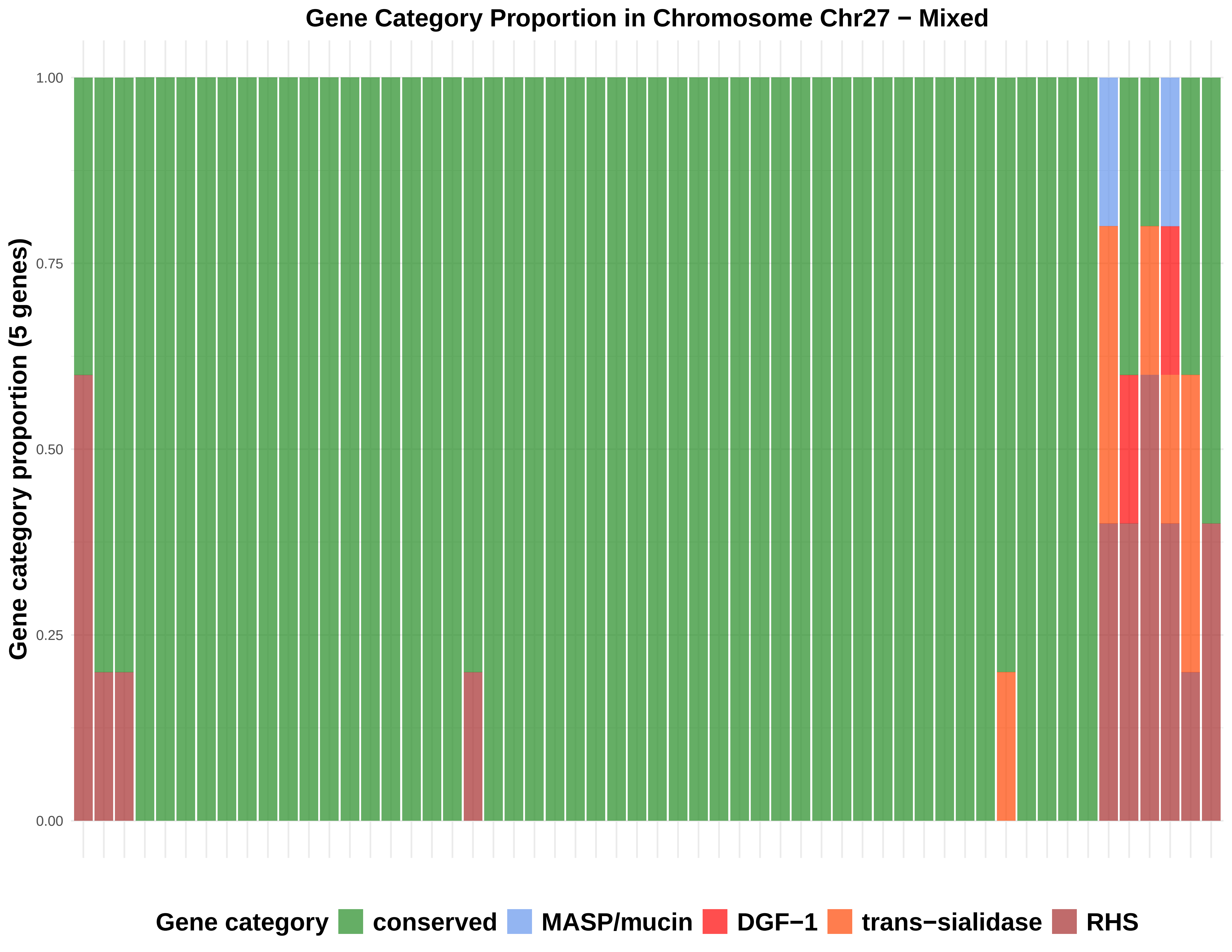

Gene Category Proportion in Chromosome Chr27 − Mixed
Gene category proportion (5 genes)
1.00
0.75
0.50
0.25
0.00
Gene category
conserved
MASP/mucin
DGF−1
trans−sialidase
RHS

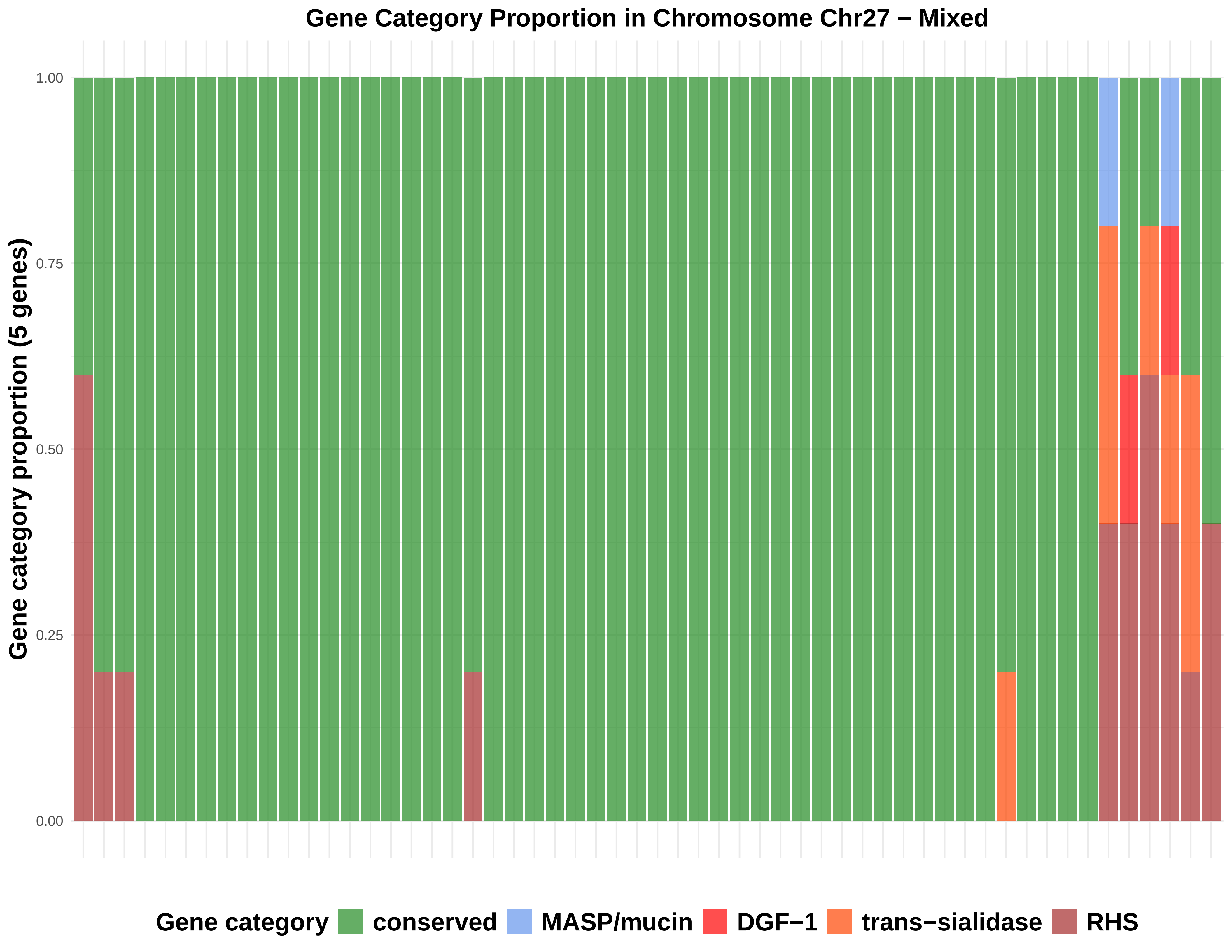

Gene Category Proportion in Chromosome Chr27 − Mixed
Gene category proportion (5 genes)
1.00
0.75
0.50
0.25
0.00
Gene category
conserved
MASP/mucin
DGF−1
trans−sialidase
RHS

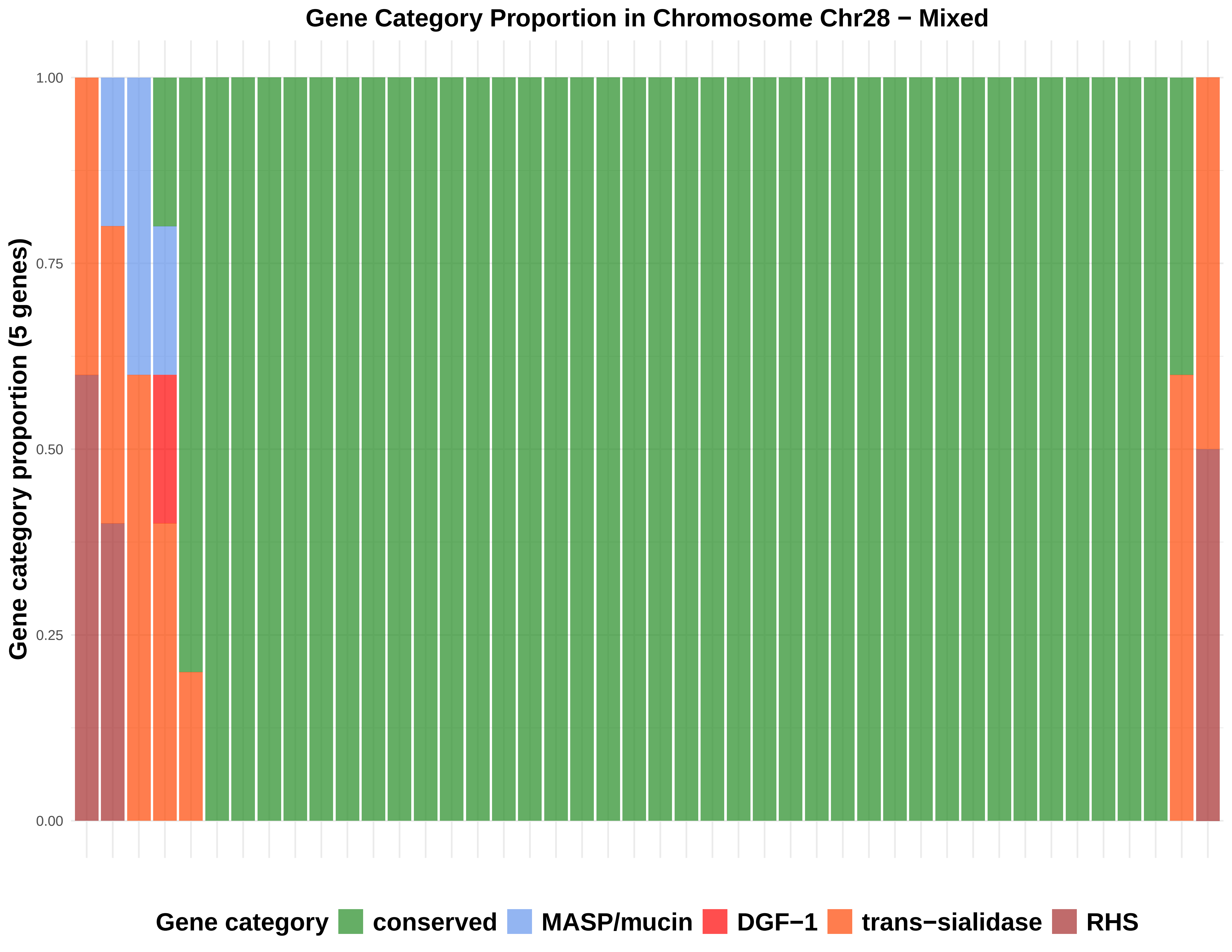
Gene Category Proportion in Chromosome Chr28 − Mixed
Gene category proportion (5 genes)
1.00
0.75
0.50
0.25
0.00
Gene category
conserved
MASP/mucin
DGF−1
trans−sialidase
RHS

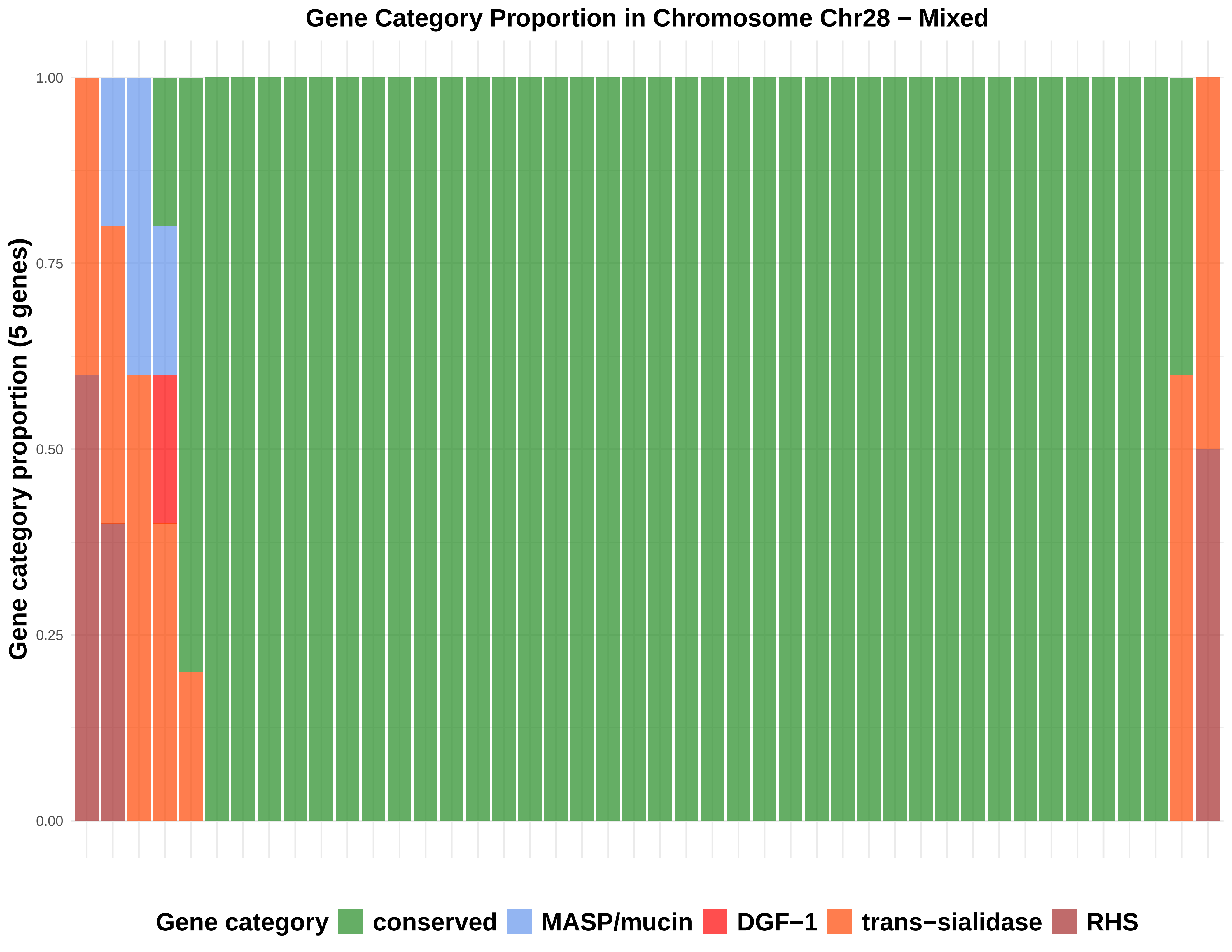

Gene Category Proportion in Chromosome Chr28 − Mixed
Gene category proportion (5 genes)
1.00
0.75
0.50
0.25
0.00
Gene category
conserved
MASP/mucin
DGF−1
trans−sialidase
RHS

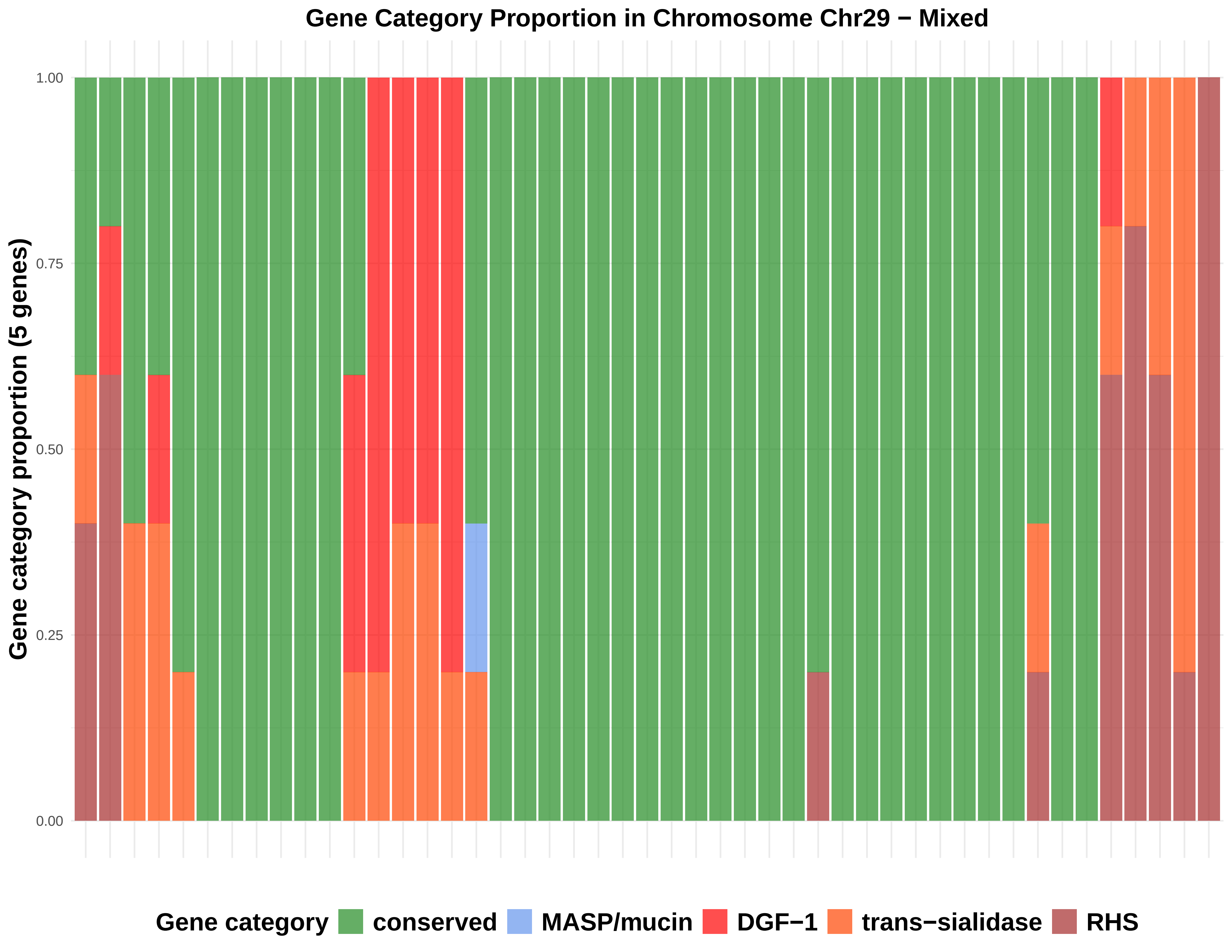

Gene Category Proportion in Chromosome Chr29 − Mixed
Gene category proportion (5 genes)
1.00
0.75
0.50
0.25
0.00
Gene category
conserved
MASP/mucin
DGF−1
trans−sialidase
RHS

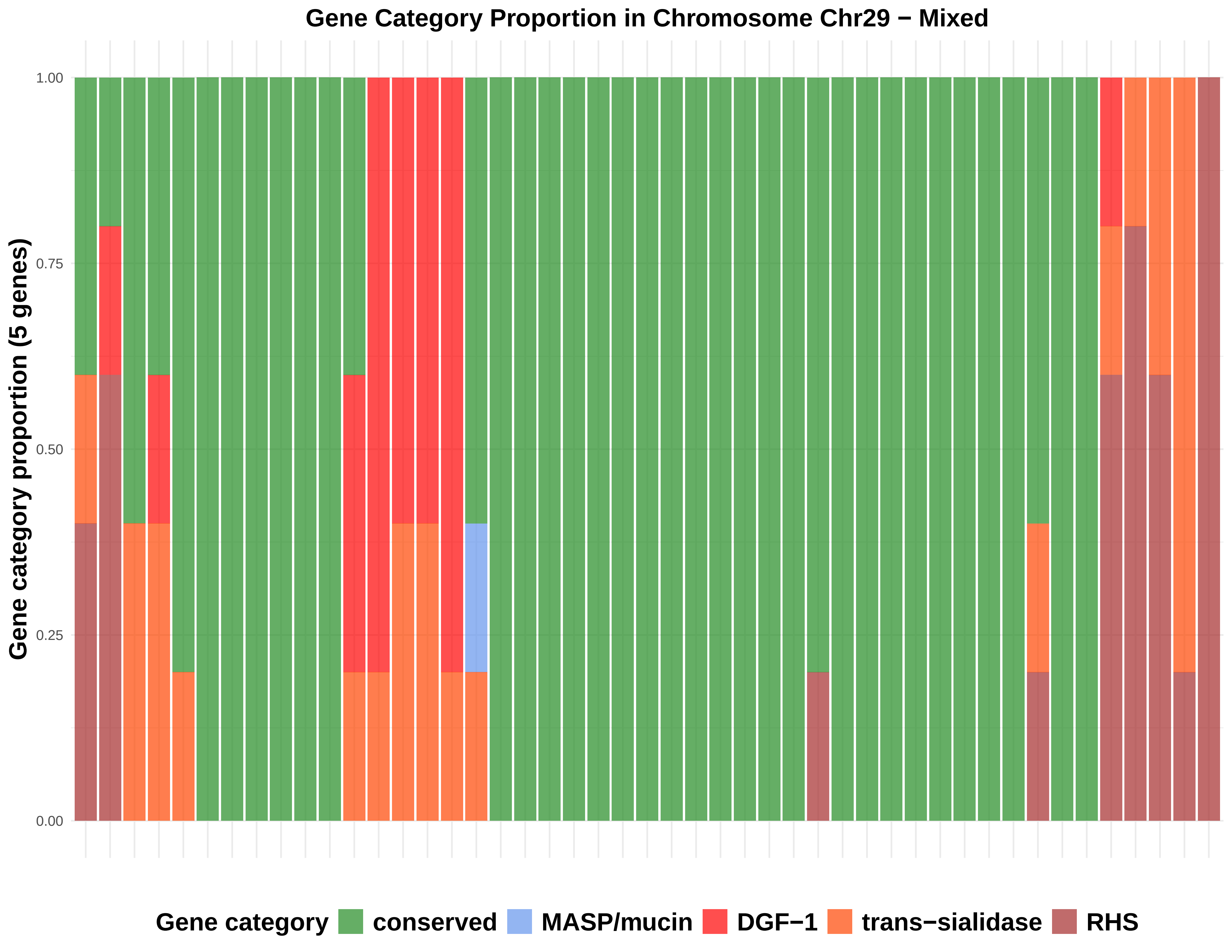

Gene Category Proportion in Chromosome Chr29 − Mixed
Gene category proportion (5 genes)
1.00
0.75
0.50
0.25
0.00
Gene category
conserved
MASP/mucin
DGF−1
trans−sialidase
RHS

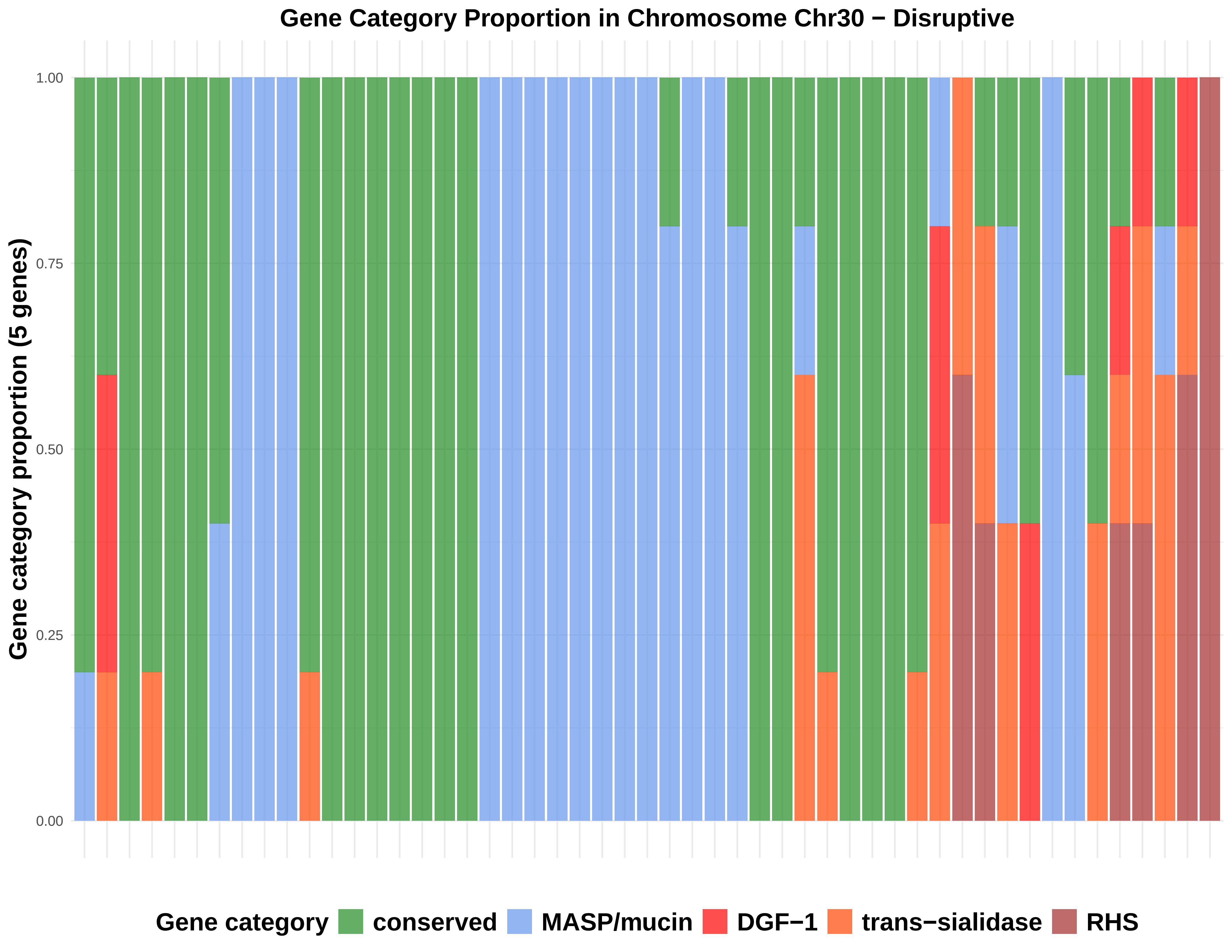

Gene Category Proportion in Chromosome Chr30 − Disruptive
Gene category proportion (5 genes)
1.00
0.75
0.50
0.25
0.00
Gene category
conserved
MASP/mucin
DGF−1
trans−sialidase
RHS

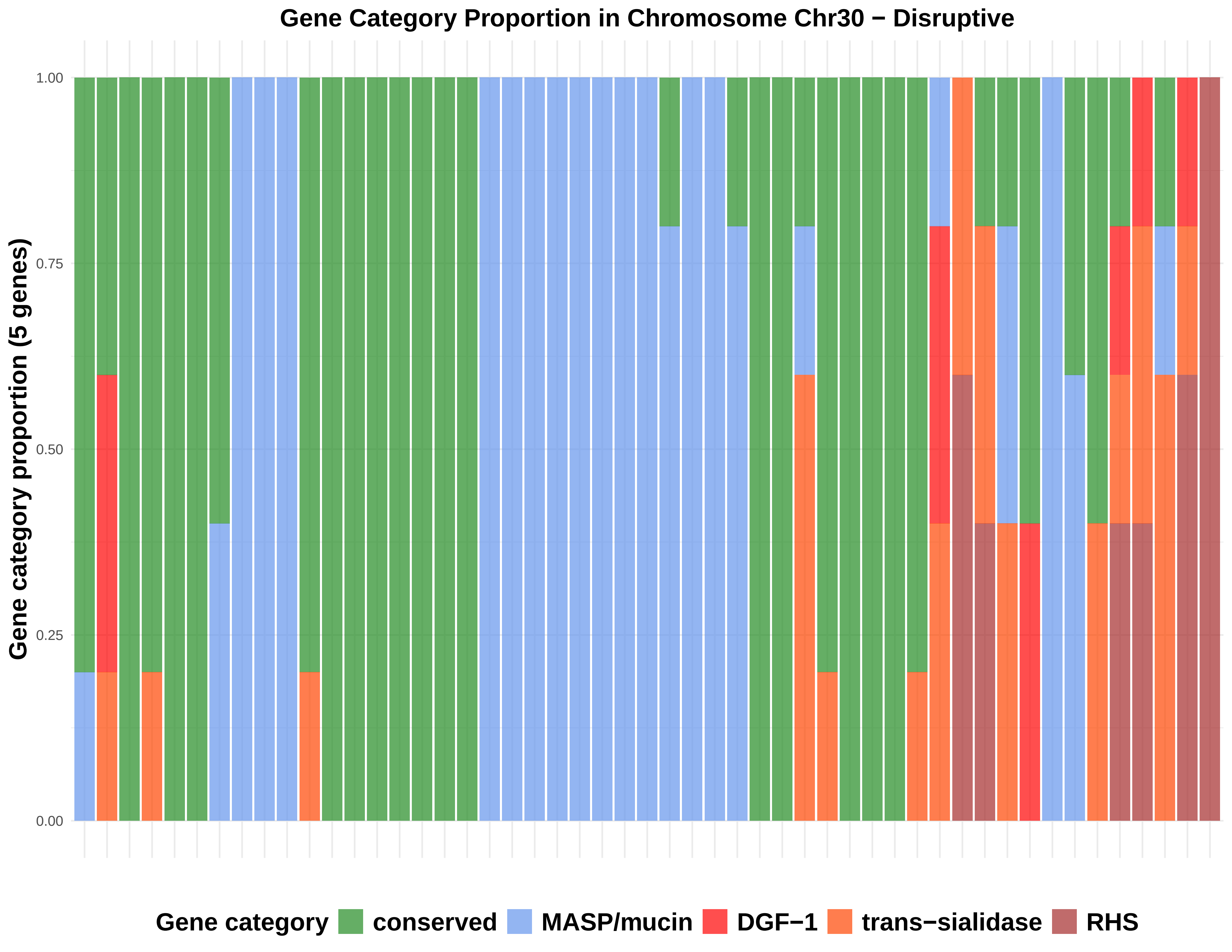
Gene Category Proportion in Chromosome Chr30 − Disruptive
Gene category proportion (5 genes)
1.00
0.75
0.50
0.25
0.00
Gene category
conserved
MASP/mucin
DGF−1
trans−sialidase
RHS

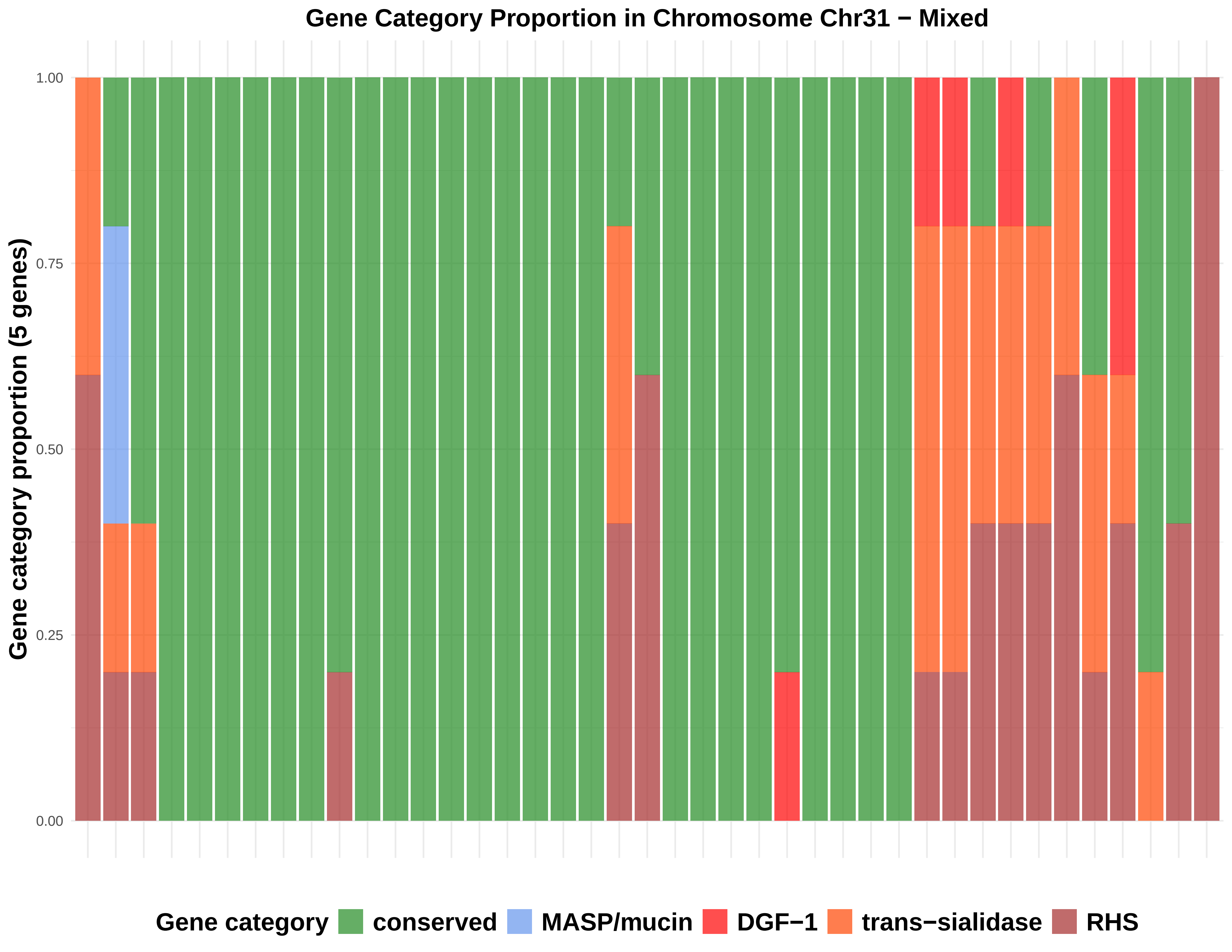

Gene Category Proportion in Chromosome Chr31 − Mixed
Gene category proportion (5 genes)
1.00
0.75
0.50
0.25
0.00
Gene category
conserved
MASP/mucin
DGF−1
trans−sialidase
RHS

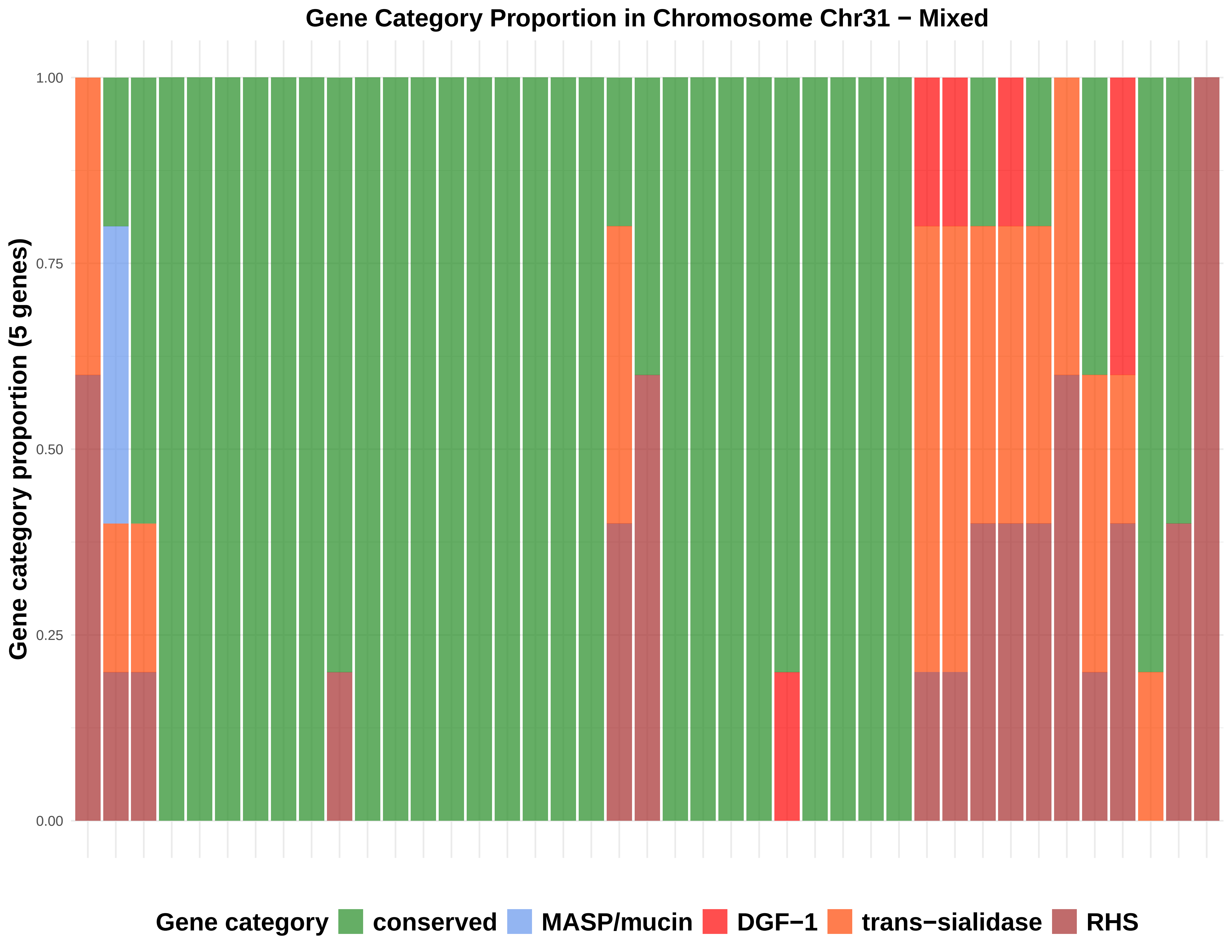

Gene Category Proportion in Chromosome Chr31 − Mixed
Gene category proportion (5 genes)
1.00
0.75
0.50
0.25
0.00
Gene category
conserved
MASP/mucin
DGF−1
trans−sialidase
RHS

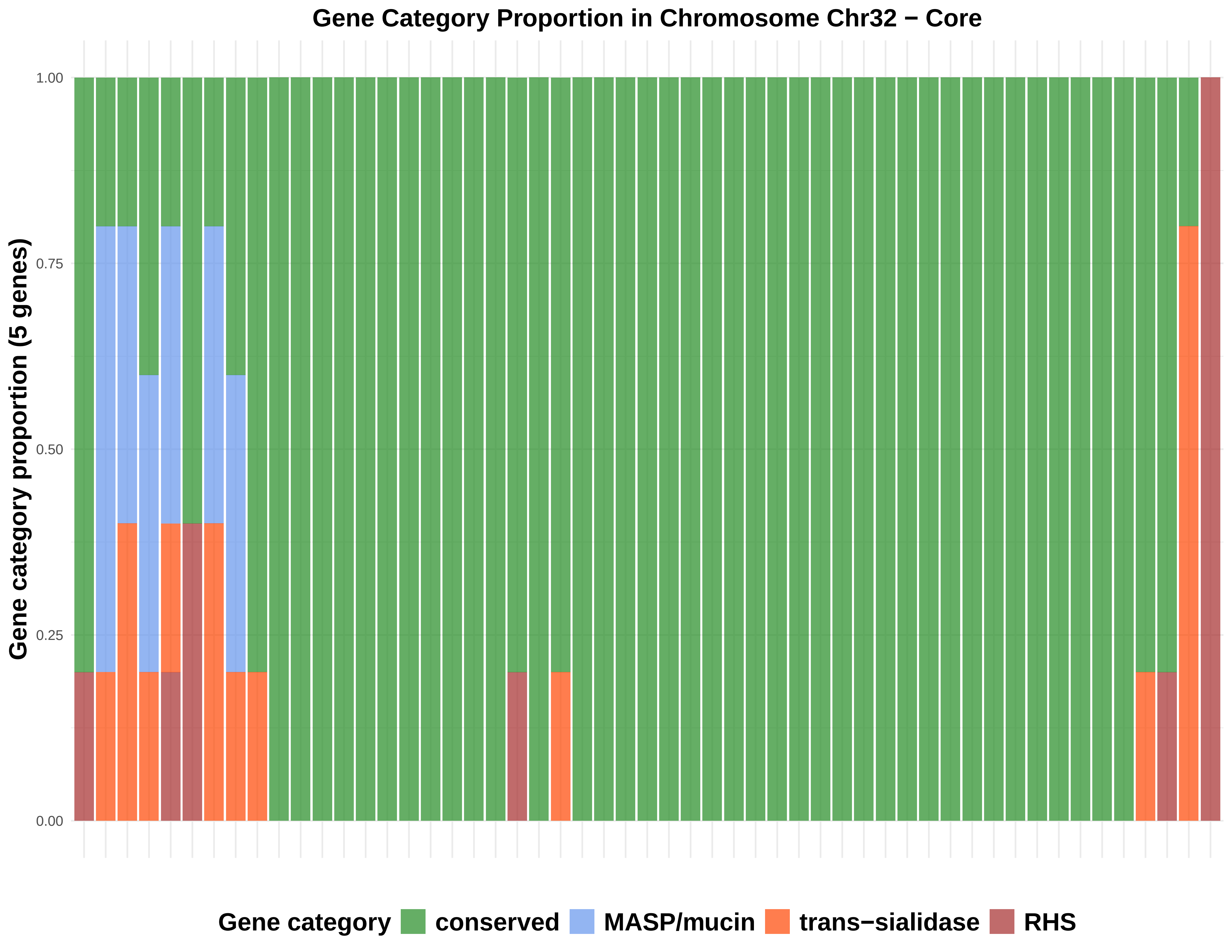
Gene Category Proportion in Chromosome Chr32 − Core
Gene category proportion (5 genes)
1.00
0.75
0.50
0.25
0.00
Gene category
conserved
MASP/mucin
trans−sialidase
RHS

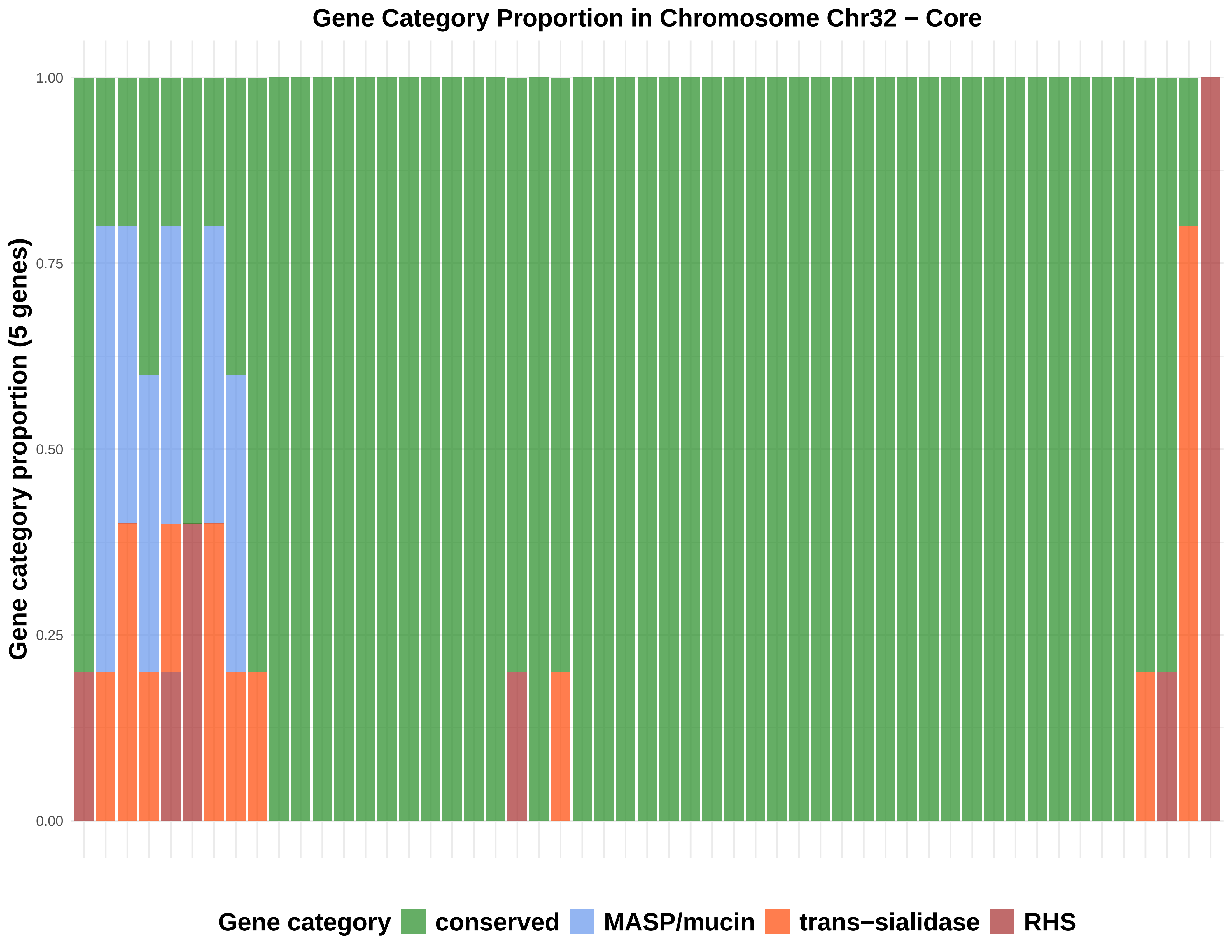
Gene Category Proportion in Chromosome Chr32 − Core
Gene category proportion (5 genes)
1.00
0.75
0.50
0.25
0.00
Gene category
conserved
MASP/mucin
trans−sialidase
RHS
