## Supplementary material for "*Trypanosoma cruzi* has 32 Chromosomes: A Telomere-to-Telomere Assembly Defines Its Karyotype": SupFigure7_BoxplotPseudogeneAndByChromosomeType.pptx

### Slide 1
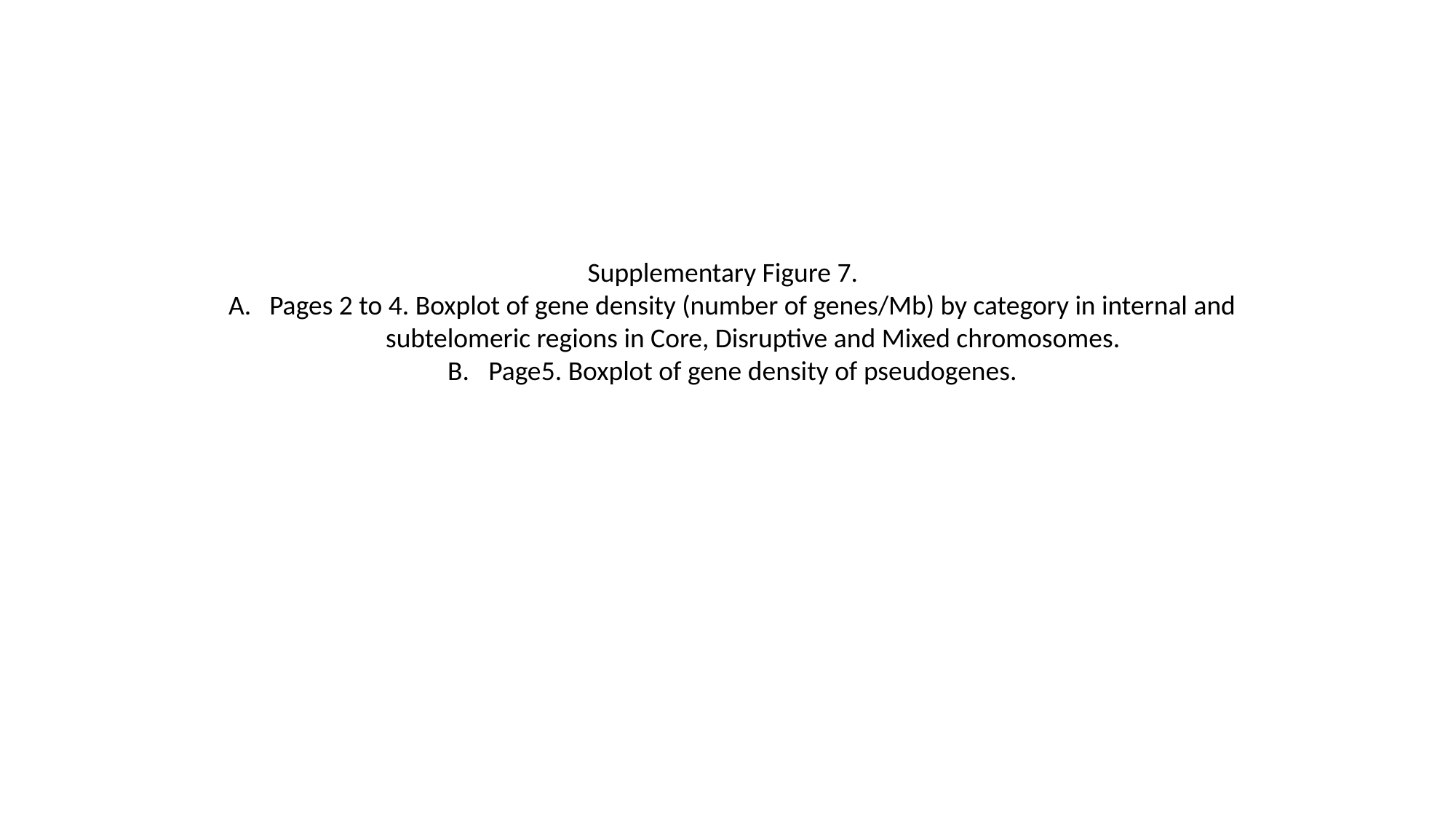

Supplementary Figure 7.
Pages 2 to 4. Boxplot of gene density (number of genes/Mb) by category in internal and subtelomeric regions in Core, Disruptive and Mixed chromosomes.
Page5. Boxplot of gene density of pseudogenes.

### Slide 2
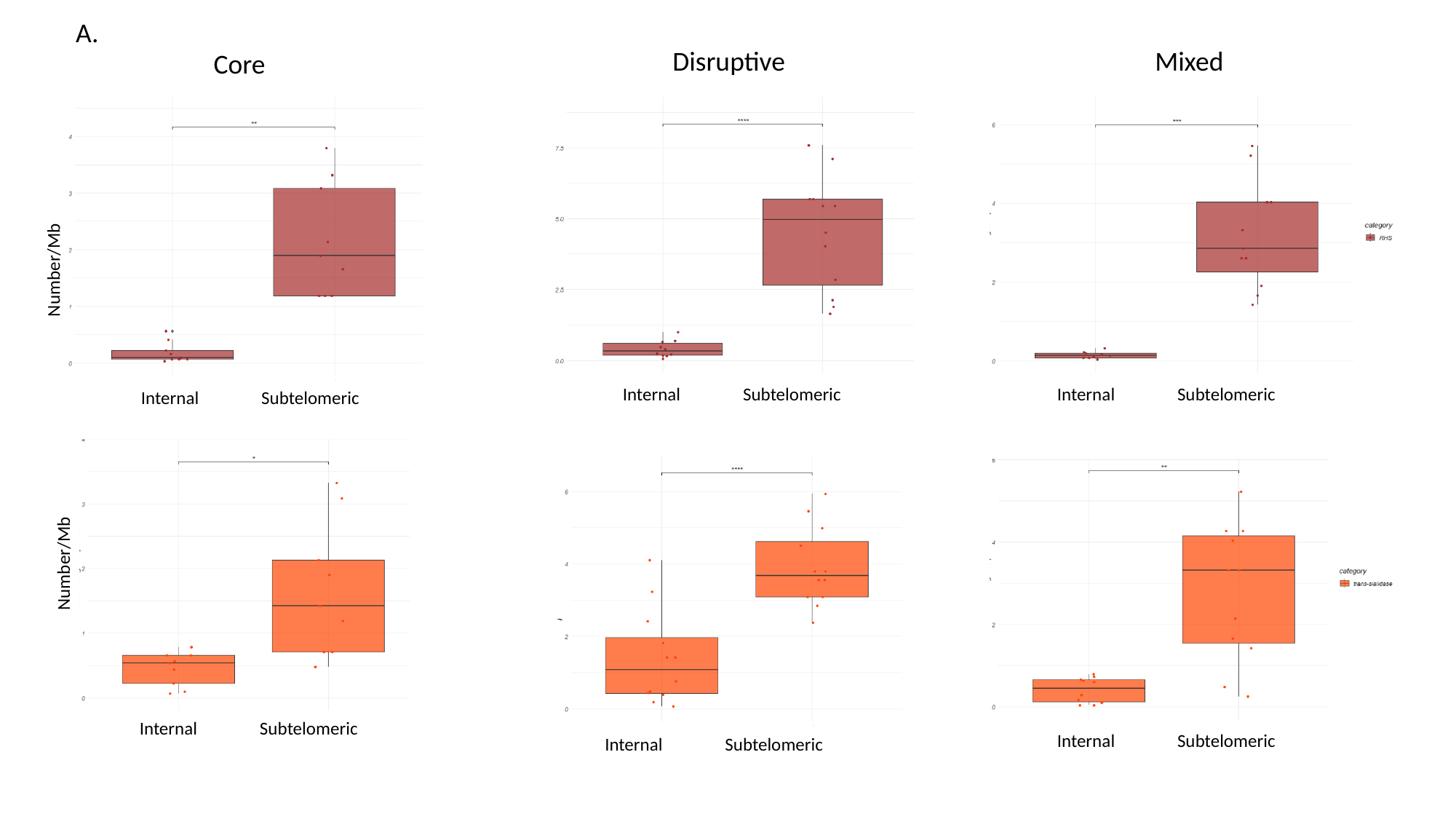

A.
Disruptive
Mixed
Core
Number/Mb
Internal Subtelomeric
Internal Subtelomeric
Internal Subtelomeric
Number/Mb
Internal Subtelomeric
Internal Subtelomeric
Internal Subtelomeric

### Slide 3
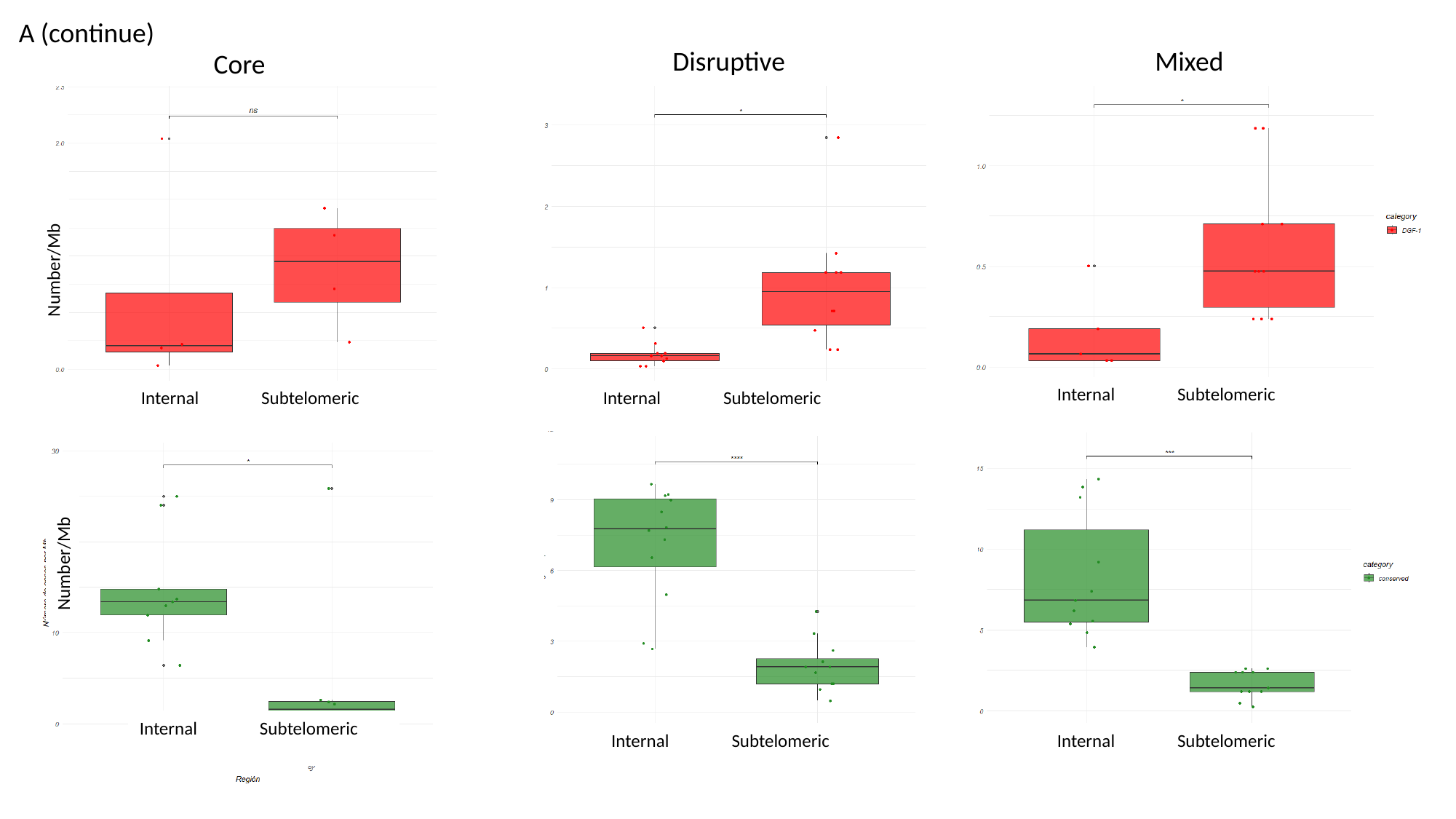

A (continue)
Disruptive
Mixed
Core
Number/Mb
Internal Subtelomeric
Internal Subtelomeric
Internal Subtelomeric
Number/Mb
Internal Subtelomeric
Internal Subtelomeric
Internal Subtelomeric

### Slide 4
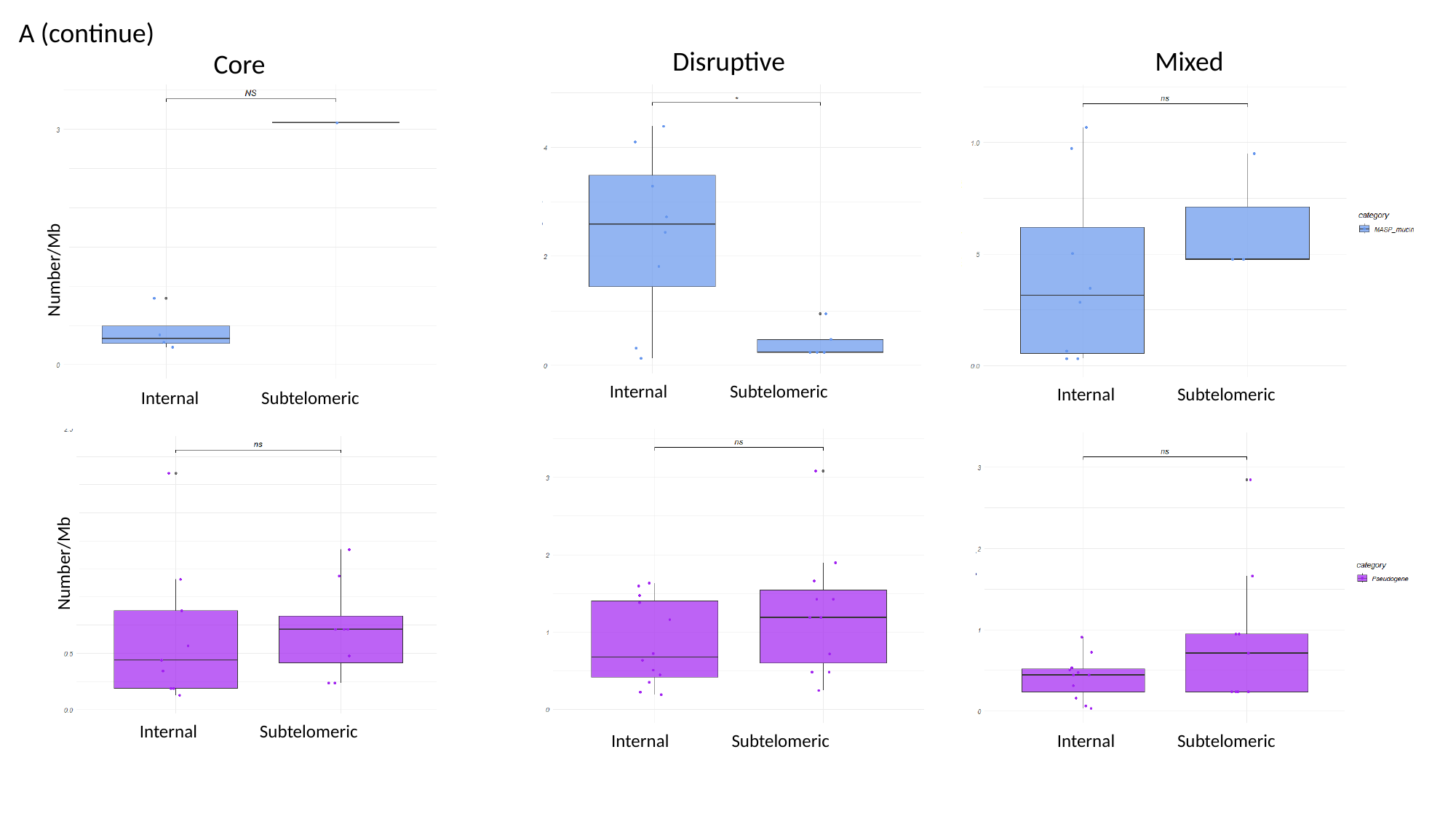

A (continue)
Disruptive
Mixed
Core
Number/Mb
Internal Subtelomeric
Internal Subtelomeric
Internal Subtelomeric
Number/Mb
Internal Subtelomeric
Internal Subtelomeric
Internal Subtelomeric

### Slide 5
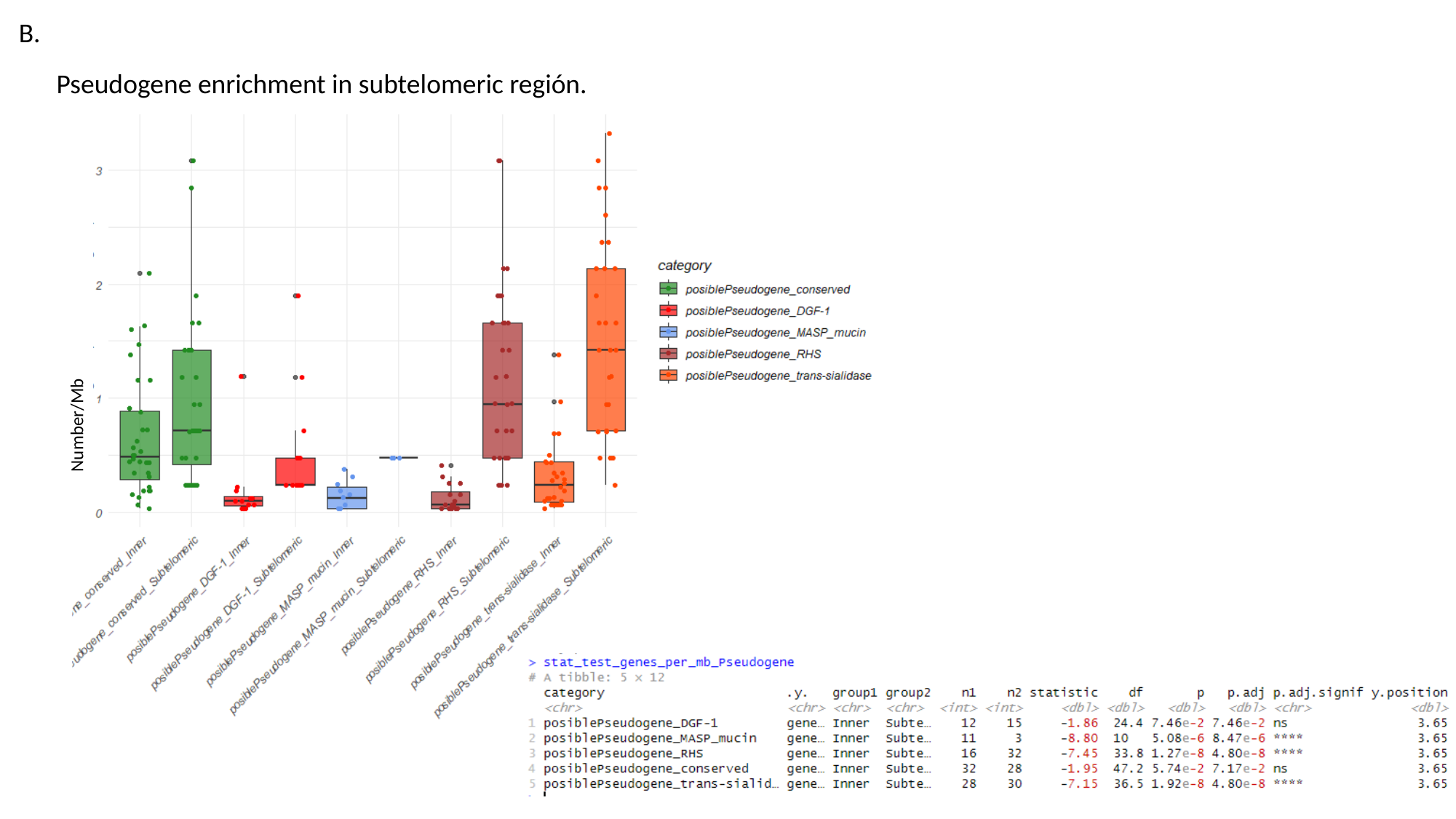

B.
Pseudogene enrichment in subtelomeric región.
Number/Mb
