## Supplementary material for "*Trypanosoma cruzi* has 32 Chromosomes: A Telomere-to-Telomere Assembly Defines Its Karyotype": SupFigure8_ETBoxplots_L1TcSire.pptx

### Slide 1
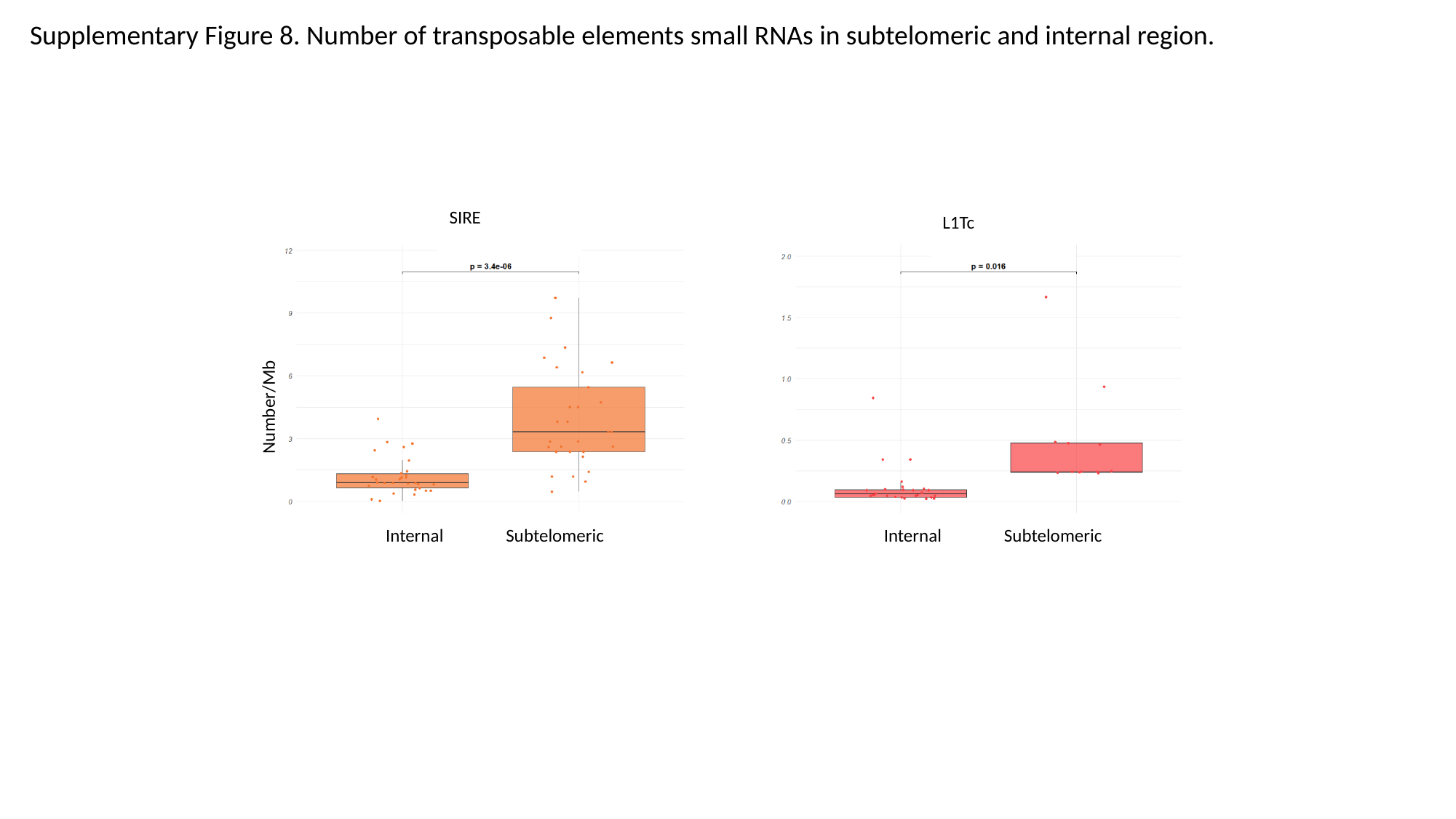

Supplementary Figure 8. Number of transposable elements small RNAs in subtelomeric and internal region.
SIRE
L1Tc
Number/Mb
Internal Subtelomeric
Internal Subtelomeric
