## Supplementary material for "*Trypanosoma cruzi* has 32 Chromosomes: A Telomere-to-Telomere Assembly Defines Its Karyotype": SupFigure9_RHSandDGF1_Trees.pptx

### Slide 1
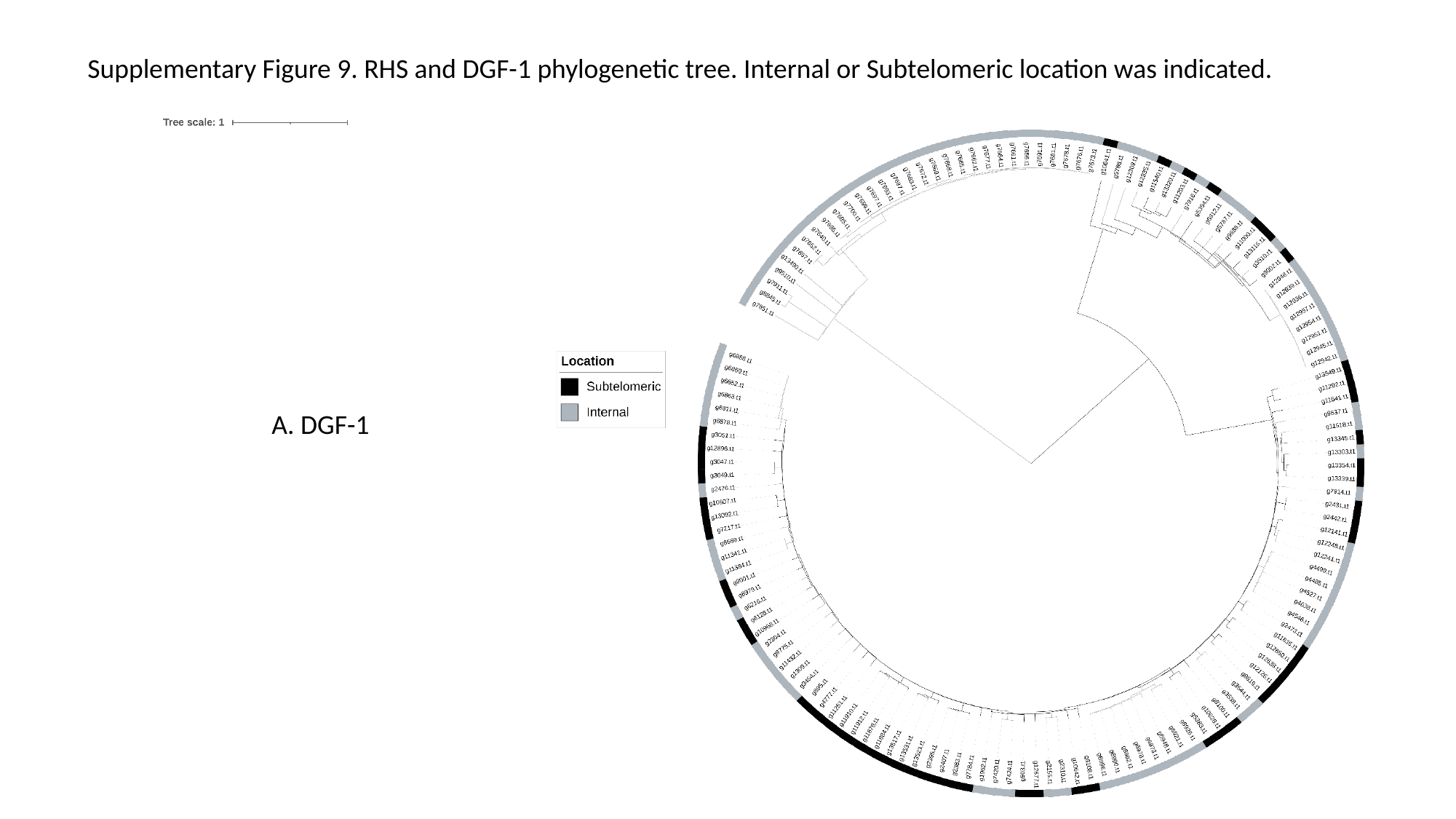

Supplementary Figure 9. RHS and DGF-1 phylogenetic tree. Internal or Subtelomeric location was indicated.
A. DGF-1

### Slide 2
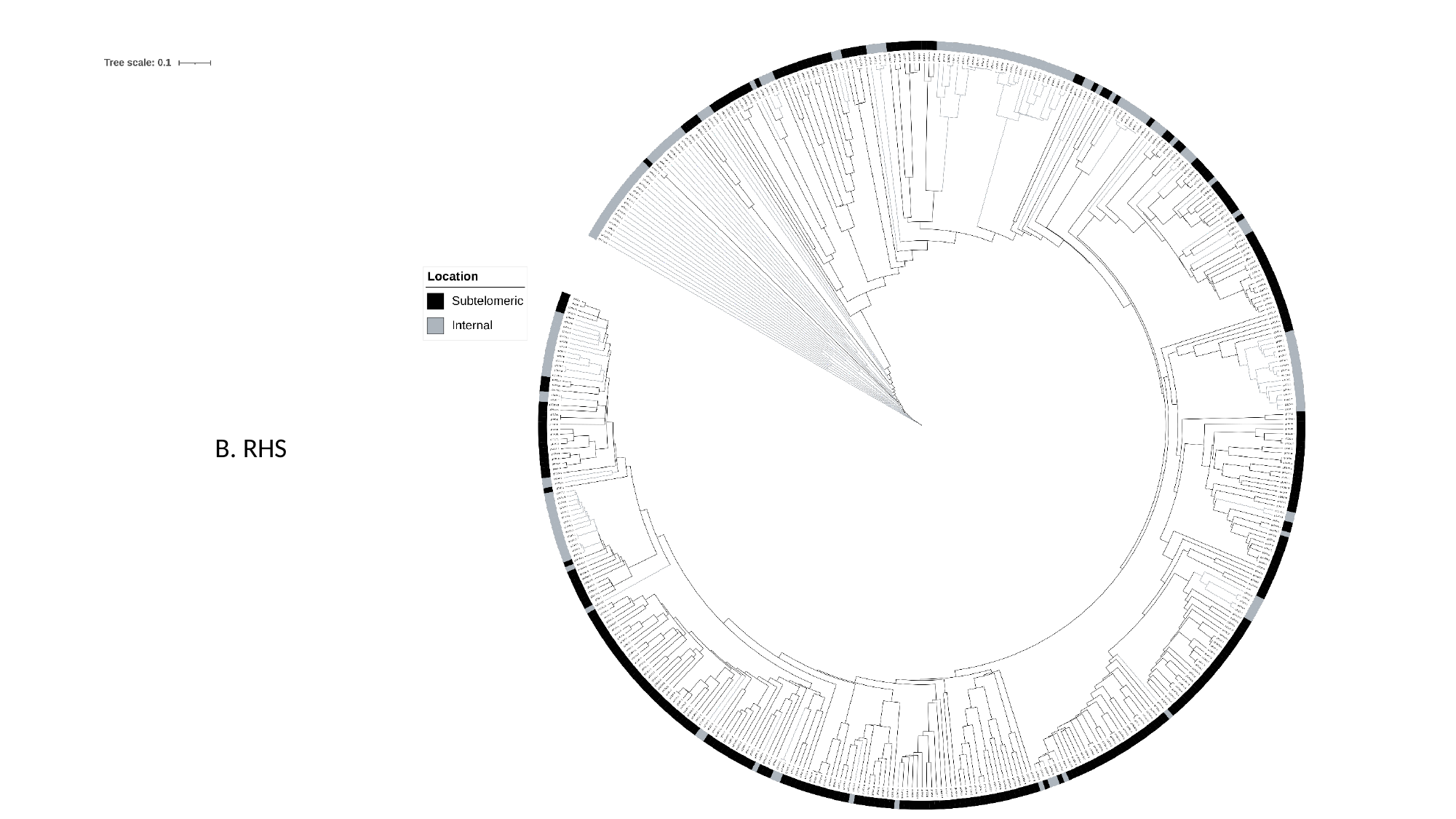

B. RHS
