## Supplementary material for "*Trypanosoma cruzi* has 32 Chromosomes: A Telomere-to-Telomere Assembly Defines Its Karyotype": SupFigureX_DensidadCromoACromo.pptx

### Slide 1
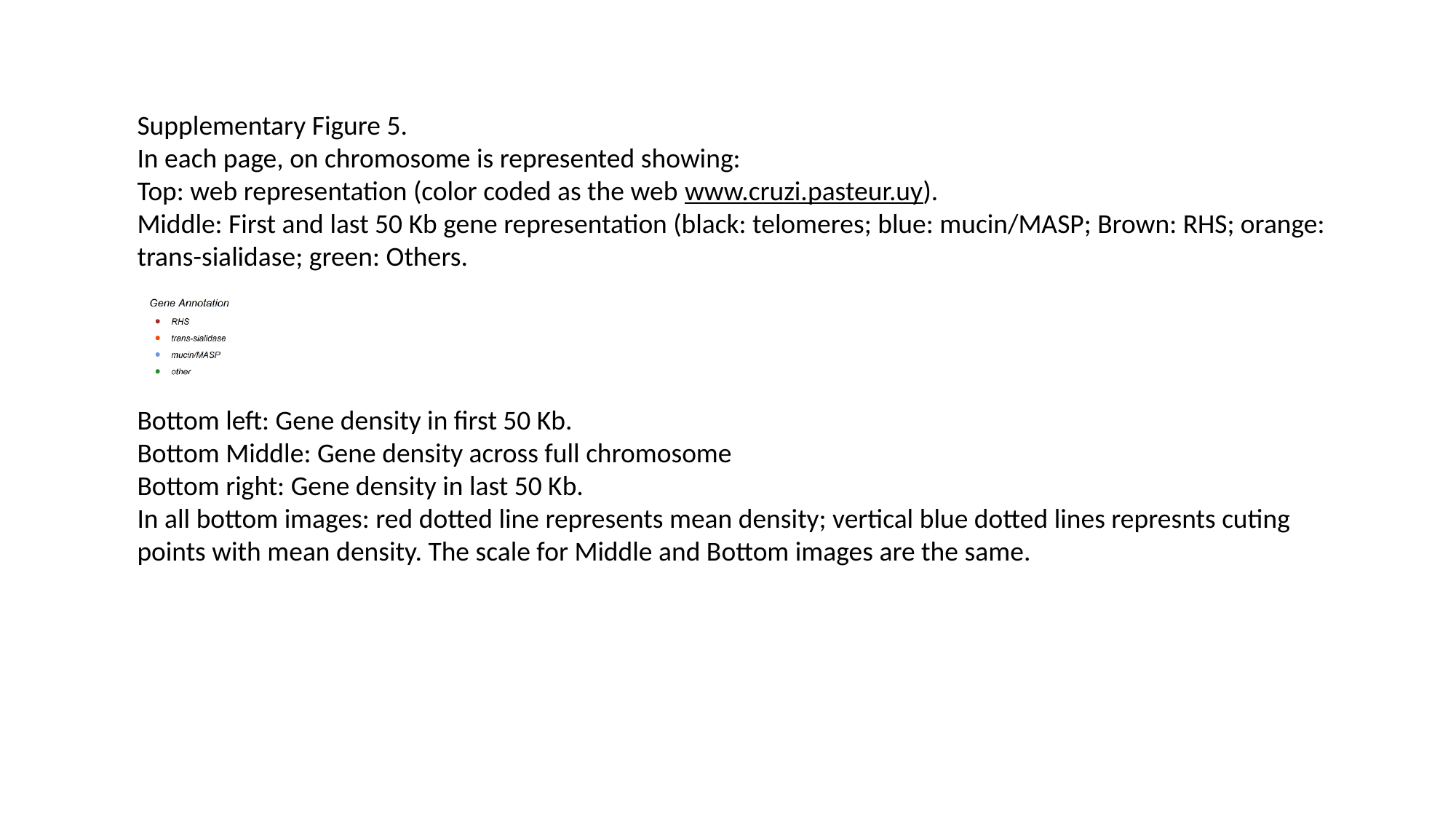

Supplementary Figure 5.
In each page, on chromosome is represented showing:
Top: web representation (color coded as the web www.cruzi.pasteur.uy).
Middle: First and last 50 Kb gene representation (black: telomeres; blue: mucin/MASP; Brown: RHS; orange: trans-sialidase; green: Others.
Bottom left: Gene density in first 50 Kb.
Bottom Middle: Gene density across full chromosome
Bottom right: Gene density in last 50 Kb.
In all bottom images: red dotted line represents mean density; vertical blue dotted lines represnts cuting points with mean density. The scale for Middle and Bottom images are the same.

### Slide 2
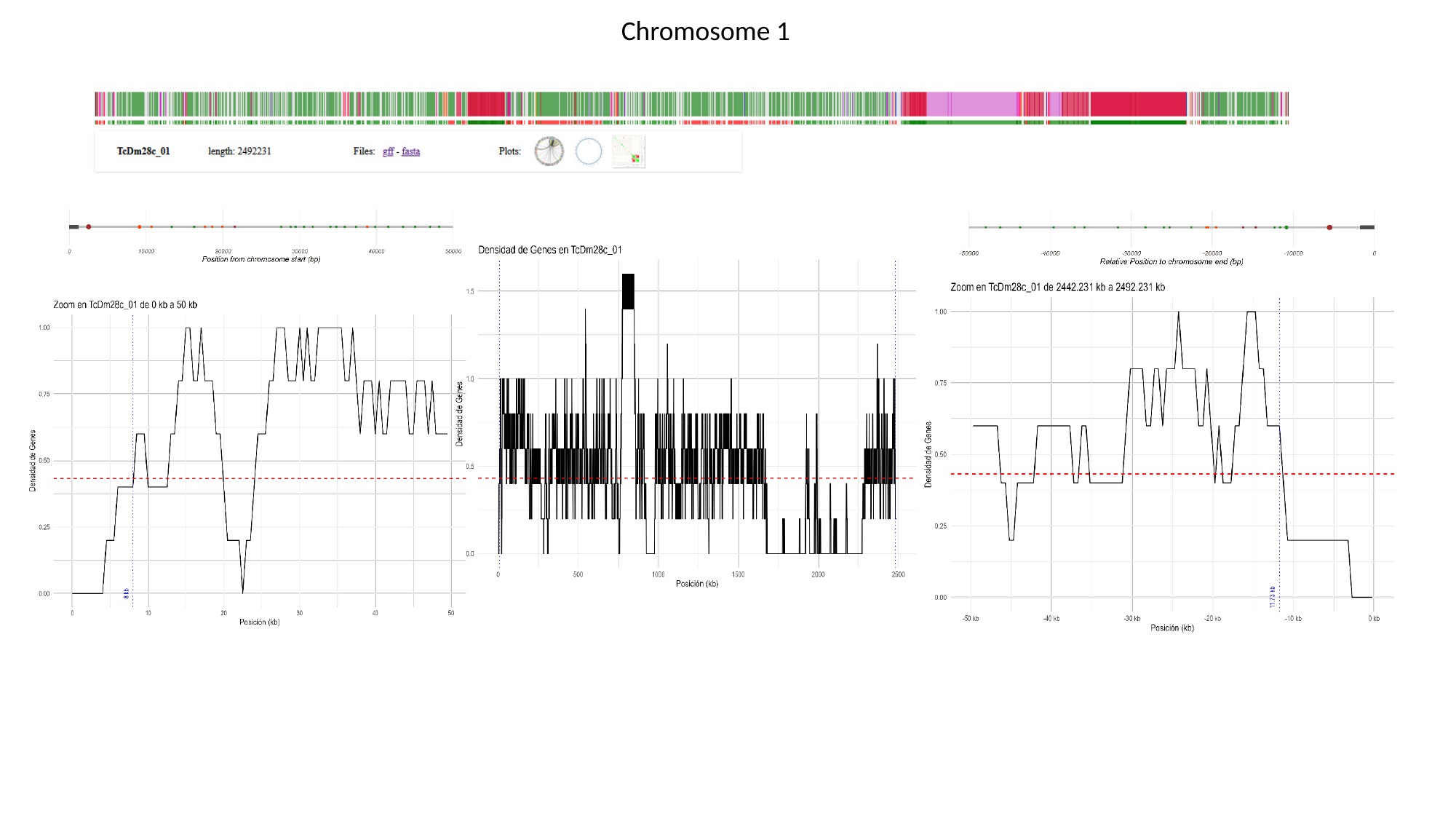

Chromosome 1

### Slide 3
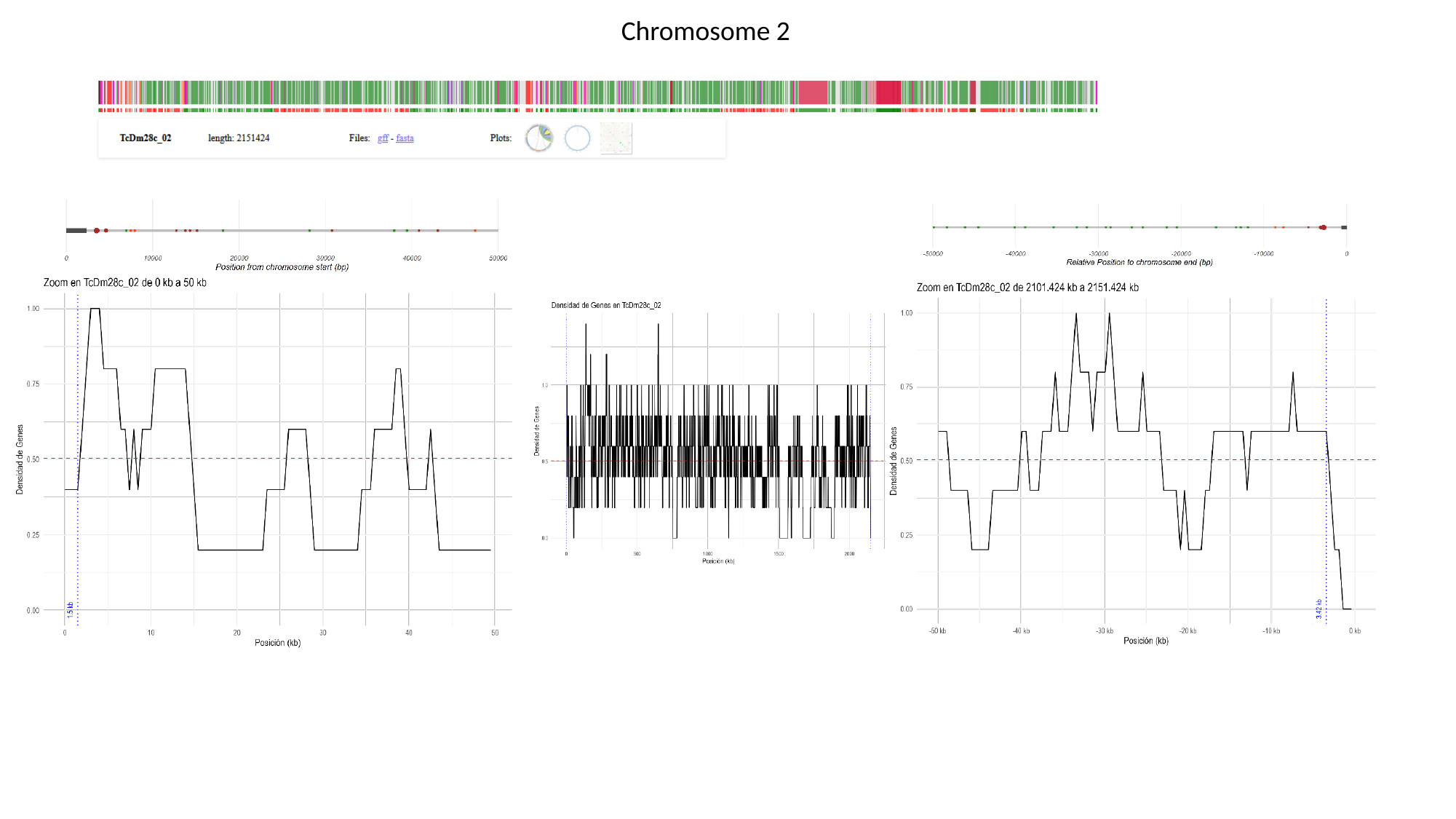

Chromosome 2

### Slide 4
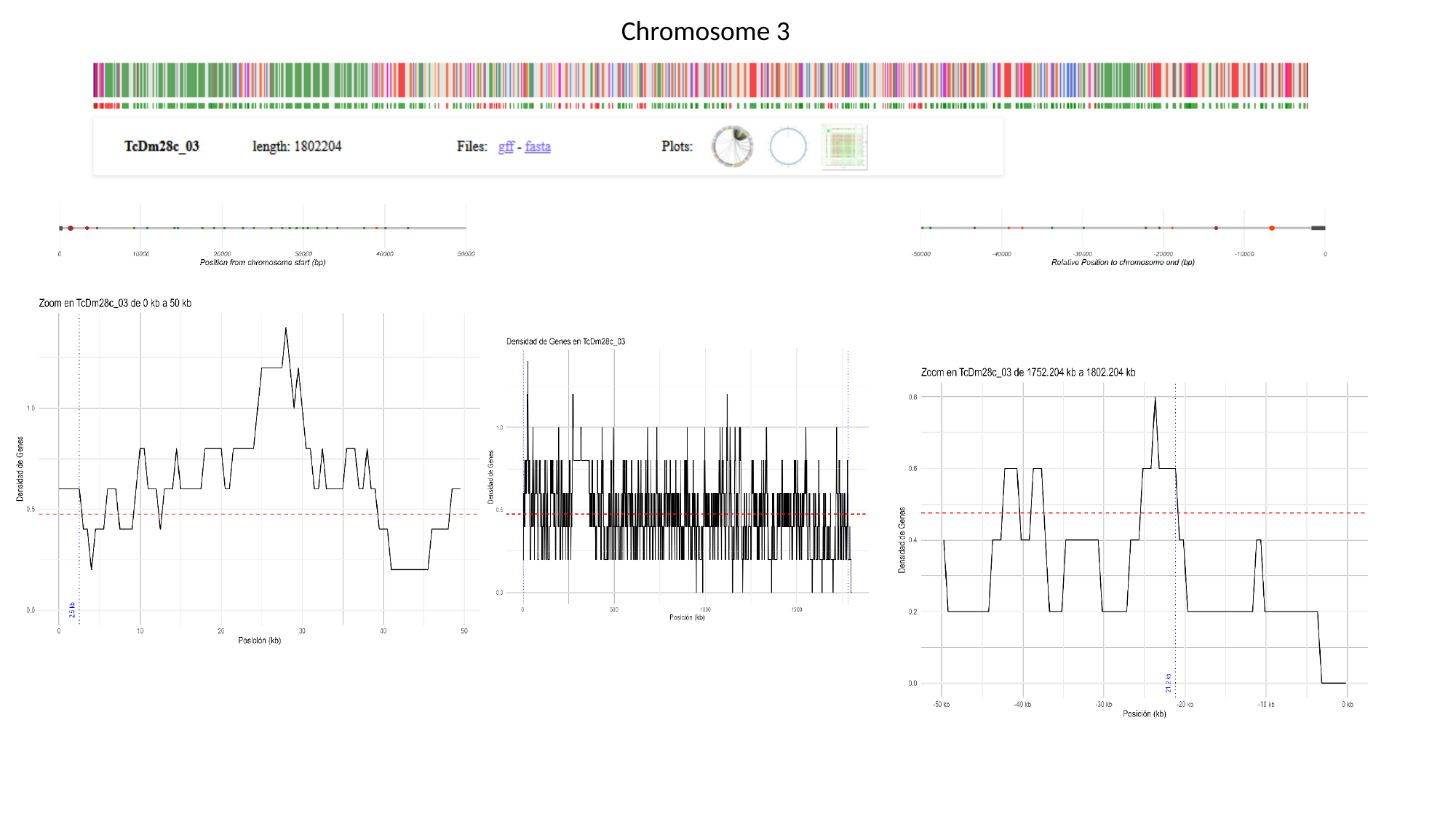

Chromosome 3

### Slide 5
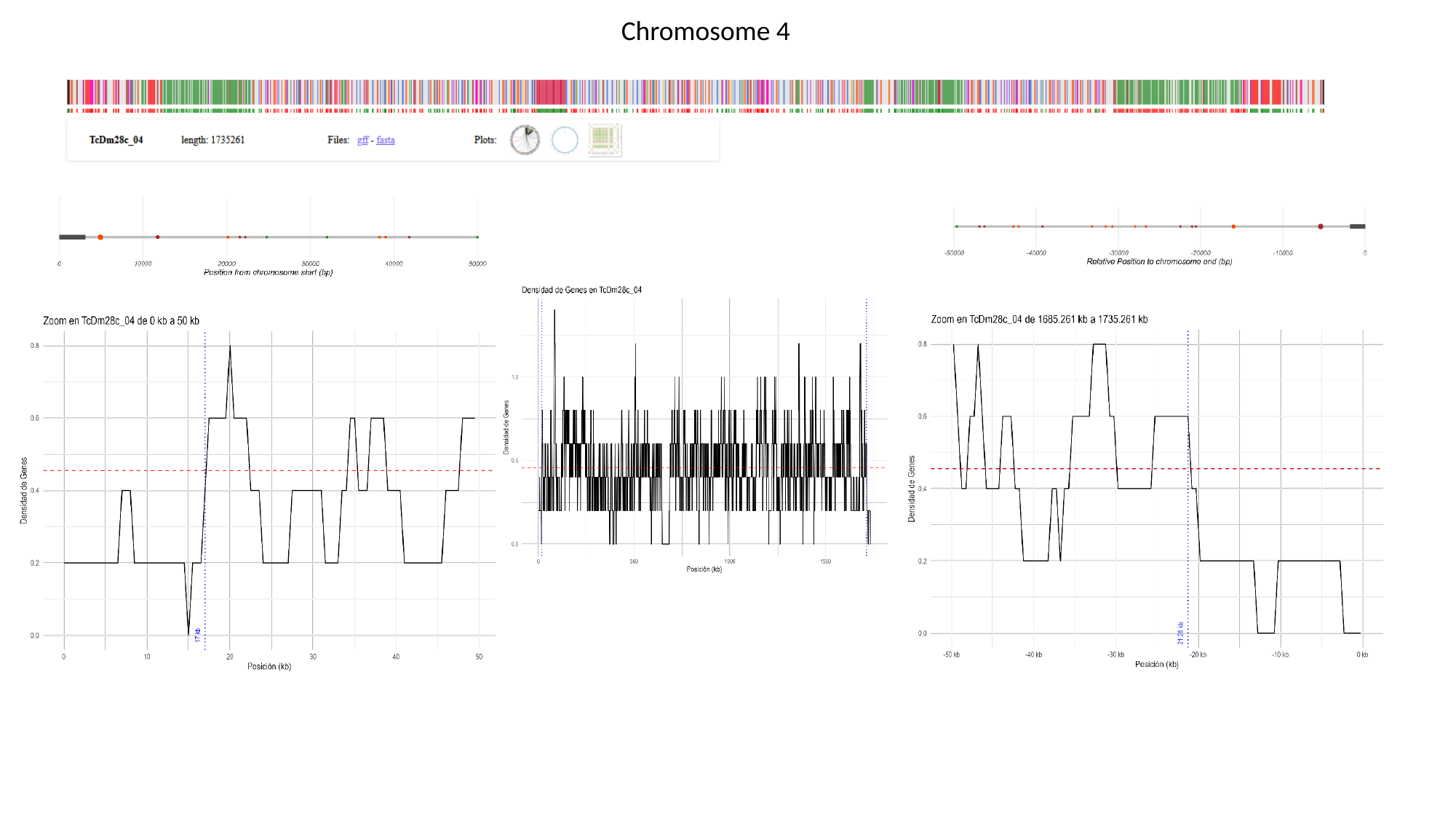

Chromosome 4

### Slide 6
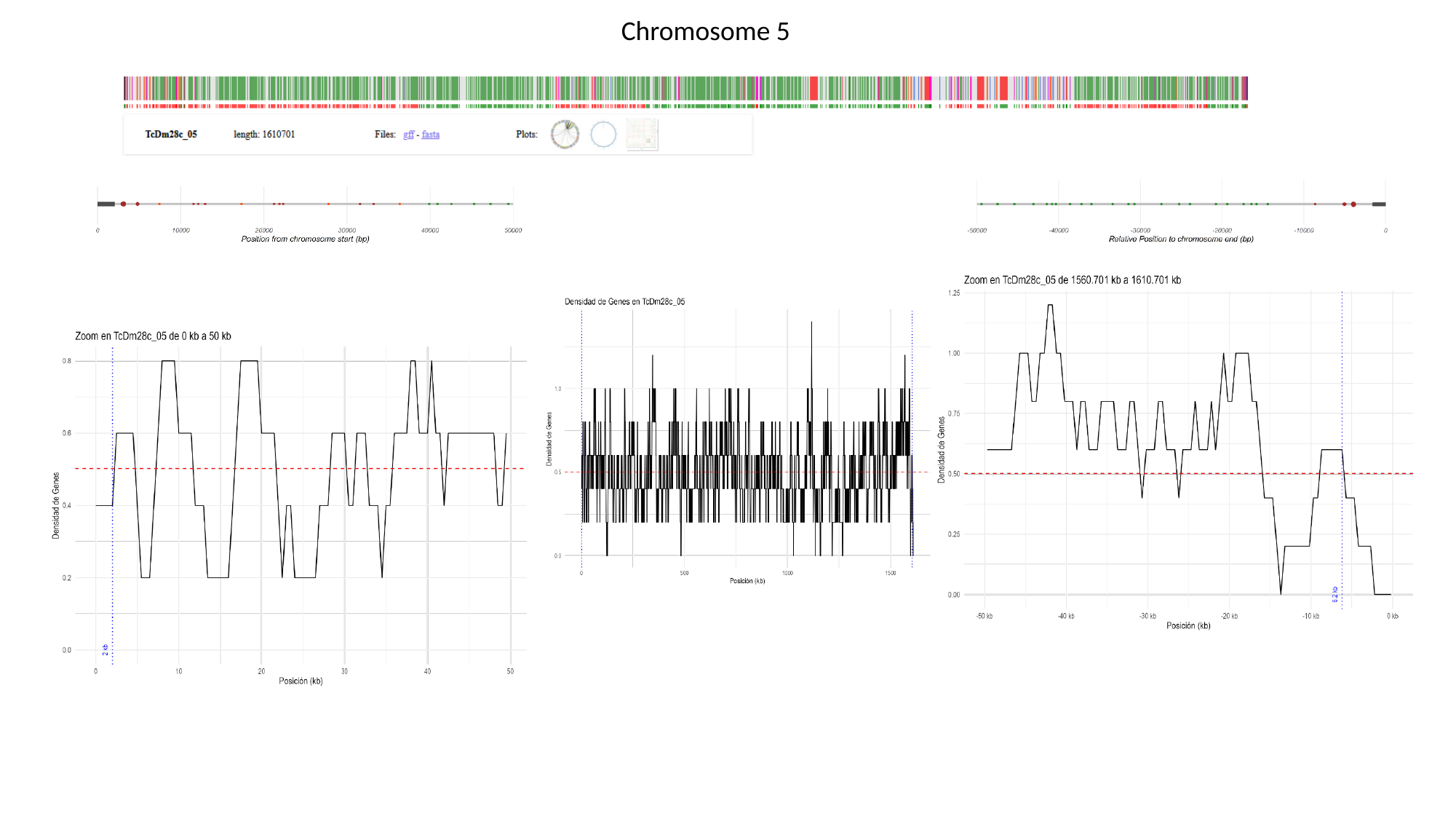

Chromosome 5

### Slide 7
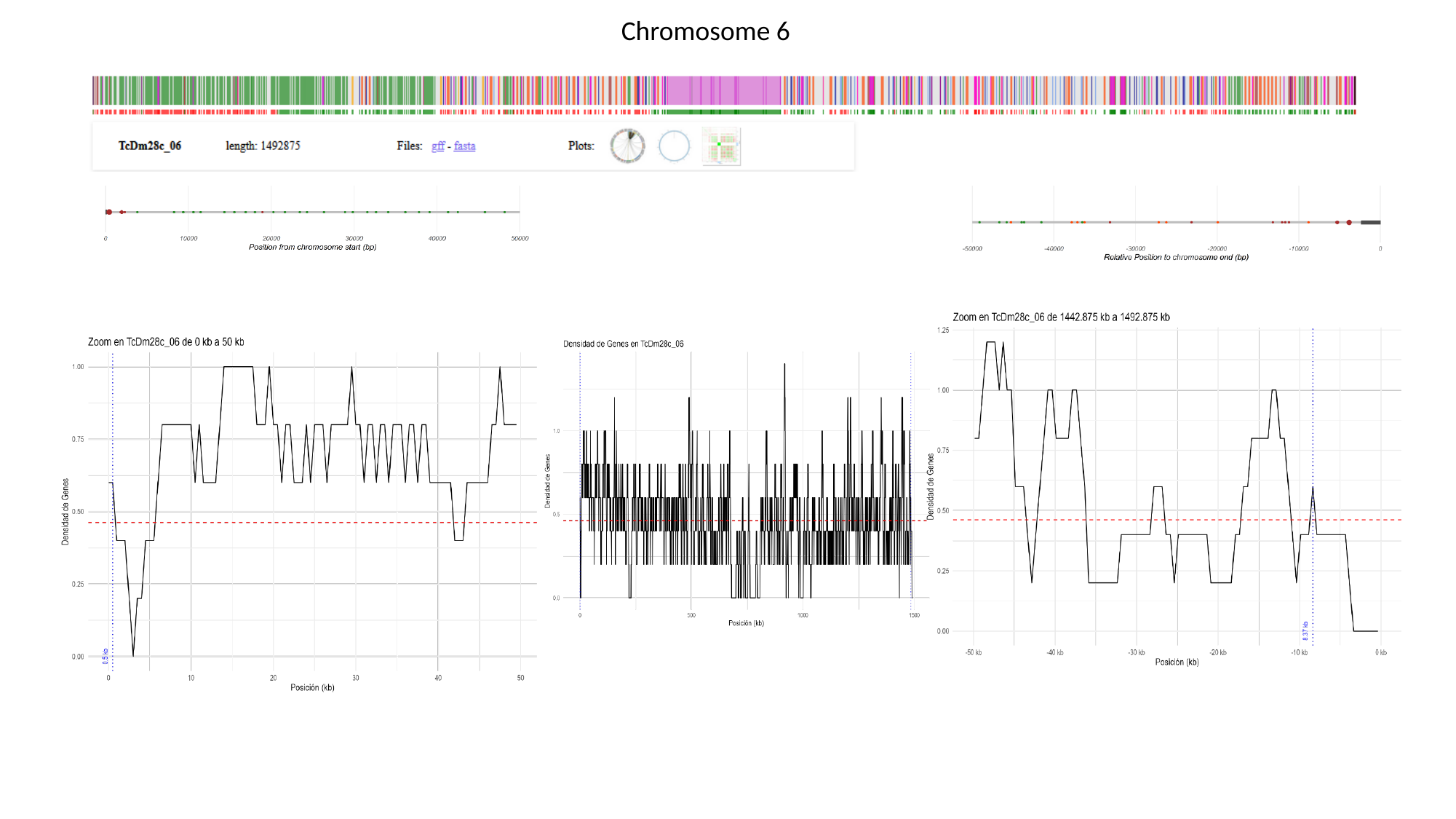

Chromosome 6

### Slide 8
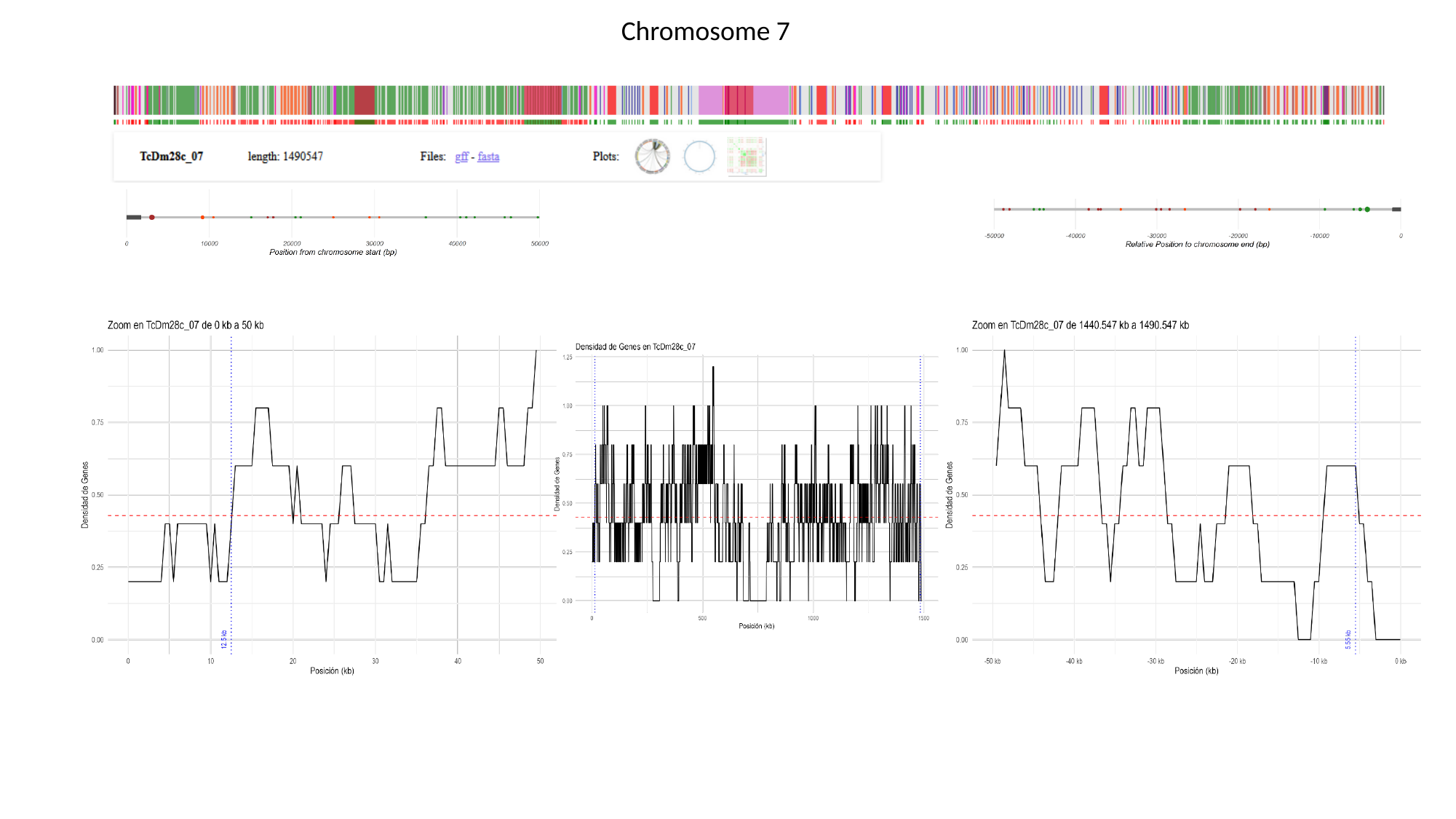

Chromosome 7

### Slide 9
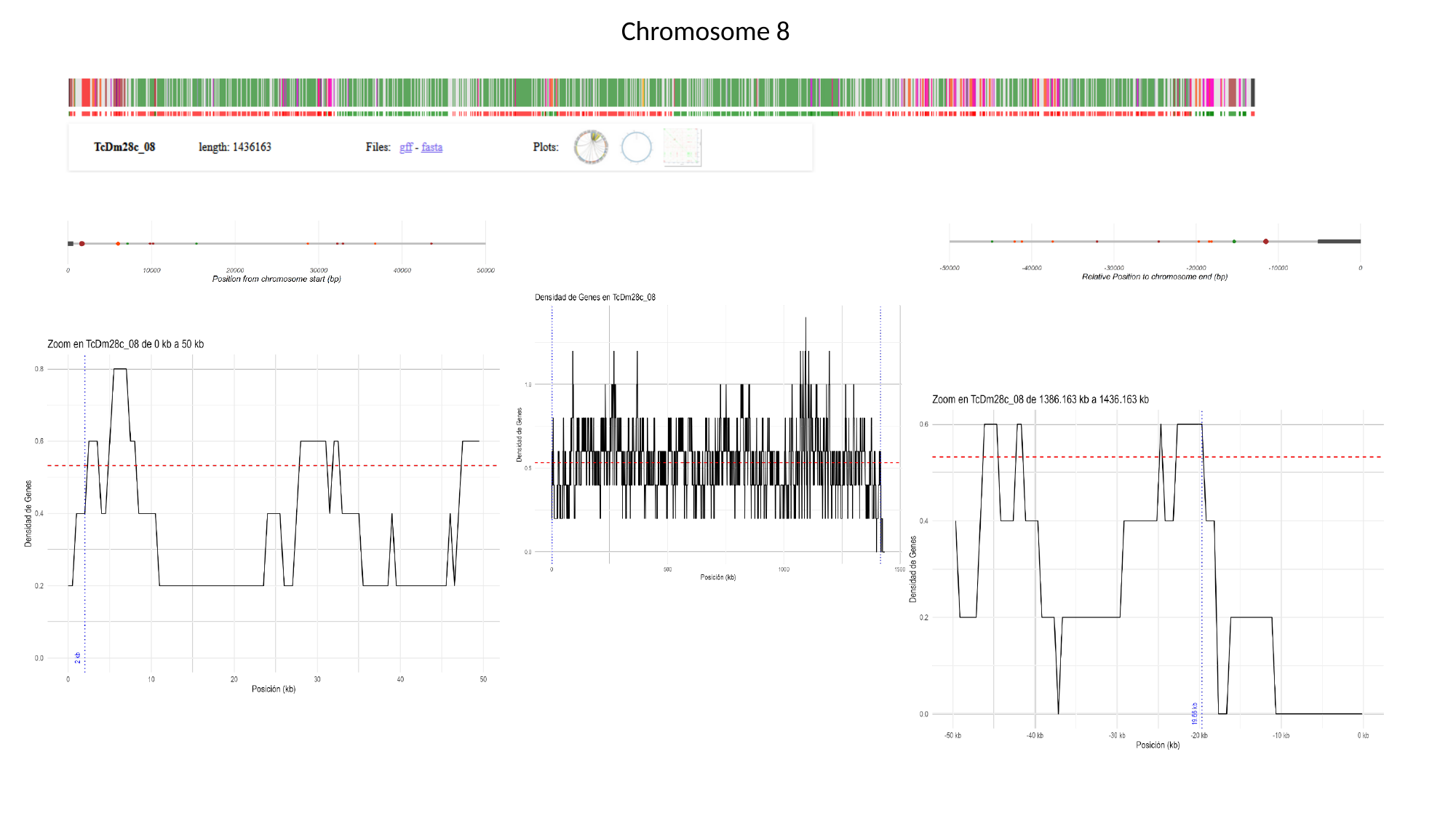

Chromosome 8

### Slide 10
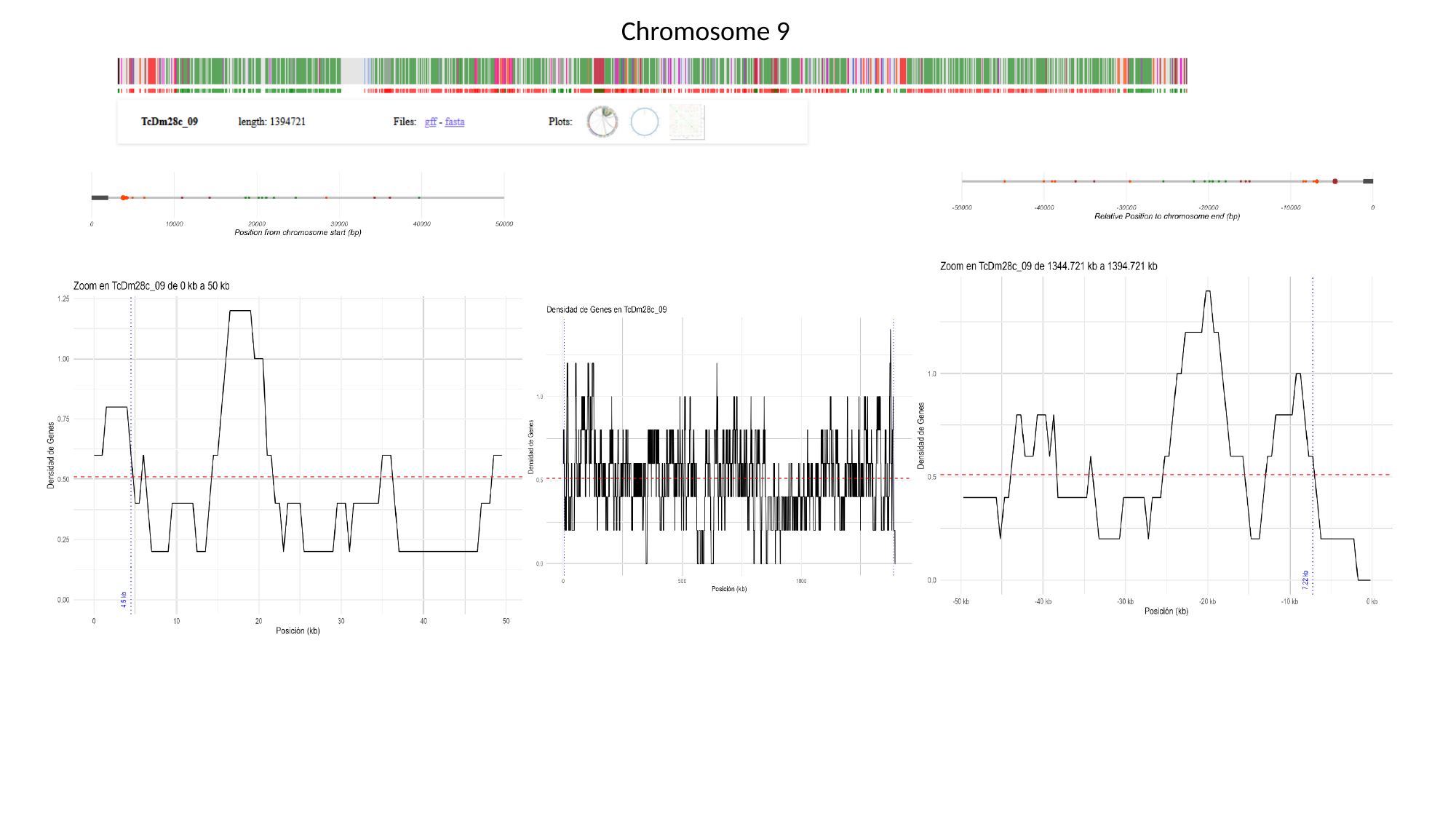

Chromosome 9

### Slide 11
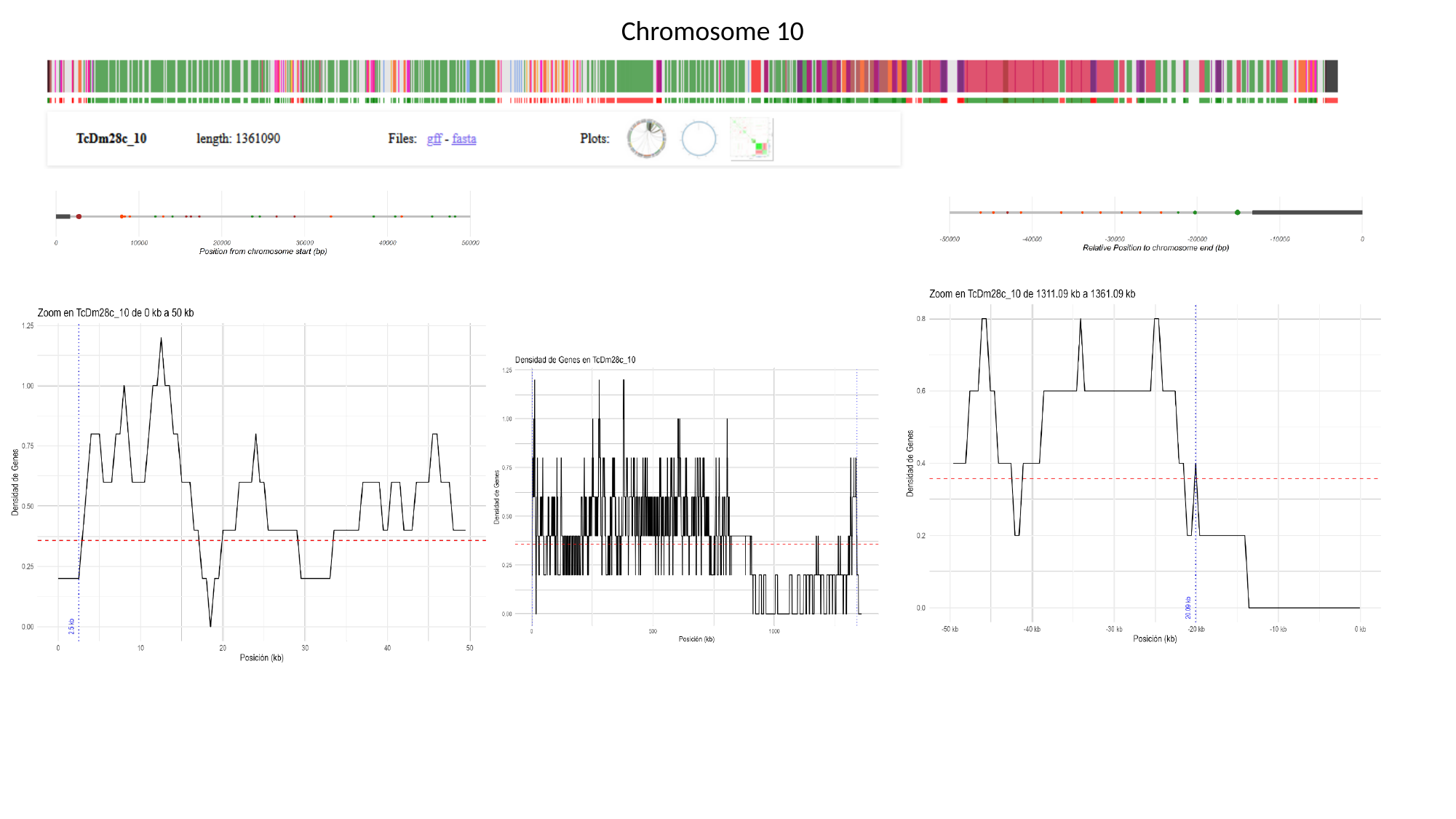

Chromosome 10

### Slide 12
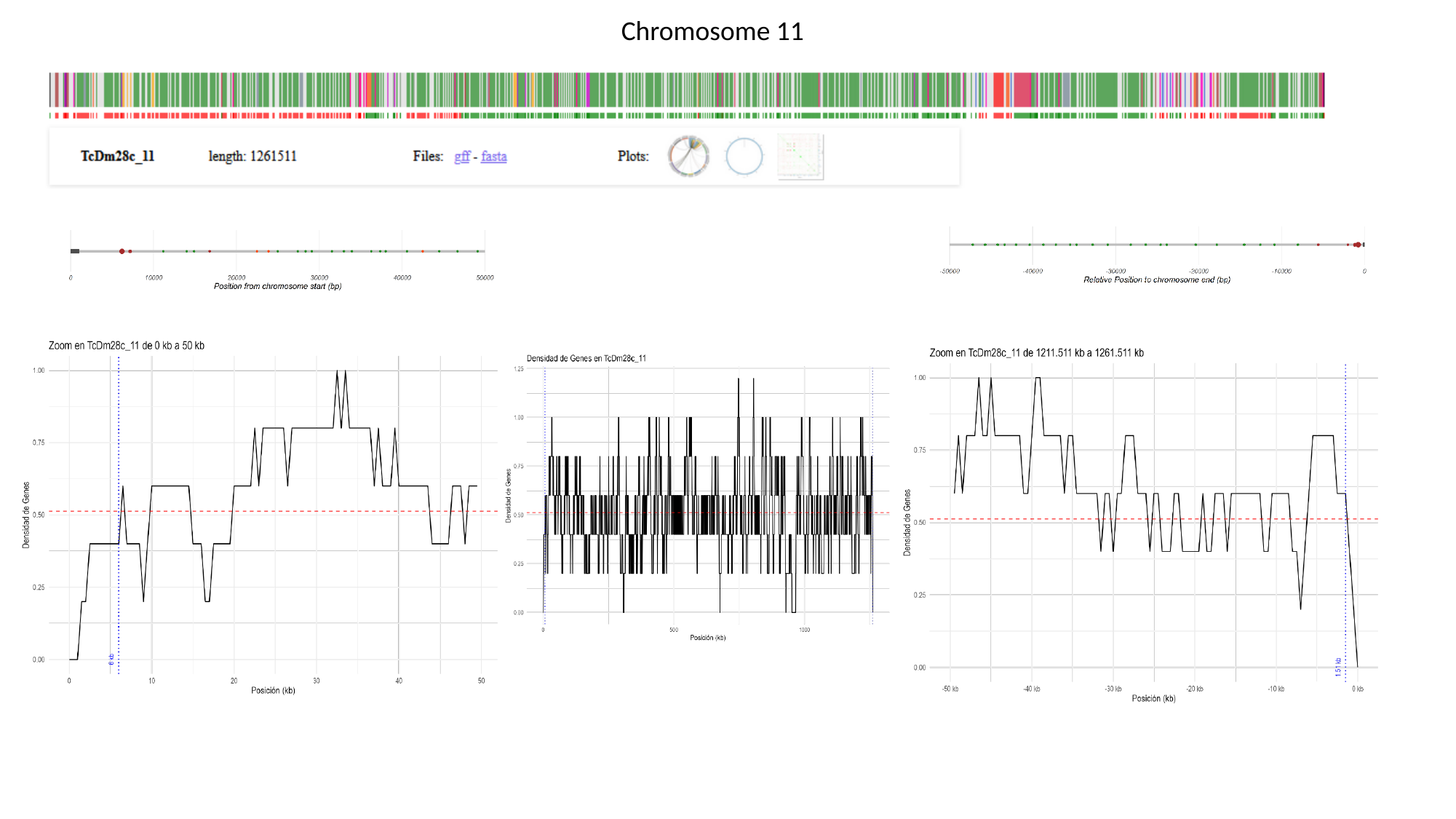

Chromosome 11

### Slide 13
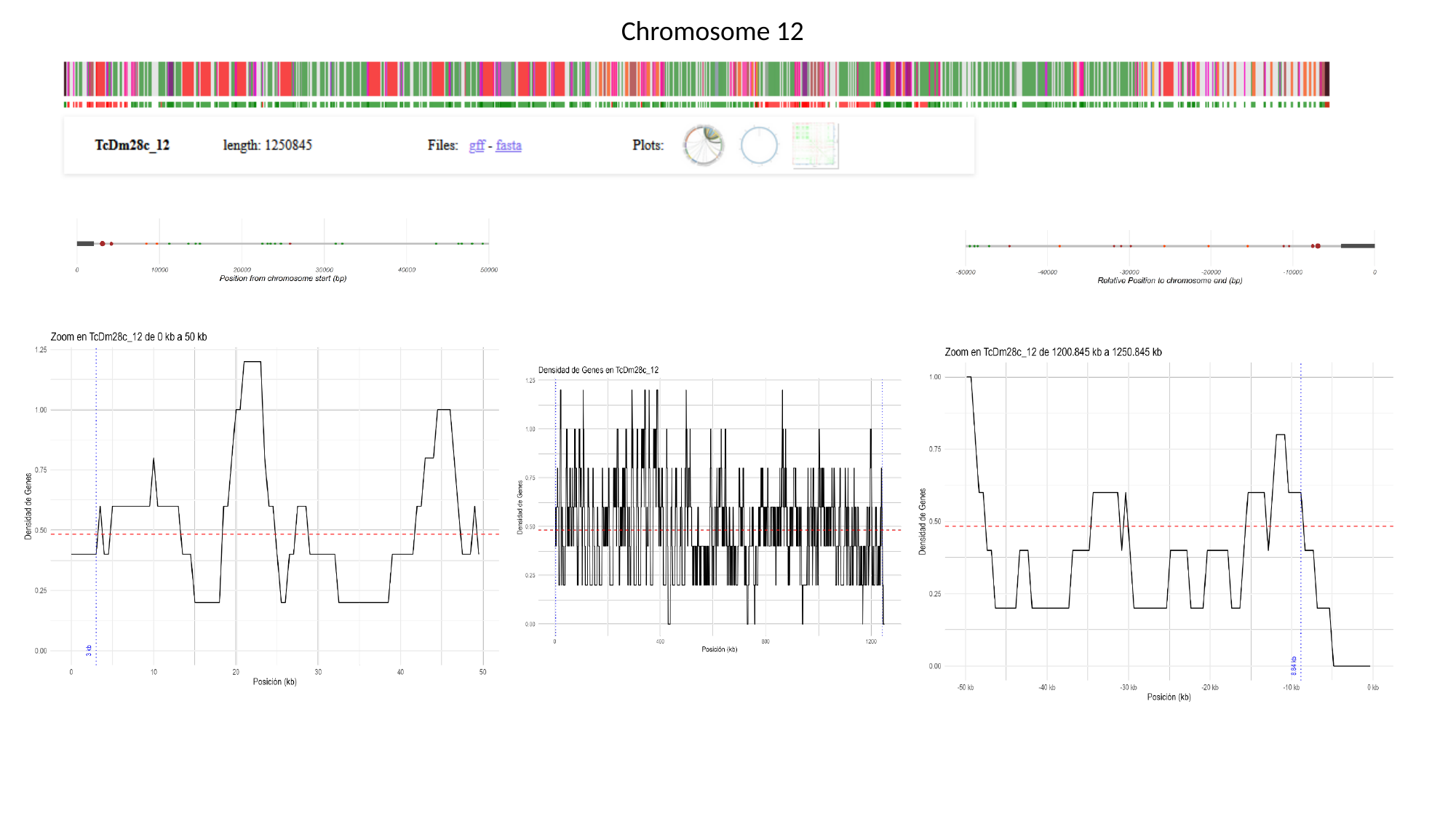

Chromosome 12

### Slide 14
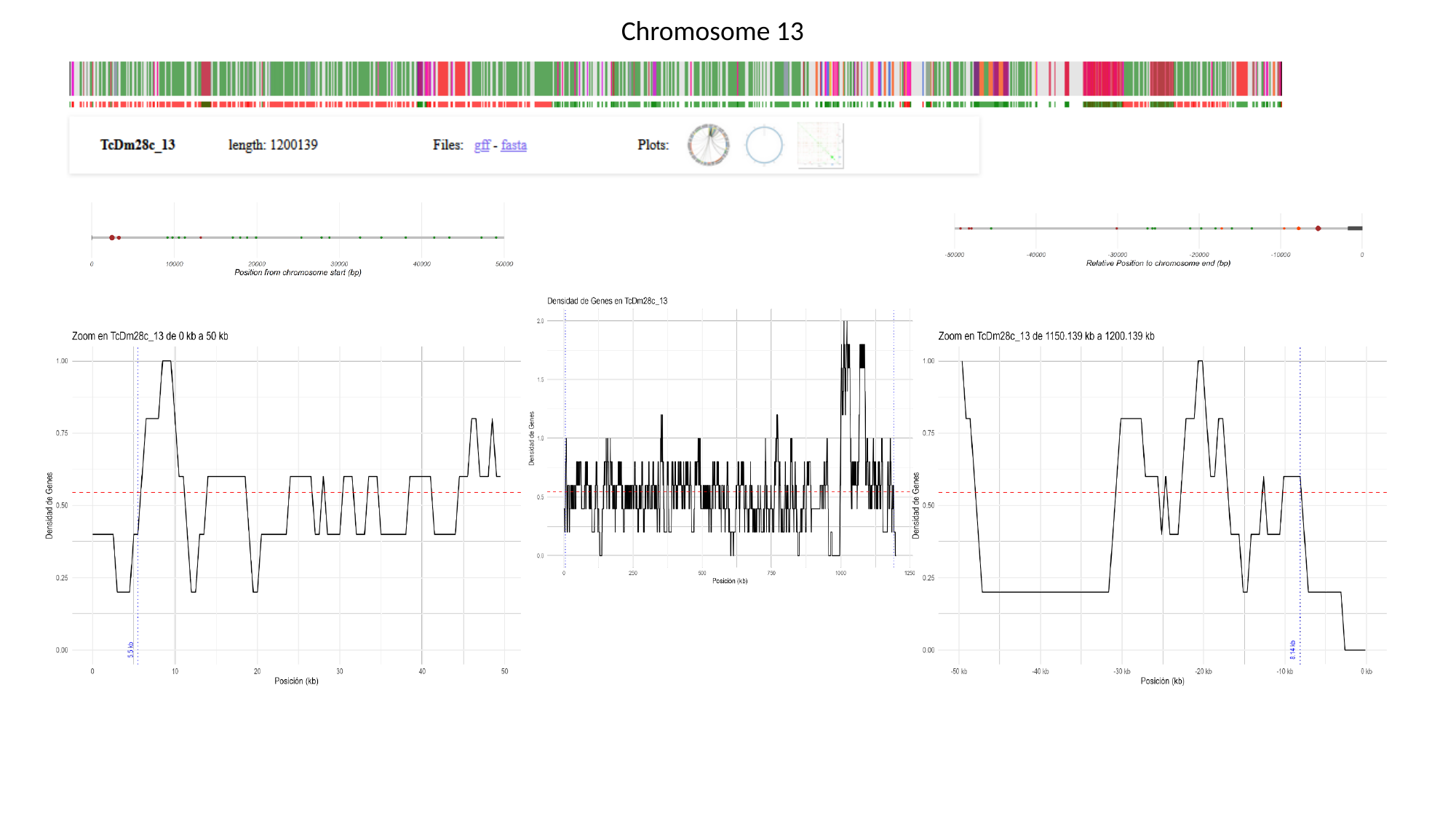

Chromosome 13

### Slide 15
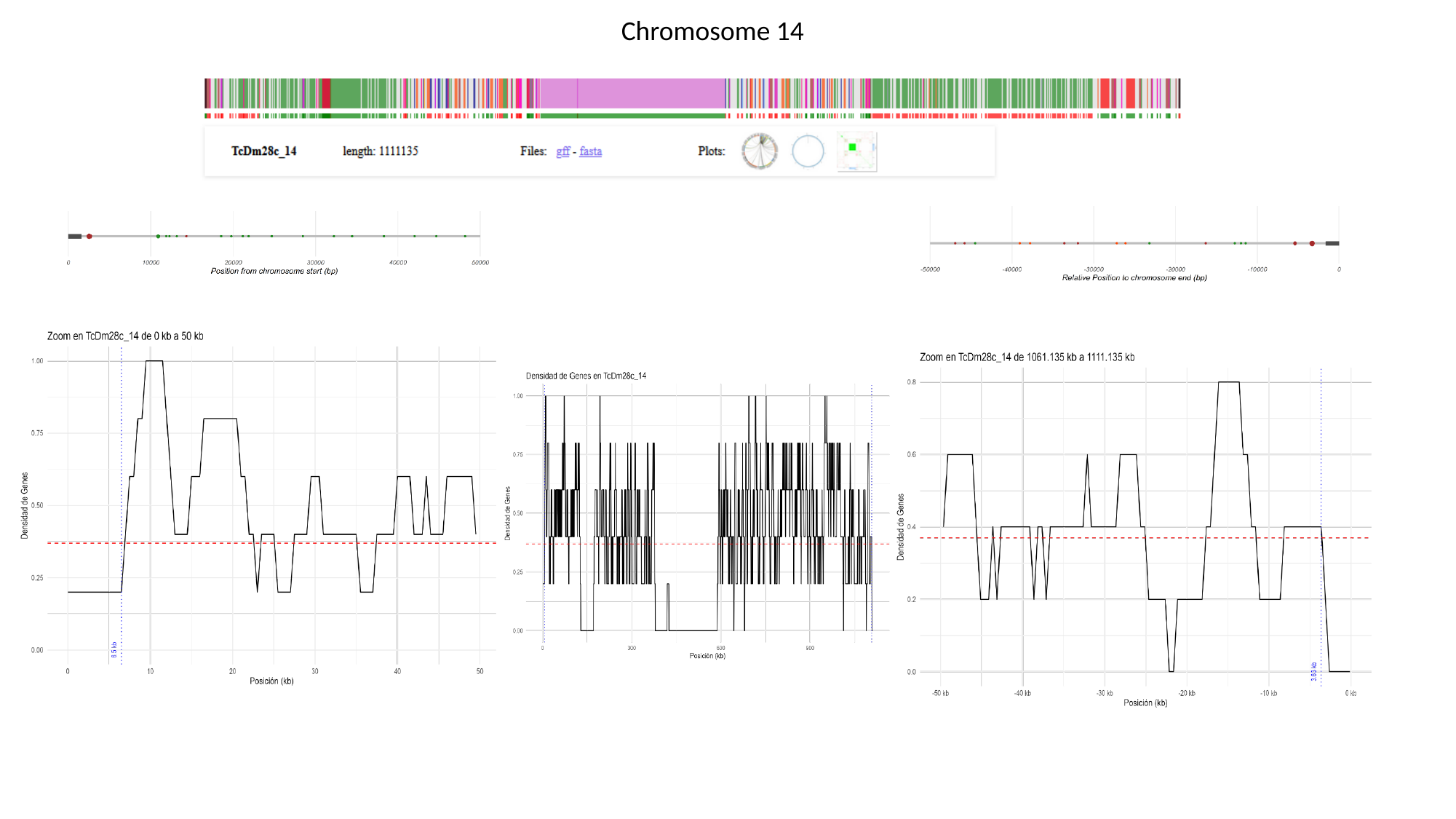

Chromosome 14

### Slide 16
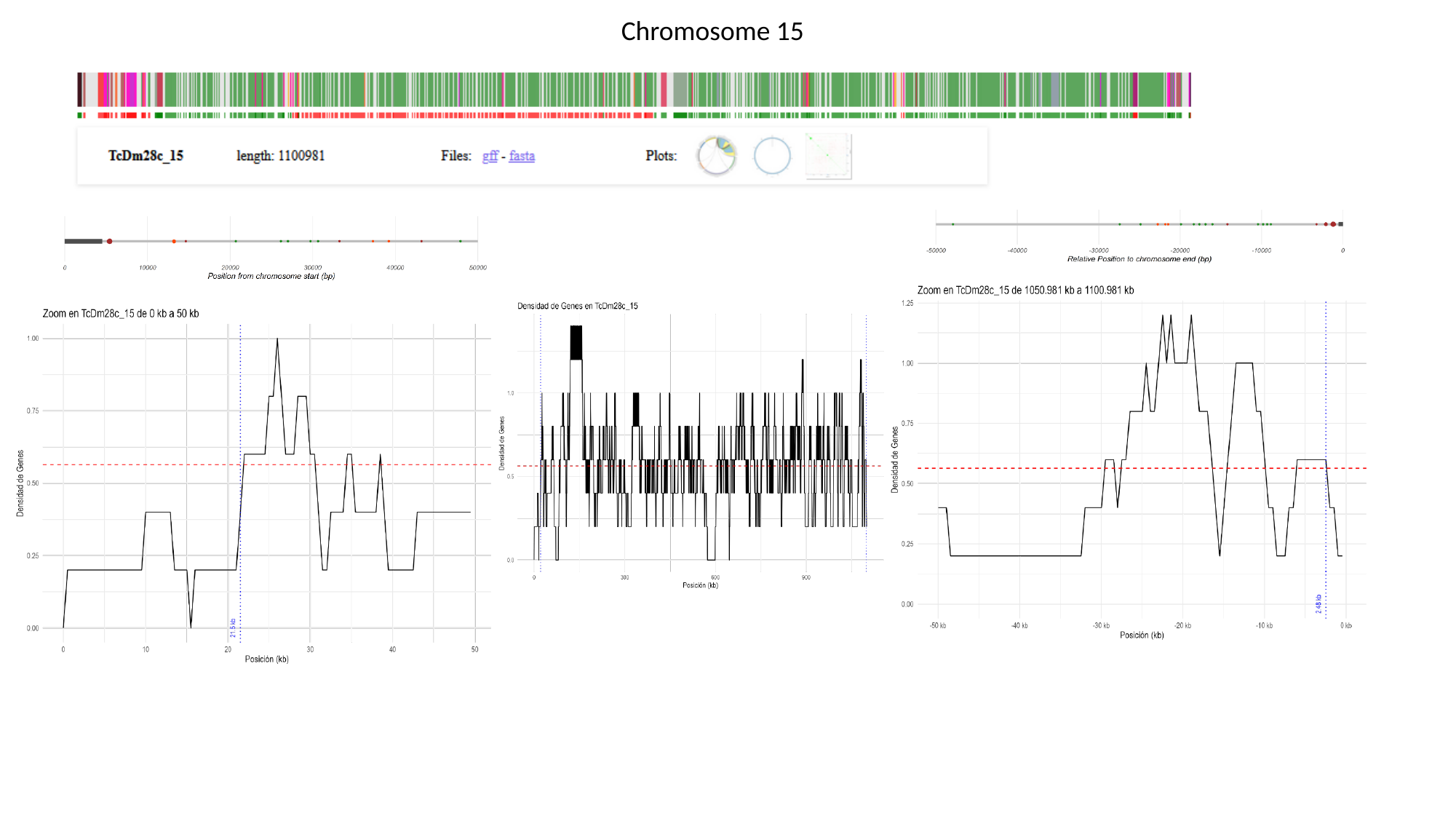

Chromosome 15

### Slide 17
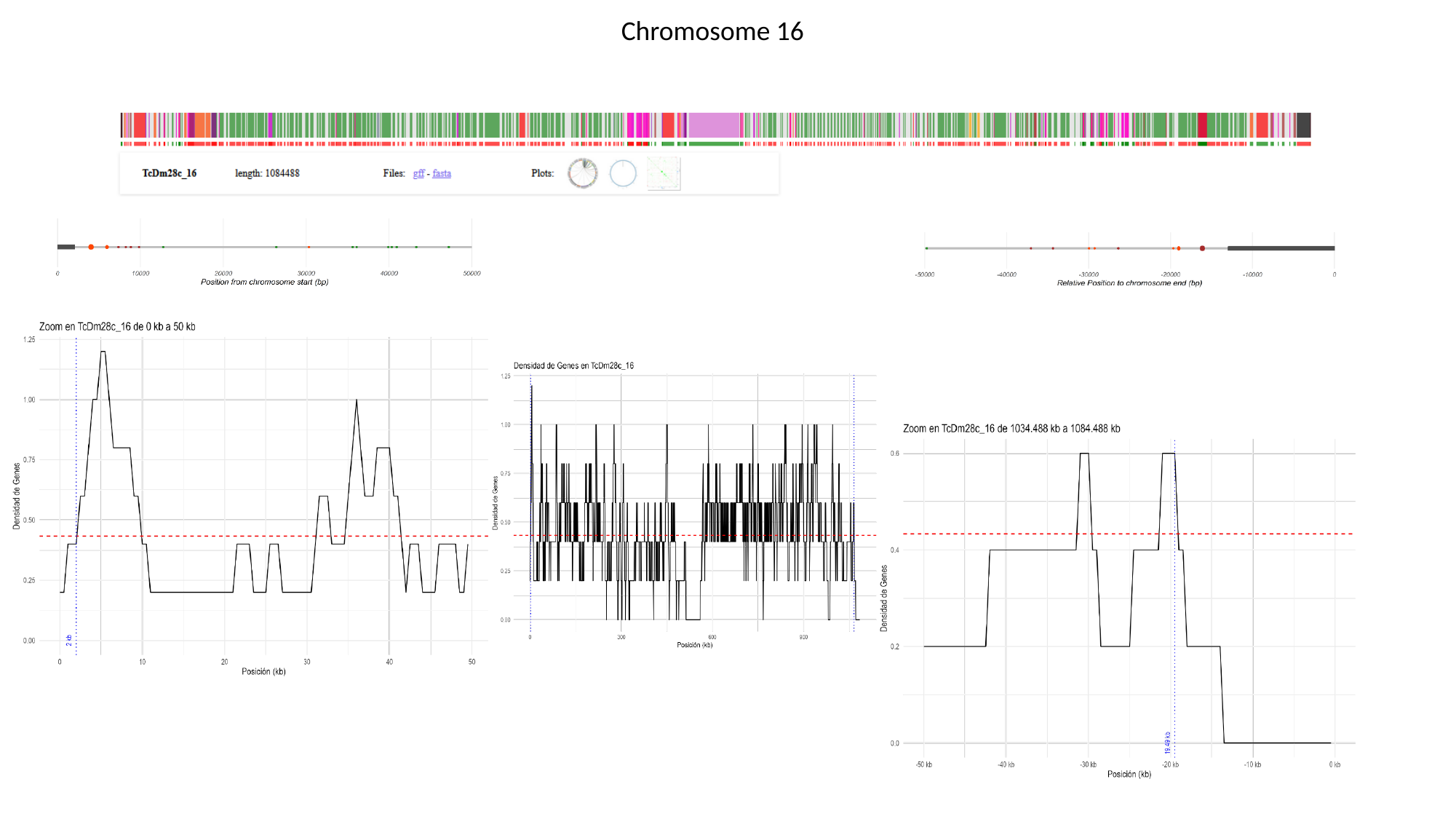

Chromosome 16

### Slide 18
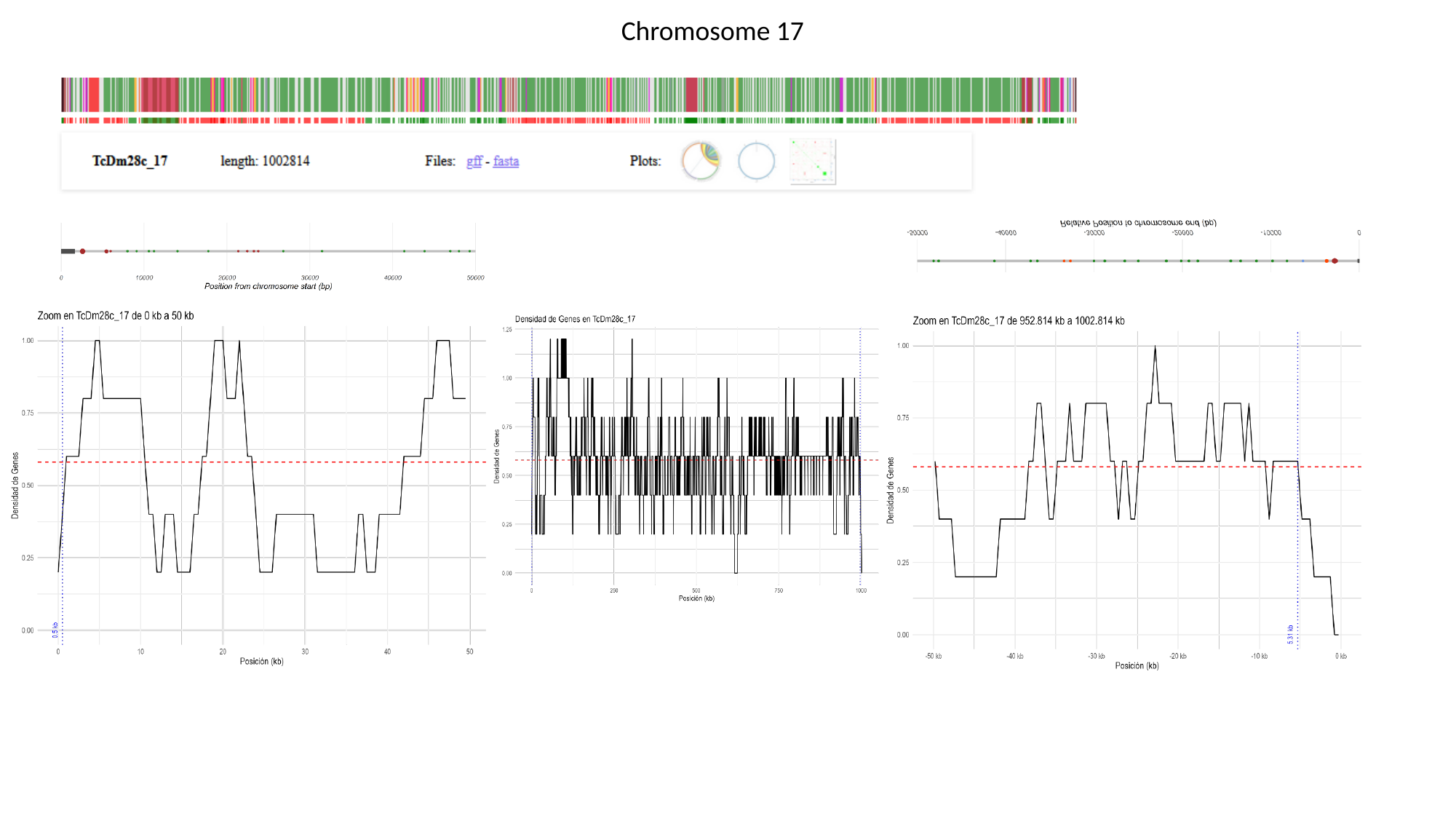

Chromosome 17

### Slide 19
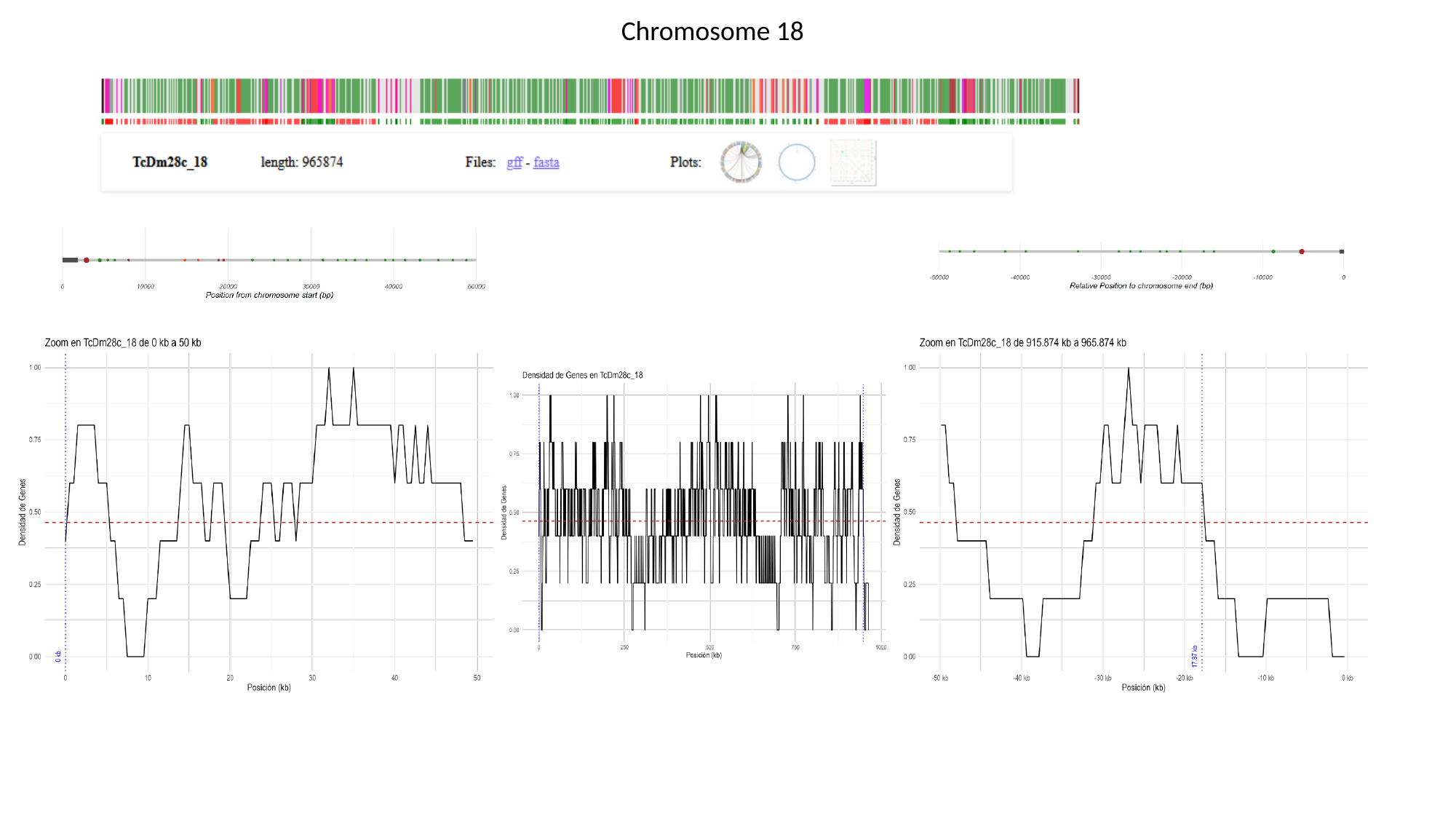

Chromosome 18

### Slide 20
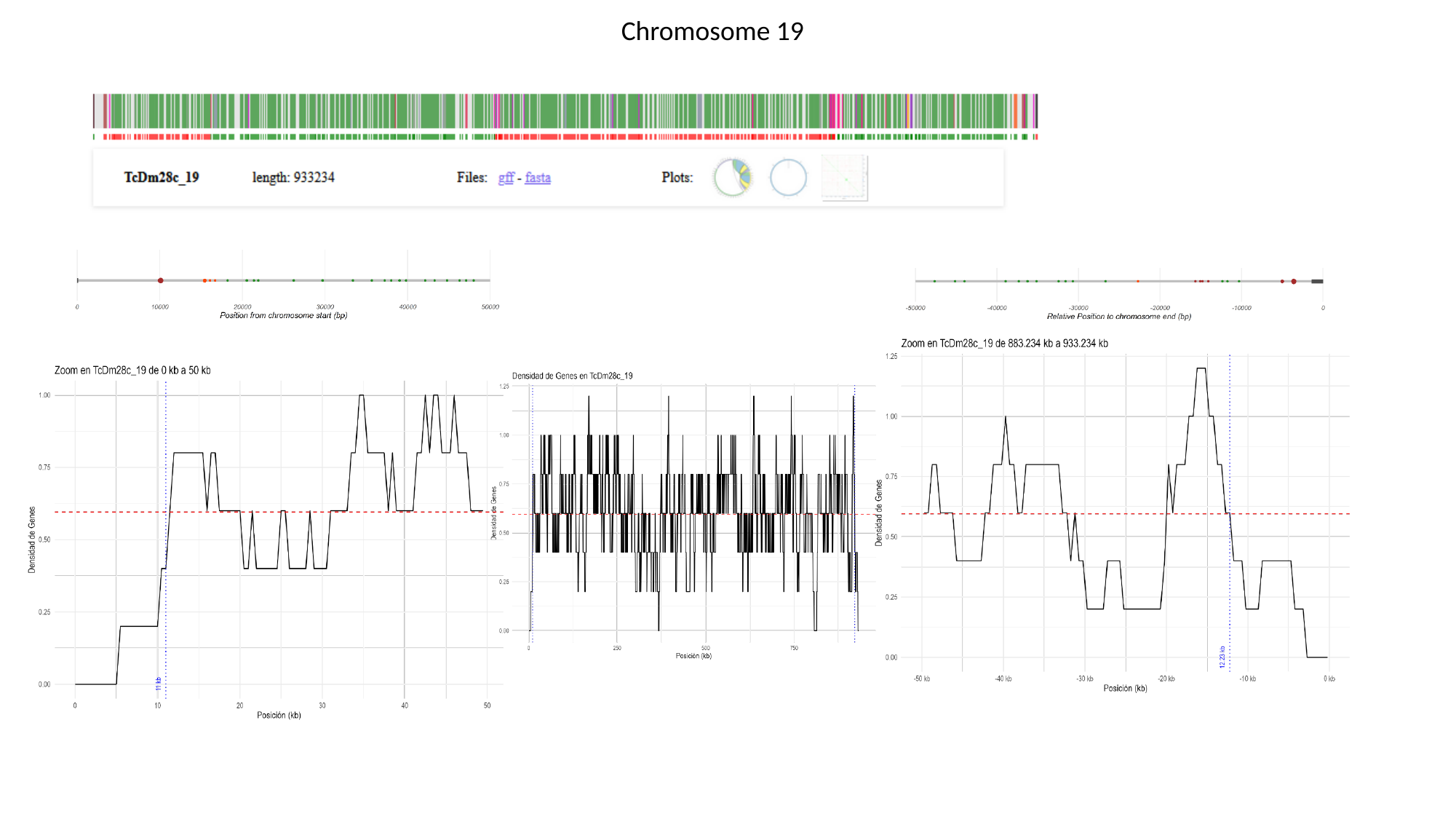

Chromosome 19

### Slide 21
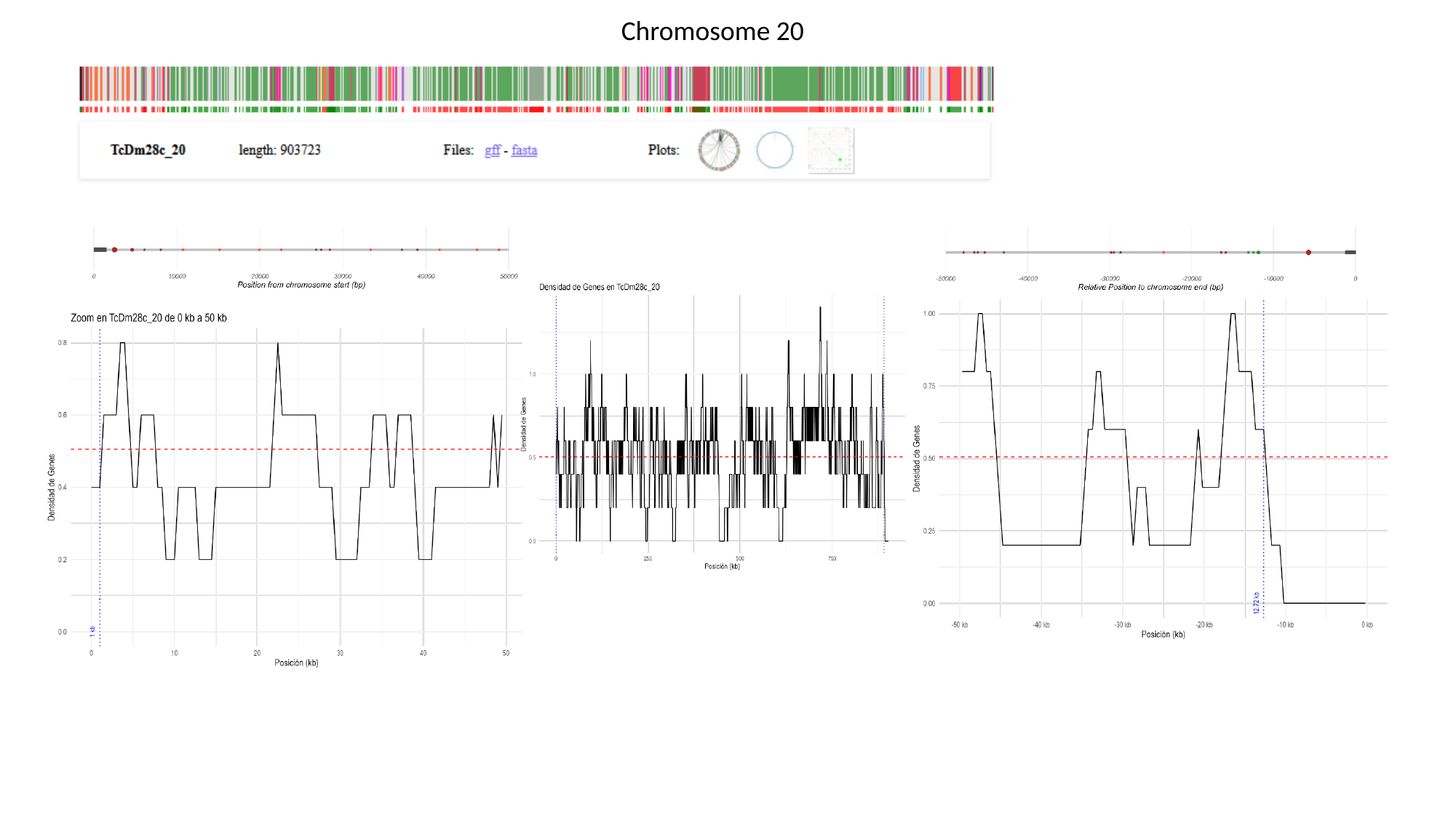

Chromosome 20

### Slide 22
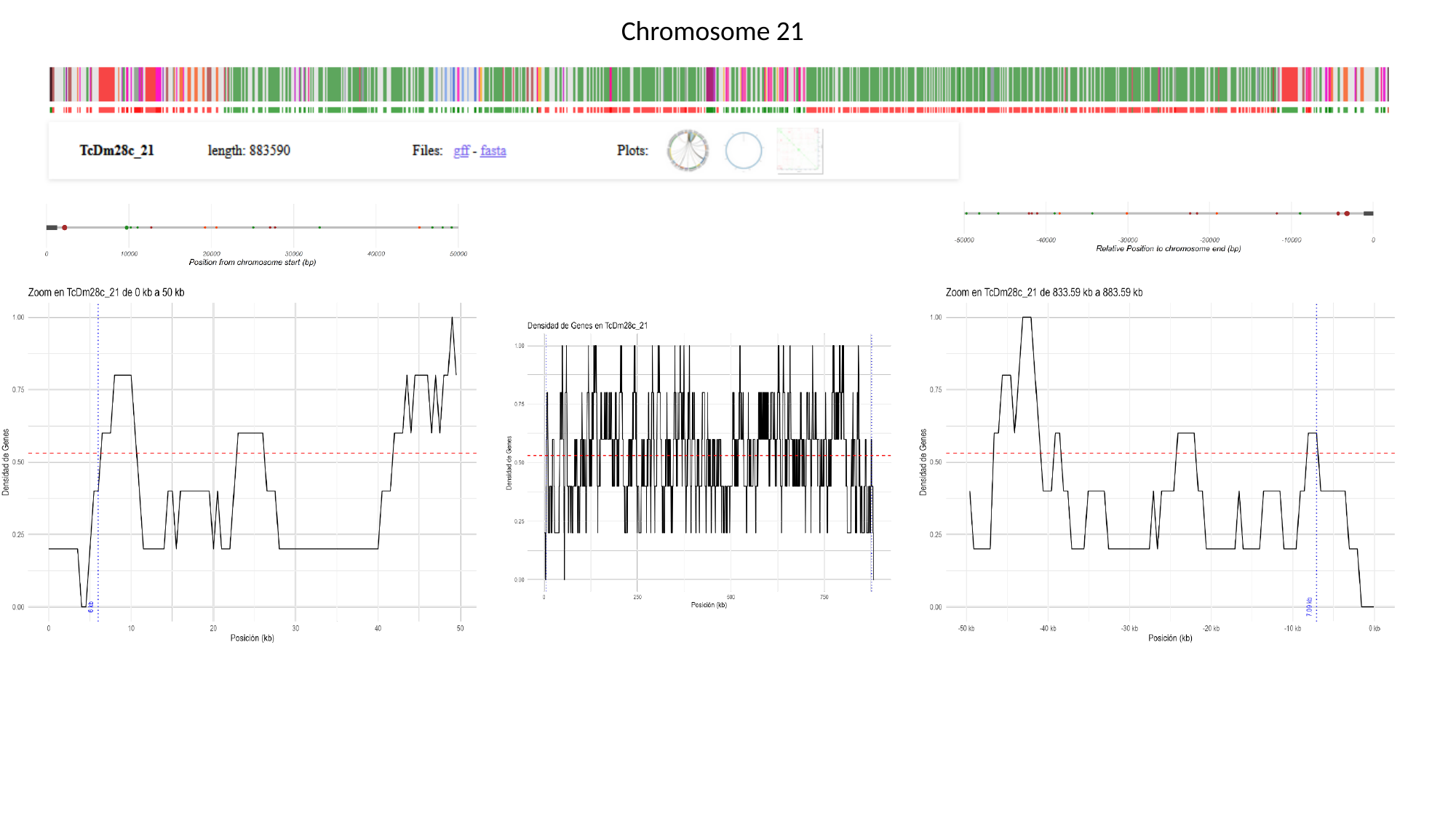

Chromosome 21

### Slide 23
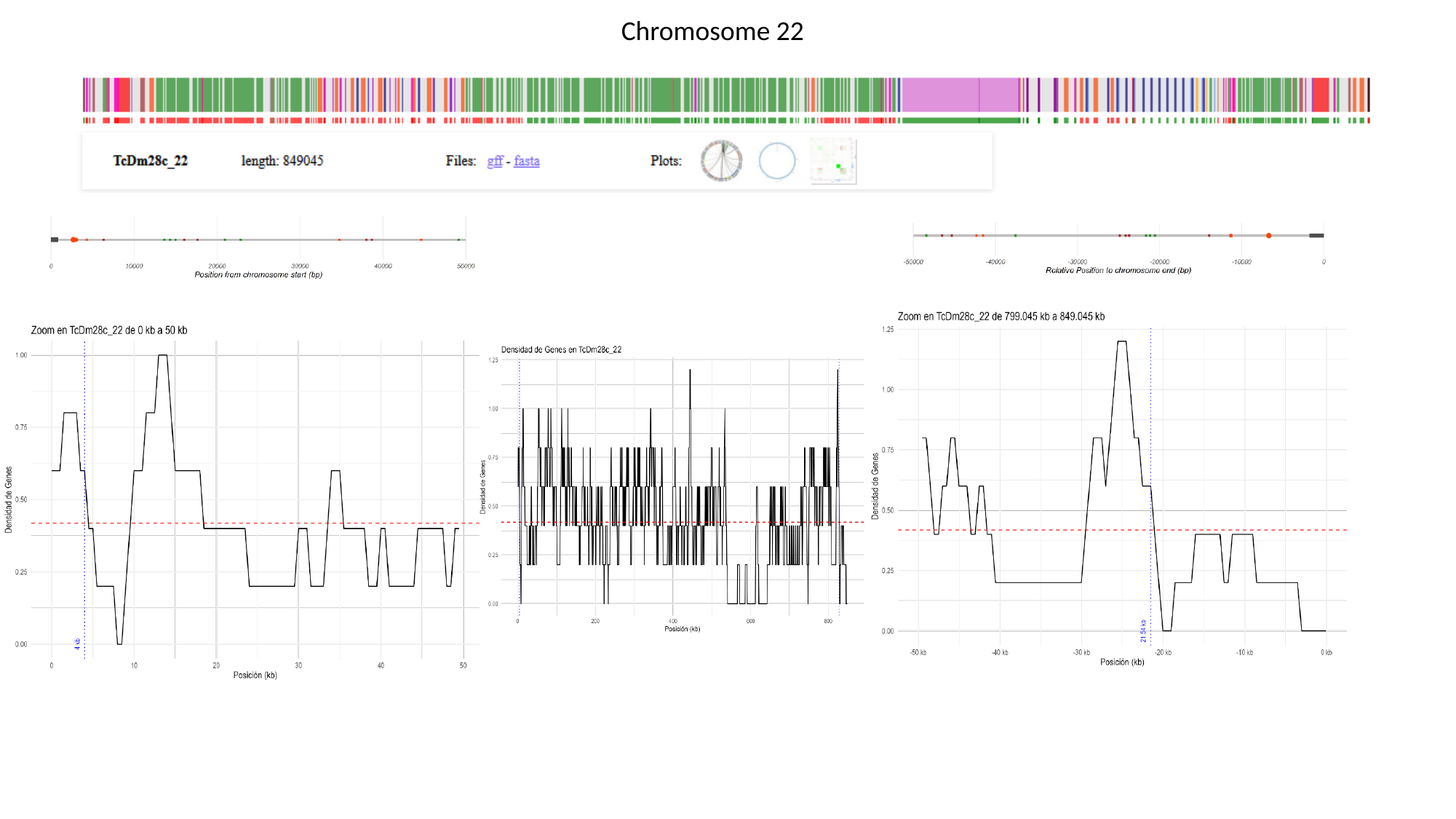

Chromosome 22

### Slide 24
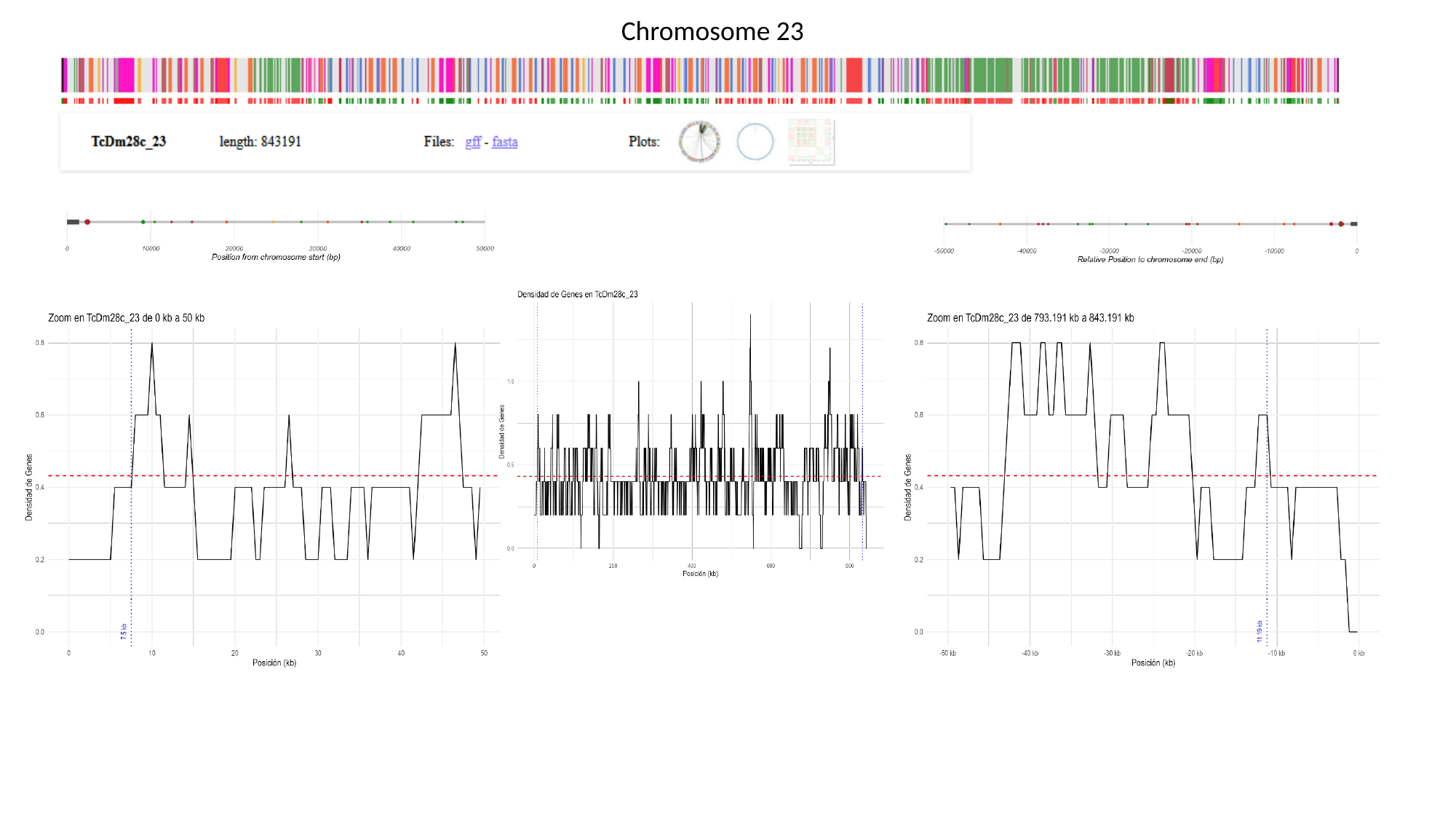

Chromosome 23

### Slide 25
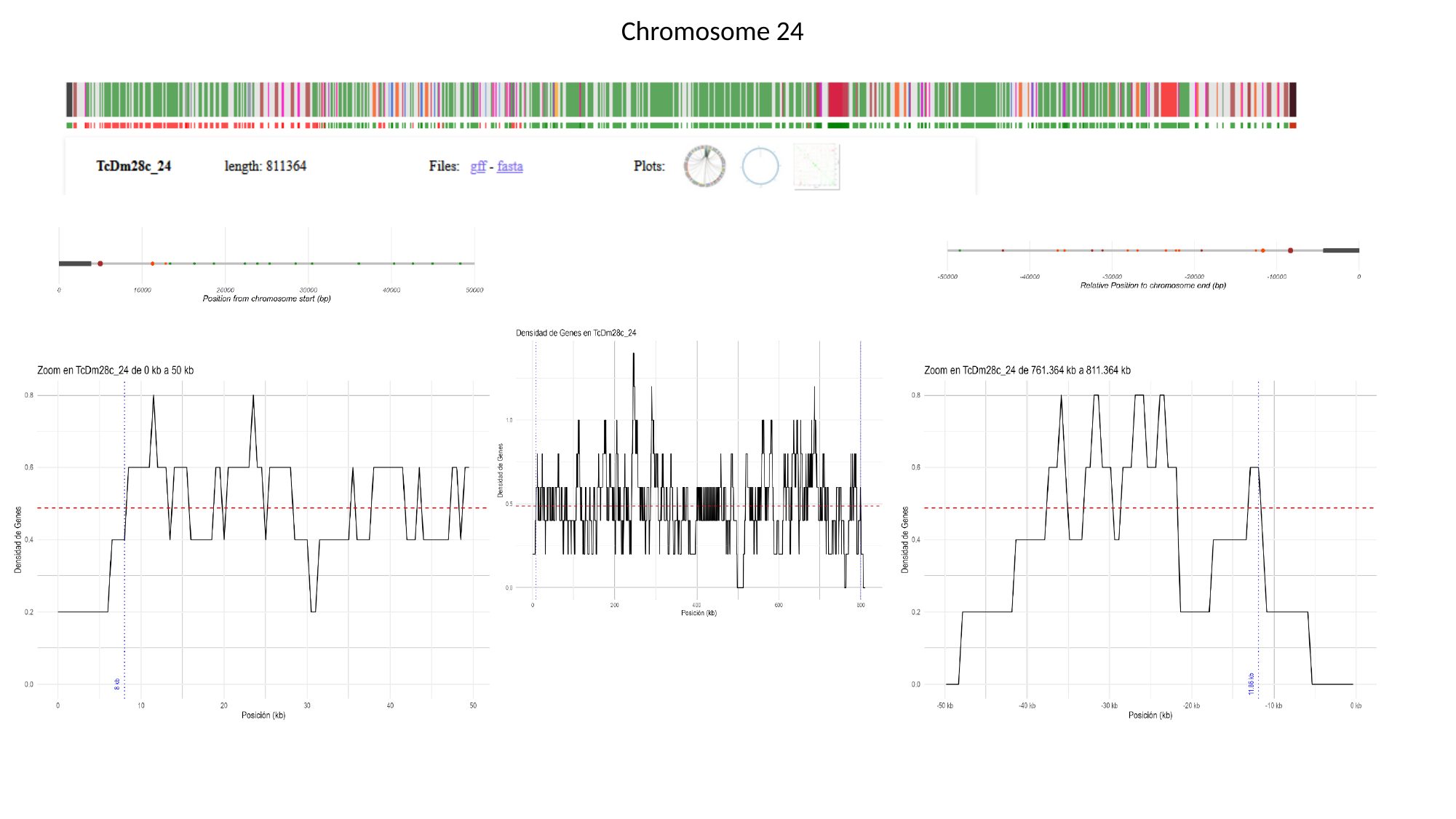

Chromosome 24

### Slide 26
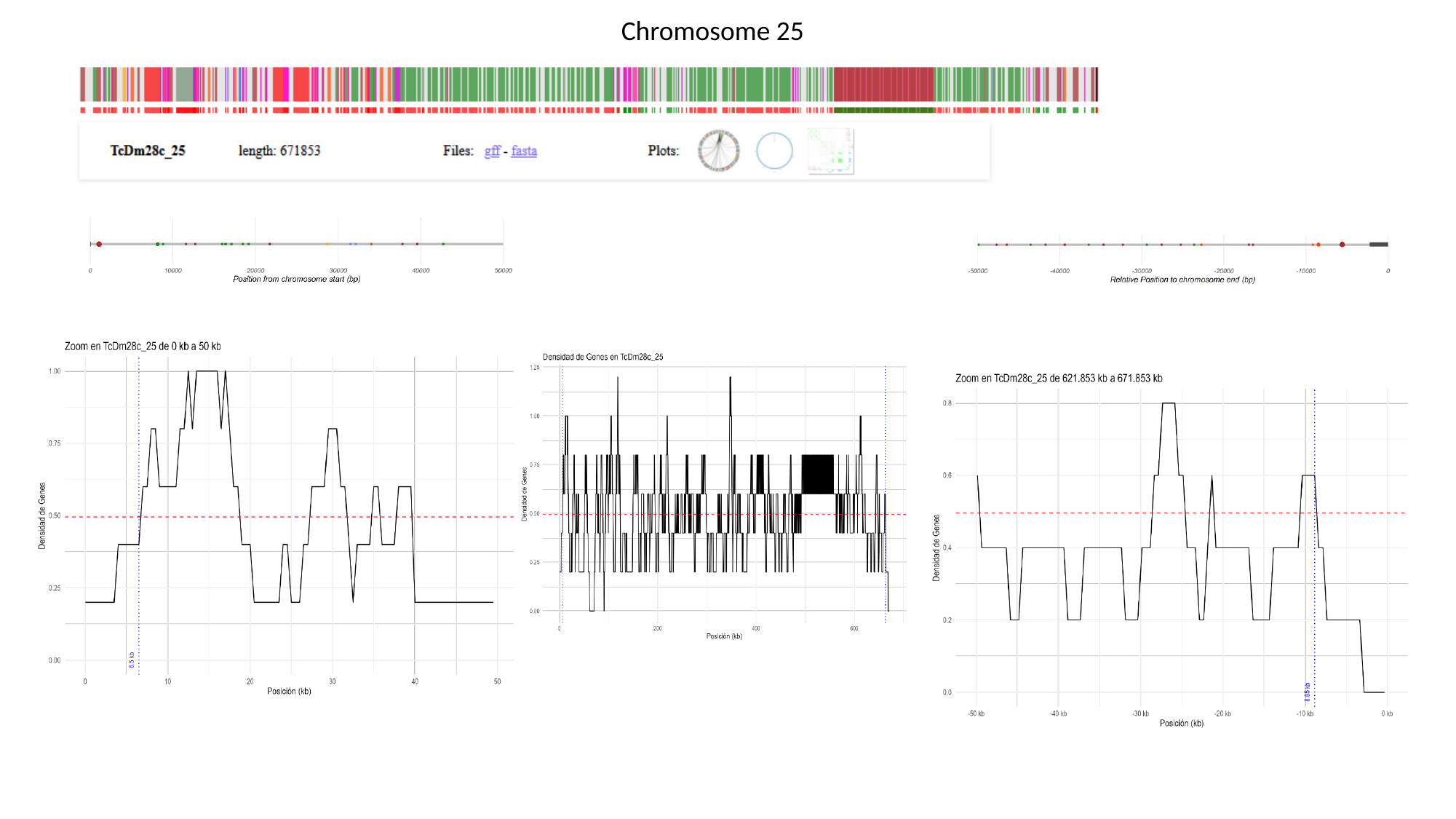

Chromosome 25

### Slide 27
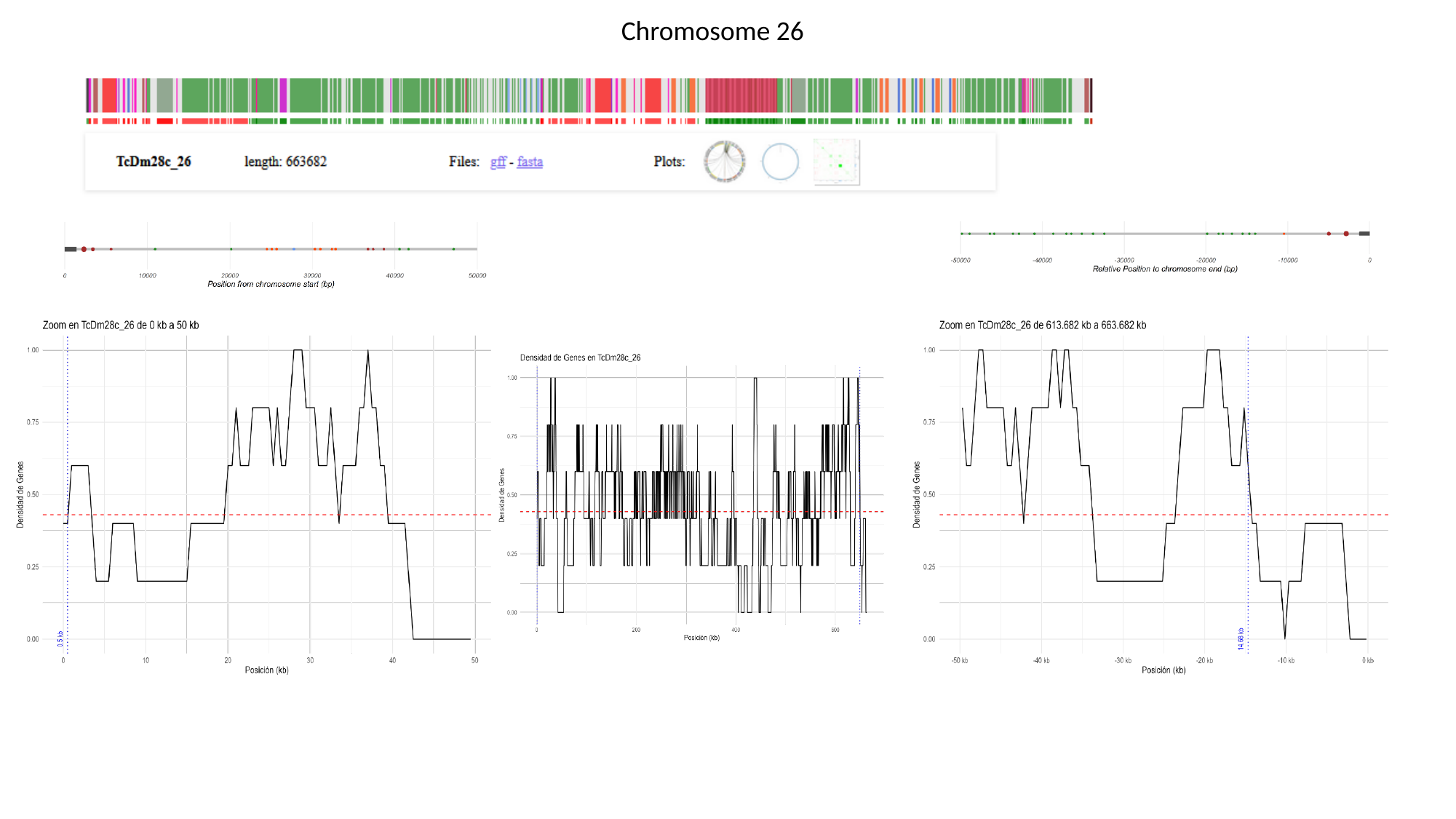

Chromosome 26

### Slide 28
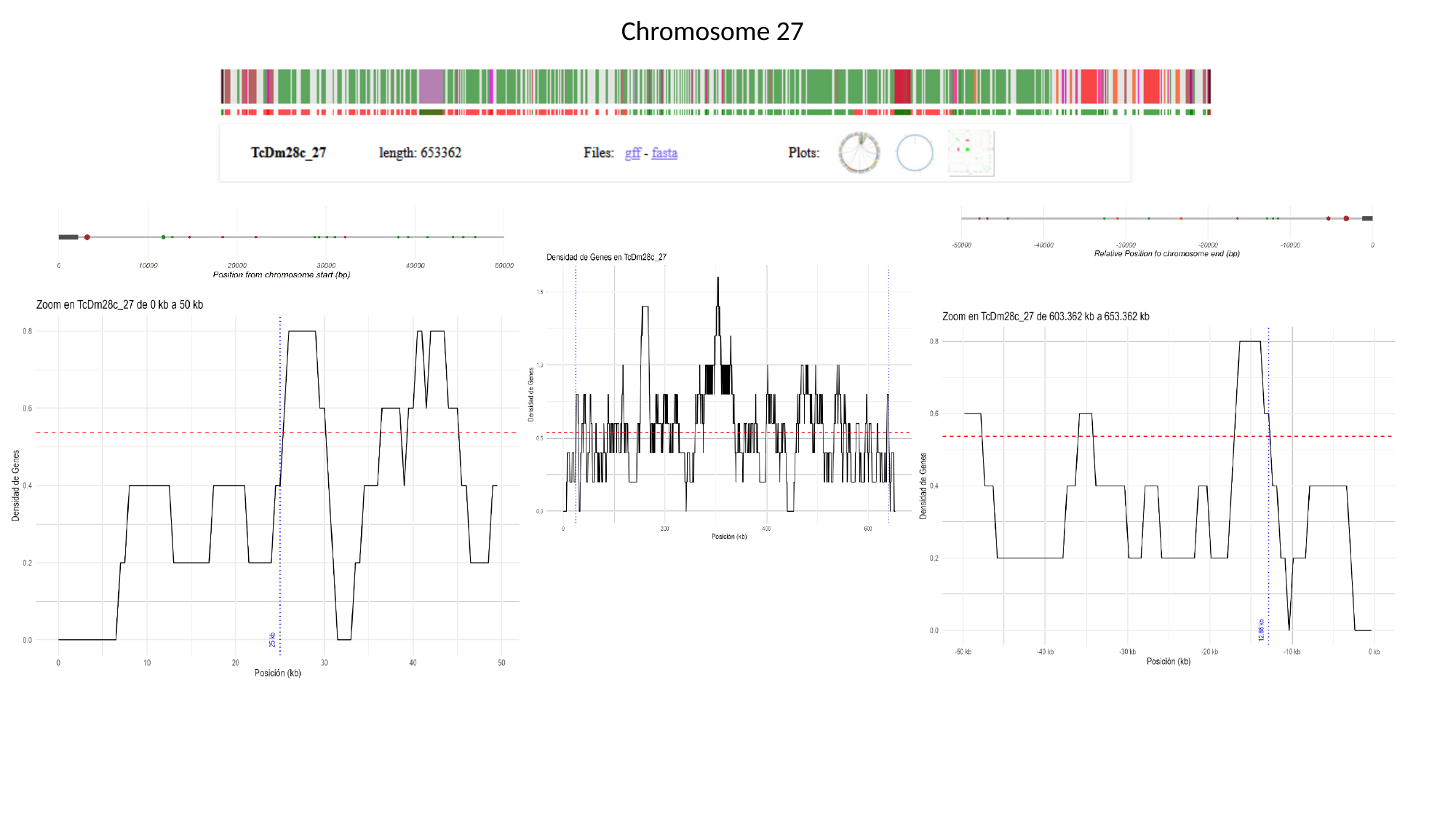

Chromosome 27

### Slide 29
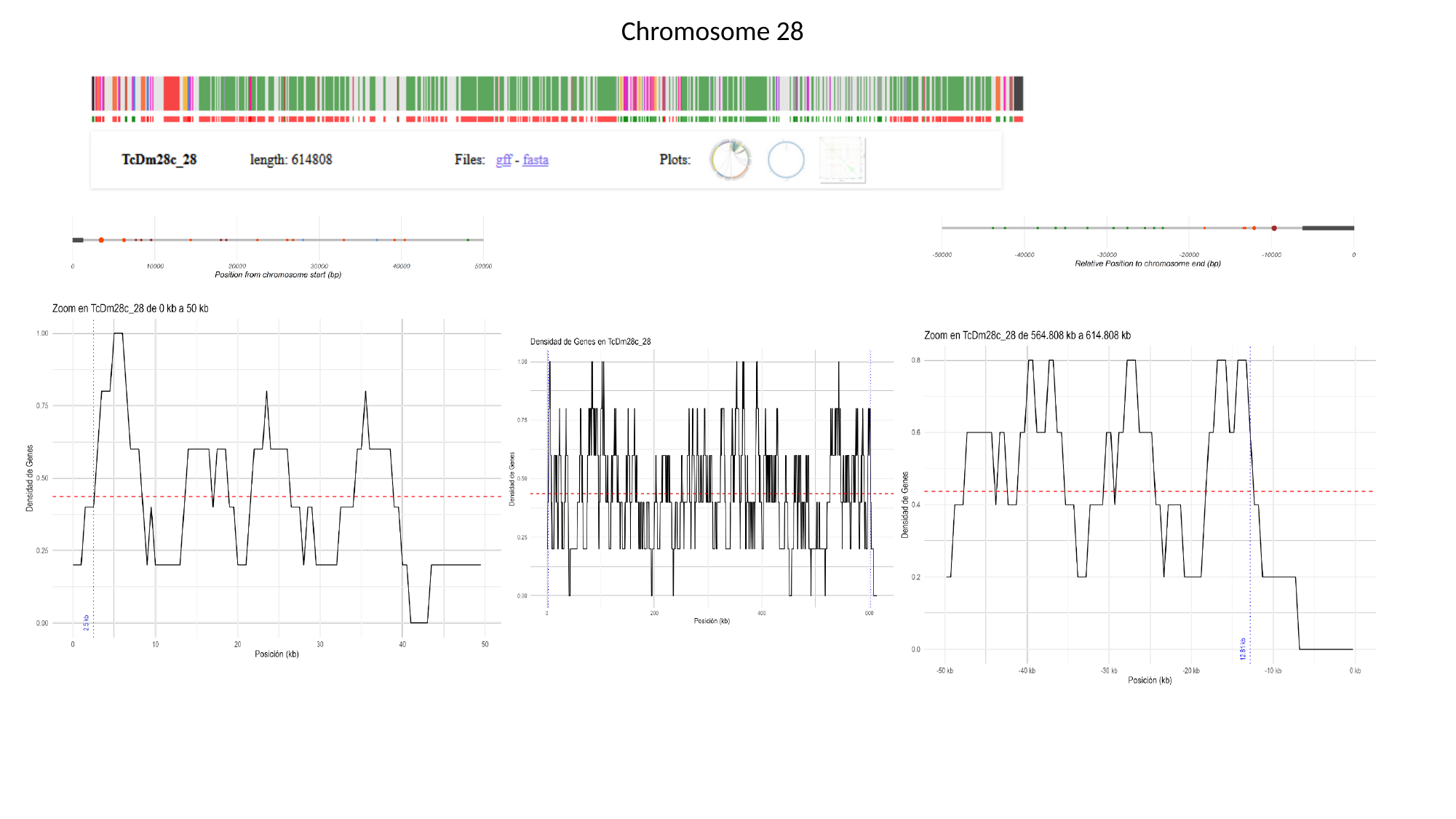

Chromosome 28

### Slide 30
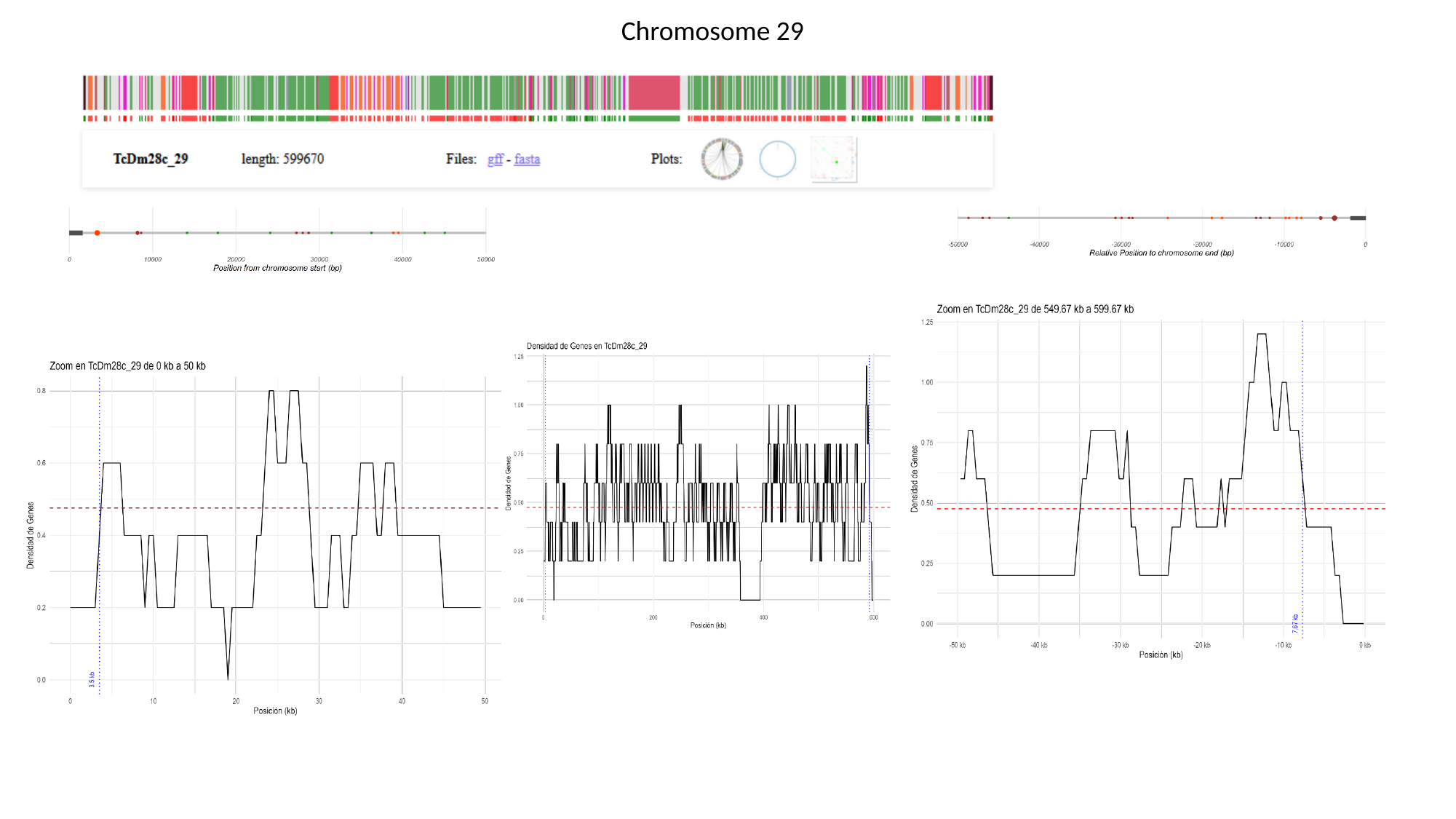

Chromosome 29

### Slide 31
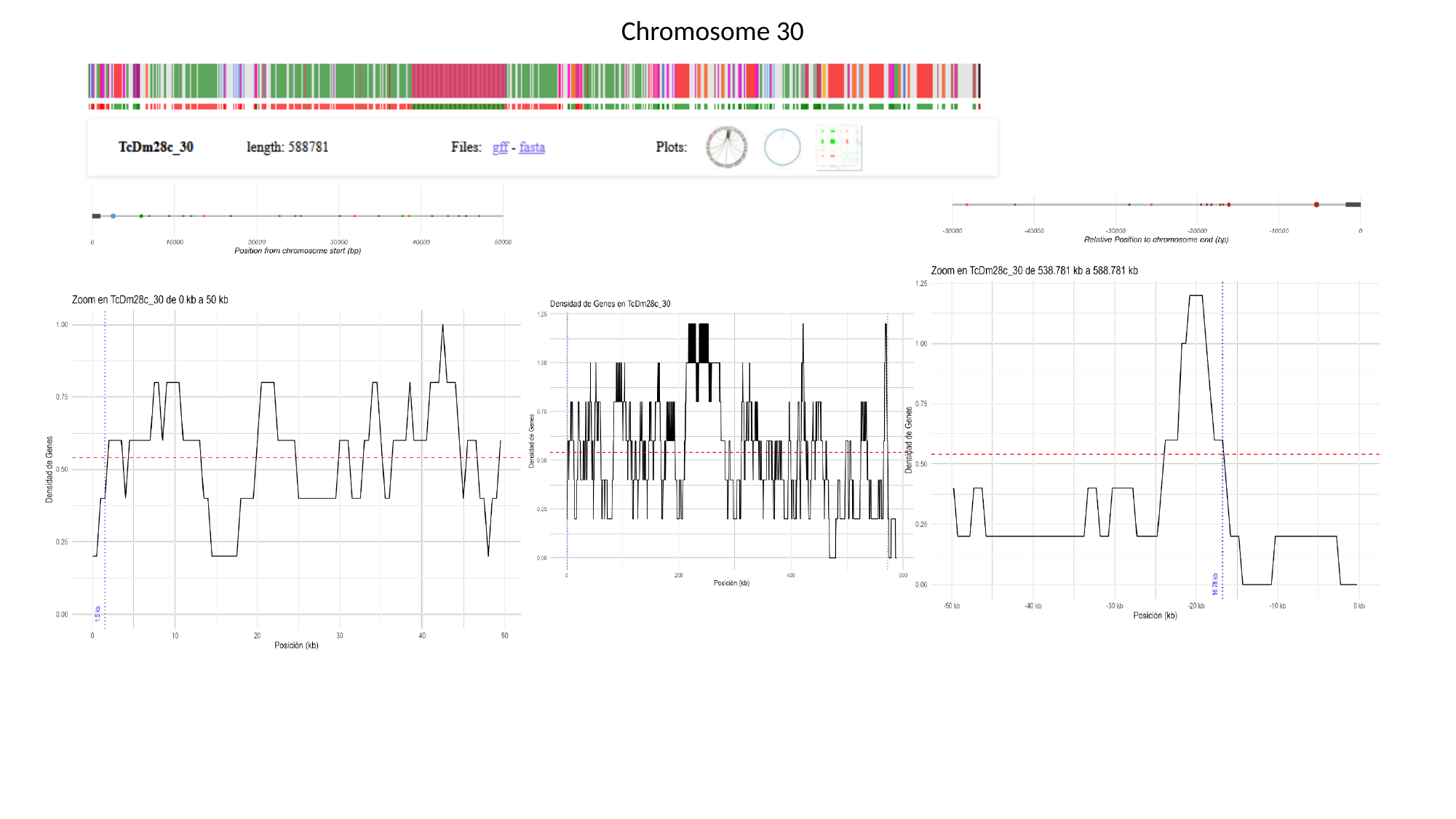

Chromosome 30

### Slide 32
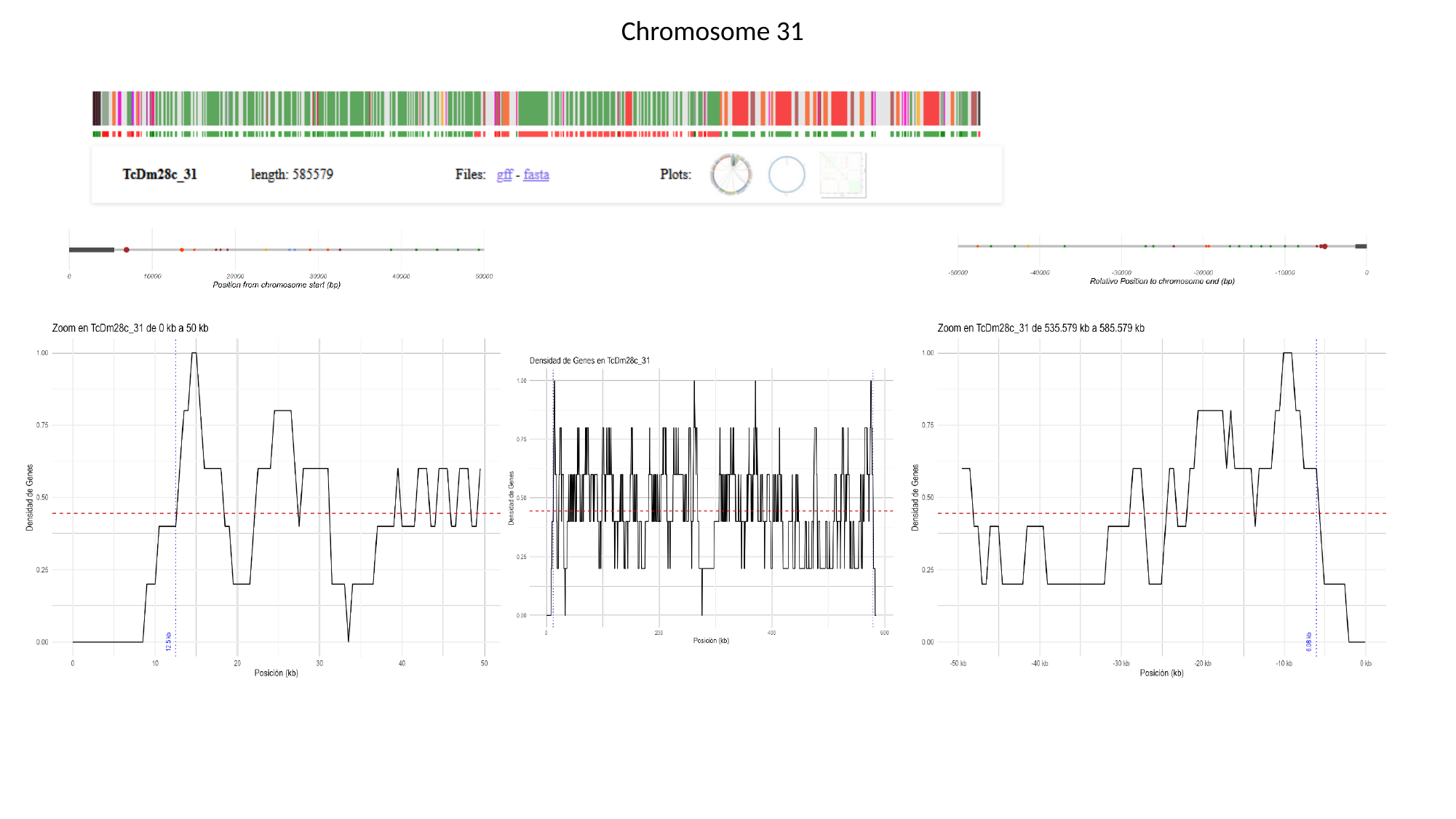

Chromosome 31

### Slide 33
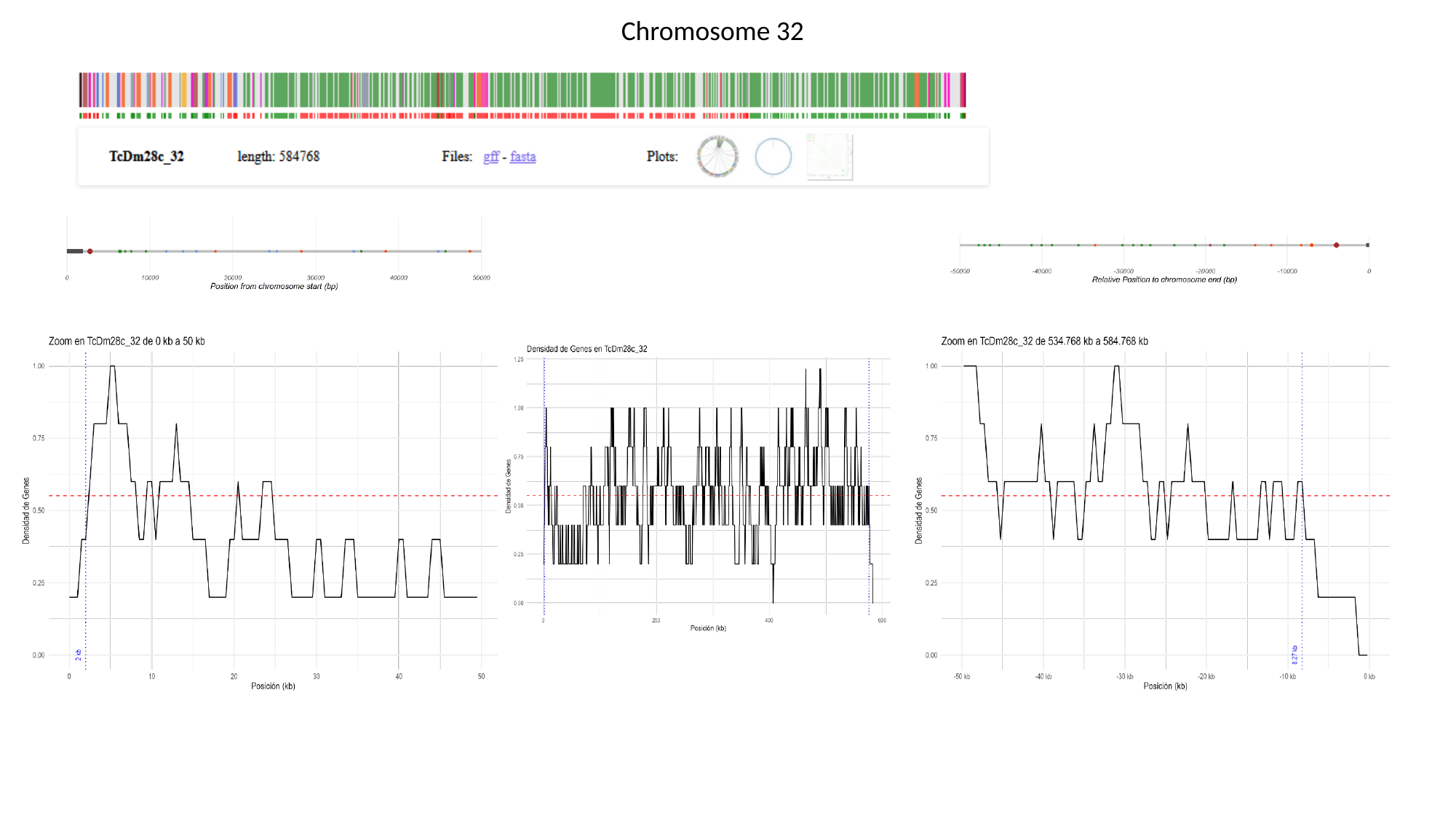

Chromosome 32
